# In-depth study of tomato and weed viromes reveals undiscovered plant virus diversity in an agroecosystem

**DOI:** 10.1101/2022.06.30.498278

**Authors:** Mark Paul Selda Rivarez, Anja Pecman, Katarina Bačnik, Olivera Maksimović Carvalho Ferreira, Ana Vučurović, Gabrijel Seljak, Nataša Mehle, Ion Gutiérrez-Aguirre, Maja Ravnikar, Denis Kutnjak

## Abstract

**Background:** In agroecosystems, viruses are well known to influence crop health and a few cause phytosanitary and economic problems, but their diversity in non-crop plants and role outside the disease perspective is less known. An extensive virome exploration that includes both crop and diverse weed plants is therefore needed to better understand roles of viruses in agroecosystems. Such unbiased exploration is possible through viromics, which could generate biological and ecological insights from immense high-throughput sequencing (HTS) data.

**Results:** Here, we implemented HTS-based viromics to explore viral diversity in tomatoes and weeds in farming areas at a nation-wide scale. We detected 125 viruses, including 79 novel species, wherein 65 were found exclusively in weeds. This spanned 21 higher-level plant virus taxa dominated by *Potyviridae*, *Rhabdoviridae*, and *Tombusviridae,* and four non-plant virus families. We detected viruses of non-plant hosts and viroid-like sequences, and demonstrated infectivity of a novel tobamovirus in plants of Solanaceae family. Diversities of predominant tomato viruses were variable, in some cases, comparable to that of global isolates of same species. We phylogenetically classified novel viruses, and showed links between a subgroup of phylogenetically-related rhabdoviruses to their taxonomically-related host plants. Ten classified viruses detected in tomatoes were also detected in weeds, which might indicate possible role of weeds as their reservoirs, and that these viruses could be exchanged between the two compartments.

**Conclusions:** We showed that even in relatively well studied agroecosystems, such as tomato farms, a large part of very diverse plant viromes can still be unknown and is mostly present in understudied non-crop plants. The overlapping presence of viruses in tomatoes and weeds implicate possible presence of virus reservoir and possible exchange between the weed and crop compartments, which may influence weed management decisions. The observed variability and widespread presence of predominant tomato viruses and the infectivity of a novel tobamovirus in solanaceous plants, provided foundation for further investigation of virus disease dynamics and their effect on tomato health. The extensive insights we generated from such in-depth agroecosystem virome exploration will be valuable in anticipating possible emergences of plant virus diseases, and would serve as baseline for further post-discovery characterization studies.

## Background

The awareness on the importance of virus diseases, especially amid an on-going COVID-19 pandemic [1], increased research interest on the exploration of virus diversity across ecosystems, assisted by high-throughput sequencing (HTS) [2–4], and by exploring global nucleotide databases [5–7]. In agroecosystems, viruses are ubiquitous microbes associated with eukaryotic hosts including crop and weed plants, fungi, oomycetes, arthropods, and nematodes, as well as prokaryotes such as bacteria [8, 9]. Thus, viruses could influence dynamics of plant populations and individual phytobiomes, directly, or through modulation of other ecological or environmental factors [10]. Due to the parasitic nature, high transmissibility and adaptability of plant pathogenic viruses [11], it was estimated that they account for half of emerging diseases in plants [12], and losses equivalent to around a quarter of expected crop yield [13]. Tomato (*Solanum lycopersicum* L.), which has the highest volume of vegetable production globally [14], is associated with more than 300 viruses including several that are frequently associated with disease symptoms and yield losses [15]. In recent years plant virologists have witnessed a spread of emerging tomato viruses, such as tomato brown rugose fruit virus (ToBRFV) [16] and tomato leaf curl New Delhi virus (ToLCNDV) [17]. To study such a high diversity of viruses, HTS has become the tool of choice. HTS-based viromics, coupled with bioinformatics tools, enable inference of biological, evolutionary, and ecological insights [18], which impact also fields of virus diagnostics and epidemiology [19].

With HTS, plant virology has greatly shifted from traditional focus on disease-associated relationships [18], to a scalable and unbiased ecosystems-level approach [3]. The study of plant virus pathogenesis and emergence with an ecological approach has also integrated the modulation of various environmental factors. Specifically, studies, which have been collectively reviewed in recent literature [20–22], demonstrated the importance of unmanaged weed plants surrounding cultivated areas in the distribution, ecology and emergence of viruses [23, 24]. In tomato, HTS has significantly advanced virus discovery, diversity, ecological and epidemiological studies [15]. Several HTS-based studies in the recent decade have contributed to the current known set of tomato viruses. A survey of tomatoes showing virus disease-like symptoms in China uncovered the presence of 21 known and one novel viruses [25]. Similar studies reported known and novel viruses associated with individual tomato plants or a collection of tomato samples using HTS [26–28]. Findings from these studies significantly contributed to the characterization of the global tomato virome and presented several insights that an HTS-based viromics survey can provide regarding the diversity, ecology and evolution of tomato viruses. In a survey of viruses in tomato and in surrounding *Solanum nigrum* from different locations in France possible exchanges of viruses were observed between wild and cultivated plant species [23]. This suggests that more extensive studies exploring the viromes of the vast diversity of weeds surrounding crop farming areas would be needed to better understand diversity and dynamics of plant viromes in an agroecosystem.

Studies that uncovered plant viromes at the ecosystem scale have focused mainly on discovery, and its epidemiological or ecological implications [23,25,29]. Knowing which species of viruses are present in an agroecosystem and to what extent they influence the host fitness landscape will greatly aid prediction of emergence [30]. To better understand plant virus diversity, ecology and possible emergence of plant virus diseases in an agroecosystem, we selected tomato as a model crop, and investigated a very diverse set of mostly broadleaf dicotyledonous volunteer plants, including weeds (designated simply ‘weeds’ hereafter) surrounding selected tomato production sites in Slovenia. We examined their viromes using an HTS-based approach, followed by virus characterization, to answer the following questions: (1) How diverse and prevalent are plant viruses and other virus-/viroid-like agents in tomatoes compared to that of surrounding weeds? (2) Are there overlaps of viruses detected in tomatoes and weeds? (3) Can we identify potential weed reservoirs of known crop viruses, and do weeds harbor an under-sampled virus diversity? (4) Can we identify some viruses in weeds, or those shared between tomatoes and weeds, which could potentially emerge in tomato? (5) What are the phylogenetic relationships of known and novel viruses to known taxa?

This study represented the largest simultaneous survey of viromes of diverse weed species and a crop (tomato) within a cropping system. Our approach can be used to gain insights on plant virus’ diversity and dynamics in the wild-cultivated agroecological interface, a known zone for virus spillovers [11,31,32]. The virome dataset could also aid in future tomato virus disease monitoring, thus possibly contributing to the prevention of future virus epidemics in tomato.

## Methods

### Sample collection and processing

Tomato and weed plants, within or in the immediate surroundings of tomato farming fields or greenhouses were collected in Slovenia during summer of 2019 and 2020. Fourteen farms at six different localities were visited, spanning central (Dol pri Ljubljani, 1 farm), west (Miren-Kostanjevica, 2 farms; Nova Gorica, 1 farm), southwest (Koper, 4 farms; Piran, 3 farms), and southeast (Novo Mesto, 3 farms) regions of Slovenia. A total of 436 samples were collected, full details of which are in Supplementary Table 1. Asymptomatic tomatoes were randomly sampled within an area, while symptomatic tomatoes and weeds were selectively sampled based on appearance of virus disease-like symptoms most commonly shown as foliar discoloration (*e.g.,* chlorosis, mosaic, yellowing), leaf deformation (*e.g.,* folding, curling), fruit deformation (*e.g.,* mottling, marbling), and systemic symptoms (*e.g.,* general stunting, rosetting), among others. Selected photographs of collected plants, with a notion of associated detected viruses, are shown in Supplementary Fig. 3 and Supplementary Table 9. Most tomato samples collected were in the early fruiting or fruit harvesting stage, while the most weed plants were sampled at the mature or flowering stage. To aliquot tissues for RNA extraction, leaves or fruits were obtained equally from different parts of the plant to account for possible uneven distribution of viruses within the plants. Leaf and fruit tissues were separately aliquoted in cases where these were collected from a single plant. Collected tissues were stored at -80°C.

### RNA extraction and sequencing

Total RNAs were extracted from individual plant samples using RNeasy Plant Mini Kit (Qiagen, USA). RNA quality and quantity were checked using QuBit fluorometer (Thermo Fisher Scientific, USA), and NanoDrop spectrophotometer (Thermo Fisher Scientific, USA). RNAs were then pooled equimolarly into composite samples based on plant type (*i.e.*, tomato or weed), health status (*i.e.*, symptomatic or asymptomatic), and the sampling location (Supplementary Table 2). A total of 67 composite samples were cleaned, concentrated and treated with DNase (RNA Clean and Concentrator, Zymo Research, USA). Alien controls from total RNAs extracted from *Phaseolus vulgaris* infected with Phaseolus vulgaris alphaendornavirus 1, Phaseolus vulgaris alphaendornavirus 2 and Phaseolus vulgaris alphaendornavirus 3 were included into each sequencing run. The 2019 and 2020 sample set were separately sent to Macrogen, Inc. (South Korea), for library preparation and high-throughput sequencing. Sequencing libraries, including depletion of ribosomal RNA using Ribo-Zero rRNA Removal Kit (Plant) (Illumina, USA), were prepared suitable for 150 bp paired-end sequencing using TruSeq RNA Library Prep Kit (Illumina, USA) and sequenced using Illumina HiSeq 2500 platform.

Total RNAs from inoculated plants (discussed below) were extracted as mentioned above and depleted of ribosomal RNA using RiboMinus^TM^ Plant Kit (Invitrogen, USA), then ligated with poly-A sequences using E. coli Poly(A) polymerase (NEB #M0276, UK). Library preparation was done using the PCR-cDNA Barcoding kit (SQK-PCB109, version 10Oct2019, Oxford Nanopore Technologies, UK), prior to sequencing using Oxford Nanopore Technologies MinION platform (Oxford Nanopore Technologies, UK) and base-calling following a previously described workflow [33].

### Sequence quality screening, trimming and virus genome assembly

Raw reads were trimmed, screened for quality and analyzed following a previously described pipeline for plant virus detection using HTS [34]. Contigs were primarily assembled from the filtered reads using CLC Genomics Workbench (GWB) v. 20 [34] (Qiagen, USA). Within the used pipeline, virus and virus-like reads and contigs were initially identified by mapping trimmed reads/contigs to virus RefSeq database (version from Jul. 2020) [35] and viral domains searches in contigs against pFam database v. 33. Candidate viral contigs were later confirmed by homology search using BLASTn against the NCBI nucleotide (nt) and BLASTx [36] against NCBI non-redundant protein (nr) databases from Dec. 2020. Assembly in metaSPAdes v. 3.14 [37] was also implemented to recover longer contigs in some cases, where these were not assembled in CLC-GWB. Consensus genome sequences of detected viruses were reconstructed using de-novo assembled contigs and/or iterative reads mapping to the most similar reference sequences obtained from NCBI GenBank. Since datasets were derived from pooled samples, consensus viral genomes were reconstructed only for the viruses with observed low to moderate population diversity (determined after manual inspection of the mapping files), indicative of infection with a single viral lineage. As a final checkup, mapping of reads of corresponding datasets to reconstructed consensus virus genomes was implemented (with 95% identity and read length fraction). Overall sequencing results and statistics, and information on sequencing read archive (SRA) metadata are given in Supplementary Table 3. The internal controls were used to check the prevalence of sequencing cross-talks. A threshold of <0.00001% of total reads in the library was set, based on the general virome composition of the internal controls, to classify sequences as either a possible contamination, crosstalk, or low titer virus infection.

For the sequences obtained from inoculated plants (discussed below) using MinION platform, quality screening was done following a customized workflow [33], that uses the programs from the NanoPack program [38] for quality screening and visualization. Reads were assembled using Racon [39]. The mapping program Minimap2 [40] run in Geneious Prime software v. 2022.1.1 (Dotmatics, USA), was used for mapping reads to viral RefSeq (version from May 2022) and consensus genomes of viruses detected in this study.

### Virus genome annotation and classification

Genomes of detected viruses were annotated using the ORF prediction tool in CLC-GWB v. 20, and checked based on known genome organization and open reading frame composition reported in the ICTV website [41], and in peer-reviewed publications. InterProScan was used to identify known and unknown protein domains [42], and in the case of potyviruses, homology alignment in BLASTp (E-val<10^-4^) with well-characterized known species was implemented to detect gene products and their start and cleavage sites [43]. Viruses were taxonomically classified based on the percent pairwise identity obtained in SDT v1. 2[44], and other genomic criteria imposed by the ICTV [41], as of Oct. 31, 2021. Multiple sequence alignments were made for each genus or family based on the recommended gene or genome segment by the ICTV. Unaligned ends were trimmed, and trimmed alignments were used for pairwise identity comparisons. Details of the genes or genome segments used, and the results of the pairwise identity comparisons are indicated in Supplementary Tables 23-42. Classifications were then confirmed by analyses of phylogenetic relationships with known virus taxa (described below).

### Genome assembly and screening for putative viroids

Viroid-like circular RNAs were assembled using the SLS-PFOR2 program [45]. BLASTn searches [36] against the NCBI nt database and BLASTx searches against NCBI nr database, from Dec. 2020, were implemented to filter out sequences from known organisms, and sequences that code for proteins (E-val<10^-4^). Filtered contigs were re-examined by remapping (95% identity and read length fraction) virtually-diced reads (generated as a part of SLS-PFOR2 pipeline) and trimmed reads to the assembled circular RNAs. Contigs with average mapping depth below 10 were manually discarded. Contigs were again filtered based on the presence of two or more rotationally identical contigs of the same length (*i.e.*, indicating the presence of (+) and (-) strands). After selecting for rotational identical contigs, presence of viroid-like secondary structure motifs (*e.g.,* avsunviroid hammerhead ribozyme and rod-like pospiviroid structures) were predicted using forna [46]. Low structural Gibbs free energy and visual inspections such as high degree of base paring, or degree of branching were the criteria used to preliminarily select for putative novel viroids.

### RT-PCR assays and Sanger sequencing

PCR primers were designed using Primer3Plus [47], and RT-PCR assays were designed to identify individual plant host(s) of selected viruses detected using HTS of pooled samples (Supplementary Table 7). RT-PCRs were performed using OneStep RT-PCR kit (Qiagen, USA) with thermocycling program conditions given in Supplementary Table 8. To confirm genome circularity of viroid-like circular RNAs, abutting primers for inverse RT-PCR were manually designed as previously described [45] (Supplementary Fig. 2). Amplicons were visualized in 1% agarose gel, stained with ethidium bromide. When PCR amplicons need to be sequenced, to confirm sequence identity or circularity in the viroid-like sequences, they were purified using QIAquick PCR purification kit (Qiagen, USA), and sent at Eurofins Genomics, Germany. Amplicon sequences were visualized and trimmed in CLC-GWB v. 20. Final sequences were remapped (95% identity and read length fraction) to the target virus genome to confirm its identity, or to putative viroid genome to confirm its circularity (Supplementary Fig. 2). Photos of identified plant hosts are in Supplementary Fig. 3.

### Phylogenetic analyses

Phylogenetic analyses were performed to investigate taxonomic positions of detected newly discovered and known viruses from different taxonomic groups (Supplementary Table 10). Conserved amino acid sequences of viruses (mainly conserved parts of RNA-dependent RNA polymerase (RdRp) (methyl transferase, helicase or CP were used in some cases) for RNA viruses, replicase (C1) protein for geminivirus and reverse transcriptase (RT) for caulimovirus) detected in this study were aligned with corresponding viral RefSeq amino acid sequences of viruses within a recognized virus family, in addition to selected BLASTp hits (E-val<10^-4^). For multiple sequence alignment, ClustalW [48] was used as implemented in CLC-GWB v. 20. Non-aligned sequences at both ends were trimmed after alignment. Realignment of trimmed sequences was done using MAFFT [49] and further processed with trimAI [50].

In the case of tobamoviruses, a nucleotide-based phylogenetic analyses were done on full genomes. Genomes of different isolates gathered from GenBank and viral RefSeq database, were aligned using MUSCLE [51], and checked for possible recombinants in RDP v. 4 [52]. All recombination events with *p*-val<10^-4^ reported by at least four methods were considered significant. Phylogenetic analyses were performed on alignments free of recombinants.

For both, protein and nucleotide alignments, maximum likelihood phylogenetic analyses were inferred in MEGA X [53], after the selection of the most suitable substitution models. Details on the sequence sets for each analysis, and the parameters used in the analyses are in Supplementary Table 10. The phylogenetic trees were visualized and edited in iToL v. 6.4 [54]. To investigate possible structuring of virus species in phylogenetic trees by associated host plants, plant host cladograms were generated in phyloT v. 2 (https://phylot.biobyte.de/) [54], based on NCBI Taxonomy. Host associations were based on information on the specific accession in NCBI GenBank of the virus isolate of tobamoviruses in Fig. 7, and all isolates of species for rhabdoviruses in Fig. 5. Connections were manually inferred between viral and plant phylogram and cladogram and visually inspected.

### Genome-wide molecular diversity analyses

Tomato viruses which have at least three isolates detected with full genomes reconstructed, including their isolates from weed samples, were included in the calculation of percent pairwise identities and polymorphisms. Full genome nucleotide sequences of selected viruses were aligned using MUSCLE [51] in MEGA X [53], and unaligned ends were trimmed thereafter. Genome segments of multi-segmented virus genomes were concatenated after trimming. Percent pairwise identities and number of polymorphic sites were calculated using SDT v. 1.2 [44]. For two newly discovered viruses, the aligned sequences were used to calculate genome-wide polymorphisms and nucleotide diversities (π) in DnaSP v. 6 [55] in sliding windows of 100 bases and step size of 5.

### Infectivity tests and transmission electron microscopy (TEM)

Further characterization studies were performed to test the infectivity and visualize the virions of newly discovered Plantago tobamovirus 1 (PTV1). Five plants each of *Solanum lycopersicum, Nicotiana benthamiana,* and *Nicotiana clevelandii,* were used in the inoculation experiments, with additional two plants each as mock inoculated controls. Plants were inoculated at the four-leaf stage. The source of inoculum for PTV1 was the original *Plantago major* infected sample from the field. Mechanical inoculation of infected plant sap, diluted 1:10 in phosphate buffer, was done on two leaves (second and third youngest), dusted with carborundum, as previously described [56]. The plants were kept in a controlled environment at 20-24°C, with 16 h daylight and 8 h darkness. Inoculated leaves were tested in pools for the presence of the target virus on 7^th^ and 14^th^ day post inoculation (dpi) in RT-PCR assays. Systemic leaves of inoculated plants were tested for the presence of the PTV1 using RT-PCR assays, wherein pooled newly grown (uninoculated) leaves at 14, 21, 28 and 35 dpi were used. TEM was done to confirm the presence of the virus particles in inoculated *N. clevelandii* plants. Tissue homogenates were applied to Formvar-coated, carbon-stabilized copper grids, and negatively stained with 1% uranyl acetate (SPI Supplies, PA, USA), before inspection in TEM. To associate the observed disease symptoms with the presence of PTV1, pool of five inoculated *N. clevelandii* plant showing virus disease-like symptoms and tested positive for PTV1 was collected at 21 dpi and analyzed using nanopore sequencing as described in section ’ RNA extraction and sequencing’ above.

## Results

### Overview of viruses detected in tomatoes and weeds and observed patterns of viral presence and diversity

Tomato (n=293), and weed (n=143, 59 species in 18 families) plant samples, were collected from six localities in Slovenia, wherein 14 farms were visited in 2019 and/or 2020 (Fig. 1a-b, for details see Supplementary Tables 1-3). Sequencing of pooled ribosomal (r)RNA-depleted total RNAs (*i.e.,* composite samples) from these samples, and subsequent bioinformatics analyses revealed 126 viruses (Fig. 1c), of which 37 are known and 55 are novel classified viruses that could be classified up to known species taxa (46 are exclusively from weeds). The remaining 33 viruses are either isolates of known viruses, which were not yet classified under established taxa in *Riboviria* (n=9), or unclassified putative novel *Riboviria,* satellite RNAs, and viroid-like circular RNAs (n=24, 19 are exclusively from weeds). Thus, the total of novel viruses discovered is 79, of which 65 were found exclusively in weed plants (Fig. 1h, for details see Supplementary Tables 4-6). Farms from the coastal region of SW Slovenia (Piran and Koper), where 51% of the samples were collected, have the highest count of virus species detected (*i.e.,* 57 viruses in Koper and 84 viruses in Piran) (Fig. 1d). The viruses were classified in 21 known plant virus taxa, and four other families that are known to be associated with other eukaryotic hosts (*Dicistroviridae, Iflaviridae, Lispiviridae,* and *Picornaviridae*) (Fig. 1e). Majority of the detected viruses were from 15 taxa of positive sense (+) single-stranded (ss) RNA viruses (n=60), followed by four families of negative sense (-) ssRNA viruses (n=17) (Fig. 1f). Ten viruses were found both in tomato and weed composite samples from several localities, eight of which are most likely plant-infecting. Some of the ten overlapping viruses are known to have wide host range, and are endemic in tomato (*i.e.,* tomato spotted wilt orthotospovirus (TSWV), cucumber mosaic virus (CMV), tomato mosaic virus (ToMV), and potato virus Y (PVY)) [15] (Fig. 1g). Viruses recently detected in tomato (*i.e.,* Solanum nigrum ilarvirus 1 (SnIV1), Ranunculus white mottle ophiovirus (RWMV), tomato matilda virus (TMaV)) [23,57–59], a known insect (Aphis glycines virus 1 (AGV1)) and a known fungal (Leveillula taurica associated rhabdo-like virus 1 (LtaRLV1)) virus, and a novel virus (plant associated tobamo-like virus 1 (PaToLV1)) were also detected in both sample types. Among the viruses detected both in tomatoes and weeds, there were several cases in which we could detect the same virus within the same location in both sample types: seven viruses (TSWV, CMV, ToMV, PaToLV1, LtaRLV1, PVY, and TMaV) with such pattern were observed.

**Figure 1.**
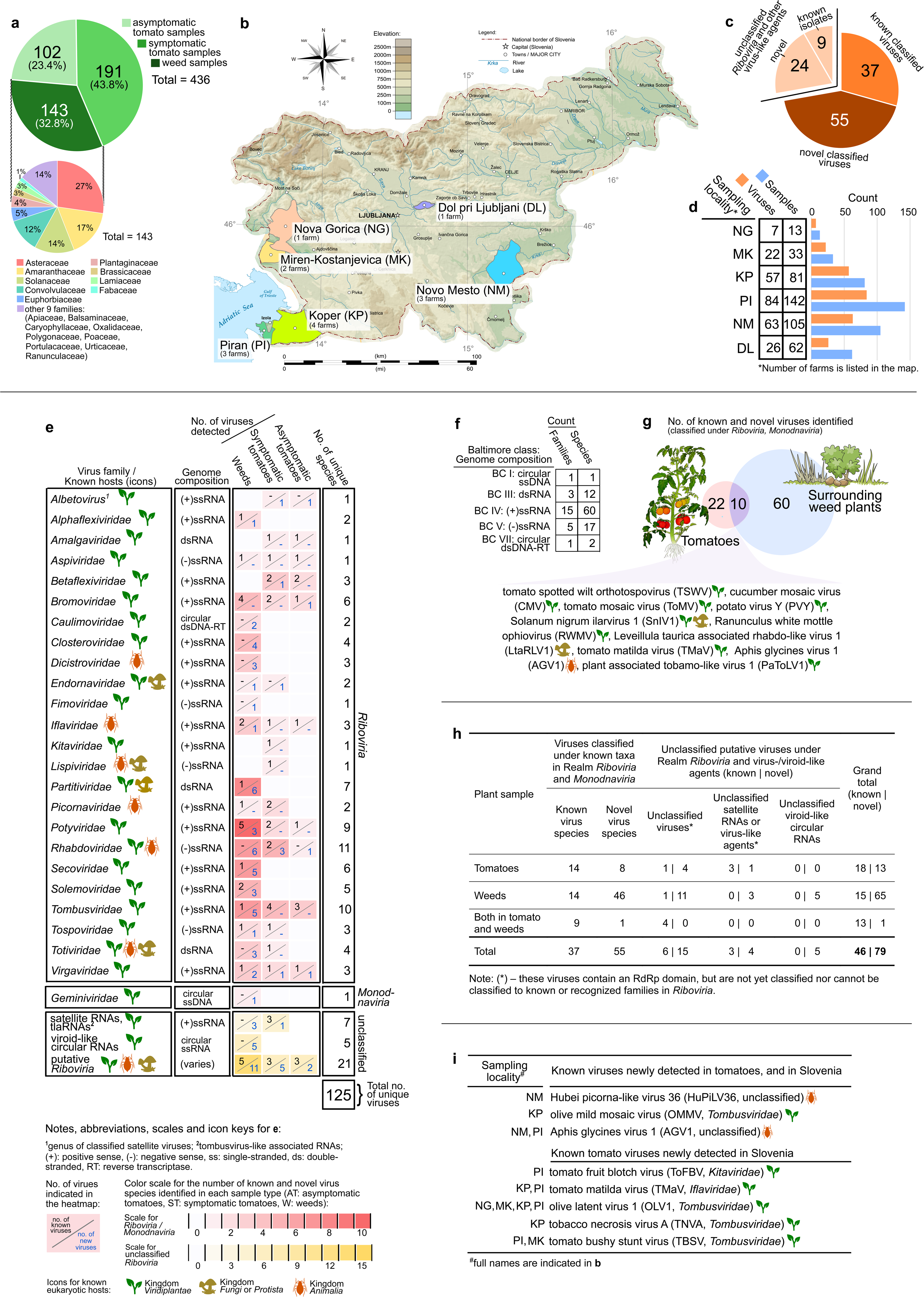
Diversity and distribution of samples and viruses detected in tomatoes and weeds surrounding tomato farms. **a** Pie chart showing proportion of sample count per sample type, and separately for weed samples, proportion of sample count per plant family. **b** Map showing the geographical locations where sampling was done. Farm codes are indicated for each locality. **c** Overall count of viruses, with (near) complete genomes, identified from this study, distributed as proportion of known, novel and unclassified *Riboviria* species. **d** Bar plot summarizing the number of samples collected, and viruses detected in each sampling locality. **e** Heatmap showing the diversity of identified viruses by genome composition, their distribution in each known virus taxa by sample type, and the known eukaryotic host (Kingdom) of each virus taxa. **f** Summary of count of identified viruses based on Baltimore class or genome composition. **g** Venn diagram showing the number of identified known and novel viruses found exclusively in tomato or weed composite samples, and the viruses found in both sample types. Credit: Images or icons used in the figure were from https://freesvg.com/, and are under Public Domain (CC0 license, https://creativecommons.org/licenses/publicdomain/). The map of Slovenia was derived from work of Andrei Nacu uploaded in Wikimedia Commons, the free media repository, under the Public Domain (CC0 license). The authors and publisher remain neutral with regard to jurisdictional claims in published maps.

Three known viruses were detected for the first time both in tomatoes and in Slovenia, and five others were detected for the first time in the country, (*i.e.*, were not reported before in local comprehensive records) [60] (Fig. 1i). Lastly, other known crop viruses were detected in composite weed samples, including two potyviruses (watermelon mosaic virus (WMV) and carrot thin leaf virus (CTLV)), ilarvirus (Prunus virus I (PrVI)), closterovirus (beet yellows virus), luteovirus (soybean dwarf virus), cucumovirus (peanut stunt virus), fabavirus (broad bean wilt virus 1), polerovirus (barley virus G), and potexvirus (white clover mosaic virus).

To get general insights into the association of detected viruses with observed disease symptoms, we have compared the number and overlap of detected viruses in sampled symptomatic and asymptomatic tomatoes. Out of the 45 viruses detected in tomato composite samples, only four (8.9%) were exclusively detected in asymptomatic tomatoes, and 23 (51.1%) exclusively in symptomatic tomatoes (Fig. 2a). Eighteen (40.0%) viruses were detected in both types of tomato samples, of which, six (*i.e.,* southern tomato virus (STV), CMV, PVY, olive latent virus 1 (OLV1), TMaV, and ToMV) were detected in at least six composite samples. A total of seven new viruses, and eight known but still unclassified arthropod, fungal and oomycete viruses were detected in tomatoes. Using RT-PCR assays (for details, see Methods and Supplementary Table 7), we confirmed the presence of a subset of mostly novel viruses in weeds and tomatoes, and identified key hosts that could be potential transmission hubs, or alternate hosts (Fig. 2b, for details see Supplementary Table 9). Some examples include: SnIV1 detected in tomatoes and in another weed host, *Physalis* sp., from a single farm, PaToLV1 detected in tomatoes and in *Convolvulus arvensis* also in a single farm, and TMaV detected in tomatoes and in three other weed species (*i.e., Chenopodium* sp., *Ranunculus repens,* and *Erigeron annuus*) that span five localities. RWMV was detected both in *Solanum nigrum,* and in four pools of tomato samples from three different localities. Twelve novel viruses (discussed in succeeding sections), including six rhabdoviruses and three tombusviruses, were detected in four Asteraceae species (*i.e., Picris echoides, Cichorium intybus, Taraxacum officinale,* and *Cirsium arvense*).

**Figure 2.**
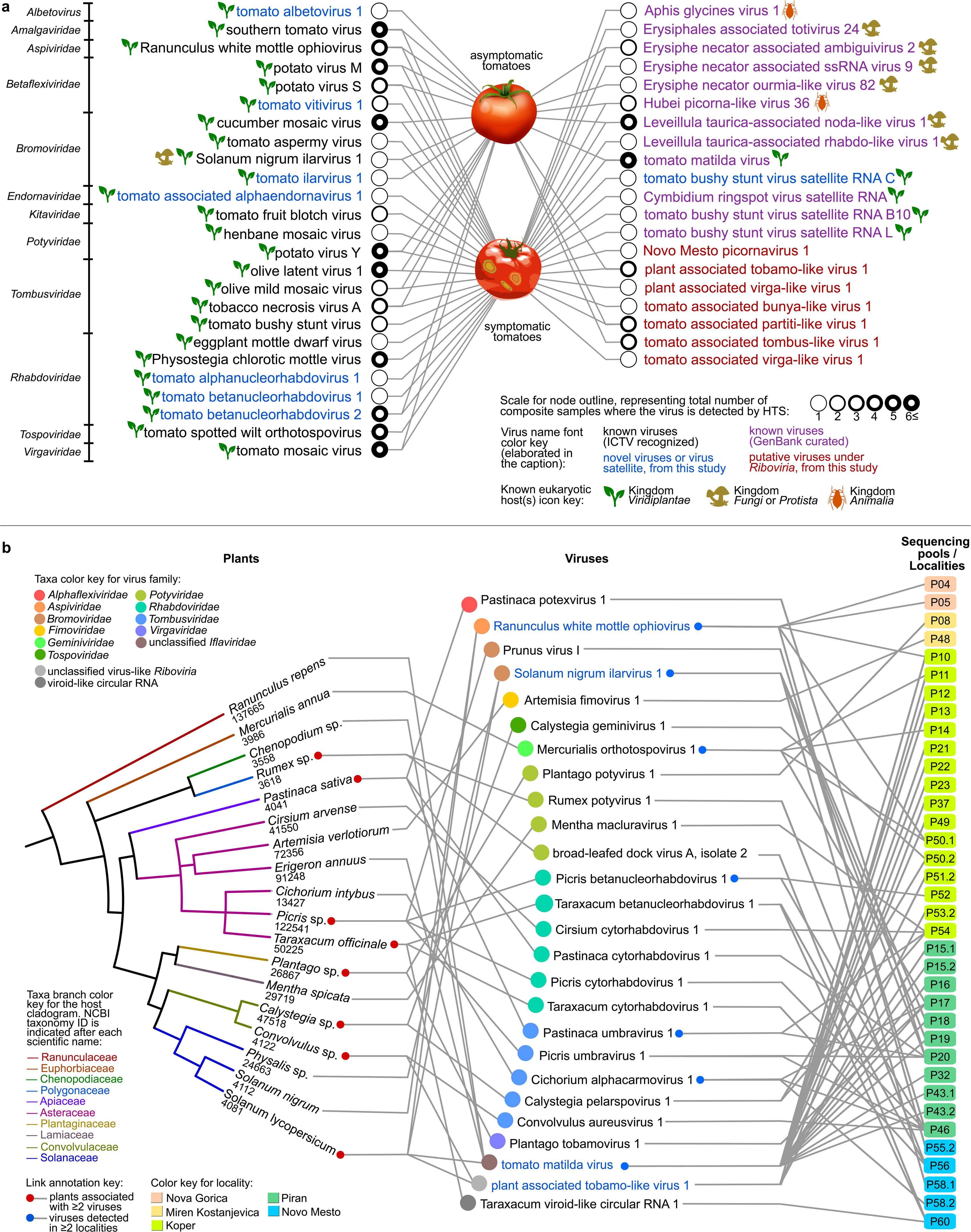
Connectivity and overlaps of viruses detected in different sample types. **a** Bipartite network showing detection of viruses in composite symptomatic and asymptomatic tomato samples from the present study. The node represents the viruses, in which outline thickness is weighted based on the number of composite samples where the virus was detected by HTS. The lines connect viruses with a sample type in which they were detected. Virus names are color coded according to classification status: known viruses are either officially recognized ICTV species (black font), or viruses that are deposited and classified in GenBank, but not ICTV-recognized (purple). Novel viruses from this study, which are classified up to the species level, are in blue font, while putative viruses without any official classification yet are in red font. **b** Plant-virus-sample pool tripartite network, showing the connectivity of the subset of viruses detected in weeds with associated hosts. Shown in the network is the subset of novel viruses detected in weeds by HTS, which were further associated with specific host species (NCBI taxonomy IDs are shown) using RT-PCR assays or deduction from HTS data (for tomato, which was not pooled with other species). Virus names in blue font are those detected in both tomato and weed samples by HTS and RT-PCR. Credit: Images and icons used in the figure were from www.freesvg.com, and are under Public Domain (CC0 license, https://creativecommons.org/licenses/publicdomain/).

To gain further insights on the population genomic diversity of the most prevalent tomato viruses, we examined populations of 12 viruses (viruses for which we were able to assemble at least three full genomes). STV, ToMV, TMaV and TSWV showed the narrowest range of pairwise nucleotide identities, lowest nucleotide diversity and number of polymorphic sites, while Physostegia chlorotic mottle virus (PhCMoV), RWMV, and potato virus S (PVS) have a moderate level of diversity (Fig 3a, Supplementary Tables 11-22). PVY, CMV, potato virus M (PVM), tomato betanucleorhabdovirus 2 (TBRV2), and tomato vitivirus 1 (TomV1) showed wide range of pairwise nucleotide identities and high number of polymorphic sites (Fig 3a, Supplementary Tables 11-22). Sliding window analysis of nucleotide diversity for two novel tomato viruses found in Slovenia (TBRV2 and TomV1), did not pinpoint specific regions of their genomes that might be more variable than others (Fig. 3b-c). Divergent lineages detected for both TBRV2 and TomV1, resulted in a high genome-wide variability depicted as fluctuations of nucleotide diversity along the genome (Fig. 3b-c).

**Figure 3.**
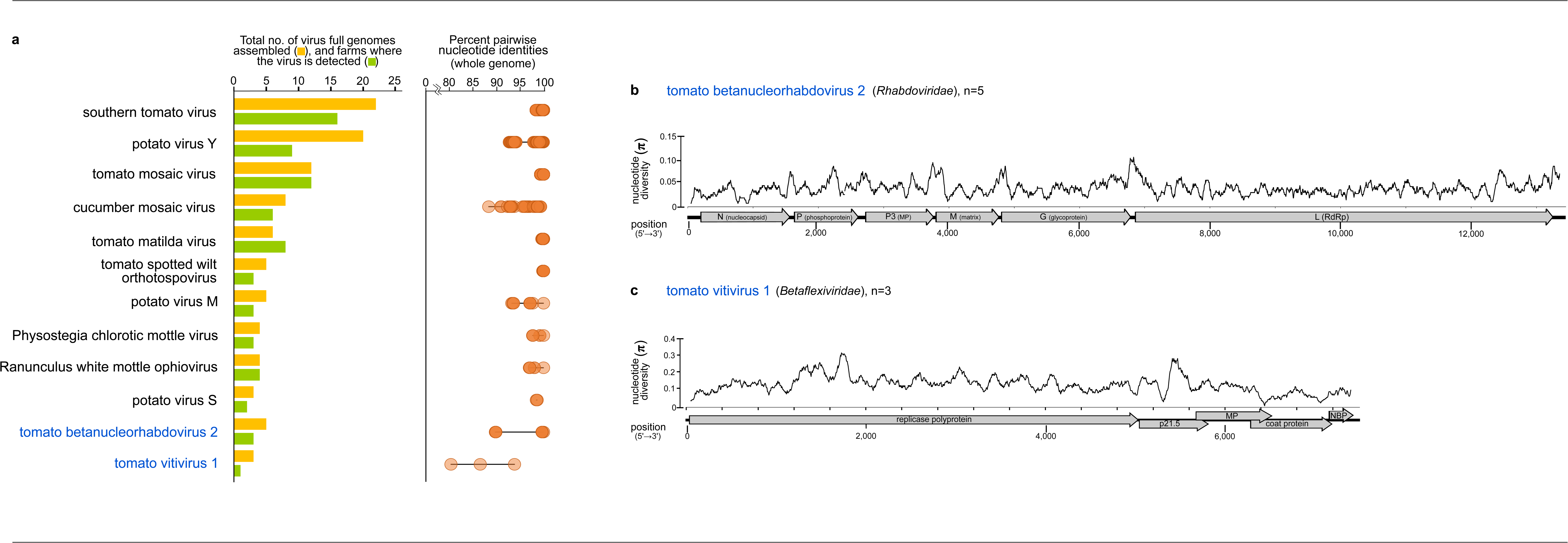
Molecular diversity and genome-wide variability of most frequently detected tomato viruses in this study. **a** Pairwise nucleotide identities of populations of tomato viruses from Slovenia. Isolates of tomato viruses from weed samples, were included in the calculation of pairwise identities. **b, c** Genome-wide nucleotide diversity of novel viruses in tomatoes from Slovenia with at least three isolates. All open reading frames and protein domain cleavage sites are shown for each virus genome.

### (-)ssRNA viruses in tomatoes and weeds: discovery of novel orthotospovirus, an ophiovirus, and a divergent bunya-like virus

Several (-)ssRNA viruses were detected in both, tomatoes and weeds. A novel orthotospovirus, Mercurialis orthotospovirus 1 (MerV1), was detected in *Mercurialis annua* (Fig. 4a) in four composite samples from three localities. Pairwise comparison of RNA-dependent RNA polymerase (RdRp) amino acid (aa) sequences showed 72.4-76.0% identity of MerV1 to related *Tospoviridae* species. MerV1 has genome segments similar to plant orthotospoviruses (Fig. 4b,e). Phylogenetic analyses based on conserved RdRp aa sequences of orthotospoviruses revealed that MerV1 is related to viruses in phylogroup C (*i.e.,* a clade of phylogenetically-related orthotospoviruses) [61] (Fig. 4i).

**Figure 4.**
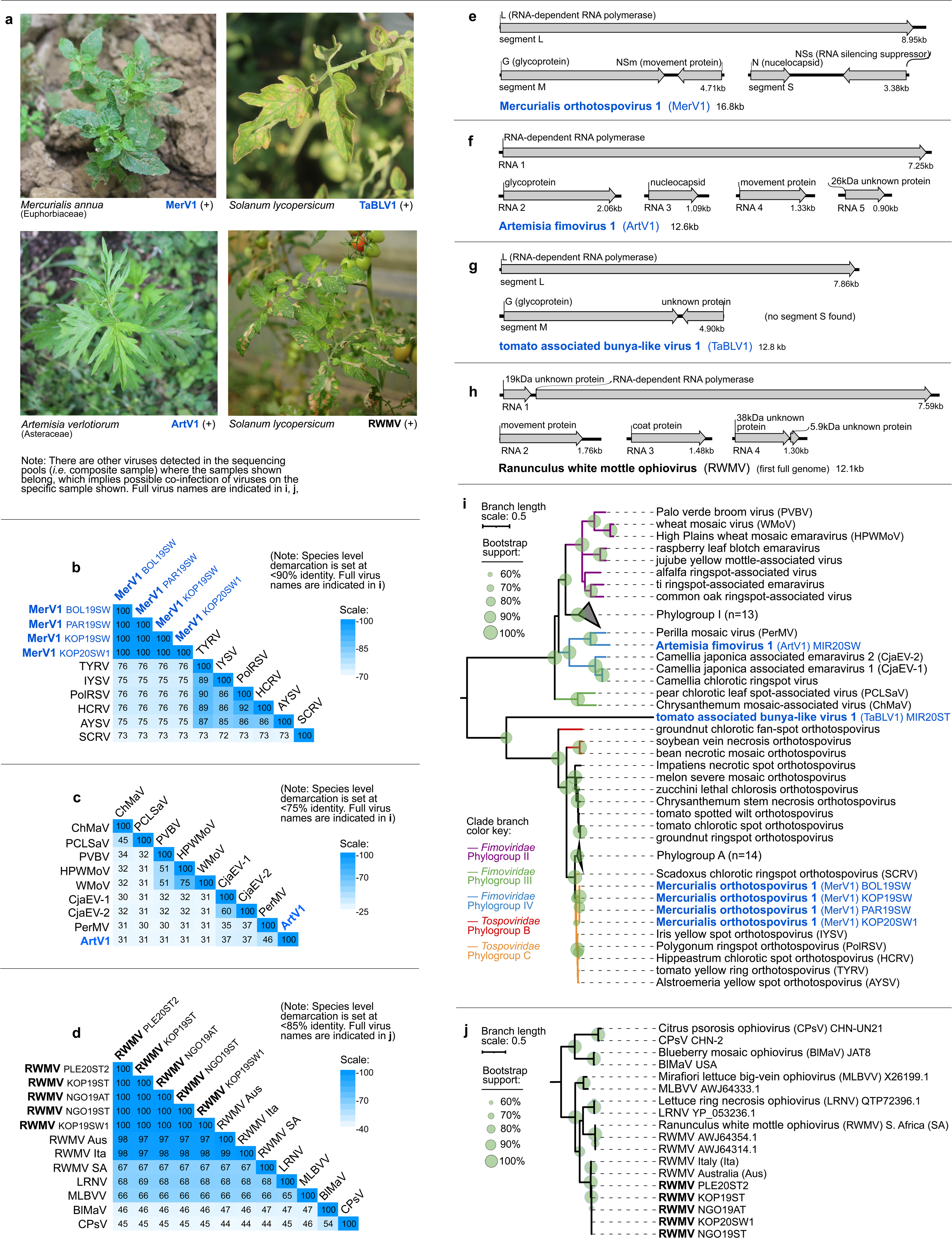
Characteristics of (-)ssRNA viruses under family *Tospoviridae, Fimoviridae*, and *Aspiviridae*, from the present study. **a** Selected photographs showing associated symptoms in field samples that tested positive for the respective viruses in RT-PCR assays. **b-d** Heatmaps showing the pairwise identities of selected viruses based on alignment and comparison of full length RdRp amino acid sequences. **e-h** Genome organization of the viruses presented herewith showing the predicted open reading frames and protein products. **i-j** Maximum likelihood phylogenetic trees based on the conserved RNA-dependent RNA polymerase (RdRp) amino acid sequences of *Tospoviridae* and *Fimoviridae* also showing tomato associated bunya-like virus 1 (**i**), and *Aspivirdae* (**j**). Branch length scale represents amino acid substitution per site. Virus names and acronyms in blue bold font are the novel viruses, while those in black bold font are known viruses from the present study. Indicated after the name or acronym are the isolate IDs of the viruses. Full virus names of those abbreviated in the pairwise identity matrices (**b-d**) can be found in the genome organization and phylogenetic trees (**e-j**).

A novel fimovirus, Artemisia fimovirus 1 (ArtV1) was detected in *Artemisia verlotiorum.* It is 30.2-46.1% identical to related *Fimoviridae* species based on RdRp aa comparison, and has genome organization similar to plant fimoviruses (Fig. 4a,c,f). Phylogenetic analyses with known fimoviruses revealed that ArtV1 is related to *Perilla mosaic virus,* making it the fourth member of the divergent phylogroup IV of *Fimoviridae* (*i.e.,* a clade of phylogenetically-related fimoviruses) [62, 63] (Fig. 4i).

A novel bunya-like virus, closely related to *Fimoviridae* and *Tospoviridae,* was detected in seven symptomatic tomatoes using RT-PCR assays, thus named tomato associated bunya-like virus 1 (TaBLV1). Pairwise comparison of RdRp aa sequences showed that TaBLV1 is only 20.8-22.4% identical to closely related bunyaviruses, however, only L- and M-like genome segments were found for TaBLV1 (L, M and S segments are characteristic of plant-infecting orthotospoviruses) (Fig. 4a,g). TaBLV1 is more closely related to orthotospoviruses based on RdRp phylogenetic analyses (Fig. 4i).

We assembled for the first time, the full genome of RWMV, which was previously detected in tomatoes and peppers in Slovenia [57] (Fig. 4a,d,h). Using RT-PCR assays, RWMV was detected in three different localities, in eight tomatoes, and one *S. nigrum*, which is a newly identified associated host. All RWMV isolates form a single clade (100% bootstrap support (BS)), but Slovenian isolates cluster separately (98.9% BS) from the Australian and Italian subclade (96.6% BS) (Fig. 4j). For details of the pairwise comparisons, see Supplementary Table 23-25. Phylogenetic trees where the subtrees in Fig. 4 were derived depicting *Tospoviridae* and *Fimoviridae* families in one tree, and *Aspiviridae* with other related viruses in one tree, are shown in Supplementary Fig. 4-01 and 4-02, where the GenBank accession numbers of the sequences used in the analyses are also indicated.

### Discovery of novel rhabdoviruses and their links to Solanaceae and Asteraceae hosts

In this study, nine novel rhabdoviruses were discovered in tomatoes and weeds. Using RT-PCR assays, tomato alphanucleorhabdovirus 1 (TARV1), tomato betanucleorhabdovirus 1 (TBRV1) and TBRV2 were confirmed in tomatoes. Five were detected in Asteraceae weeds:

*P. echoides* (Picris betanucleorhabdovirus 1 (PBRV1), Picris cytorhabdovirus 1 (PiCRV1)), *T. officinale* (Taraxacum betanucleorhabdovirus 1 (TarBRV1) and Taraxacum cytorhabdovirus 1 (TCRV1)), *Cirsium arvense* (Cirsium cytorhabdovirus 1 (CCRV1)). One cytorhabdovirus was detected in *Pastinaca sativa* (Apiaceae) (Pastinaca cytorhabdovirus 1 (PaCRV1)). TarBRV1 and TCRV1 were found to be co-infecting a single *T. officinale* sample, and PBRV1 and PiCRV1 co-infecting one *P. echoides* sample (Fig. 5a). Viruses were distinguished based on pairwise comparison of full genome nucleotide (nt) sequences (Fig. 5b-d). All novel rhabdoviruses have genomes typical of their genus, except for the highly divergent PiCRV1, which might have a putative bipartite genome, or the two contigs were not assembled together in our analyses (Fig. 5e). PiCRV1 is only 29.1-33.5% identical to closely related cytorhabdoviruses, based on comparison of RdRp aa sequences (Supplementary Table 26-B).

Investigation of virus-host relationships was done by drawing links between rhabdovirus phylogram and plant host cladogram (Fig. 5f). This revealed that rhabdoviruses from certain clades are often associated with Solanaceae and Asteraceae plant hosts. In particular, a clade of closely related alphanucleorhabdoviruses, including one discovered from this study (potato yellow dwarf virus, joa yellow blotch virus, eggplant mottled dwarf virus, PhCMoV, TARV1), was associated with eight plant species from family Solanaceae, both experimentally and as natural hosts. A few associations of betanucleorhabdoviruses discovered in weeds (PBRV1, TarBV1), and in tomatoes (TBRV1, TBRV2) with solanaceous plants were also found. Two novel betanucleorhabdoviruses (PBRV1, TarBV1) from this study and one known virus (Sonchus yellow net virus) were associated with three Asteraceae plants. For details of the pairwise comparisons, see Supplementary Tables 26-28. An identical rhabdovirus phylogenetic is shown in Supplementary Fig. 4-03, where the GenBank accession numbers of the sequences used in the analyses are indicated.

### Detection of a kitavirus in tomatoes and other known and novel *Martellivirales* species

The recently described tomato fruit blotch virus (ToFBV, *Kitaviridae*) [28,64,65], was detected in two symptomatic tomato leaf samples from Slovenia (Fig. 6a). Infectivity and possible vector of ToFBV were not reported yet, and it is currently classified in the genus *Blunervirus.* Pairwise comparison of conserved RdRp aa sequences of blunerviruses (Fig. 6b), revealed high molecular divergence among members, with only 30.9-41.3% pairwise identity between species. Phylogenetic analyses using conserved RdRp aa sequences showed that all known ToFBV isolates from a single clade (100% BS) (Fig. 6k).

**Figure 5.**
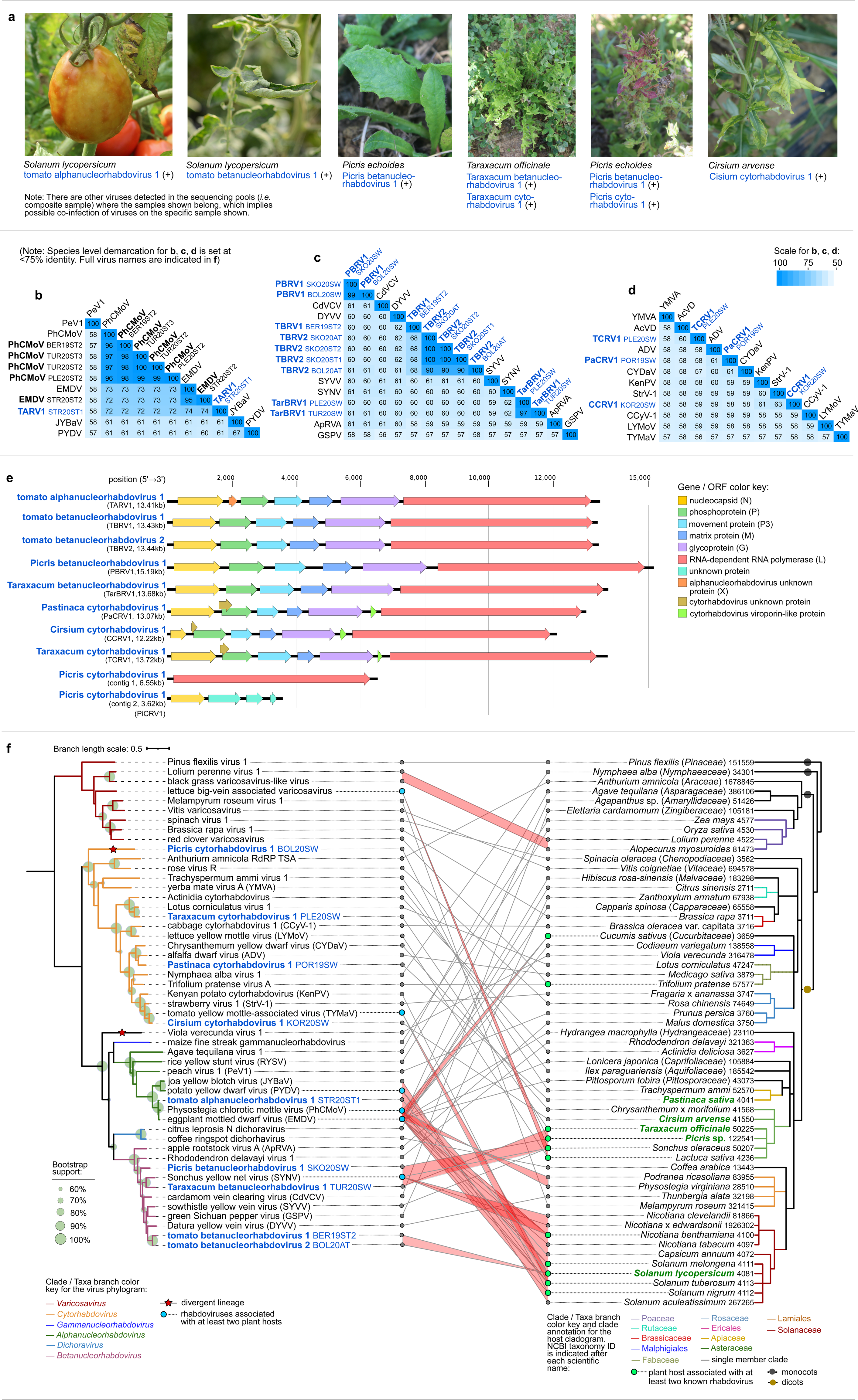
Characteristics of novel (-)ssRNA viruses classified in family *Rhabdoviridae* from the present study. **a** Photographs of samples that tested positive for selected novel rhabdoviruses in RT-PCR assays. **b-d** Heatmaps showing the pairwise identities of species in genus *Alphanucleorhabdovirus* (**b**), *Betanucleorhabdovirus* (**c**), and *Cytorhabdovirus* (**d**), based on full genome length nucleotide sequences. Virus acronyms in blue bold font are the novel rhabdoviruses, while those in black bold font are known rhabdoviruses from the present study. Indicated after virus names and acronyms are the isolate IDs. **e** Genome organization of novel rhabdoviruses discovered in the present study. Genome lengths are shown to scale. Open reading frames (ORFs) and the proteins they code for are color coded accordingly. **f** Co- phylogenetic tree (tanglegram) showing the phylogenetic relationships of novel rhabdoviruses among known species (left tree), which are linked with associated plant host(s) indicated in GenBank, shown on the right tree. Numerous links of well-supported clade of viruses to taxonomically-related plants are highlighted in red. Maximum likelihood phylogenetic tree of rhabdoviruses was constructed **based** on the conserved amino acid sequence of the RdRp. Branch length scale represents amino acid substitution per site. The host cladogram was made in phyloT (www.phylot.biobyte.de). In the virus tree, viruses associated with at least two plant hosts are indicated by a blue circle. In the host tree, green circles are used to indicate plants that are associated with two or more rhabdoviruses, and the host species names in bold green font are those that have representative samples in the present study. The clades are separately color coded and annotated for each tree. Full virus names of those abbreviated in the pairwise identity matrices (**b-d**) can be found in the genome organization and phylogenetic trees (**e, f**).

**Figure 6.**
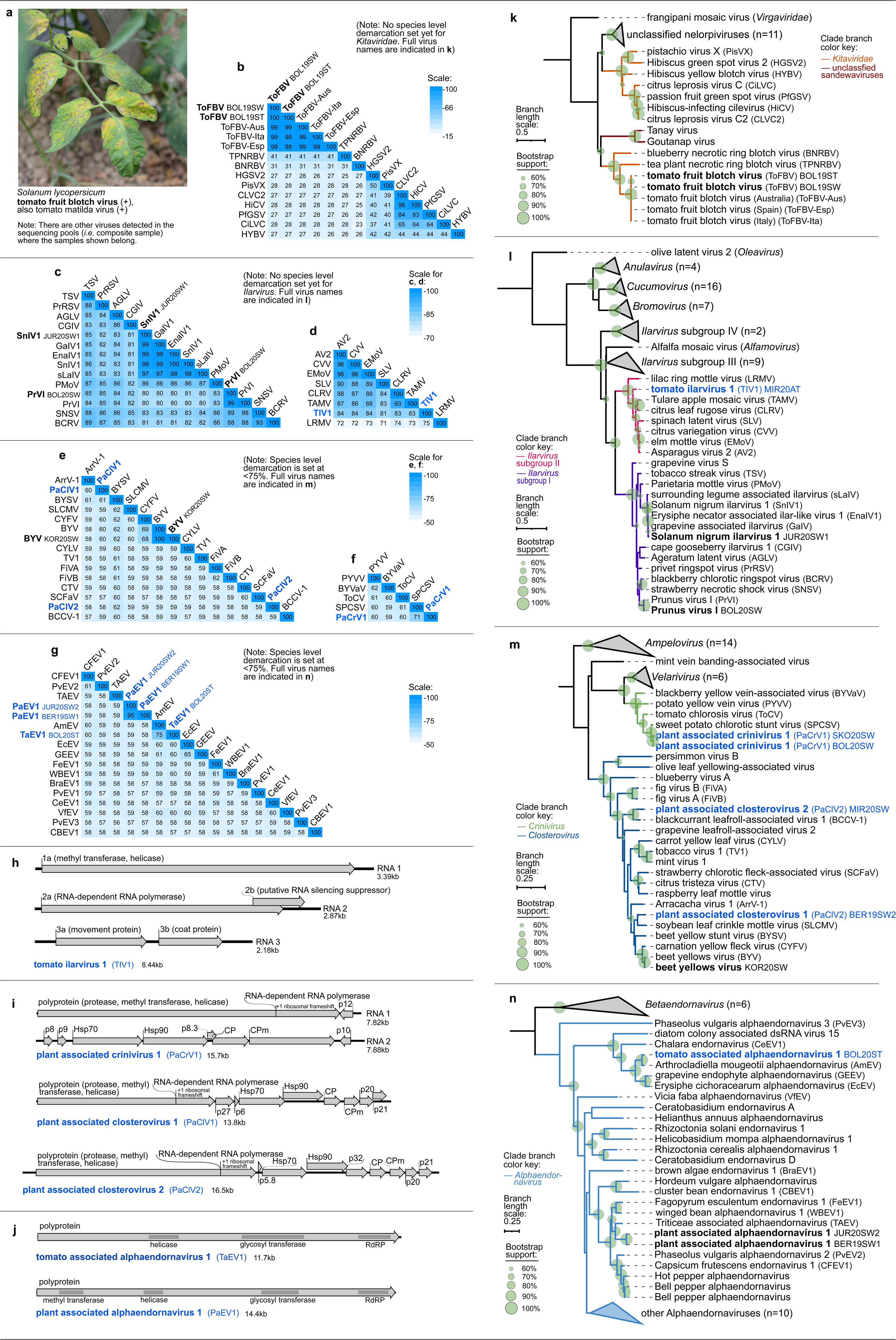
Characteristics of (+)ssRNA viruses in families *Kitaviridae*, *Bromoviridae*, *Closteroviridae*, and *Endornaviridae* from the present study. **a** Photograph of one of the symptomatic tomato samples that tested positive for ToFBV in RT-PCR assays. **b-g** Heatmaps showing the pairwise identities to distinguish different *Kitaviridae* (**b**) and *Bromoviridae* (**c, d**) species based on alignment and comparison of RdRp amino acid sequences, members of *Closterovirus* genus (**e**) based on full genome nucleotide sequences, members of *Crinivirus* genus (**f**) based on full RNA1 segment nucleotide sequence, and members of *Alphaendornavirus* (**g**) based on full genome nucleotide sequences **h-j** Genomes of the novel viruses, with the predicted open reading frames, protein domains, and genome size. **k-n** Maximum likelihood phylogenetic trees constructed based on the conserved amino acid sequence of the RdRp of (**k**) *Kitaviridae*, (**l**) *Bromoviridae*, (**m**) *Closteroviridae*, and (**n**) *Endornaviridae*. Branch length scale represents amino acid substitution per site. Indicated after the acronym are the isolate IDs of the viruses. Virus names and acronyms in blue bold font are the novel viruses, while those in black bold font are known viruses from the present study. Full virus names of those abbreviated in the pairwise identity matrices (**b-g**) can be found in the genome organization and phylogenetic trees (**h-n**).

**Figure 7.**
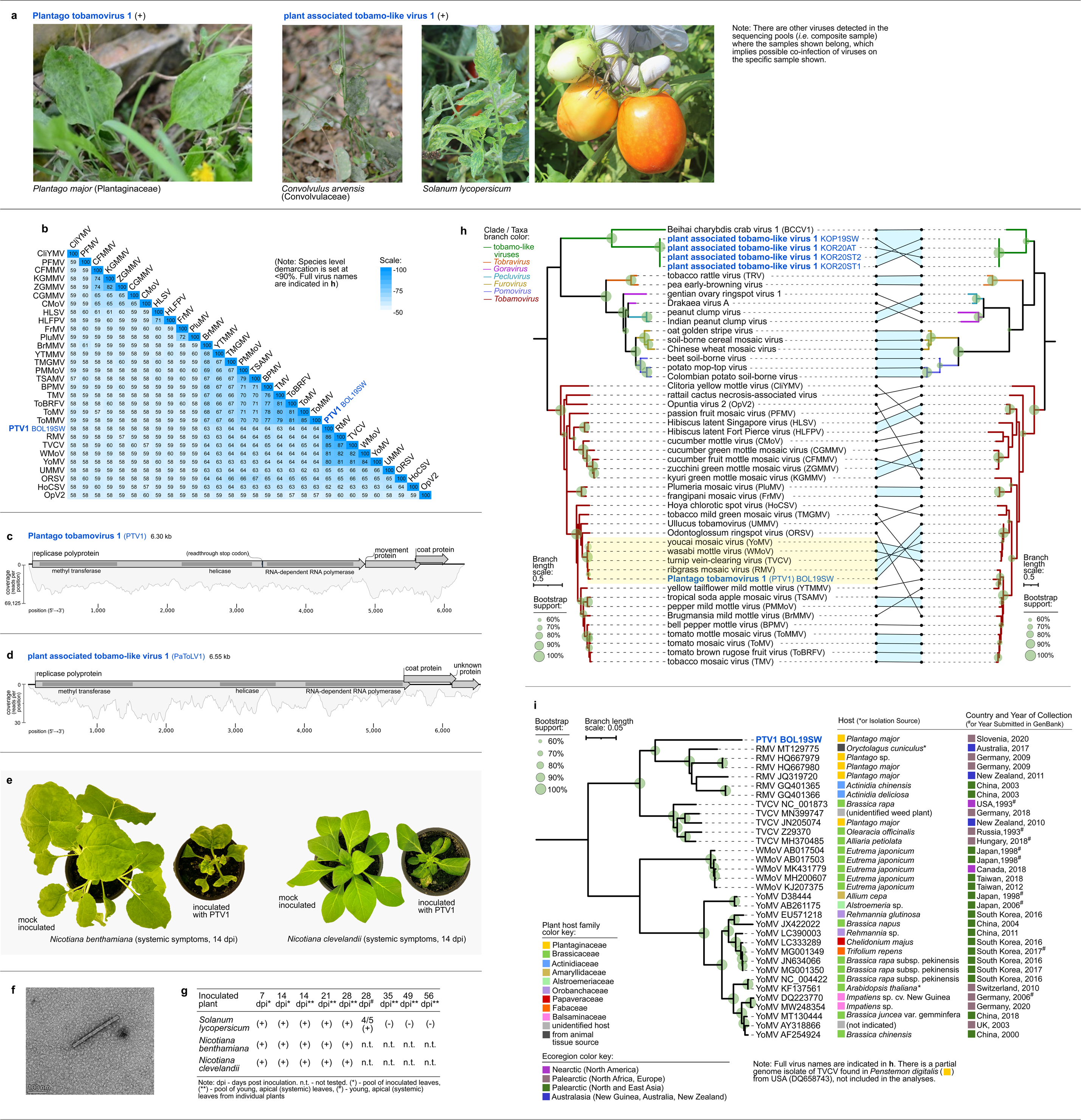
Characteristics of (+)ssRNA viruses in family *Virgaviridae* from the present study. **a** Selected photographs of samples that tested positive in RT-PCR assays for the viruses presented herewith. **b** Heatmap of pairwise identities of *Tobamovirus* (*Virgaviridae*) species based on full genome nucleotide sequences. **c, d** Genome organization and read coverage of novel viruses showing the open reading frames, and the proteins they code for. **e** Photographs of the PTV1 inoculated symptomatic plants and mock inoculated control plants. **f** Electronmicrograph showing particle morphology of PTV1. **g** RT-PCR assays of test plants inoculated with Plantago tobamovirus 1 (PTV1). **h** Maximum likelihood phylogenetic trees constructed based on the conserved amino acid sequence of methyl transferase-helicase (left) and coat protein (CP) (right). The phylogenetic congruence of the methyl transferase-helicase-based tree (left) is demonstrated by line connecting identical viruses to the CP-based tree (right). Congruent clades with >60% bootstrap support are highlighted (light blue). The clade of viruses closely related to PTV1 is highlighted in yellow. **i** Maximum likelihood phylogenetic tree, based on full genome nucleotide sequences, of all the isolates of a subgroup of tobamoviruses including PTV1. Virus names and acronyms in blue bold font are the novel viruses from the present study. Indicated after the acronym are the isolate IDs of the viruses. Branch length scale represents amino acid (**h**) or nucleotide substitution (**i**) per site. The hosts or isolation sources of the viruses as indicated in NCBI GenBank, and the countries and dates of collection are indicated. Full virus names of those abbreviated in (**b**) and (**i**) are indicated in (**c-d**) and (**h**).

SnIV1, a recently described ilarvirus, was detected in *Physalis* sp., which is the sixth distinct source association with the virus, aside from three other plant species (*S. lycopersicum, S. nigrum*, and an unidentified legume plant) [23, 29], and grapevines infected with a fungus (*Erysiphe necator*) and an oomycete (*Plasmopara viticola*) [66, 67]. PrVI, recently discovered in sweet cherry [68], was detected in our study in *P. echoides*, co-infected with PiCRV1. Pairwise comparison of RdRp aa sequences of the isolates of SnIV1 and PrVI from this study showed that they are >98% identical to other isolates of each species (Fig. 6c). A novel subgroup II ilarvirus, tomato ilarvirus 1 (TIV1), was detected in an asymptomatic tomato sample, and is at most 83.3% identical to closely related ilarviruses based on RdRp aa sequence comparison (Fig. 6d,h).

Three new *Closteroviridae* species were detected in weeds through HTS (Fig. 6e,f,i).Two new alphaendornaviruses with their typical protein domains were also discovered, one of which is from a pool of symptomatic tomatoes (Fig. 6g,j). Novel ilarvirus, closterovirus and alphaendornavirus all form distinct phylogenetic clades with related viruses (>60% BS) (Fig. 6l-n). For details of the pairwise comparisons, see Supplementary Tables 29-33. Expanded versions of the collapsed phylogenetic trees in Fig. 6 are shown in Supplementary Fig. 4-04, 05, 06, and 07, where the GenBank accession numbers of the sequences used in the analyses are indicated. Details of InterPro domain search are in Supplementary Tables 43-44.

### Characteristics of a new tobamovirus, and a divergent tobamo-like virus species

A new tobamovirus, Plantago tobamovirus 1 (PTV1), was discovered in *Plantago major* (Fig. 7a), and percent pairwise identities based on whole genome nt sequences revealed that PVT1 is 80.4-85.7% identical to closely related tobamoviruses, which is below the current species demarcation at <90% (Fig. 7b). PTV1 has genome composition and organization typical of tobamoviruses (Fig. 7c). Maximum likelihood phylogenetic analyses using conserved methyl transferase-helicase aa sequences showed that PTV1 is closely related to ribgrass mosaic virus (RMV), turnip vein clearing virus (TVCV), youcai mosaic virus (YoMV), and wasabi mottle virus (WMoV) (Fig. 7h, highlighted in yellow). Phylogenetic analyses using full nt genome sequences of all known isolates from this tobamovirus clade including RMV, TVCV, YoMV, WMoV revealed relatedness but distinct divergence of PTV1 from the RMV clade (Fig. 7i). By superimposing information on host and country of origin (Fig. 7l), we found that aside from PTV1, *P. major* (Plantaginaceae) is a common host for RMV from Germany, and for RMV and TVCV in another eco-geographically distinct country, New Zealand [69, 70]. Systemic infection with disease symptoms, which was confirmed by RT-PCR, was observed in *Nicotiana benthamiana* and *Nicotiana clevelandii* that were mechanically-inoculated with PTV1-infected *P. major* leaf tissues. Inoculated tomato plants remained asymptomatic, however, PTV1 was detected by RT-PCR in systemic leaves until 28 dpi but not at 35 dpi and beyond (Fig. 7e-g, Supplementary Fig. 5), which might be a consequence of either transient infection or cross-contamination during the experiment. Transmission electron microscopy of a *N. clevelandii* infected plant revealed PTV1 virions that are rigid rods, typical for tobamoviruses (Fig. 7f). HTS of the same plant confirmed the presence of PTV1 in systemic tissues (Supplementary Fig. 5).

A tobamo-like virus, PaToLV1, was detected in symptomatic tomatoes and in *Convolvulus arvensis* (Fig. 7a). Pairwise identity analysis using RdRp aa sequences showed that PaToLV1 is <31% identical to Plumeria mosaic virus (*Tobamovirus*), tobacco rattle virus (*Tobravirus*) and Beihai chrarybdis crab virus 1 (BCCV1, unclassified *Riboviria*) (Supplementary Table 35). PaToLV1 has genome structure similar to BCCV1, except for an additional unknown open reading frame (ORF) (Fig. 7d). Phylogenetic analyses based on the RdRp and CP aa sequences both revealed that PaToLV1 isolates form a monophyletic clade, which is clustered together with BCCV1 (Fig. 7h). For details of the pairwise comparisons, see Supplementary Table 34-35. Detailed version of the *Virgaviridae* phylogenetic trees in Fig. 7 are shown in Supplementary Fig. 4-08-A (methyl transferase, helicase-based) and Supplementary Fig. 4-08-B (coat protein-based), where the GenBank accession numbers of the sequences used in the analyses are indicated. Details of InterPro domain search for virus genomes are in Supplementary Tables 45-46.

### Characteristics of novel *Potyviridae* species, and crop-infecting potyviruses found in weeds

Aside from PVY, two known potyviruses (WMV and CTLV), previously detected in Slovenian crops and ornamentals [71, 72], were detected in pools of weed plants from this study. Full genomes of these viruses were detected in composite samples from five different tomato farms, encompassing two localities in the Primorska region (SW Slovenia: Koper,

Piran). Henbane mosaic virus, previously found in tomatoes from Slovenia[56], was detected in a composite sample of open field-grown symptomatic tomatoes. Three novel members of *Potyviridae* (Mentha macluravirus 1 (MenMV1), Plantago potyvirus 1 (PlaPV1) and Rumex potyvirus 1 (RumPV1)) were discovered in weeds showing virus disease-like symptoms (Fig. 8a). Pairwise comparison of full-length (nt, aa) polyprotein ORF distinguished the new species from the known ones based on molecular demarcation criteria for *Potyviridae* (<82% aa, <76% nt) [73] (Fig. 8b-e). The genome and polyprotein ORF of novel potyviruses were characterized, and all protein domains, typical of macluraviruses and potyviruses were found (Fig. 8f). Phylogenetic analyses based on conserved RdRp aa sequence placed MenMV1 in a clade with artichoke latent virus and Narcissus latent virus (100% BS) (Fig. 8g). PlaPV1 was placed in a clade with cucurbit vein banding virus (83.1% BS), and RumPV1 was placed in a clade with lotus latent virus and Calystegia hederacea virus (100% BS).

**Figure 8.**
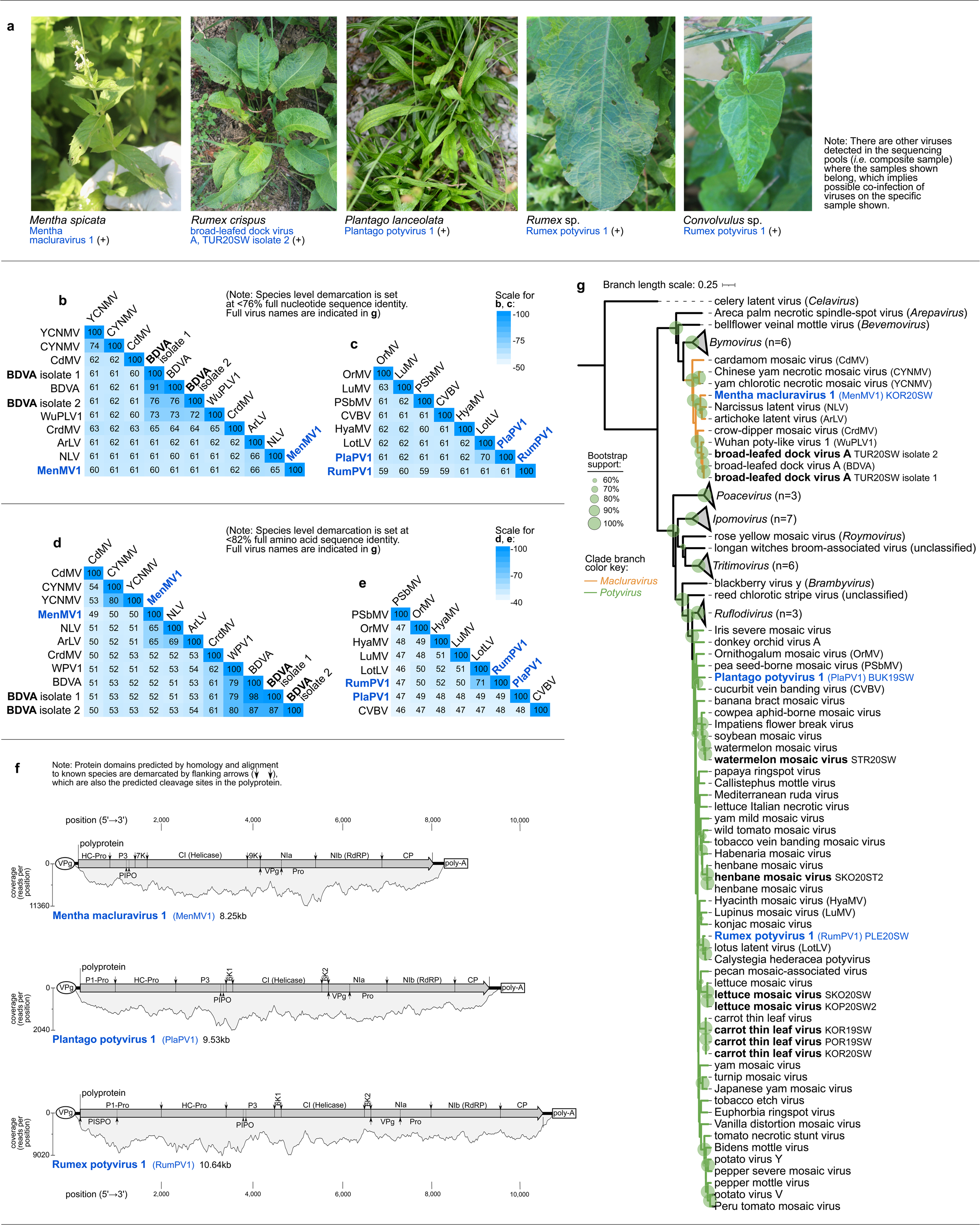
Characteristics of (+)ssRNA viruses from family *Potyviridae* from the present study. **a** Selected photographs of samples that tested positive in RT-PCR assays for the viruses presented herewith. **b-e** Heatmaps of the pairwise identities of *Macluravirus* and *Potyvirus* species based on alignment and comparison of full genome nucleotide sequences (**b-c**) and based on full polyprotein amino acid sequence (**d-e**). **f** Genome organization with predicted cistrons and read coverage for novel potyviruses from this study. **g** Maximum likelihood phylogenetic tree based on the alignment of conserved RdRp amino acid sequence of RefSeq species from the *Potyviridae* family. Branch length scale represents amino acid substitution per site. Virus names and acronyms in blue bold font are the novel viruses, while those in black bold font are known viruses from the present study. Indicated after the acronym are the isolate IDs of the viruses. Full virus names of those abbreviated in the pairwise identity matrices (**b-e**) can be found in the genome organization and phylogenetic trees (**f-g**).

Two different lineages of broad-leafed dock virus A (BDVA), were also detected. A lineage with a considerably low pairwise identity compared with original isolate (86.6% aa, 76.0% nt) was found in *Rumex crispus*. For details of the pairwise comparisons, see Supplementary Tables 36-37. An expanded version of the collapsed *Potyviridae* phylogenetic tree in Fig. 8, is shown in Supplementary Fig. 4-09, where the GenBank accession numbers of the sequences used in the analyses are indicated. Details of protein domain and cleavage site search for virus genomes are in Supplementary Table 47.

### Known tomato tombusviruses, and discovery of new *Tombusviridae* species and tombusvirus-like associated RNAs in weeds

Genomes of three known members of *Tombusviridae* (tomato bushy stunt virus (TBSV), tobacco necrosis virus A (TNVA), olive latent virus 1 (OLV1)), and partial genome of olive mild mosaic virus (OMMV), were assembled from sequences derived from composite tomato samples. Five new viruses classified in four different *Tombusviridae* genera were discovered in weeds showing virus disease-like symptoms (Fig. 9a). These include two new umbraviruses (Pastinaca umbravirus 1 (PasUV1) and Picris umbravirus 1 (PicUV1)), one new aureusvirus (Convolvulus aureusvirus 1 (ConAV1)), one new pelarspovirus (Calystegia pelarspovirus 1 (CalPV1)), and one new alphacarmovirus (Cichorium alphacarmovirus 1 (CicAV1)). Three sequences, with similarity to self-replicating, coat-dependent RNA replicons, called tombusvirus-like associated RNAs [74], were also detected in composite weed samples. Pairwise comparisons of full genome nt or RdRp (replicase) aa sequences were done, whichever is appropriate for species demarcation (Fig. 9b-f). Genome organization typical for the genera was determined for the new *Tombusviridae* members. Novel unknown ORFs were detected in Picris umbravirus 1 and CalPV1 (Fig. 9g). Genome compositions and organizations of new tombusvirus-like associated RNAs were similar to those previously described [74], except for the two plant associated tombusvirus-like RNA 1 isolates, in which ambisense ORFs were detected. Phylogenetic analyses placed the novel tombusvirus-like associated RNAs in a clade with Arracacha latent virus E associated RNA, a member of subgroup III tombusvirus-like associated RNAs (78.7% BS) [74]. Novel tombusviruses were likewise placed in clades with known species of their respective genera (>60% BS) (Fig. 9h). For details of the pairwise comparisons, see Supplementary Tables 38-42. An expanded version of the collapsed *Tombusviridae* phylogenetic tree in Fig. 9, with providence virus (*Carmotetraviridae*) as outgroup, is shown in Supplementary Fig. 4-10, where the GenBank accession numbers of the sequences used in the analyses are also indicated.

**Figure 9.**
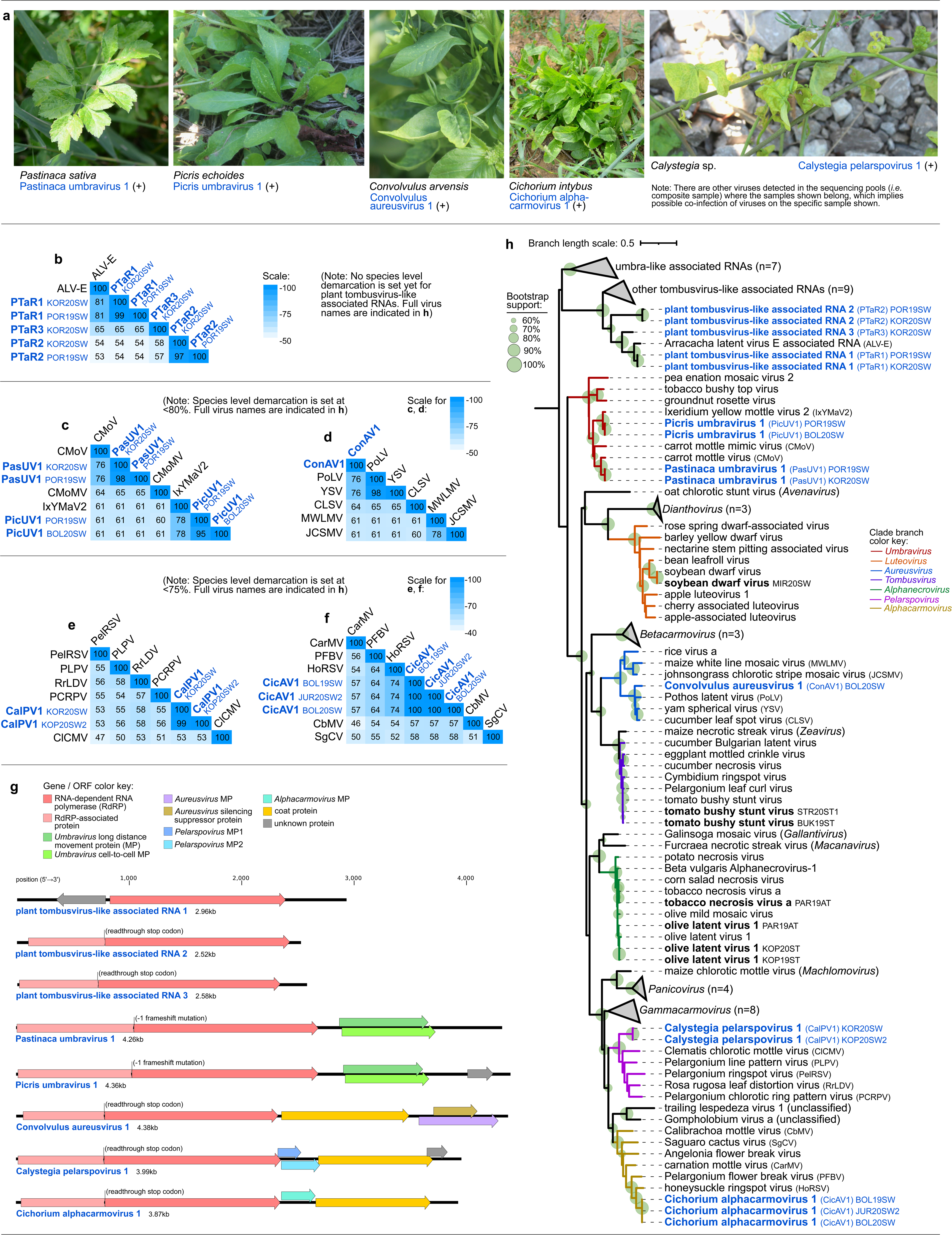
Characteristics of (+)ssRNA viruses in family *Tombusviridae* from the present study. **a** Selected photographs of samples that tested positive in RT-PCR assays for the viruses presented herewith. **b-f** Heatmaps of pairwise identities of plant tombusvirus-like associated RNAs based on alignment and comparison of RdRp amino acid sequences (**b**), *Umbravirus* species based on full genome nucleotide sequences (**c**), *Aureusvirus* species (**d**), *Pelarspovirus* species (**e**), and *Alphacarmovirus* species (**f**) based on RdRp amino acid sequences. **g** Genome organization of novel *Tombusviridae* species and tombusvirus-like associated RNAs from this study. Open reading frames (ORFs) and the proteins they code for are color coded accordingly. **h** Maximum likelihood phylogenetic tree based on the conserved amino acid sequence of RdRp. Virus names and acronyms in blue bold font are the novel viruses, while those in black bold font are known viruses from the present study. Indicated after the acronym are the isolate IDs of the viruses. Branch length scale represents amino acid substitution per site. Full virus names of those abbreviated in the pairwise matrices in (**b-f**) are in the phylogenetic tree (**h**).

### Discovery of other numerous viruses, satellite RNAs, and viroid-like circular RNAs

Aside from viruses mentioned above, a diverse set of 37 classified known and novel viruses, 21 putative virus-like *Riboviria* species (*i.e.,* cannot be classified in known virus taxa), and four known and new satellite RNAs were also detected (Supplementary Table 5). Genome organization of these viruses and phylogenetic trees including the viruses are shown in Supplementary Fig. 1 and 4. Aside from STV, other dsRNA viruses were discovered, including six new *Partitiviridae* members from weeds, two of which were detected in 2019 and 2020 at the same farm. Three new *Totiviridae* members were discovered, two of which are phylogenetically related to plant-associated members of the family. Circular DNA viruses were discovered, including two new caulimoviruses found in weeds, and partial genome of a new geminivirus (Calystegia geminivirus 1), from a symptomatic *Calystegia* sp. sample co-infected with CalPV1. To our knowledge, this could be one of the few geminiviruses in wild plants from Europe, since the majority of geminiviruses were found in Asia and the Americas [75]. Several other (+)ssRNA viruses were also discovered and classified under *Alphaflexiviridae* (n=1), *Dicistroviridae* (n=3)*, Iflaviridae* (n=1), *Secoviridae* (n=5), and *Solemoviridae* (n=3). The new alphaflexivirus (Pastinaca potexvirus 1) was detected in a *P. sativa* sample co-infected with PasUV1 and PaCRV1. One new satellite virus (tomato albetovirus 1) and new satellite RNA (TBSV satellite RNA C), associated with *Tombusviridae* species [76, 77], were detected in tomatoes. Six putative virus-like sequences detected were previously associated with arthropod [4], fungi [66], or oomycete [67] hosts, while other 15 are highly divergent ones with unidentified hosts. Using a custom discovery pipeline based on SLS-PFOR2 [45], we assembled and screened 4249 circular RNAs, and identified 344 viroid-like candidates (Supplementary Table 6). Five viroid-like circular RNAs were preliminarily selected, based on predicted secondary structure, %GC content, and degree of branching. Genome circularity was confirmed for Taraxacum viroid-like circular RNA 1 (Supplementary Fig. 2), which was detected in a *T. officinale* sample co-infected with TarBV1.

## Discussion

Virus diversity in specific crops is relatively well-studied compared to weed plants or crop’s wild relatives [20, 21]. A few studies used HTS to uncover virus diversity and possible exchanges in the cultivated and wild compartments of agroecosystems [23, 29]. Here, we detected 37 known and 55 novel viruses, which were classified in established virus taxa, and 33 unclassified *Riboviria* members. This indicates a highly understudied diversity of novel viruses in an agroecosystem, especially in weeds that might serve as reservoirs of viruses that could infect crops. This study represents the largest survey of viromes of diverse weed species within tomato agroecosystems or any cropping system. Forty-six novel viruses classified in known taxa, and additional 19 novel unclassified viruses, were detected in weeds, some of which are also present in tomatoes (*e.g.,* PaToLV1, LtaRLV1). Overall, the viromes were dominated by 60 (+)ssRNA viruses (under *Martellivirales, Potyviridae*, and *Tombusviridae*) and 17 (-)ssRNA viruses (under *Mononegavirales*). Another study looked at the exchanges of viruses between tomato and a wild relative, *S. nigrum,* in France, wherein PVY and SnIV1 were found to be present in both, cultivated and wild compartments [23]. Here we observed the same pattern, finding PVY and SnIV1 in tomato and a weed (*Physalis* sp.), and in addition, we identified new weed hosts of viruses found in tomatoes, such as RWMV in *S. nigrum*. We also detected in weed samples known viruses of other crops, which were previously detected in the country (*e.g.,* WMV, CTLV). The detection of crop-infecting viruses in weed plants implies the possible role of weeds as wild reservoirs and alternate hosts for such viruses [78].

In this study, around two-thirds (60/92) of the known and novel classified viruses detected were found exclusively in weeds surrounding tomato farms, 22 were found exclusively in tomatoes, and 10 viruses were found both in weed and tomato samples. The shared viruses include both generalist viruses (*i.e.*, viruses with known wide host range that includes wild plants) some of which are also important pathogens of tomatoes [15] (*e.g.,* TSWV, CMV, PVY). Overall, a small proportion of viruses was associated with plants from different families, which might suggest a wide host range for these viruses. Associated host plants were further confirmed using RT-PCR for 36 out of all 125 viruses detected, of which, only three (*i.e.,* TMaV, PaToLV, and Rumex potyvirus 1) were associated with at least two plants from different botanical families. Such small proportion of generalist viruses is consistent with previous hypotheses that most viruses evolved and settled as specialists that are only adapted to a few hosts (*i.e.,* have narrow host ranges), because of the fitness cost and evolutionary constraints associated to a generalist lifestyle [79]. This also implies presence of possible barriers for host-switching that could be established in the course of virus-host co-evolution [11]. Nevertheless, in this study, we collected just a portion of larger plant diversity and did not confirm plant infectivity of these viruses. Thus, post-discovery characterization and extending sampling and analysis to other crops and weeds, will help confirm these hypotheses.

Viruses infecting plants may not cause any disease symptoms, because of a rather neutral or beneficial interaction with its host, latent infection, host resistance, or modulation of environmental factors [10, 80]. A previous study of tomato viruses in China focused only on symptomatic samples [25]. In our study, HTS and RT-PCR assays further revealed viruses that are shared between tomatoes and weeds, and those that were detected in both asymptomatic and symptomatic tomatoes. In tomatoes, 45 different viruses were detected, 18 of which were found in both asymptomatic and symptomatic tomatoes. This observation has an important implication when aiming to capture a significant breadth of virus diversity through sampling across ecosystem scales. This observation also gave important insights on possible symptom masking due to virus-tomato-environment interactions leading to latency, or due to host resistance or tolerance.

Monitoring virus diversity and evolutionary dynamics of economically important and widespread viruses could help anticipate emergence of new variants, thus preventing disease outbreaks [30]. Here, we investigated the molecular diversity of viruses based on full genome nucleotide identities, and variability among virus isolates based on genome-wide polymorphic sites and nucleotide diversity scanning. These were determined on the most prevalent viruses in tomatoes, wherein some were detected for the first time in Slovenian tomatoes and are present in several localities, such as TMaV. Low diversity was observed among the isolates of ToMV and the persistent STV, which is consistent with previous reports on their global diversity [81, 82]. TSWV, in contrast to its considerable diversity at the global scale [83], showed low diversity in the samples analyzed here.. Sequenced genomes of PVY, CMV, PVM, PhCMoV, RWMV showed moderate levels of variability. PhCMoV is a recently detected, and widespread pathogen of tomato and other crops in Europe, where considerable level of diversity was observed among the isolates [84, 85]. RWMV was recently detected in Slovenian peppers and tomatoes, where isolates showed moderate diversity as well [57]. Moreover, two new viruses (TBRV2, TomV1) associated with tomatoes, showed high level of variability, which might imply that these viruses are circulating in the area for a long time already and/or are undergoing diversifying evolution. Further diversity and evolution studies of viruses found in tomatoes in our virome survey, such as ToFBV, PhCMoV, TMaV, SnIV1, RWMV, TBRV2, and TomV1 would bring insightful information about the nature of the observed variations, *e.g.*, if they are the result of neutral processes or are subjected to different kind of selection pressures and/or possible ongoing adaptation to tomato.

We observed high diversity of novel viruses at higher taxonomical levels. The most illustrative example is detection of 11 rhabdoviruses, of which 9 are newly discovered species. Three new rhabdoviruses phylogenetically-related to plant rhabdoviruses were associated with tomato, which increases the count of known tomato rhabdoviruses from two to five. In weeds, six new rhabdoviruses were detected, primarily associated with Asteraceae species. Our observations added to the discoveries made in other studies, *e.g.,* a recent discovery of 27 new viruses that are phylogenetically-related to plant rhabdoviruses through homology searches in the transcriptome shotgun assembly database of NCBI [86]. Some rhabdoviruses are known to replicate in its arthropod vector as well [87], thus, further infectivity tests are needed to confirm their infectivity and pathogenicity in associated plant hosts.

Among the most economically important viruses of tomato, tobamoviruses are the most problematic in recent years, due to the emergence of ToBRFV [16]. We detected a novel tobamovirus (PTV1) in *Plantago major* surrounding a greenhouse farm in Slovenia. We demonstrated the systemic infection of PTV1 in various Solanaceae hosts, but more extensive biological and genetic characterization is needed to understand its biological characteristics. Nevertheless, such discovery calls for further research on risks associated with tobamoviruses as one of the major causes of disease outbreaks in tomato and other solanaceous crops [15]. A novel tobamo-like virus (PaToLV1) was also detected in tomatoes and in *Convolvulus arvensis*. A previous study showed the evolutionary relationship of the most similar known virus, BCCV1, to *Virgaviridae* and *Martellivirales* families [4], which could also be the case for PaToLV1.

Moreover, besides viruses, we also attempted to uncover new viroid-like agents in our dataset, with the aim to contribute to their possibly still undiscovered diversity [45]. We obtained five candidate viroids, wherein one was experimentally confirmed to be circular and structurally similar to members of *Avsunviroidae* due to the presence of a hammerhead ribozyme motif [88]. These results suggest that further research in this direction and development of new bioinformatics tools for viroid discovery within sequencing datasets could uncover even larger hidden diversity of these non-protein coding pathogenic agents.

Aside from studying virus diversity and evolution, understanding of host-virus co-evolutionary relationships is also essential in predicting possible virus emergence through host-switching [89]. We observed an association of a clade of closely related alphanucleorhabdoviruses with plant species from family Solanaceae. However, the general topology of tanglegram suggested discordant evolutionary historic events between plant rhabdoviruses and their hosts, which is similar to what was previously shown in animal/insect rhabdoviruses [89]. Likewise, we found frequent association of different but related viruses with a single plant host. This is the case of *Plantago major,* which was associated with PTV1, RMV, and TVCV isolates from four different countries from two very distinct eco-geographical regions. This subgroup was originally associated with brassica hosts [90] and recently, as reported in this study, with several Plantaginaceae and Actinidiaceae plants. This observation indicates a possibility of virus spread through introduction of plants, followed by diversifying evolution through a mix of confounding environmental and host-related factors [91].

## Conclusions

Viruses, viroids and other virus-like agents are well-studied in crops, but known to a much lesser extent in non-crop plants. Their phytosanitary implications and economic importance in crops such as tomato [15], resulted to an increased interest to discover novel species. Here, we presented an extensive viromics study, where we uncovered the vast diversity of viromes of plant species in defined agroecosystems linked with tomato production. Similar systematic surveys in various crop agroecosystems, which could also include fungal, oomycete, arthropod or nematode vectors and environmental media, such as soil and water should, in the future, provide more virome data from still undescribed compartments of agroecosystems. Collectively, this will help uncover fraction of the unknown viruses in the global virosphere [92], and including their yet unknown ecological interactions and functions [8, 9]. Overall, our study contributed valuable information that will, in future studies, help predict possible virus emergence in the wild-cultivated agroecological interface, a zone of possible virus spillovers and disease outbreaks [11,31,32].

## Supplementary information

**Additional File 1: Supplementary Table 1.** Individual plant samples, their species names, corresponding plant families and sampling locations. **Supplementary Table 2.** Composite samples used in high-throughput sequencing, their composition, year of collection, farm type and other associated information. **Supplementary Table 3.** Sequence read archive (SRA) metadata of the sequencing experiments, read counts, and other associated information.

**Additional File 2: Supplementary Table 4.** List summarizing all viruses detected in the composite samples. **Supplementary Table 5.** List of analyses done in the reconstruction and identification of individual virus genomes. **Supplementary Table 6.** List of circular RNAs assembled using a custom pipeline based on SLS-PFOR2, with filtering using BLASTn and BLASTx searches.

**Additional File 3: Supplementary Figure 1**. Genome organization of novel viruses, first full genomes of known viruses showing known and putative open reading frames and the protein they code for and predicted secondary structures of selected viroid-like circular RNAs detected in this study.

**Additional File 4: Supplementary Table 7.** RT-PCR primers and PCR conditions used in confirmation of associated plant hosts of selected viruses and putative viroids. **Supplementary Table 8.** RT-PCR thermocycling conditions used in the detection of selected viruses and putative viroids in associated plant hosts. **Supplementary Figure 2**. Orientation and annealing sites of the primers designed for the amplification of the circular genome of Taraxacum viroid-like circular RNA 1, and results of RT-PCR confirmation of the circular genome. **Supplementary Table 9.** List of confirmed associated plant hosts of selected viruses and putative viroids.

**Additional File 5: Supplementary Figure 3**. Representative field photos of confirmed associated plant hosts of selected viruses and putative viroids.

**Additional File 6: Supplementary Table 10.** Model selection and other parameters used for phylogenetic analyses. **Supplementary Figure 4.** Phylogenetic trees up to the virus family or order level showing relationships of identified known and novel viruses in this study.

**Additional File 7 (Data Source): Supplementary Table 11-22.** Summary of all results from the DnaSP v. 6 analyses.

**Additional File 8 (Data Source): Supplementary Table 23-42.** Summary of all pairwise identity values from the SDT v. 1.2 analyses.

**Additional File 9 (Data Source): Supplementary Table 43-47.** Summary of all results from the protein domain prediction using InterPro scans, and homology alignments with known species of potyviruses to identify start and cleavage sites of proteins.

**Additional File 10: Supplementary Figure 5.** RT-PCR, TEM and nanopore sequencing results on the confirmation of infectivity of Plantago tobamovirus 1 in solanaceous hosts.

## Declarations

### Authors’ contributions

M.R., D.K., and I.G.A. acquired funding. D.K., M.R., and M.P.S.R. formulated and designed the study. D.K. and M.R. supervised the study. M.P.S.R. did the laboratory and greenhouse work, the data analyses, and wrote the first draft of the manuscript. K.B. did the electron microscopy, and A.P. did a part of the nanopore sequencing of inoculated plants and data analyses. The rest of the authors significantly helped in sampling and sample processing. All authors contributed to editing the final manuscript.

## Supporting information

Additonal-File-01_Sample-Phenodata_Sequencing-Metadata

Additonal-File-02_All-Viruses-Lists

Additional-File-03_Genome Organization

Additional-File-04_RT-PCR Confirmations of Selected Viruses

Additional-File-05_Field Photos of Associated Plant Hosts of Viruses

Additional-File-06_Phylogenetic-Analyses-Trees

Additonal-File-07_Data-Source_Nucleotide-Diversity

Additonal-File-08_Data-Source_Pairwise-Similarity-Scores

Additonal-File-09_Data-Source_InterPro,Pfam-Domain-Searches-Genome-annotation

Additonal-File-10_Supp-Figure-5_RT-PCR-tests-HTS-PTV1

## Acknowledgements

The authors would like to thank Zala Kogej, Živa Lengar, Meta Ješelnik, Lija Fajdiga, Miha Kitek, and Anja Cerovšek for their help in sampling and sample processing, Živa Ramšak and Henrik Krnec for helping in setting up bioinformatics programs used in viroid discovery, and Magda Tušek Žnidarič for guidance in the electron microscopy experiment. Finally, the authors would like to thank Benjamin Lee and Uri Neri for insightful discussions on viroid discovery, and the Plant Virus Ecology Journal Club led by Carolyn Malmstrom for the informative and stimulating discussions on virology that contributed useful insights to this study.

## Availability of data and material

Raw sequencing reads were submitted in the NCBI Sequence Read Archive (SRA), with the BioProject accession number PRJNA772045. All sample and sequencing metadata, and SRA accession numbers are in Supplementary Tables 1-3. All data related to virus genome assembly and identification, and their NCBI GenBank accession numbers are in Supplementary Tables 4-6. Description of other supplementary material are provided in the ‘Supplementary information’ section of this paper. High quality version of tables and figures and supplementary files are available in *Figshare* (link: https://doi.org/10.6084/m9.figshare.20200769).

## Ethics approval and consent to participate

Not applicable.

## Consent for publication

Not applicable.

