## Additional-File-03_Genome Organization for "In-depth study of tomato and weed viromes reveals undiscovered plant virus diversity in an agroecosystem"

### SUPPLEMENTARY INFORMATION

### Additonal File 03

**Supplementary Figure 1.** Genome organization of novel viruses or first full genomes of known viruses, showing known and putative open reading frames and the protein it codes for, and predicted secondary structures of selected viroid-like circular RNAs detected in this study.

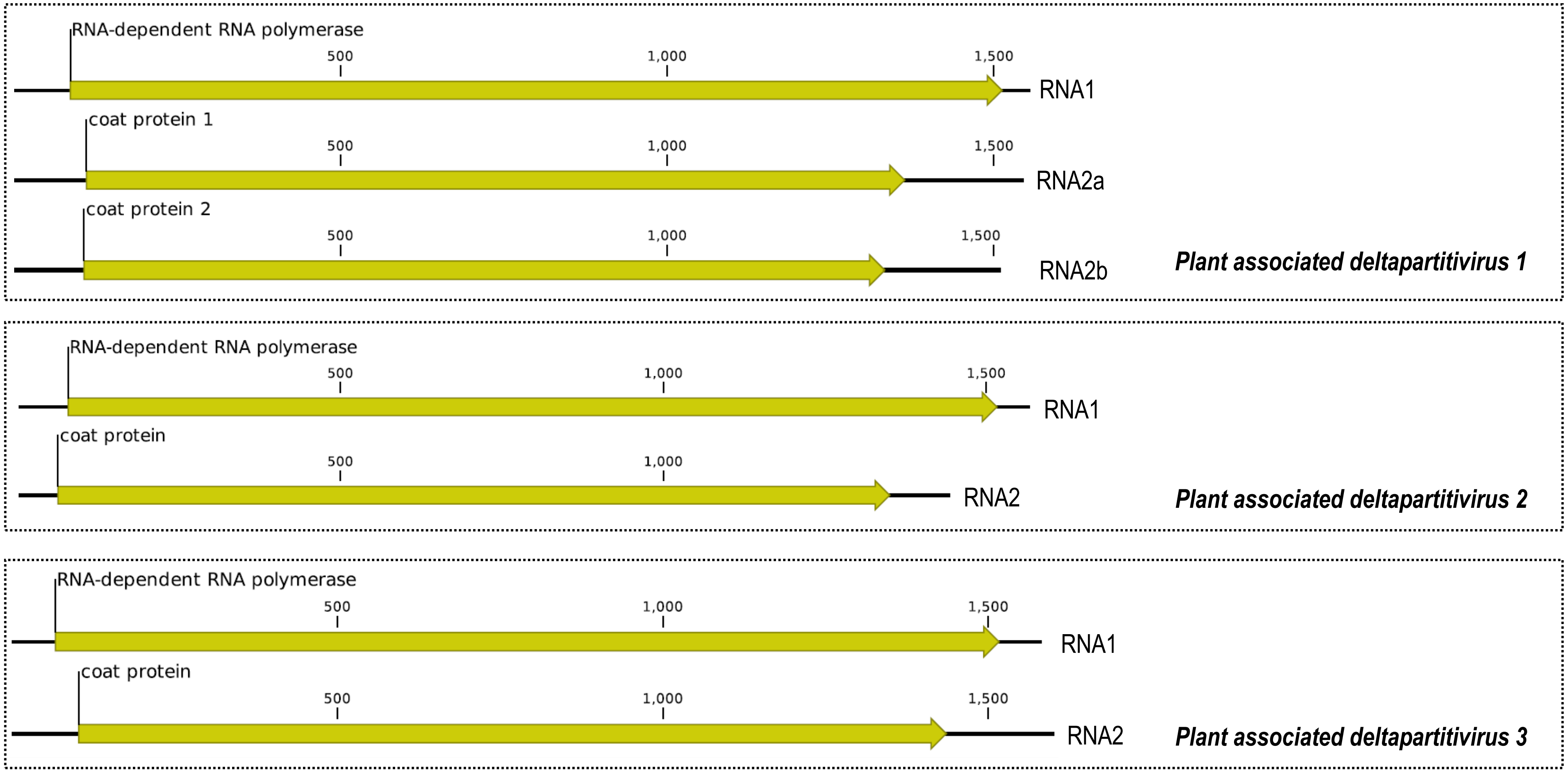


**Supplementary Figure 1-01.** Genomes of new virus species discovered under family *Partitiviridae,* order *Durnavirales.*

**Supplementary Figure 1.** Genome organization of novel viruses or first full genomes of known viruses, showing known and putative open reading frames and the protein it codes for, and predicted secondary structures of selected viroid-like circular RNAs detected in this study. **Note**: Genome length in number of bases are shown with a scale. For full information on genome length, protein domains, *etc.,* please refer to Supplementary Table 5, and the corresponding accession in GenBank.

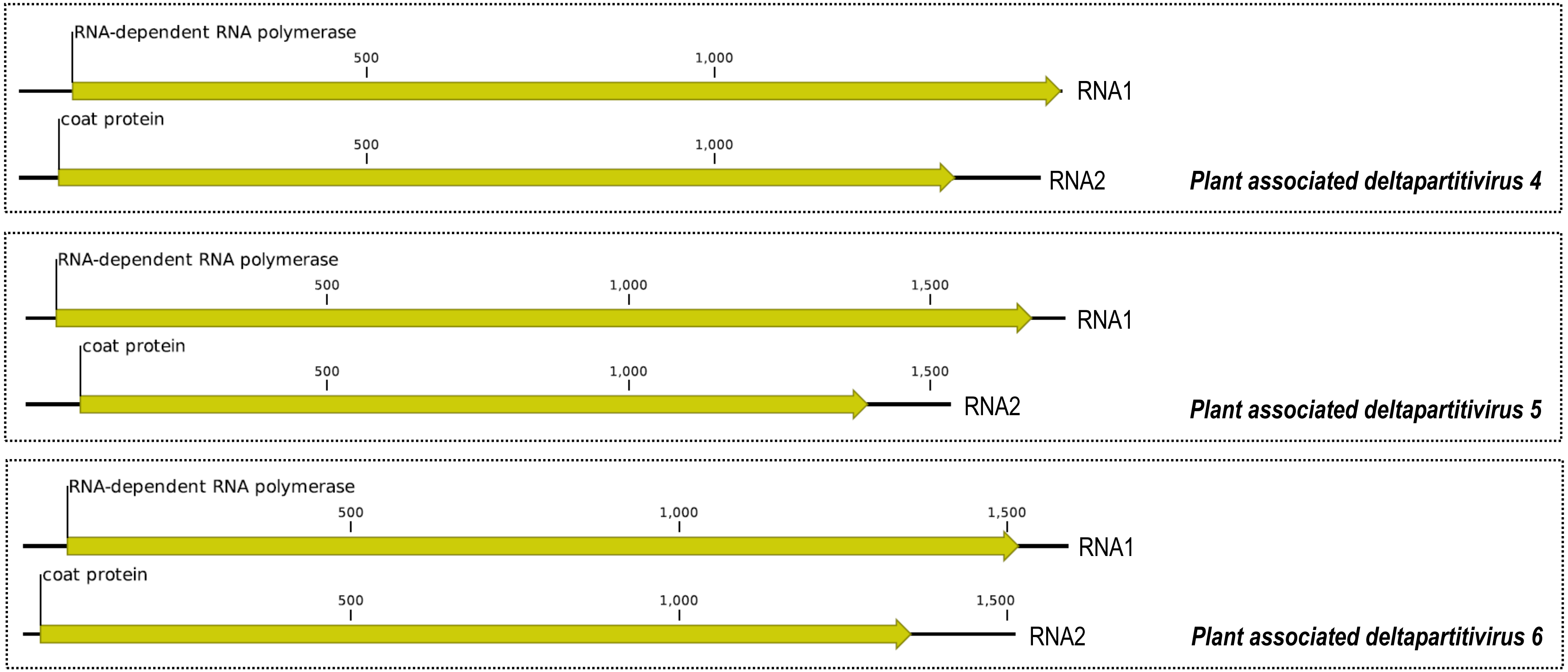


**Supplementary Figure 1-01** (continued)**.** Genomes of new virus species discovered under family *Partitiviridae,* order *Durnavirales.*

**Supplementary Figure 1** (continued)**.** Genome organization of novel viruses or first full genomes of known viruses, showing known and putative open reading frames and the protein it codes for, and predicted secondary structures of selected viroid-like circular RNAs detected in this study. **Note**: Genome length in number of bases are shown with a scale. For full information on genome length, protein domains, *etc.,* please refer to Supplementary Table 5, and the corresponding accession in GenBank.

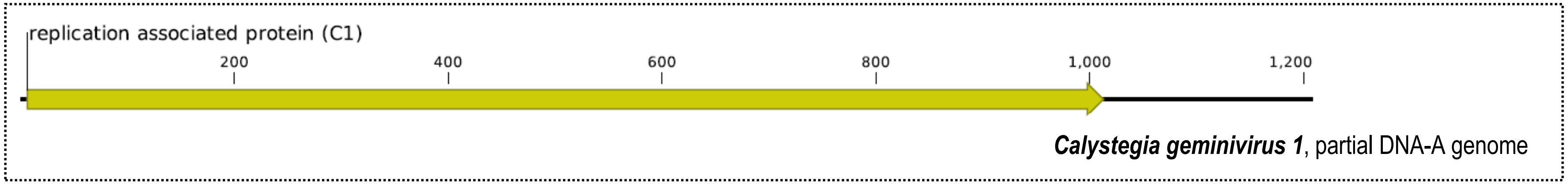


**Supplementary Figure 1-02.** Genomes of new virus species discovered under *Geminiviridae* (order *Geplafuvirales*).

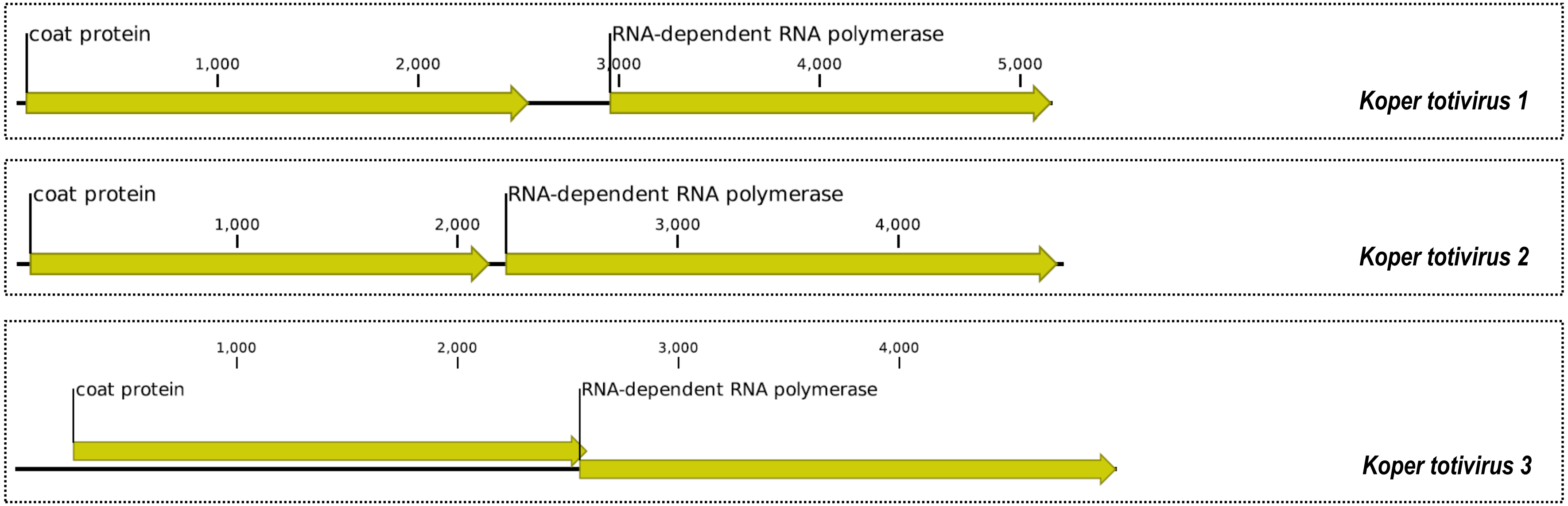


**Supplementary Figure 1-03.** Genomes of new virus species discovered under *Totiviridae* (order *Ghabrivirales*).

**Supplementary Figure 1** (continued)**.** Genome organization of novel viruses or first full genomes of known viruses, showing known and putative open reading frames and the protein it codes for, and predicted secondary structures of selected viroid-like circular RNAs detected in this study. **Note**: Genome length in number of bases are shown with a scale. For full information on genome length, protein domains, *etc.*, please refer to Supplementary Table 5, and the corresponding accession in GenBank.

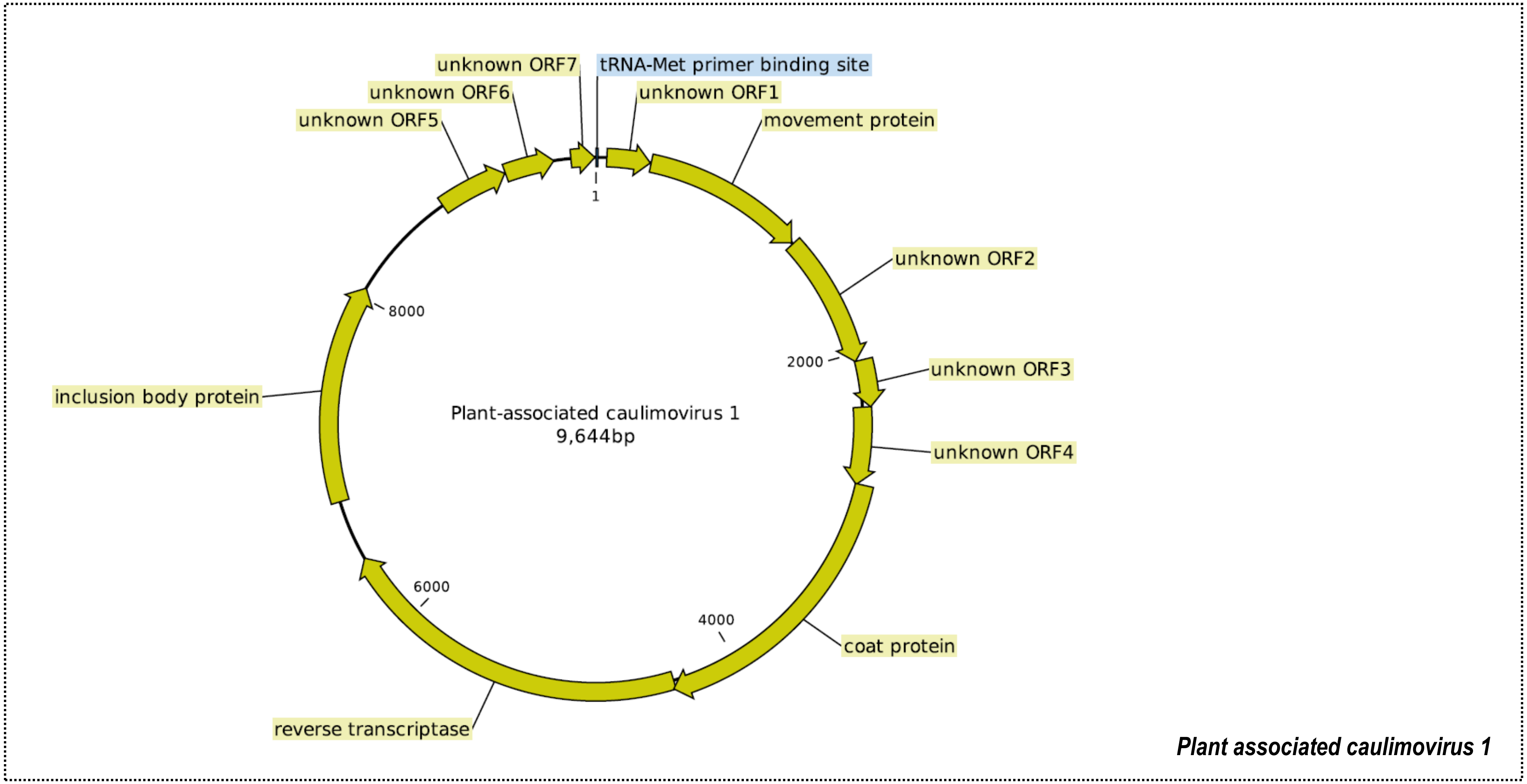


**Supplementary Figure 1-04.** Genomes of new virus species discovered under family *Caulimoviridae* (order *Ortelivirales*).

**Supplementary Figure 1** (continued)**.** Genome organization of novel viruses or first full genomes of known viruses, showing known and putative open reading frames and the protein it codes for, and predicted secondary structures of selected viroid-like circular RNAs detected in this study. **Note**: Genome length in number of bases are shown with a scale. For full information on genome length, protein domains, *etc.,* please refer to Supplementary Table 5, and the corresponding accession in GenBank.

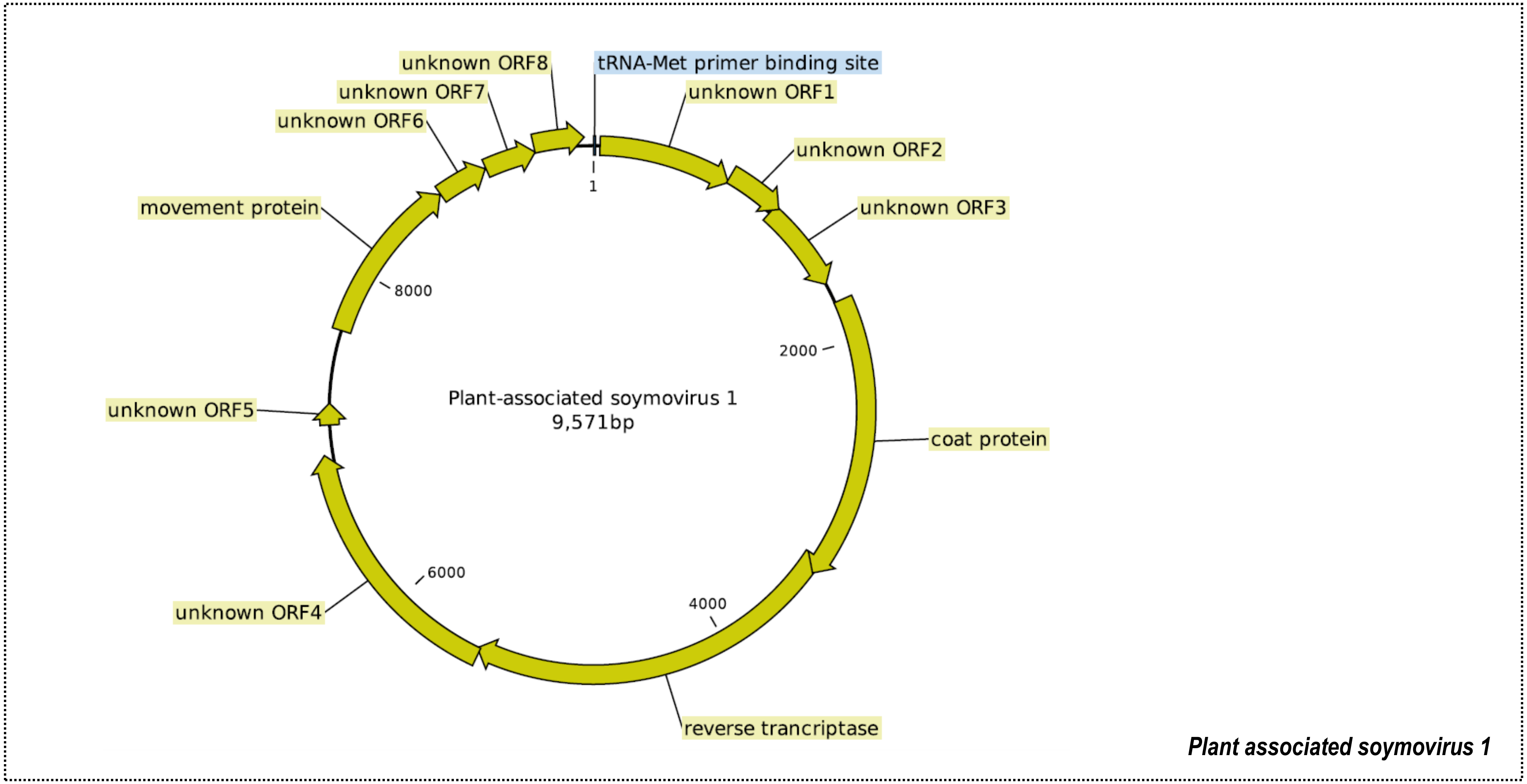


**Supplementary Figure 1-04** (continued)**.** Genomes of new virus species discovered under family *Caulimoviridae* (order *Ortelivirales*).

**Supplementary Figure 1** (continued)**.** Genome organization of novel viruses or first full genomes of known viruses, showing known and putative open reading frames and the protein it codes for, and predicted secondary structures of selected viroid-like circular RNAs detected in this study. **Note**: Genome length in number of bases are shown with a scale. For full information on genome length, protein domains, *etc.,* please refer to Supplementary Table 5, and the corresponding accession in GenBank.

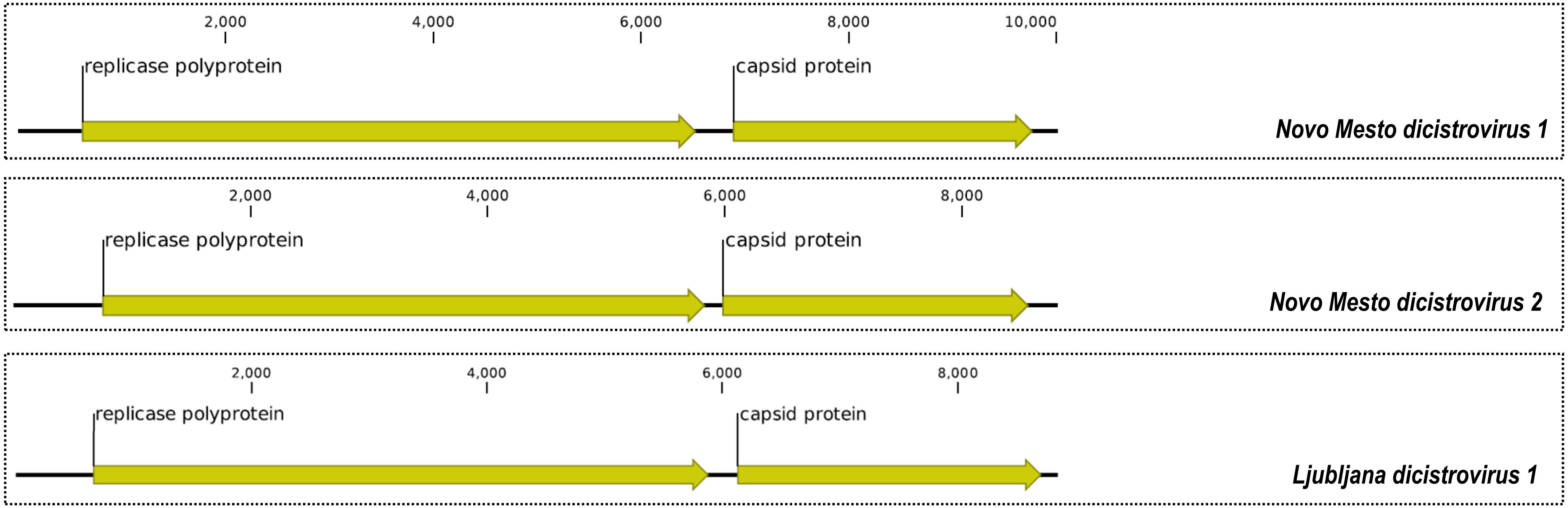


**Supplementary Figure 1-05.** Genomes of new virus species discovered under family *Dicistroviridae* (order *Picornavirales*).

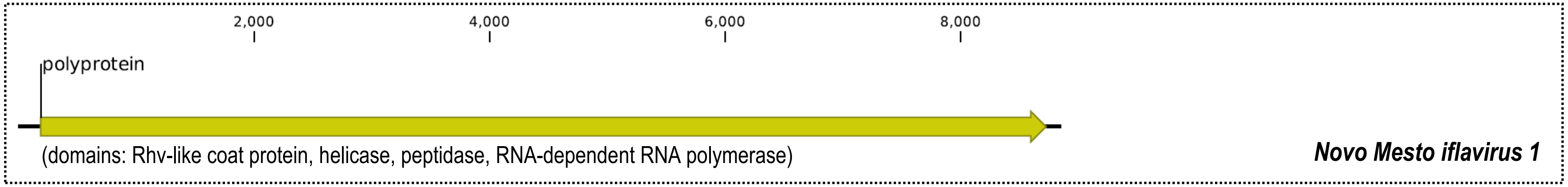


**Supplementary Figure 1-06.** Genomes of new virus species discovered under family *Iflaviridae* (order *Picornavirales*).

**Supplementary Figure 1** (continued)**.** Genome organization of novel viruses or first full genomes of known viruses, showing known and putative open reading frames and the protein it codes for, and predicted secondary structures of selected viroid-like circular RNAs detected in this study. **Note**: Genome length in number of bases are shown with a scale. For full information on genome length, protein domains, *etc.,* please refer to Supplementary Table 5, and the corresponding accession in GenBank.

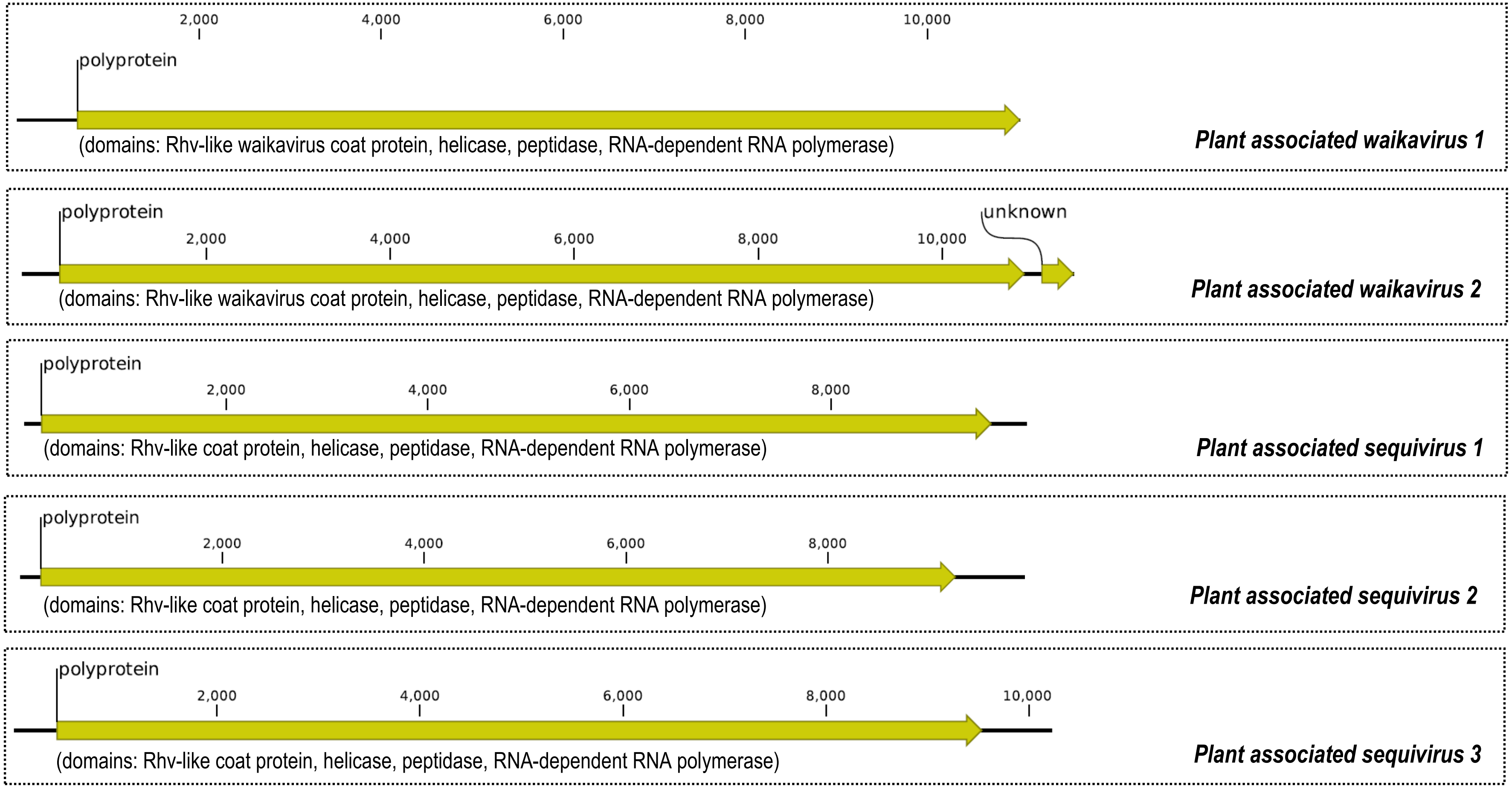


**Supplementary Figure 1-07.** Genomes of new virus species discovered under family *Secoviridae* (order *Picornavirales*).

**Supplementary Figure 1** (continued)**.** Genome organization of novel viruses or first full genomes of known viruses, showing known and putative open reading frames and the protein it codes for, and predicted secondary structures of selected viroid-like circular RNAs detected in this study. **Note**: Genome length in number of bases are shown with a scale. For full information on genome length, protein domains, *etc.,* please refer to Supplementary Table 5, and the corresponding accession in GenBank.

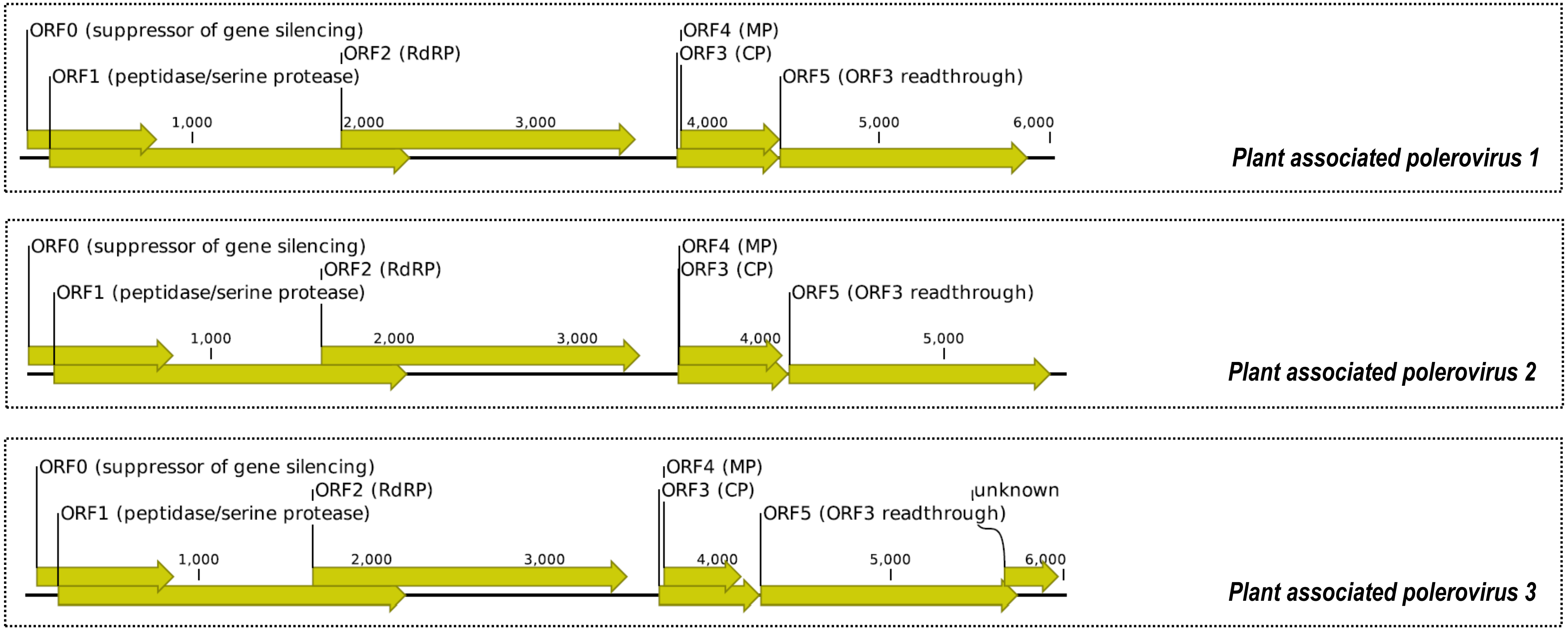


**Supplementary Figure 1-08.** Genomes of new virus species discovered under genus *Polerovirus,* family *Solemoviridae* (order *Sobelivirales*)*.* **Note:** RdRP - RNA-dependent RNA polymerase, MP - movement protein, CP - coat protein.

**Supplementary Figure 1** (continued)**.** Genome organization of novel viruses or first full genomes of known viruses, showing known and putative open reading frames and the protein it codes for, and predicted secondary structures of selected viroid-like circular RNAs detected in this study. **Note**: Genome length in number of bases are shown with a scale. For full information on genome length, protein domains, *etc.,* please refer to Supplementary Table 5, and the corresponding accession in GenBank.

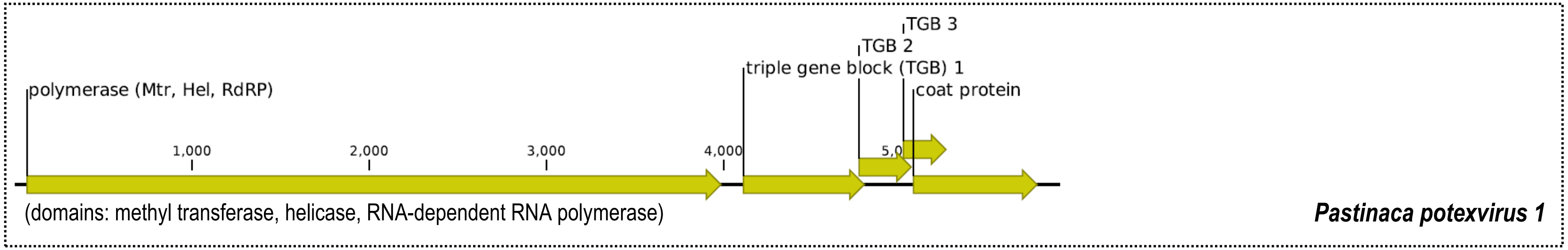


**Supplementary Figure 1-09.** Genome of new virus species discovered under genus *Potexvirus,* family *Alphaflexiviridae* (order *Tymovirales*).

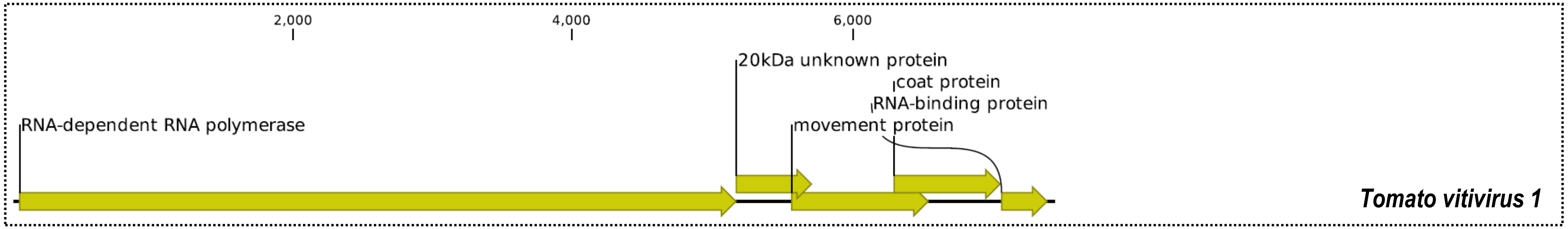


**Supplementary Figure 1-10.** Genome of new virus species discovered under genus *Vitivirus,* family *Betaflexiviridae* (order *Tymovirales*).

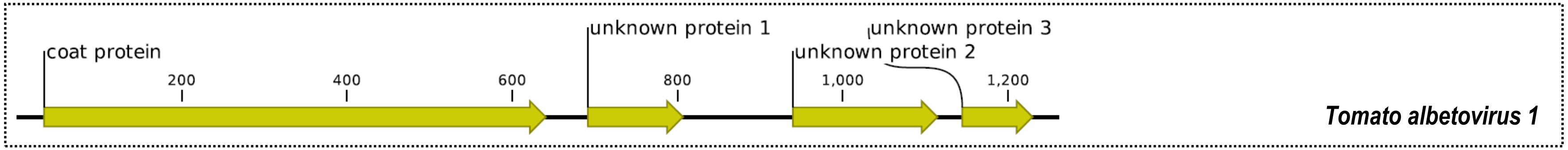


**Supplementary Figure 1-11.** Genome of new virus species discovered under genus *Albetovirus,* under unclassified satellite viruses.

**Supplementary Figure 1** (continued)**.** Genome organization of novel viruses or first full genomes of known viruses, showing known and putative open reading frames and the protein it codes for, and predicted secondary structures of selected viroid-like circular RNAs detected in this study. **Note**: Genome length in number of bases are shown with a scale. For full information on genome length, protein domains, *etc.,* please refer to Supplementary Table 5, and the corresponding accession in GenBank.

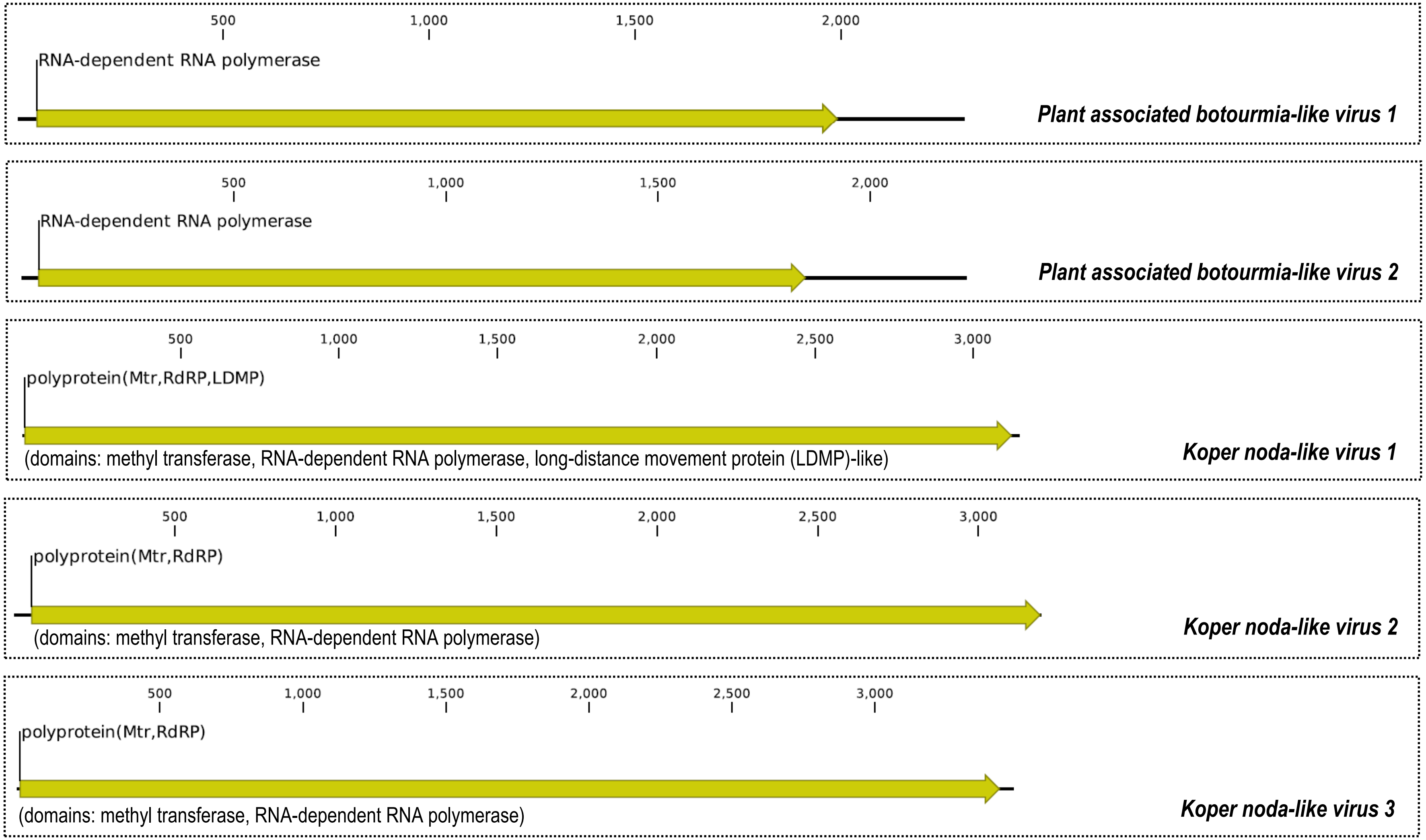


**Supplementary Figure 1-12.** Genome of new but unclassified virus species discovered under realm *Riboviria.*

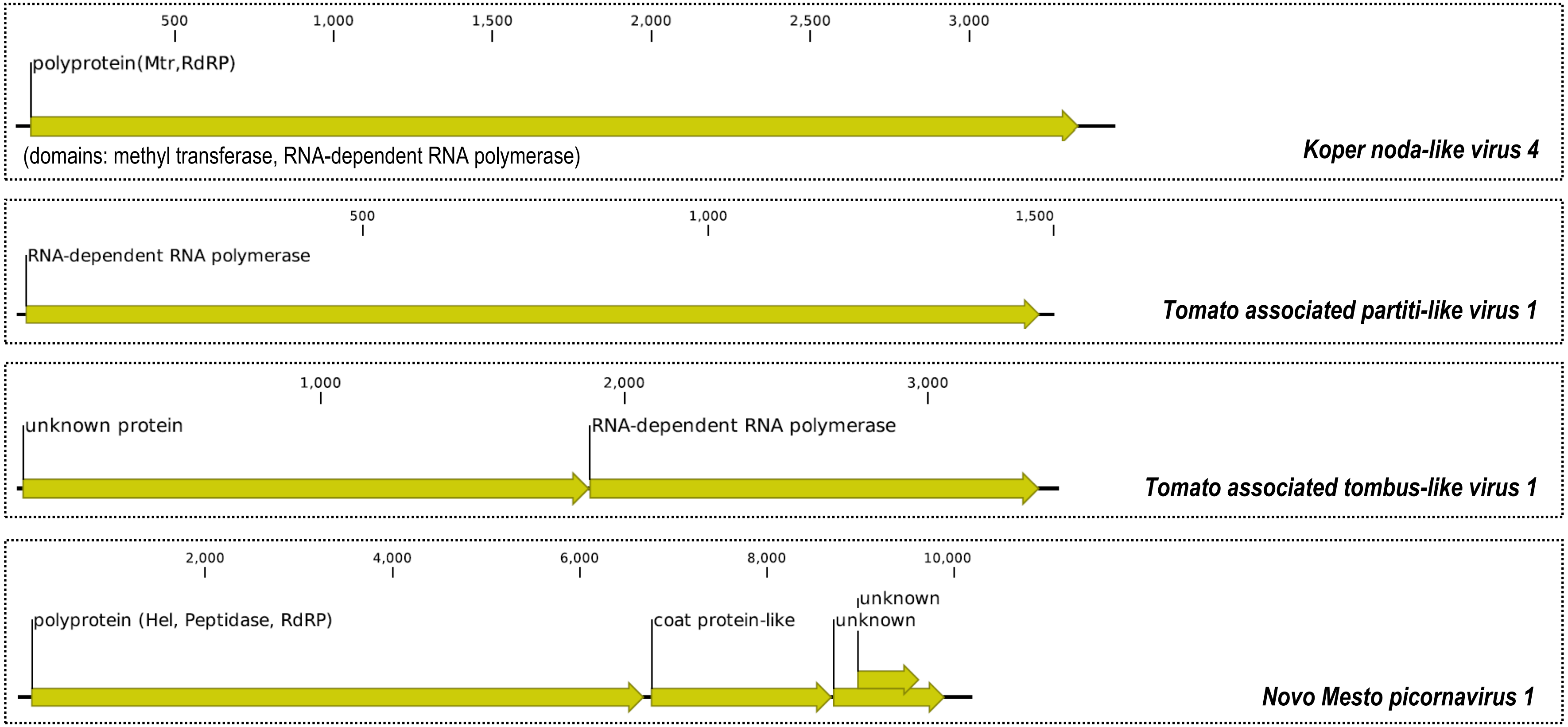


**Supplementary Figure 1-12** (continued)**.** Genome of new but unclassified virus species discovered under Realm *Riboviria.*

**Supplementary Figure 1** (continued)**.** Genome organization of novel viruses or first full genomes of known viruses, showing known and putative open reading frames and the protein it codes for, and predicted secondary structures of selected viroid-like circular RNAs detected in this study. **Note**: Genome length in number of bases are shown with a scale. For full information on genome length, protein domains, *etc.,* please refer to Supplementary Table 5, and the corresponding accession in GenBank.

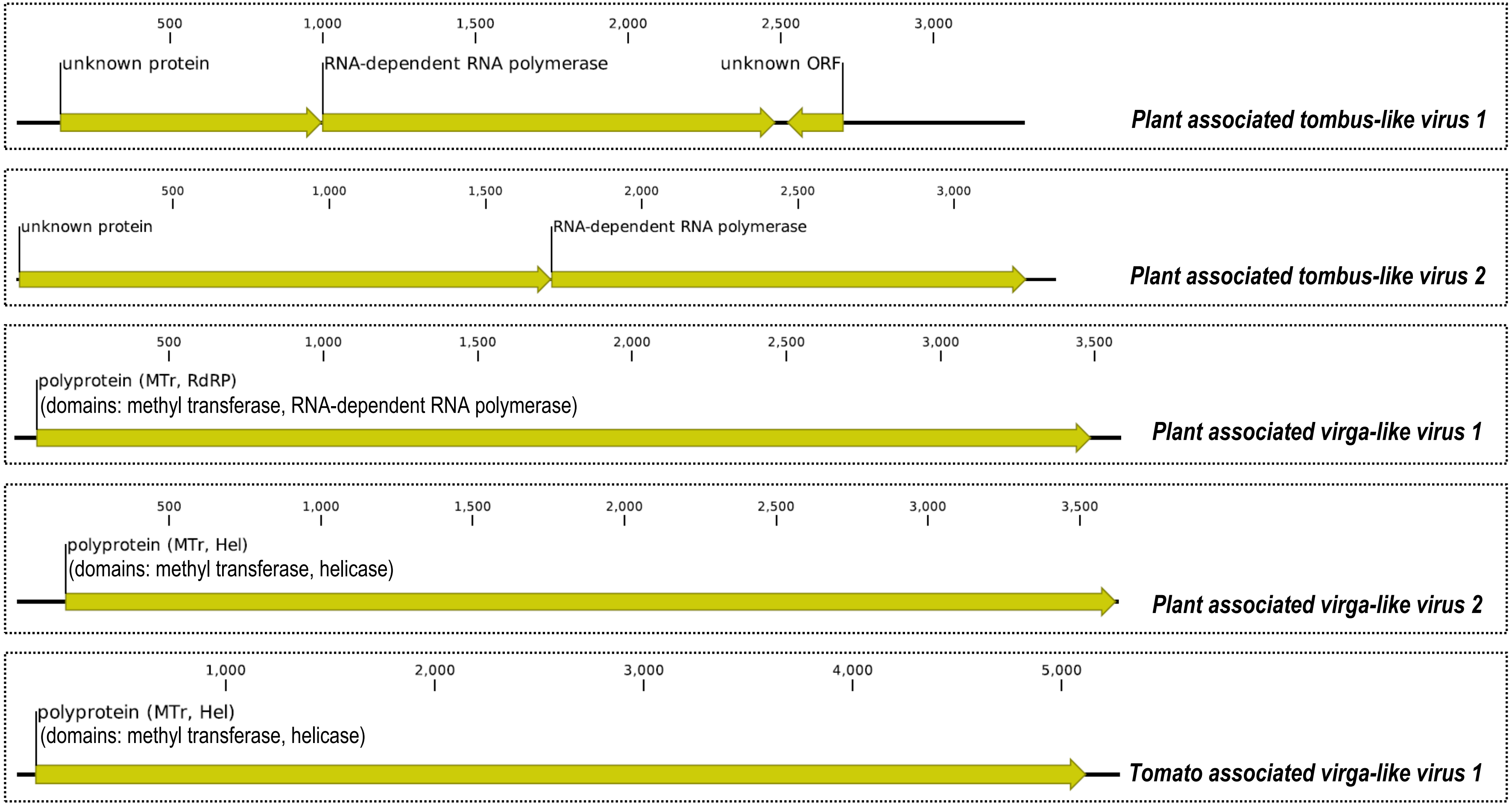


**Supplementary Figure 1-12** (continued)**.** Genome of new but unclassified virus species discovered under Realm *Riboviria.*

**Supplementary Figure 1** (continued)**.** Genome organization of novel viruses or first full genomes of known viruses, showing known and putative open reading frames and the protein it codes for, and predicted secondary structures of selected viroid-like circular RNAs detected in this study. **Note**: Genome length in number of bases are shown with a scale. For full information on genome length, protein domains, *etc.,* please refer to Supplementary Table 5, and the corresponding accession in GenBank.

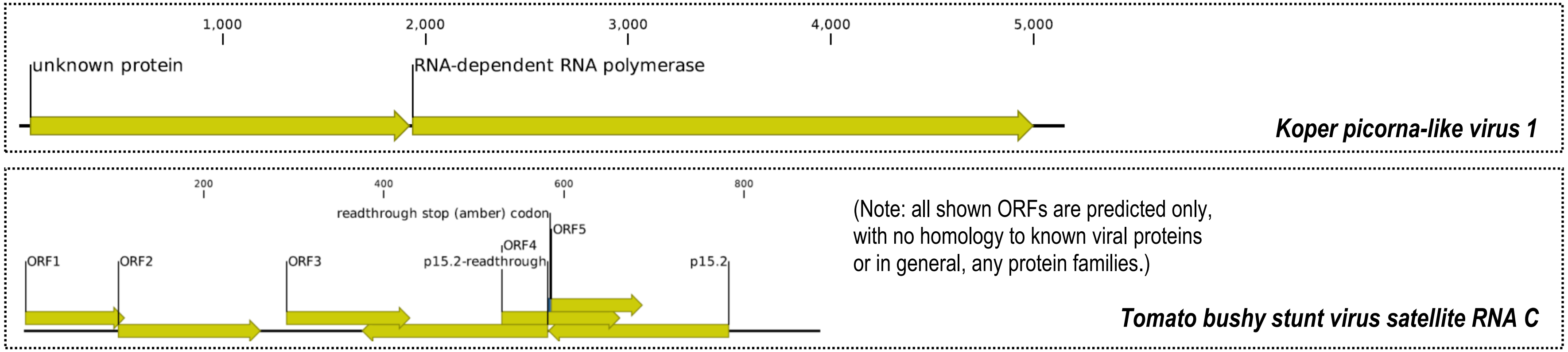


**Supplementary Figure 1-12** (continued)**.** Genome of new but unclassified virus species discovered under Realm *Riboviria.*

**Supplementary Figure 1** (continued)**.** Genome organization of novel viruses or first full genomes of known viruses, showing known and putative open reading frames and the protein it codes for, and predicted secondary structures of selected viroid-like circular RNAs detected in this study. **Note**: Genome length in number of bases are shown with a scale. For full information on genome length, protein domains, *etc.,* please refer to Supplementary Table 5, and the corresponding accession in GenBank.

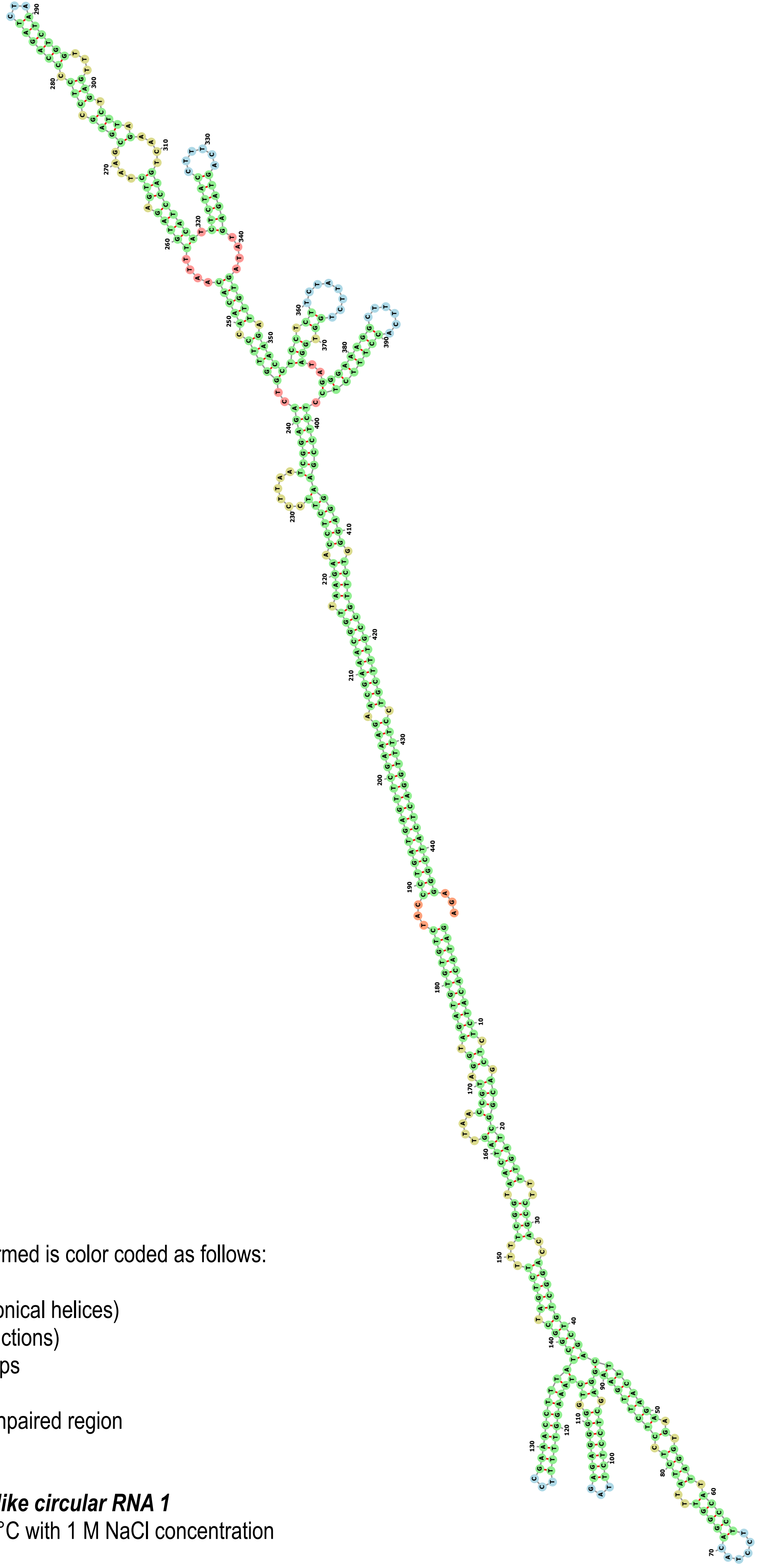

Type of structure formed is color coded as follows:

**Green**: Stems (canonical helices)
**Red**: Multiloops (junctions)
**Yellow**: Interior Loops
**Blue**: Hairpin loops
**Orange**: 5' and 3' unpaired region

***Taraxacum viroid-like circular RNA 1***
dG = -194.35, at 37°C with 1 M NaCl concentration

**Supplementary Figure 1-13**. Predicted secondary structures of selected viroid-like circular RNAs detected in this study.

**Supplementary Figure 1** (continued)**.** Genome organization of novel viruses, or first full genomes of known viruses showing known and putative open reading frames and the protein it codes for, and predicted secondary stuctures of viroid-like circular RNAs detected in this study. **Note**: Genome length in number of bases are shown with a scale. For full information on genome length, protein domains, etc., please refer to Supplementary Table 5 (of the Supplementary Information), and the corresponding accession in GenBank.

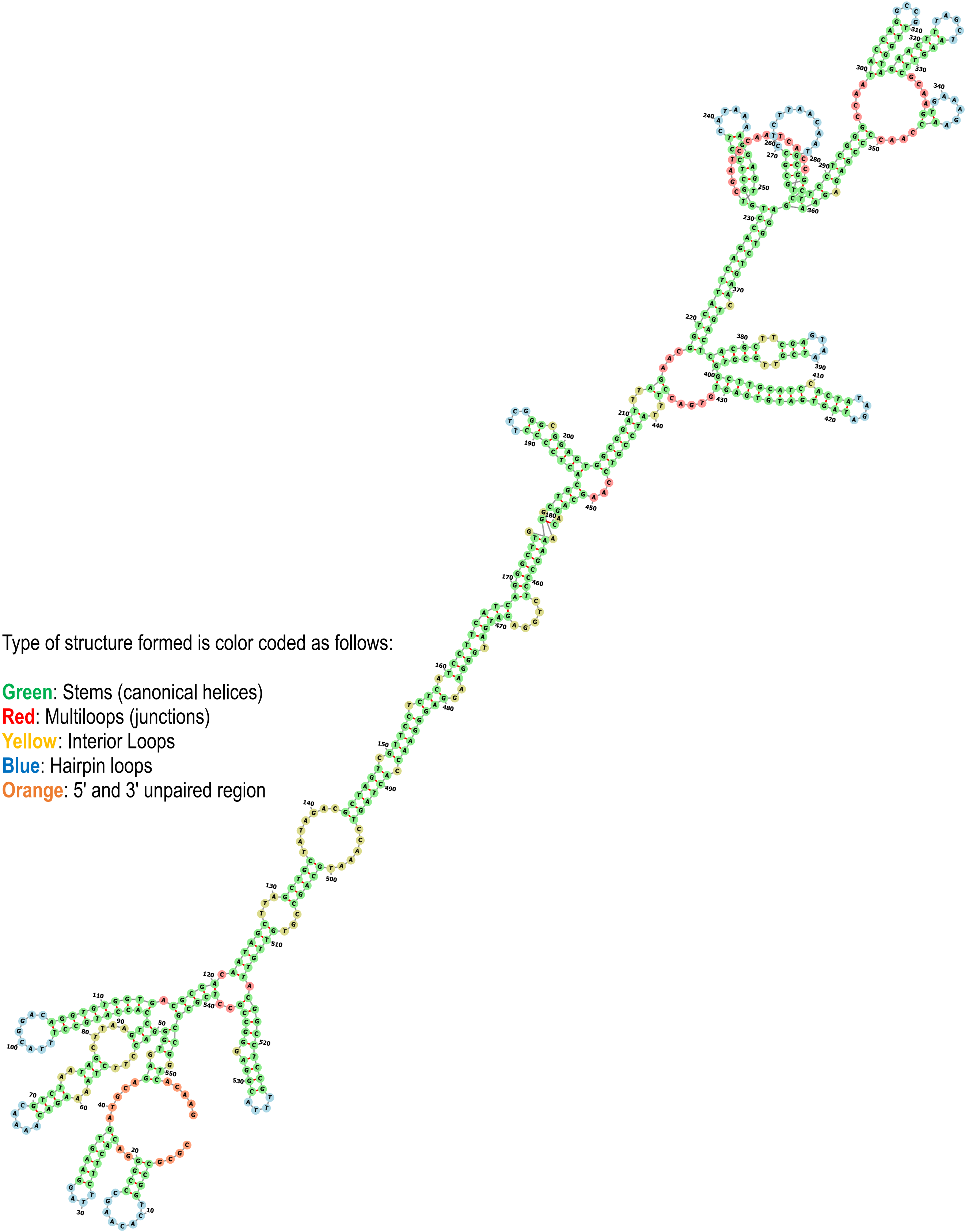

Type of structure formed is color coded as follows:

**Green**: Stems (canonical helices)
**Red**: Multiloops (junctions)
**Yellow**: Interior Loops
**Blue**: Hairpin loops
**Orange**: 5' and 3' unpaired region

***Plant associated viroid-like circular RNA 1*** dG = -181.90, at 37°C with 1 M NaCl concentration

**Supplementary Figure 1-13** (continued). Predicted secondary structures of selected viroid-like circular RNAs detected in this study.

**Supplementary Figure 1** (continued)**.** Genome organization of novel viruses, or first full genomes of known viruses showing known and putative open reading frames and the protein it codes for, and predicted secondary stuctures of viroid-like circular RNAs detected in this study. **Note**: Genome length in number of bases are shown with a scale. For full information on genome length, protein domains, etc., please refer to Supplementary Table 5 (of the Supplementary Information), and the corresponding accession in GenBank.

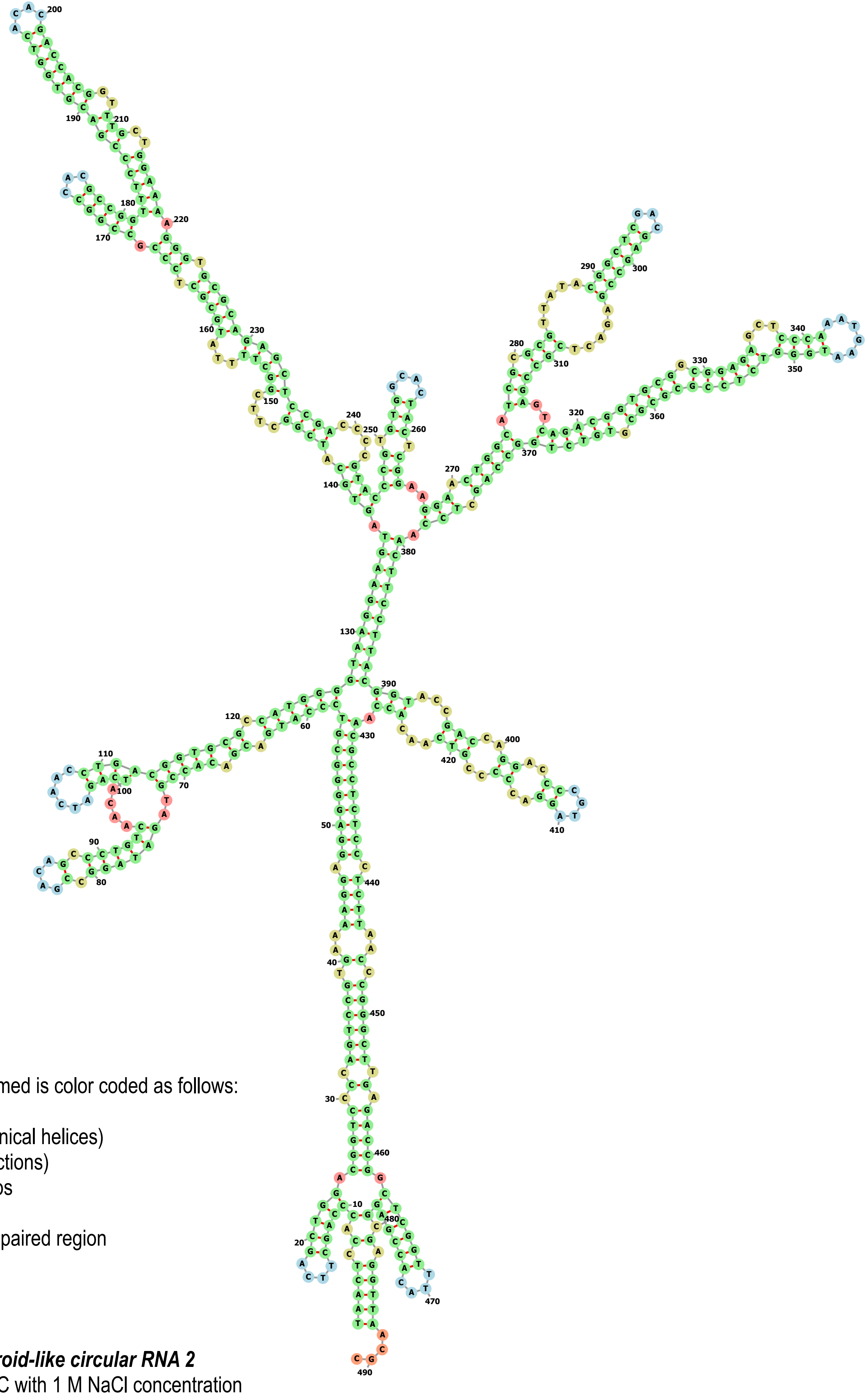

Type of structure formed is color coded as follows:

**Green**: Stems (canonical helices)
**Red**: Multiloops (junctions)
**Yellow**: Interior Loops
**Blue**: Hairpin loops
**Orange**: 5' and 3' unpaired region

***Plant associated viroid-like circular RNA 2***
dG = -247.60, at 37°C with 1 M NaCl concentration

**Supplementary Figure 1-13** (continued). Predicted secondary structures of selected viroid-like circular RNAs detected in this study.

**Supplementary Figure 1** (continued)**.** Genome organization of novel viruses, or first full genomes of known viruses showing known and putative open reading frames and the protein it codes for, and predicted secondary stuctures of viroid-like circular RNAs detected in this study. **Note**: Genome length in number of bases are shown with a scale. For full information on genome length, protein domains, etc., please refer to Supplementary Table 5 (of the Supplementary Information), and the corresponding accession in GenBank.

Type of structure formed is color coded as follows:

**Green**: Stems (canonical helices)
**Red**: Multiloops (junctions)
**Yellow**: Interior Loops
**Blue**: Hairpin loops
**Orange**: 5' and 3' unpaired region

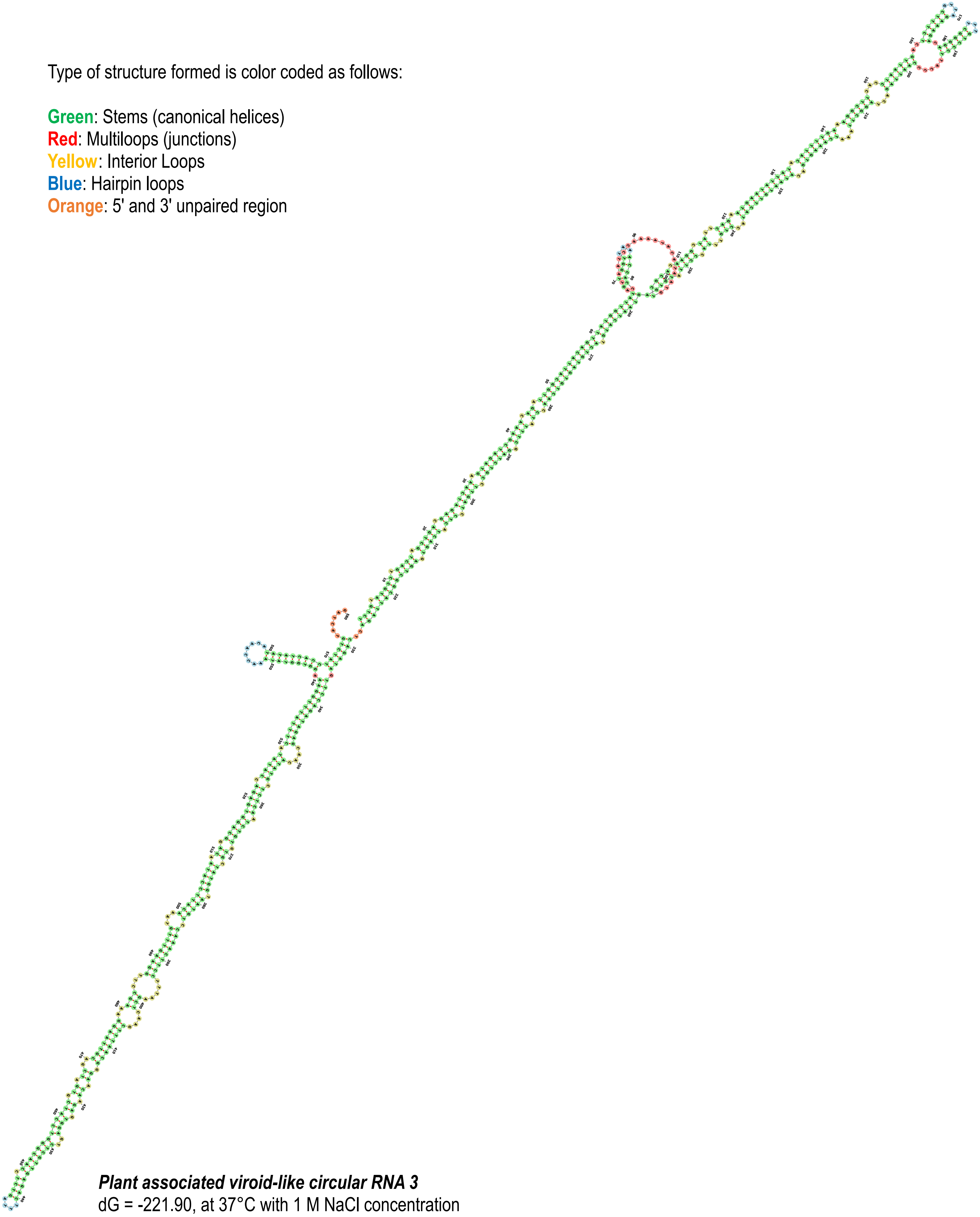

***Plant associated viroid-like circular RNA 3***
dG = -221.90, at 37°C with 1 M NaCl concentration

**Supplementary Figure 1-13** (continued). Predicted secondary structures of selected viroid-like circular RNAs detected in this study.

**Supplementary Figure 1** (continued**)**. Genome organization of novel viruses, or first full genomes of known viruses showing known and putative open reading frames and the protein it codes for, and predicted secondary stuctures of viroid-like circular RNAs detected in this study. **Note**: Genome length in number of bases are shown with a scale. For full information on genome length, protein domains, etc., please refer to Supplementary Table 5 (of the Supplementary Information), and the corresponding accession in GenBank.

Type of structure formed is color coded as follows:

**Green**: Stems (canonical helices)
**Red**: Multiloops (junctions)
**Yellow**: Interior Loops
**Blue**: Hairpin loops
**Orange**: 5' and 3' unpaired region

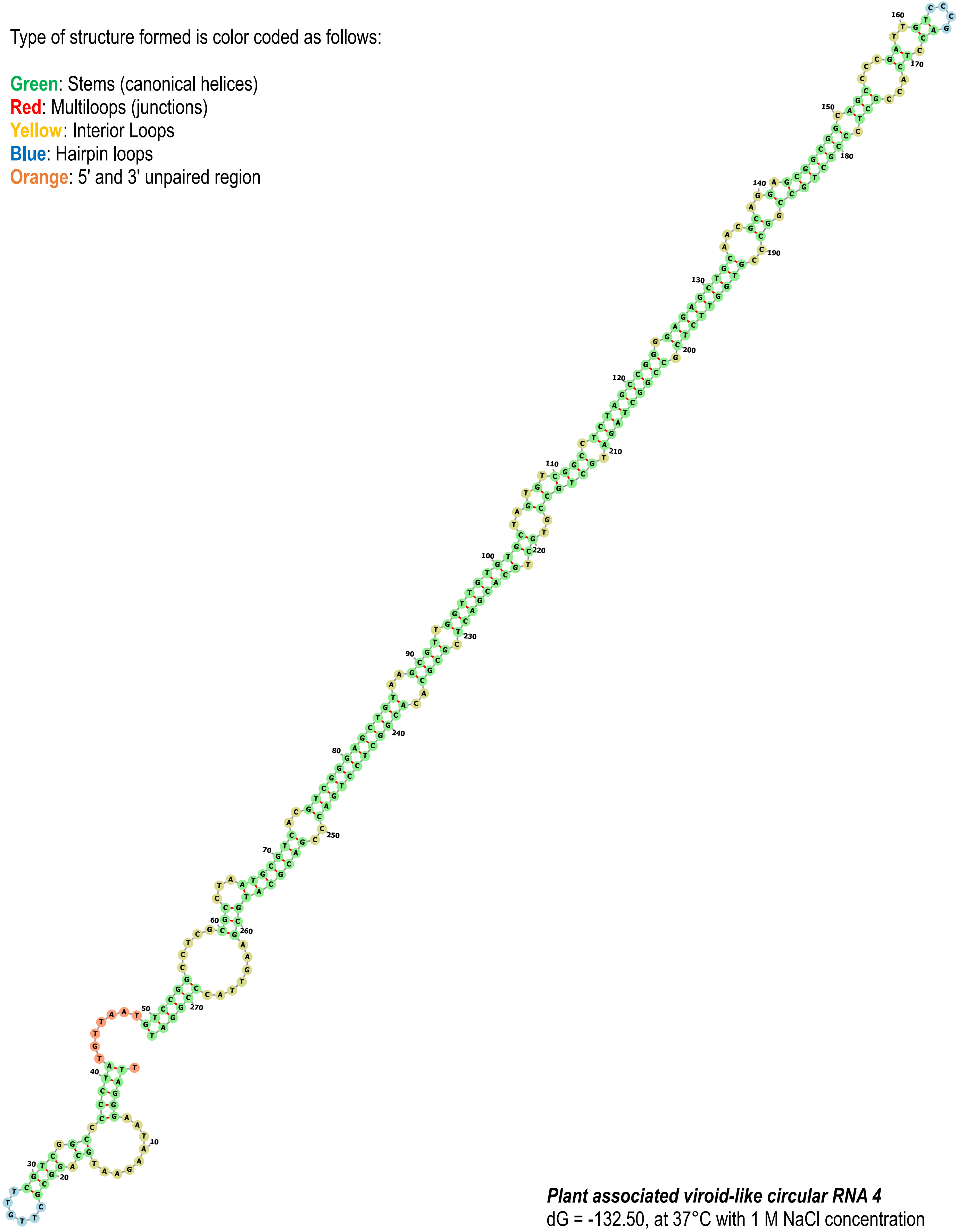

***Plant associated viroid-like circular RNA 4***
dG = -132.50, at 37°C with 1 M NaCl concentration

**Supplementary Figure 1-13** (continued). Predicted secondary structures of selected viroid-like circular RNAs detected in this study.

**Supplementary Figure 1** (continued)**.** Genome organization of novel viruses, or first full genomes of known viruses showing known and putative open reading frames and the protein it codes for, and predicted secondary stuctures of viroid-like circular RNAs detected in this study. **Note**: Genome length in number of bases are shown with a scale. For full information on genome length, protein domains, etc., please refer to Supplementary Table 5 (of the Supplementary Information), and the corresponding accession in GenBank.
