## Additional-File-04_RT-PCR Confirmations of Selected Viruses for "In-depth study of tomato and weed viromes reveals undiscovered plant virus diversity in an agroecosystem"

### SUPPLEMENTARY INFORMATION

### Additional File 04

**Supplementary Table 7.** RT-PCR primers and PCR conditions used in confirmation of associated plant hosts of selected viruses and putative viroids. **Supplementary Table 8.** RT-PCR thermocycling conditions used in the detection of selected viruses and putative viroids in associated plant hosts. **Supplementary Figure 2.** Orientation and annealing sites of the primers designed for the amplification of the circular genome of Taraxacum viroid-like circular RNA 1, and results of RT-PCR confirmation of the circular genome. **Supplementary Table 9**. List of confirmed associated plant hosts of selected viruses and putative viroids.

**Supplementary Table 7.** RT-PCR primers and PCR conditions used in confirmation of associated plant hosts of selected viruses and a putative viroid. Note: GenBank accession numbers of the species indicated above are in Supplementary Table 5.

| No. | Target virus name | Primer ID | Strand | Sequence (5'→3') | Length (bases) | Target genome region (start, end position / gene(s)) | | | Annealing tempera-ture. (°C) | Amplicon size (bases) |
|---|---|---|---|---|---|---|---|---|---|---|
| | | | | | | Start | End | Gene | | |
| 01 | Artemesia fimovirus 1 | NFimo1-F<br>NFimo1-R | forward<br>reverse | ACATGGAAGTGACAGGTTCTCC<br>ATAGTTGACGCCTACCCATTGC | 22<br>22 | 5889 | 6435 | RdRP | 57 | 547 |
| 02 | Calystegia geminivirus 1 | NGemi1-F<br>NGemi1-R | forward<br>reverse | CAGAACCTCAACCTTCCATTCC<br>CCAATCAGCTCTTTCCAGTGC | 22<br>21 | 248 | 817 | replicase | 57 | 570 |
| 03 | Plantago potyvirus 1 | NPoty1-F<br>NPoty1-R | forward<br>reverse | AGGCTAGAGATCGCAAACTTGG<br>CAACTTTATCCGTGCTCTGTGG | 22<br>22 | 5687 | 6550 | polyprotein | 57 | 864 |
| 04 | Mentha macluravirus 1 | NPoty2-F<br>NPoty2-R | forward<br>reverse | ACATACGGCTCAGCTTTCTTCC<br>GGCACAGAGAACTTCAACATCG | 22<br>22 | 2474 | 3306 | polyprotein | 57 | 833 |
| 05 | Rumex potyvirus 1 | NPoty3-F<br>NPoty3-R | forward<br>reverse | GTTGACCTAACCCCTCACAACC<br>CACTACTGGTAGCCCACACTGC | 22<br>22 | 7098 | 7733 | polyprotein | 57 | 636 |
| 06 | broad-leafed dock virus A, isolate 2 | NPoty4-F<br>NPoty4-R | forward<br>reverse | GGATGAGGAATATGGAGCTTGG<br>TACTAGGCGGTGGAAGAAAACC | 22<br>22 | 7502 | 8178 | polyprotein | 57 | 677 |
| 07 | Pastinaca umbravirus 1 | NTomb1-F<br>NTomb1-R | forward<br>reverse | GCGAGACTGTCTGTACCACTGC<br>CAGTCGTACCCCCTCTAACTGG | 22<br>22 | 2504 | 3354 | RdRP-MP | 57 | 851 |
| 08 | Picris umbravirus 1 | NTomb2-F<br>NTomb2-R | forward<br>reverse | TCGTGCTGTATAGGGTTCATGG<br>CAAACTCCCCAAATGGACTACC | 22<br>22 | 1846 | 2839 | RdRP | 57 | 994 |
| 09 | Convolvulus aureusvirus 1 | NTomb3-F<br>NTomb3-R | forward<br>reverse | GAAAGTCAGCCGAATTGTAGGG<br>CAATCCAAGTTGGTCCTTCTCC | 22<br>22 | 1088 | 1814 | RdRP | 57 | 727 |
| 10 | Calystegia pelarspovirus 1 | NTomb4-F<br>NTomb4-R | forward<br>reverse | GCCGAATGAGGGTATGTTTAGG<br>GTTCCGCTGTTCTTCAACTGG | 22<br>21 | 1019 | 1594 | RdRP | 57 | 576 |
| 11 | Cichorium alphacarmovirus 1 | NTomb6-F<br>NTomb6-R | forward<br>reverse | GGAGAAACATGAGGAACGAACC<br>AGAAAACCCTTTCCAGGAGACC | 22<br>22 | 322 | 989 | RdRP | 57 | 668 |
| 12 | Pastinaca potexvirus 1 | NAflex1-F<br>NAflex1-R | forward<br>reverse | CACAGTGGATGAGGATGTAGCC<br>TTCGTAAGCTGAGCTGAGTTGG | 22<br>22 | 3565 | 4542 | polypro-tein-TGB | 57 | 978 |
| 13 | plant associated tobamo-like virus 1 | NVirga1-F<br>NVirga1-R | forward<br>reverse | CTTCACCTGTCTCAGTGAGGAC<br>TATGAGTTGCGATGGGTAGACG | 22<br>22 | 115 | 522 | replicase | 55 | 408 |
| 14 | Plantago tobamovirus 1 | NVirga2-F<br>NVirga2-R | forward<br>reverse | AACGCACTATCCGAGCTATCTG<br>TACAGCCACCCTAAACCATGTC | 22<br>22 | 1720 | 2421 | replicase | 55 | 702 |

| | | | | | | | | | | |
|---|---|---|---|---|---|---|---|---|---|---|
| 15 | Mercurialis orthotospovirus 1 | NTospo1-F<br>NTospo1-R | forward<br>reverse | TAGAGCCGAAGATGTTGTGGAC<br>GTCAGCGACCATTAAGCCTTTG | 22<br>22 | 1661 | 2291 | L gene (RdRP) | 55 | 631 |
| 16 | tomato associated bunya-like virus 1 | NTospo2-F<br>NTospo2-R | forward<br>reverse | CGAAAGGAGGCGATAGTGATGC<br>CTGCGTCATCCCTACCTGATAC | 22<br>22 | 7109 | 7615 | L gene (RdRP) | 56 | 507 |
| 17 | tomato vitivirus 1 | NBflexi1-F<br>NBflexi1-R | forward<br>reverse | TCTTTCCCCTCTTGATCTGTGC<br>GTGAACCCTGAATTGGTTGAGC | 22<br>22 | 1057 | 1540 | movement protein | 55 | 484 |
| 18 | Prunus virus I | NBromo1-F<br>NBromo1-R | forward<br>reverse | AAGTTTCGAGACCTTTGCGTTG<br>CTCAAACACACTTCCGCTTCAG | 22<br>22 | 826 | 1699 | movement protein | 55 | 874 |
| 19 | tomato ilarvirus 1 | NBromo2-F<br>NBromo2-R | forward<br>reverse | ACATGGCGTTAGATGGTAGGTC<br>AAATTCGCAGACAAGGTTCGTG | 22<br>22 | 945 | 1824 | movement protein | 55 | 880 |
| 20 | Ranunculus white mottle ophiovirus | RWMV-F<br>RWMV-R | forward<br>reverse | TGTGTGTTCATCTCTCTTCTGC<br>ACAGGGAAGTGAATCACACCTA | 22<br>22 | 804 | 1296 | coat protein | 54 | 493 |
| 21 | tomato betanucleorhabdo-virus 1 | NRhabdo1-F<br>NRhabdo1-R | forward<br>reverse | GACGGTAGGTTACAATCTCC<br>TGATAGGGCTAGGATATGGG | 20<br>20 | 12078 | 12589 | L gene (RdRP) | 55 | 512 |
| 22 | Pastinaca cytorhabdovirus 1 | NRhab3-F<br>NRhab3-R | forward<br>reverse | GAGGAAAAAGTCTGTCATGGAC<br>GCAAGGTAATAATAGCACTCGG | 22<br>22 | 1395 | 1925 | L gene (RdRP) | 52 | 531 |
| 23 | tomato betanucleorhabdo-virus 2 | NRhab4-F<br>NRhab4-R | forward<br>reverse | TTCCTGTTCATTATCACAATGC<br>GTTAGTTGACCAAGAGTACCAG | 22<br>22 | 4279 | 4874 | L gene (RdRP) | 51 | 596 |
| 24 | Picris betanucleorhabdo-virus 1 | NRhab5-F<br>NRhab5-R | forward<br>reverse | ATTGTTACACGATATTGCTGGG<br>TTCCTCATATCTCCACCTCAAC | 22<br>22 | 3300 | 3816 | L gene (RdRP) | 52 | 517 |
| 25 | Cirsium cytorhabdovirus 1 | NRhab6-F<br>NRhab6-R | forward<br>reverse | TTTAGTTAGATCATTCACGGCG<br>TGAGGTCCCTTGATAATCGATC | 22<br>22 | 2204 | 2742 | L gene (RdRP) | 52 | 539 |
| 26 | Taraxacum betanucleorhabdovirus 1 | NRhab7-F<br>NRhab7-R | forward<br>reverse | ATAGTTCGGACAGATCAAGGAG<br>ATCTCAAATGTTTGCCACTCTC | 22<br>22 | 5196 | 5757 | L gene (RdRP) | 52 | 562 |
| 27 | Picris cytorhabdovirus 1 | NRhab8-F<br>NRhab8-R | forward<br>reverse | TCGACCAAAAGATAACAACGAC<br>CTTTGAAAATCACTAGTCCGGG | 22<br>22 | 5798 | 6389 | L gene (RdRP) | 52 | 592 |
| 28 | Taraxacum cytorhabdovirus 1 | NRhab9-F<br>NRhab9-R | forward<br>reverse | CTGTATGTGGTGAAGTCAATGG<br>TCTCATTCTTTTCGCTTCTTCG | 22<br>22 | 2799 | 3347 | L gene (RdRP) | 52 | 549 |
| 29 | tomato alphanucleorhabdo-virus 1 | NRhab10-F<br>NRhab10-R | forward<br>reverse | GATTGTATTTCCCACTACGGACAAC<br>CATACCATCATCACATAGTGTTGGC | 25<br>25 | 3182 | 4230 | L gene (RdRP) | 55 | 1049 |
| 30 | Leveillula taurica associated rhabdo-like virus 1 | NRhab2-F<br>NRhab2-R | forward<br>reverse | CCACATTATGACACAAGACCAG<br>TAAGCTTTGTTACCTAACGCAC | 22<br>22 | 330 | 844 | L gene (RdRP) | 52 | 515 |
| 31 | eggplant mottled dwarf alphanucleorhabdovirus | EMDV-F<br>EMDV-R | forward<br>reverse | TACTCATTACACAAAGAGAAGC<br>CGGTATAGTTATACTAGCAGCA | 22<br>22 | 143 | 699 | L gene (RdRP) | 51 | 557 |

| | | | | | | | | | | |
|---|---|---|---|---|---|---|---|---|---|---|
| 32 | Physostegia chlorotic mottle alphanucleorhabdovirus | PhCMoV-F<br>PhCMoV-R | forward<br>reverse | ATAGTGACATTCTGTTTGACCG<br>CCCATACTACCCATTATTCTGC | 22<br>22 | 2810 | 3463 | L gene (RdRP) | 52 | 654 |
| 33 | tomato fruit blotch virus | ToFBV-R3-F<br>ToFBV-R3-R | forward<br>reverse | GTGGTTATTATGGATATACCTGCG<br>GAGAGAACACAAAACAAGAAGC | 24<br>22 | 654 | 1371 | coat protein | 52 | 718 |
| 34 | Solanum nigrum ilarvirus 1 | SnIV-R3-F<br>SnIV-R3-R | forward<br>reverse | GTATGAAAACTTCAACCTCTCC<br>ATATAGCTACCCAGAAATCAGC | 22<br>22 | 1248 | 1916 | coat protein | 52 | 669 |
| 35 | tomato matilda virus | TMaV-F<br>TMaV-R | forward<br>reverse | ACTAGCCGTTATATTTAGTGGG<br>CTACTATACTGAGAACTCCTTTCC | 22<br>24 | 5293 | 5900 | polyprotein | 52 | 608 |
| 36 | Taraxacum viroid-like circular RNA 1 | NVrd1-F-LP<br>NVrd1-R-LP | forward<br>reverse | TCGGCTAGTCTTTTTCGGTAAC<br>AGGTGAAAGCCTTTCCCTATCC | 22<br>22 | 138 | 392 | not applicable | 55 | 255 |
| | (Note: LP - linear amplification, C - circular amplification) | NVrd1-F-C<br>NVrd1-R-C | primer 1<br>primer 2 | GGCTCGCTTAGACTCTACAAATTG<br>CTGGTTTGAGTCTTAGAACTGACC | 24<br>24 | 277<br>(+) strand | 291<br>(-) strand | not applicable | 55 | 433 |

**Supplementary Table 8.** RT-PCR thermocycling conditions used in the detection of selected viruses and putative viroids in associated plant hosts.

| | | |
|---|---|---|
| Reverse transcription step (2 sub-steps): | 30 min at 50ºC<br>15 min at 95ºC | |
| PCR step (3 sub-steps): | 0.5 min at 94ºC<br>0.5 min at annealing temperature (ºC) in **Supplementary Table 7**<br>1.0 min at 72ºC | 35 cycles |
| Extension step: | 10 min at 72ºC | |

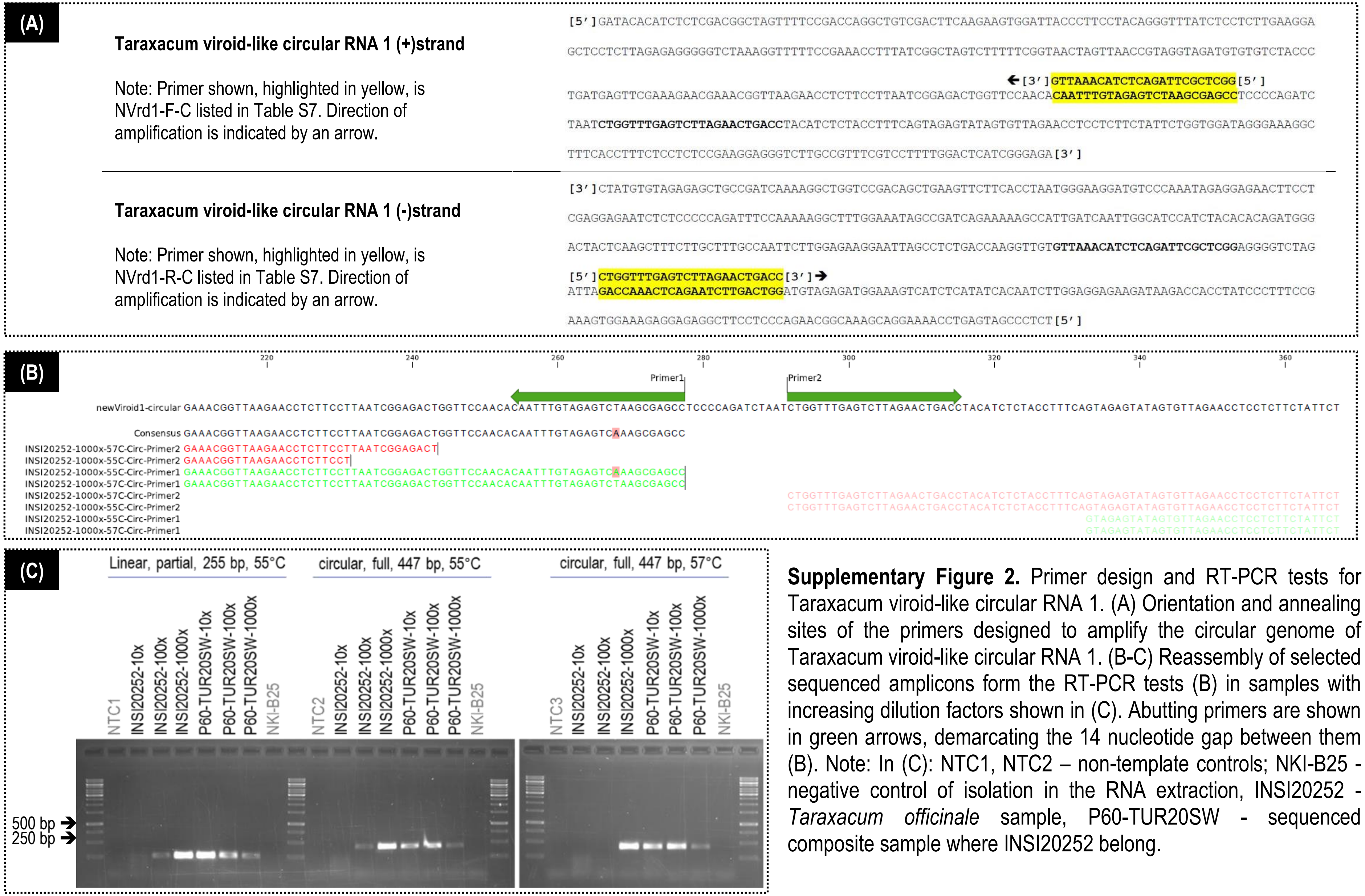


**Supplementary Figure 2.** Primer design and RT-PCR tests for Taraxacum viroid-like circular RNA 1. (A) Orientation and annealing sites of the primers designed to amplify the circular genome of Taraxacum viroid-like circular RNA 1. (B-C) Reassembly of selected sequenced amplicons form the RT-PCR tests (B) in samples with increasing dilution factors shown in (C). Abutting primers are shown in green arrows, demarcating the 14 nucleotide gap between them (B). Note: In (C): NTC1, NTC2 – non-template controls; NKI-B25 - negative control of isolation in the RNA extraction, INSI20252 - *Taraxacum officinale* sample, P60-TUR20SW - sequenced composite sample where INSI20252 belong.

**Supplementary Table 9.** List of confirmed associated plant hosts of selected viruses. For information related to the individual plant samples and the composite samples, please refer to Supplementary Table 1 and 2. Note: In most of the cases infection of multiple viruses in individual plants with cannot be excluded, except if stated otherwise.

| No. | Virus name | Sample ID | Tissue | Plant species name | Plant family | Symptoms (if any) |
|---|---|---|---|---|---|---|
| 01 | Artemesia fimovirus 1 | INSI20094 | leaf | *Artemisia verlotiorum* | Asteraceae | mild leaf yellowing |
| 02 | Calystegia geminivirus 1 | INSI20122 | leaf | *Calystegia* sp. | Convolvulaceae | leaf yellowing with crumpling deformation |
| 03 | Plantago potyvirus 1 | INSI19064 | leaf | *Plantago lanceolata* | Plantaginaceae | interveinal leaf yellowing with deformation |
| 04 | Mentha macluravirus 1 | INSI20169 | leaf | *Mentha spicata* | Lamiaceae | mild leaf yellowing with a few necrotic spots |
| 05 | Rumex potyvirus 1 | INSI20188 | leaf | *Rumex* sp. | Polygonaceae | ring-like chlorotic lesions and red spots on leaves |
| | | INSI20199 | leaf | *Convolvulus* sp. | Convolvulaceae | leaf chlorosis |
| 06 | Broad-leafed dock virus A, isolate 2 | INSI20253 | leaf | *Rumex crispus* | Polygonaceae | leaf chlorosis with mottling |
| 07 | Pastinaca umbravirus 1 | INSI19137 | leaf | *Pastinaca sativa* | Apiaceae | necrotic spots, mosaic on leaves |
| | | INSI19156 | leaf | *Pastinaca sativa* | Apiaceae | irregular leaf chlorosis |
| | | INSI20168 | leaf | *Pastinaca sativa* | Apiaceae | irregular leaf chlorosis |
| 08 | Picris umbravirus 1 | INSI19136 | leaf | *Picris echoides* | Asteraceae | necrotic leaf lesions |
| | | INSI20082 | leaf | *Picris echoides* | Asteraceae | chlorotic leaf spots |
| 09 | Convolvulus aureusvirus 1 | INSI20072 | leaf | *Convolvulus arvensis* | Convolvulaceae | necrotic leaf lesions |
| | | INSI20073 | leaf | *Convolvulus arvensis* | Convolvulaceae | leaf curling with mild yellowing |
| 10 | Calystegia pelarspovirus 1 | INSI20122 | leaf | *Calystegia* sp. | Convolvulaceae | leaf yellowing with crumpling deformation |
| | | INSI20163 | leaf | *Calystegia* sp. | Convolvulaceae | necrotic leaf lesions with yellow halo |
| 11 | Cichorium alphacarmovirus 1 | INSI19123 | leaf | *Cichorium intybus* | Asteraceae | leaf mosaic, leaf dwarfing and deformation |
| | | INSI20074 | leaf | *Cichorium intybus* | Asteraceae | necrotic leaf lesions |
| | | INSI20075 | leaf | *Picris echoides* | Asteraceae | leaf mosaic |
| | | INSI20078 | leaf | *Picris echoides* | Asteraceae | redness of leaves, necrosis on flowers |
| | | INSI20082 | leaf | *Picris echoides* | Asteraceae | chlorotic leaf spots |
| 12 | Pastinaca potexvirus 1 | INSI19137 | leaf | *Pastinaca sativa* | Apiaceae | necrotic spots, mosaic on leaves |
| 13 | Plant associated tobamo-like virus 1 | INSI19082 | leaf | *Convolvulus arvensis* | Convolvulaceae | with powdery mildew colonization / infection |
| | | INSI20147 | leaf | *Solanum lycopersicum* | Solanaceae | none (asymptomatic) |
| | | INSI20152 | leaf | *Solanum lycopersicum* | Solanaceae | none (asymptomatic) |
| | | INSI20157 | leaf | *Solanum lycopersicum* | Solanaceae | none (asymptomatic) |
| | | INSI20158 | leaf | *Solanum lycopersicum* | Solanaceae | none (asymptomatic) |

| | | | | | | |
|---|---|---|---|---|---|---|
| | | INSI20159 | leaf | *Solanum lycopersicum* | Solanaceae | none (asymptomatic) |
| | | INSI20146 | leaf | *Solanum lycopersicum* | Solanaceae | interveinal leaf yellowing |
| | | INSI20148 | leaf | *Solanum lycopersicum* | Solanaceae | interveinal leaf yellowing |
| | | INSI20149 | leaf | *Solanum lycopersicum* | Solanaceae | interveinal leaf yellowing |
| | | INSI20150 | leaf | *Solanum lycopersicum* | Solanaceae | interveinal leaf yellowing |
| | | INSI20151 | leaf | *Solanum lycopersicum* | Solanaceae | interveinal leaf yellowing |
| | | INSI20154 | fruit | *Solanum lycopersicum* | Solanaceae | irregular yellow discolorations on fruits |
| | | INSI20155 | fruit | *Solanum lycopersicum* | Solanaceae | irregular yellow discolorations on fruits |
| | | INSI20156 | fruit | *Solanum lycopersicum* | Solanaceae | irregular yellow discolorations on fruits |
| | | INSI20156F | fruit | *Solanum lycopersicum* | Solanaceae | irregular yellow discolorations on fruits |
| 14 | Plantago tobamovirus 1 | INSI19124 | leaf | *Plantago major* | Plantaginaceae | leaf mosaic |
| 15 | Mercurialis orthotospovirus 1 | INSI19080 | leaf | *Mercurialis annua* | Euphorbiaceae | leaf chlorosis with slight deformation |
| | | INSI19098 | leaf | *Mercurialis annua* | Euphorbiaceae | leaf chlorosis with deformation and necrotic lesions |
| | | INSI19121 | leaf | *Mercurialis annua* | Euphorbiaceae | none (asymptomatic) |
| | | INSI20111 | leaf | *Mercurialis annua* | Euphorbiaceae | leaf deformation |
| | | INSI20112 | leaf | *Mercurialis annua* | Euphorbiaceae | necrotic leaf spots, shot-hole symtpoms |
| 16 | Tomato associated bunya-like virus 1 | INSI20083 | leaf | *Solanum lycopersicum* | Solanaceae | leaf yellowing with necrotic lesions |
| | | INSI20084 | leaf | *Solanum lycopersicum* | Solanaceae | leaf yellowing |
| | | INSI20085 | leaf | *Solanum lycopersicum* | Solanaceae | leaf yellowing with necrotic lesions |
| | | INSI20088 | leaf | *Solanum lycopersicum* | Solanaceae | leaf deformation |
| | | INSI20088F | fruit | *Solanum lycopersicum* | Solanaceae | yellow discoloration |
| | | INSI20089 | leaf | *Solanum lycopersicum* | Solanaceae | leaf yellowing with necrotic lesions |
| | | INSI20090 | leaf | *Solanum lycopersicum* | Solanaceae | leaf yellowing with necrotic lesions |
| 17 | Tomato vitivirus 1 | INSI20124 | leaf | *Solanum lycopersicum* | Solanaceae | necrotic leaf lesions |
| | | INSI20124F | fruit | *Solanum lycopersicum* | Solanaceae | yellow discoloration on fruits, necrosis on sepals |
| 18 | Prunus virus I | INSI20078 | leaf | *Picris echoides* | Asteraceae | red discolorations on leaf and sepal tips |
| 19 | Tomato ilarvirus 1 | INSI20087 | leaf | *Solanum lycopersicum* | Solanaceae | none (asymptomatic) |
| 20 | Ranunculus white mottle ophiovirus | INSI19037 | leaf | *Solanum lycopersicum* | Solanaceae | necrotic leaf spots |
| | | INSI19040 | leaf | *Solanum lycopersicum* | Solanaceae | necrotic leaf spots, discoloration on fruits |
| | | INSI19042 | leaf | *Solanum lycopersicum* | Solanaceae | necrotic leaf spots |
| | | INSI19043 | leaf | *Solanum lycopersicum* | Solanaceae | mild leaf chlorosis |
| | | INSI19073 | leaf | *Solanum lycopersicum* | Solanaceae | none (asymptomatic) |
| | | INSI19074 | leaf | *Solanum lycopersicum* | Solanaceae | leaf yellowing with necrotic lesions |

| | | | | | | |
|---|---|---|---|---|---|---|
| | | INSI19102 | leaf | *Solanum nigrum* | Solanaceae | none (asymptomatic) |
| | | INSI20177 | leaf | *Solanum lycopersicum* | Solanaceae | leaf folding |
| | | INSI20177F | fruit | *Solanum lycopersicum* | Solanaceae | fruit deformation, cracking |
| 21 | Tomato betanucleorhabdovirus 1 | INSI19008 | leaf | *Solanum lycopersicum* | Solanaceae | leaf folding |
| 22 | Pastinaca cytorhabdovirus 1 | INSI19137 | leaf | *Pastinaca sativa* | Apiaceae | necrotic spots, mosaic on leaves |
| 23 | Tomato betanucleorhabdovirus 2 | INSI20041 | leaf | *Solanum lycopersicum* | Solanaceae | none (asymptomatic) |
| | | INSI20068 | leaf | *Solanum lycopersicum* | Solanaceae | none (asymptomatic) |
| | | INSI20123 | leaf | *Solanum lycopersicum* | Solanaceae | leaf yellowing and deformation, uneven fruit ripening |
| | | INSI20132 | leaf | *Solanum lycopersicum* | Solanaceae | leaf yellowing and deformation |
| | | INSI20133 | leaf | *Solanum lycopersicum* | Solanaceae | none (asymptomatic) |
| 24 | Picris betanucleorhabdovirus 1 | INSI20078 | leaf | *Picris echoides* | Asteraceae | red discolorations on leaf and sepal tips |
| | | INSI20080 | leaf | *Picris echoides* | Asteraceae | red discolorations on leaf and sepal tips |
| | | INSI20138 | leaf | *Picris echoides* | Asteraceae | cholorotic and necrotic spots |
| 25 | Cirsium cytorhabdovirus 1 | INSI20161 | leaf | *Cirsium arvense* | Asteraceae | leaf yellowing (near leaf lamina) |
| 26 | Taraxacum betanucleorhabdovirus 1 | INSI20194 | leaf | *Taraxacum officinale* | Asteraceae | systemic leaf chlorosis |
| | | INSI20252 | leaf | *Taraxacum officinale* | Asteraceae | interveinal leaf chlorosis |
| 27 | Picris cytorhabdovirus 1 | INSI20080 | leaf | *Picris echoides* | Asteraceae | red discolorations on leaf and sepal tips |
| | | INSI20081 | leaf | *Picris echoides* | Asteraceae | leaf mosaic |
| | | INSI20082 | leaf | *Picris echoides* | Asteraceae | red lesions on leaves and midribs |
| 28 | Taraxacum cytorhabdovirus 1 | INSI20194 | leaf | *Taraxacum officinale* | Asteraceae | systemic leaf chlorosis |
| 29 | Tomato alphanucleorhabdovirus 1 | INSI20029 | leaf | *Solanum lycopersicum* | Solanaceae | fruit mottling, showing large irregular shape yellow dents |
| | | INSI20030 | leaf | *Solanum lycopersicum* | Solanaceae | fruit mottling, showing large irregular shape yellow dents |
| | | INSI20030F | fruit | *Solanum lycopersicum* | Solanaceae | fruit mottling, showing large irregular shape yellow dents |
| 30 | Leveillula taurica associated rhabdo-like virus 1 | INSI19051 | leaf | *Solanum lycopersicum* | Solanaceae | leaves with powdery mildew colonization, necrotic spots |
| | | INSI19053 | leaf | *Solanum lycopersicum* | Solanaceae | leaves with powdery mildew colonization, necrotic spots |
| | | INSI19054 | leaf | *Solanum lycopersicum* | Solanaceae | leaves with powdery mildew colonization, necrotic spots |
| | | INSI19055 | leaf | *Solanum lycopersicum* | Solanaceae | leaves with powdery mildew colonization, necrotic spots |
| | | INSI19056 | leaf | *Solanum lycopersicum* | Solanaceae | leaves with powdery mildew colonization, necrotic spots |
| | | INSI19058 | leaf | *Solanum lycopersicum* | Solanaceae | leaves with powdery mildew colonization, necrotic spots |
| 31 | Eggplant mottled dwarf alphanucleorhabdovirus | INSI20038 | leaf | *Solanum lycopersicum* | Solanaceae | necrotic spots on leaves |
| | | INSI20038F | fruit | *Solanum lycopersicum* | Solanaceae | fruit mottling, showing circular yellow dents |
| | | INSI20039 | leaf | *Solanum lycopersicum* | Solanaceae | necrotic spots on leaves |
| | | INSI20039F | fruit | *Solanum lycopersicum* | Solanaceae | fruit mottling, showing circular yellow dents |

| | | | | | | |
|---|---|---|---|---|---|---|
| 32 | Physostegia chlorotic mottle alphanucleorhabdovirus | INSI19009 | leaf | *Solanum lycopersicum* | Solanaceae | fruit and leaf mottling and yellowing, leaf folding |
| | | INSI20177 | leaf | *Solanum lycopersicum* | Solanaceae | leaf folding |
| | | INSI20177F | fruit | *Solanum lycopersicum* | Solanaceae | fruit deformation, cracking |
| | | INSI20239F | fruit | *Solanum lycopersicum* | Solanaceae | fruit and leaf mottling and yellowing |
| | | INSI20242F | fruit | *Solanum lycopersicum* | Solanaceae | fruit and leaf mottling and yellowing |
| 33 | Tomato fruit blotch virus | INSI19101 | leaf | *Solanum lycopersicum* | Solanaceae | leaf yellowing with necrotic lesions |
| | | INSI19122 | leaf | *Solanum lycopersicum* | Solanaceae | leaf yellowing with necrotic lesions |
| 34 | Solanum nigrum ilarvirus 1 | INSI19127 | leaf | *Solanum lycopersicum* | Solanaceae | none (asymptomatic) |
| | | INSI19133 | leaf | *Solanum lycopersicum* | Solanaceae | none (asymptomatic) |
| | | INSI19134 | leaf | *Solanum lycopersicum* | Solanaceae | mild yellowing and leaf twisting and deformations |
| | | INSI20216 | leaf | *Physalis* sp. | Solanaceae | uneven leaf yellowing |
| 35 | Tomato matilda virus | INSI19073 | leaf | *Solanum lycopersicum* | Solanaceae | none (asymptomatic) |
| | | INSI19085 | leaf | *Solanum lycopersicum* | Solanaceae | none (asymptomatic) |
| | | INSI19086 | leaf | *Solanum lycopersicum* | Solanaceae | none (asymptomatic) |
| | | INSI19087 | leaf | *Solanum lycopersicum* | Solanaceae | none (asymptomatic) |
| | | INSI19088 | leaf | *Solanum lycopersicum* | Solanaceae | none (asymptomatic) |
| | | INSI19089 | leaf | *Solanum lycopersicum* | Solanaceae | none (asymptomatic) |
| | | INSI19091 | leaf | *Solanum lycopersicum* | Solanaceae | none (asymptomatic) |
| | | INSI19092 | leaf | *Solanum lycopersicum* | Solanaceae | none (asymptomatic) |
| | | INSI19093 | leaf | *Solanum lycopersicum* | Solanaceae | none (asymptomatic) |
| | | INSI19081 | leaf | *Solanum lycopersicum* | Solanaceae | corky symptoms on fruits, leaf yellwoing |
| | | INSI19094 | leaf | *Solanum lycopersicum* | Solanaceae | yellow spots on leaves |
| | | INSI19103 | leaf | *Solanum lycopersicum* | Solanaceae | none (asymptomatic) |
| | | INSI19104 | leaf | *Solanum lycopersicum* | Solanaceae | none (asymptomatic) |
| | | INSI19106 | leaf | *Solanum lycopersicum* | Solanaceae | none (asymptomatic) |
| | | INSI19108 | leaf | *Solanum lycopersicum* | Solanaceae | none (asymptomatic) |
| | | INSI19110 | leaf | *Solanum lycopersicum* | Solanaceae | none (asymptomatic) |
| | | INSI19111 | leaf | *Solanum lycopersicum* | Solanaceae | none (asymptomatic) |
| | | INSI19112 | leaf | *Solanum lycopersicum* | Solanaceae | none (asymptomatic) |
| | | INSI19113 | leaf | *Solanum lycopersicum* | Solanaceae | none (asymptomatic) |
| | | INSI19100 | leaf | *Solanum lycopersicum* | Solanaceae | leaf yellowing with necrotic lesions |
| | | INSI19101 | leaf | *Solanum lycopersicum* | Solanaceae | leaf yellowing with necrotic lesions |
| | | INSI19114 | leaf | *Solanum lycopersicum* | Solanaceae | necrotic leaf lesions |
| | | INSI19115 | leaf | *Solanum lycopersicum* | Solanaceae | leaf chlorosis with necrotic lesions |

| | | | | |
|---|---|---|---|---|
| INSI19116 | leaf | *Solanum lycopersicum* | Solanaceae | leaf yellowing with necrotic lesions |
| INSI19117 | leaf | *Solanum lycopersicum* | Solanaceae | leaf yellowing with necrotic lesions |
| INSI19122 | leaf | *Solanum lycopersicum* | Solanaceae | leaf yellowing with necrotic lesions |
| INSI19126 | leaf | *Solanum lycopersicum* | Solanaceae | leaf yellowing with necrotic lesions |
| INSI19128 | leaf | *Solanum lycopersicum* | Solanaceae | leaf yellowing with necrotic lesions |
| INSI19129 | leaf | *Solanum lycopersicum* | Solanaceae | yellow spots on leaves |
| INSI19134 | leaf | *Solanum lycopersicum* | Solanaceae | mild yellowing and leaf twisting and deformations |
| INSI19135 | leaf | *Solanum lycopersicum* | Solanaceae | leaf yellowing with necrotic lesions |
| INSI19127 | leaf | *Solanum lycopersicum* | Solanaceae | none (asymptomatic) |
| INSI19130 | leaf | *Solanum lycopersicum* | Solanaceae | none (asymptomatic) |
| INSI19131 | leaf | *Solanum lycopersicum* | Solanaceae | none (asymptomatic) |
| INSI19132 | leaf | *Solanum lycopersicum* | Solanaceae | none (asymptomatic) |
| INSI19133 | leaf | *Solanum lycopersicum* | Solanaceae | none (asymptomatic) |
| INSI19141 | leaf | *Solanum lycopersicum* | Solanaceae | none (asymptomatic) |
| INSI19142 | leaf | *Solanum lycopersicum* | Solanaceae | none (asymptomatic) |
| INSI19143 | leaf | *Solanum lycopersicum* | Solanaceae | none (asymptomatic) |
| INSI19144 | leaf | *Solanum lycopersicum* | Solanaceae | none (asymptomatic) |
| INSI19147 | leaf | *Solanum lycopersicum* | Solanaceae | none (asymptomatic) |
| INSI19148 | leaf | *Solanum lycopersicum* | Solanaceae | none (asymptomatic) |
| INSI19139 | leaf | *Solanum lycopersicum* | Solanaceae | leaf yellowing with necrotic lesions |
| INSI19149 | leaf | *Solanum lycopersicum* | Solanaceae | leaf yellowing |
| INSI19151 | leaf | *Solanum lycopersicum* | Solanaceae | none (asymptomatic) |
| INSI19154 | leaf | *Solanum lycopersicum* | Solanaceae | leaf mottling, dwarfing and twisting |
| INSI19155 | leaf | *Solanum lycopersicum* | Solanaceae | leaf mottling, dwarfing and twisting |
| INSI19140 | leaf | *Chenopodium* sp. | Chenopodiaceae | yellow spots on leaves |
| INSI19150 | leaf | *Chenopodium* sp. | Chenopodiaceae | yellow spots on leaves |
| INSI19157 | leaf | *Erigeron annuus* | Asteraceae | necrotic lesions on leaves |
| INSI19158 | leaf | *Ranunculus repens* | Ranunculaceae | leaf mosaic |
| INSI20040 | leaf | *Solanum lycopersicum* | Solanaceae | none (asymptomatic) |
| INSI20041 | leaf | *Solanum lycopersicum* | Solanaceae | none (asymptomatic) |
| INSI20042 | leaf | *Solanum lycopersicum* | Solanaceae | none (asymptomatic) |
| INSI20043 | leaf | *Solanum lycopersicum* | Solanaceae | none (asymptomatic) |
| INSI20044 | leaf | *Solanum lycopersicum* | Solanaceae | none (asymptomatic) |

| | | | | | | |
|---|---|---|---|---|---|---|
| | | INSI20030 | leaf | *Solanum lycopersicum* | Solanaceae | necrotic leaf spots |
| | | INSI20031 | leaf | *Solanum lycopersicum* | Solanaceae | necrotic leaf spots |
| | | INSI20033 | leaf | *Solanum lycopersicum* | Solanaceae | necrotic leaf spots |
| | | INSI20035 | leaf | *Solanum lycopersicum* | Solanaceae | purpling of leaf lamina |
| | | INSI20036 | leaf | *Solanum lycopersicum* | Solanaceae | interveinal leaf yellowing |
| | | INSI20037 | leaf | *Solanum lycopersicum* | Solanaceae | necrotic leaf spots, necrosis on leaf lamina |
| | | INSI20038 | leaf | *Solanum lycopersicum* | Solanaceae | necrotic leaf spots, necrosis on leaf lamina |
| | | INSI20055 | leaf | *Solanum lycopersicum* | Solanaceae | purpling of leaf lamina |
| | | INSI20108 | leaf | *Solanum lycopersicum* | Solanaceae | leaf deformation |
| | | INSI20126 | leaf | *Solanum lycopersicum* | Solanaceae | necrotic leaf spots |
| | | INSI20127 | leaf | *Solanum lycopersicum* | Solanaceae | necrotic leaf spots, necrosis on leaf lamina |
| | | INSI20128 | leaf | *Solanum lycopersicum* | Solanaceae | necrotic leaf spots, necrosis on leaf lamina |
| | | INSI20129 | leaf | *Solanum lycopersicum* | Solanaceae | necrotic leaf spots, necrosis on leaf lamina |
| | | INSI20132 | leaf | *Solanum lycopersicum* | Solanaceae | leaf deformation and dwarfing |
| 36 | Taraxacum viroid-like circular RNA 1 | INSI20252 | leaf | *Taraxacum officinale* | Asteraceae | interveinal leaf chlorosis |
