## Additional-File-05_Field Photos of Associated Plant Hosts of Viruses for "In-depth study of tomato and weed viromes reveals undiscovered plant virus diversity in an agroecosystem"

### SUPPLEMENTARY INFORMATION

### Additional File 05

**Supplementary Figure 3.** Representative field photos of confirmed associated plant hosts of selected viruses and putative viroids.

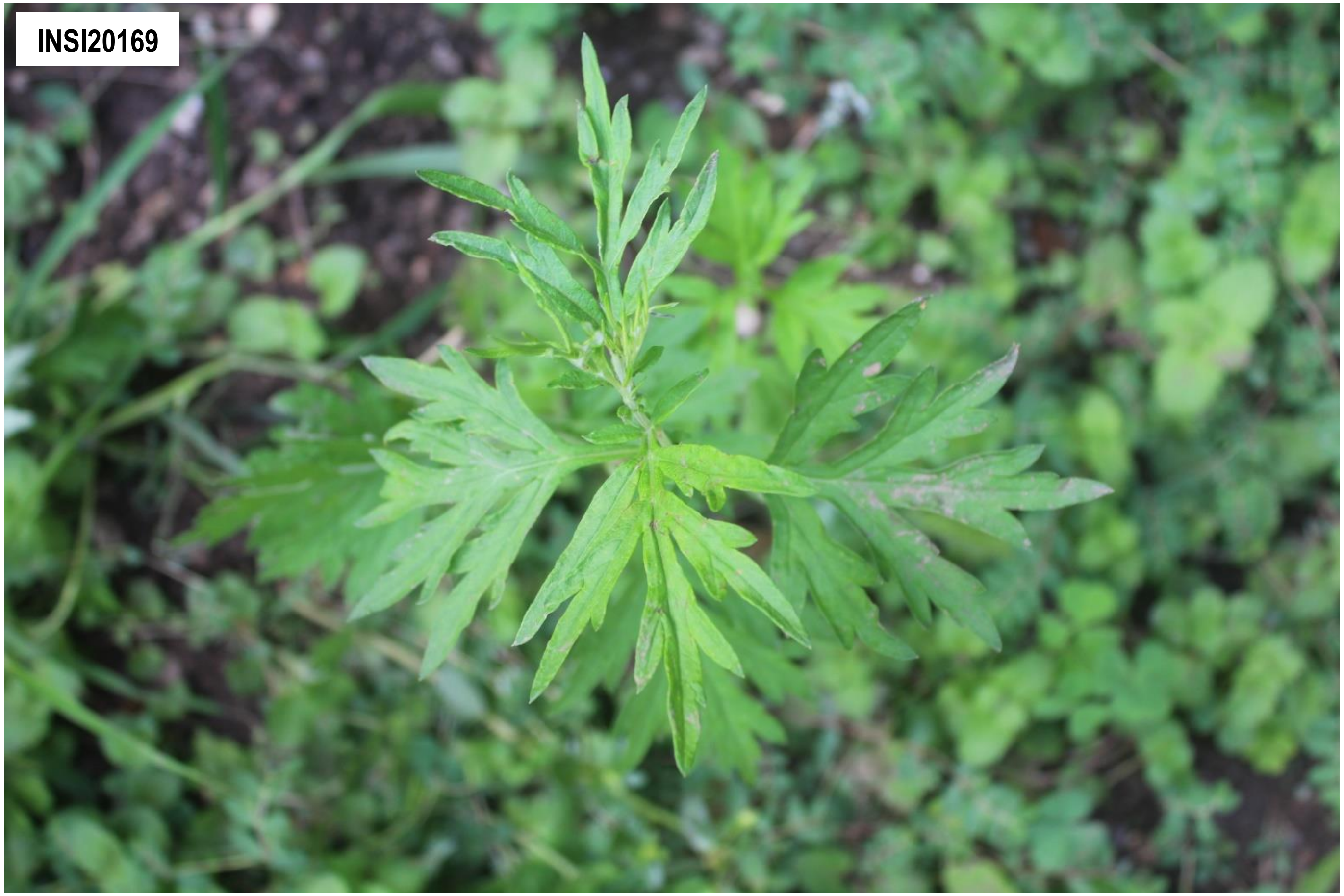


**Supplementary Figure 3-01.** Artermisia fimovirus 1. The confirmed associated plant host(s) shown is/are *Artemisia verlotiorum* (Asteraceae).

**Supplementary Figure 3.** Representative field photos of confirmed associated plant hosts of selected viruses and putative viroids. **Note**: There are other viruses detected in the sequencing pools (*i.e.* composite sample) where the samples shown belong, which implies possible co-infection of viruses on the specific sample shown. Thus, it is difficult to associate the symptom phenotype with the detection of a single specific virus. Complete information on the individual samples can be found in Supplementary Table 1 and the description of symptoms of the associated plant hosts are summarized in Supplementary Table 9.

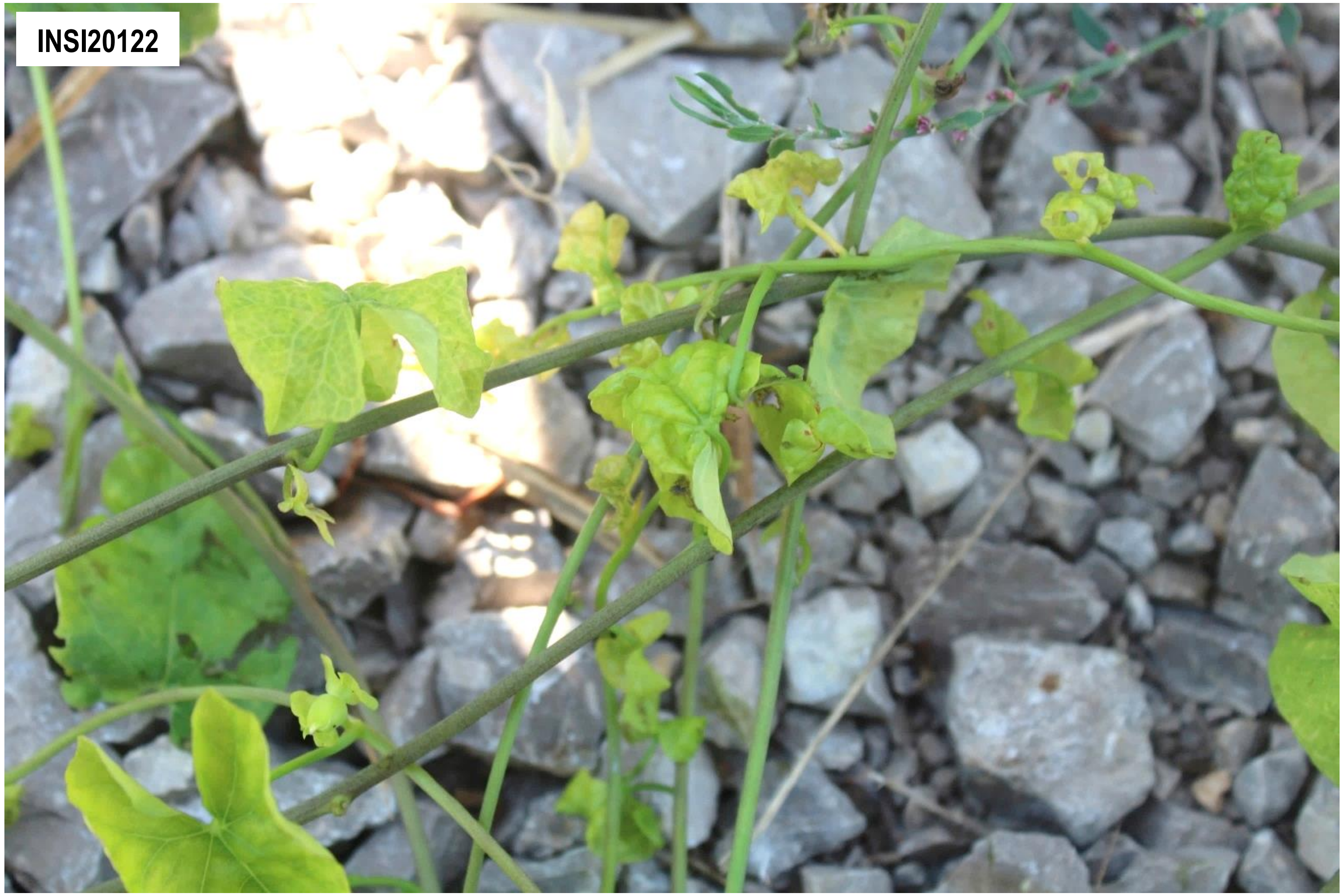


**Supplementary Figure 3-02.** Calystegia geminivirus 1 and Calystegia pelarspovirus 1. The confirmed associated plant host(s) shown is/are *Calystegia sp.* (Convolvulaceae).

**Supplementary Figure 3** (continued)**.** Representative field photos of confirmed associated plant hosts of selected viruses and putative viroids. **Note**: There are other viruses detected in the sequencing pools (*i.e.* composite sample) where the samples shown belong, which implies possible co-infection of viruses on the specific sample shown. Thus, it is difficult to associate the symptom phenotype with the detection of a single specific virus. Complete information on the individual samples can be found in Supplementary Table 1 and the description of symptoms of the associated plant hosts are summarized in Supplementary Table 9.

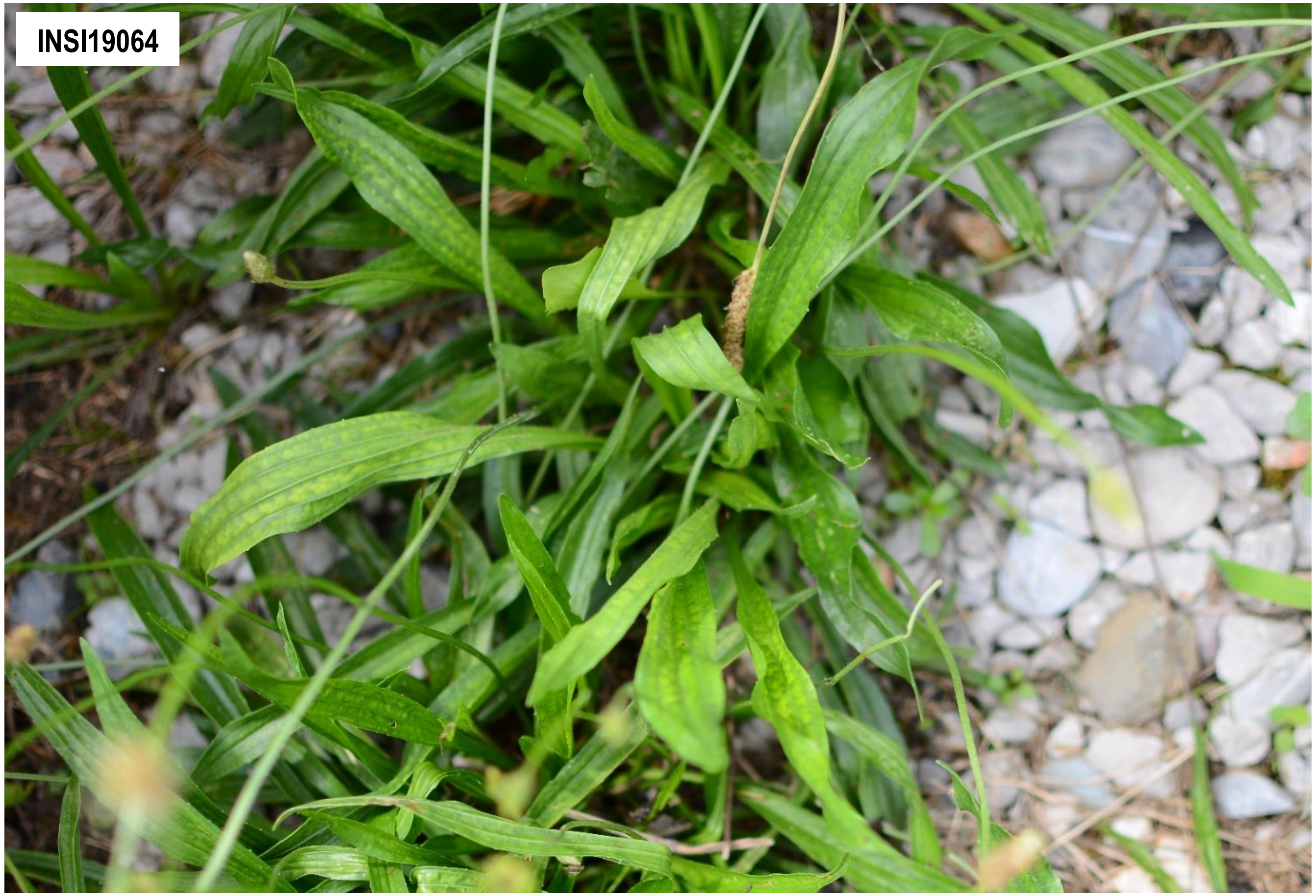


**Supplementary Figure 3-03.** Plantago potyvirus 1. The confirmed associated plant host(s) shown is/are *Plantago lanceolata* (Plantaginaceae).

**Supplementary Figure 3** (continued)**.** Representative field photos of confirmed associated plant hosts of selected viruses and putative viroids. **Note**: There are other viruses detected in the sequencing pools (*i.e.* composite sample) where the samples shown belong, which implies possible co-infection of viruses on the specific sample shown. Thus, it is difficult to associate the symptom phenotype with the detection of a single specific virus. Complete information on the individual samples can be found in Supplementary Table 1 and the description of symptoms of the associated plant hosts are summarized in Supplementary Table 9.

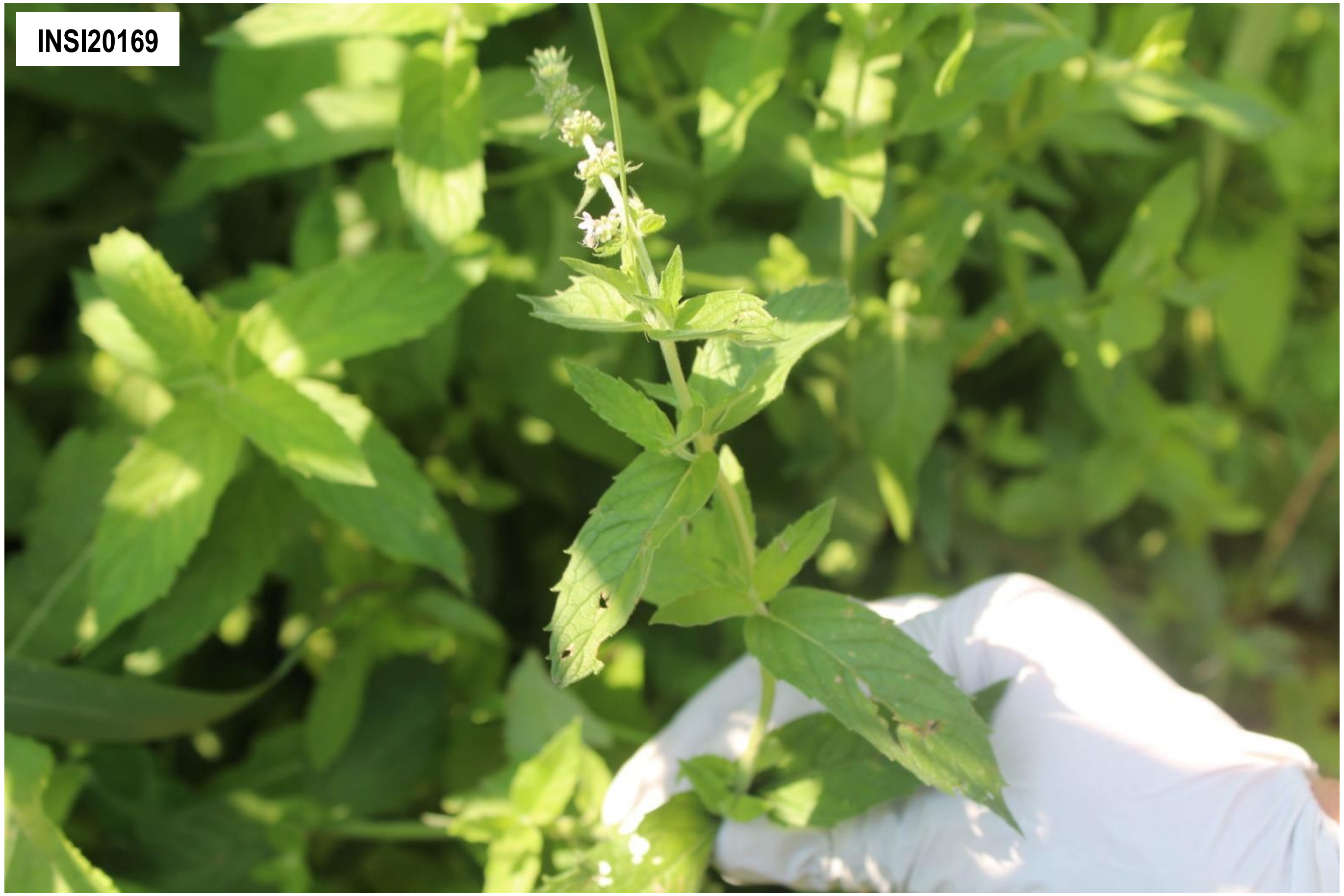


**Supplementary Figure 3-04.** Mentha macluravirus 1. The confirmed associated plant host(s) shown is/are *Mentha spicata* (Lamiaceae).

**Supplementary Figure 3** (continued)**.** Representative field photos of confirmed associated plant hosts of selected viruses and putative viroids. **Note**: There are other viruses detected in the sequencing pools (*i.e.* composite sample) where the samples shown belong, which implies possible co-infection of viruses on the specific sample shown. Thus, it is difficult to associate the symptom phenotype with the detection of a single specific virus. Complete information on the individual samples can be found in Supplementary Table 1 and the description of symptoms of the associated plant hosts are summarized in Supplementary Table 9.

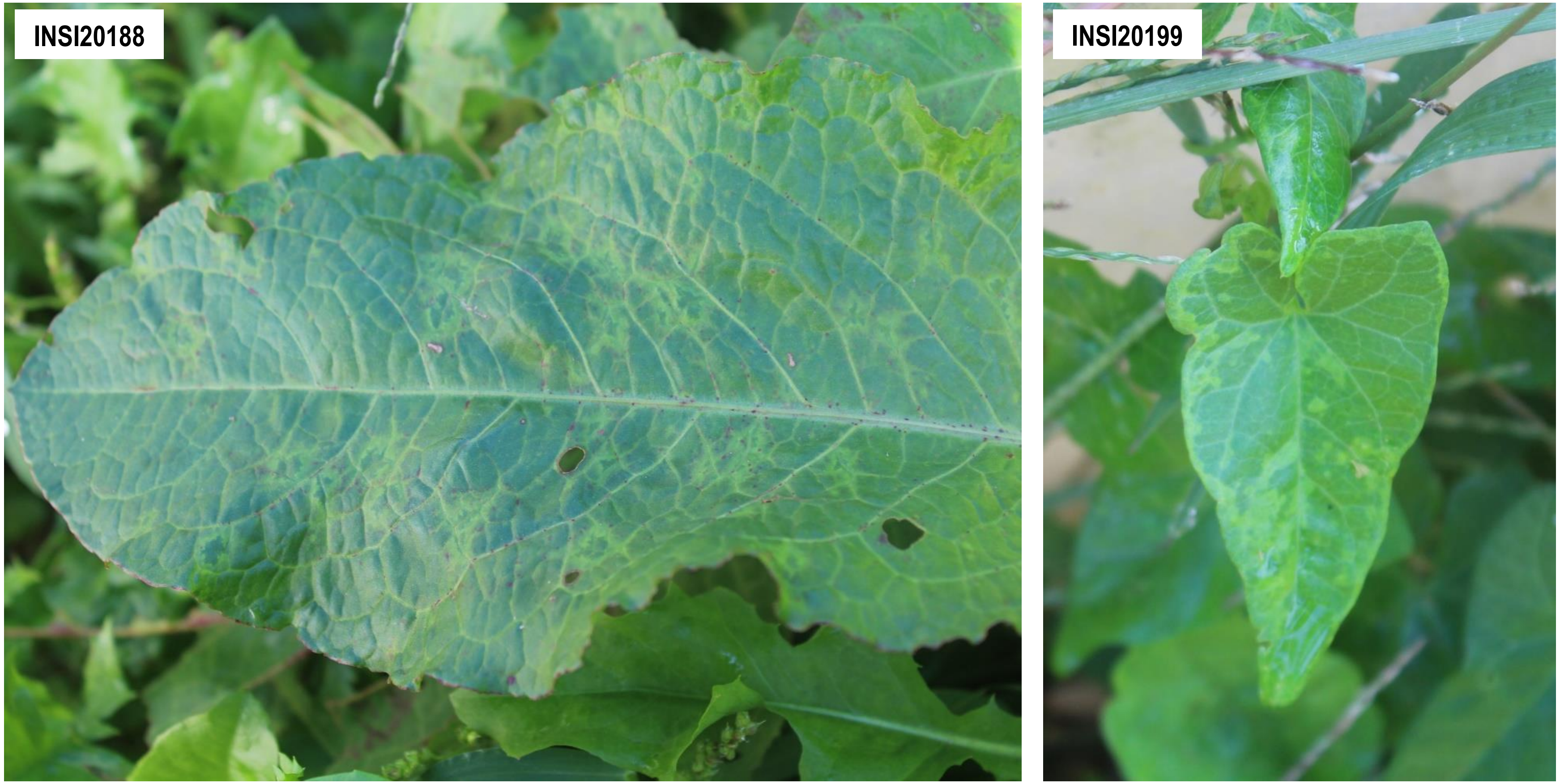


**Supplementary Figure 3-05.** Rumex potyvirus 1. The confirmed associated plant host(s) shown is/are *Rumex* sp. (Polygonaceae) (INSI20188) and *Convolvulus* sp. (Convolvulaceae) (INSI20199).

**Supplementary Figure 3** (continued)**.** Representative field photos of confirmed associated plant hosts of selected viruses and putative viroids. **Note**: There are other viruses detected in the sequencing pools (*i.e.* composite sample) where the samples shown belong, which implies possible co-infection of viruses on the specific sample shown. Thus, it is difficult to associate the symptom phenotype with the detection of a single specific virus. Complete information on the individual samples can be found in Supplementary Table 1 and the description of symptoms of the associated plant hosts are summarized in Supplementary Table 9.

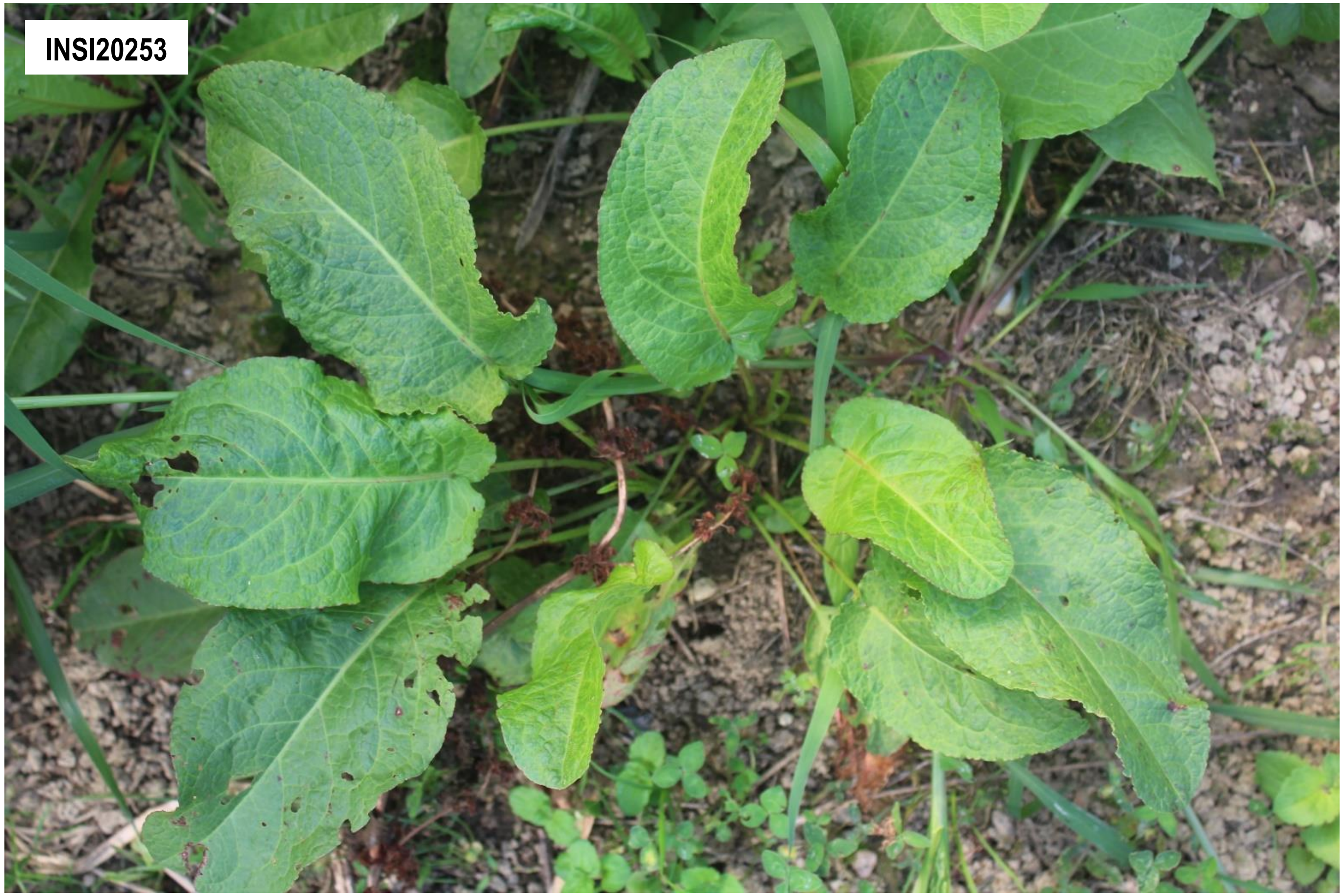


**Supplementary Figure 3-06.** Broad-leafed dock virus A, isolate 2. The confirmed associated plant host(s) shown is/are *Rumex crispus* (Polygonaceae).

**Supplementary Figure 3** (continued)**.** Representative field photos of confirmed associated plant hosts of selected viruses and putative viroids. **Note**: There are other viruses detected in the sequencing pools (*i.e.* composite sample) where the samples shown belong, which implies possible co-infection of viruses on the specific sample shown. Thus, it is difficult to associate the symptom phenotype with the detection of a single specific virus. Complete information on the individual samples can be found in Supplementary Table 1 and the description of symptoms of the associated plant hosts are summarized in Supplementary Table 9.

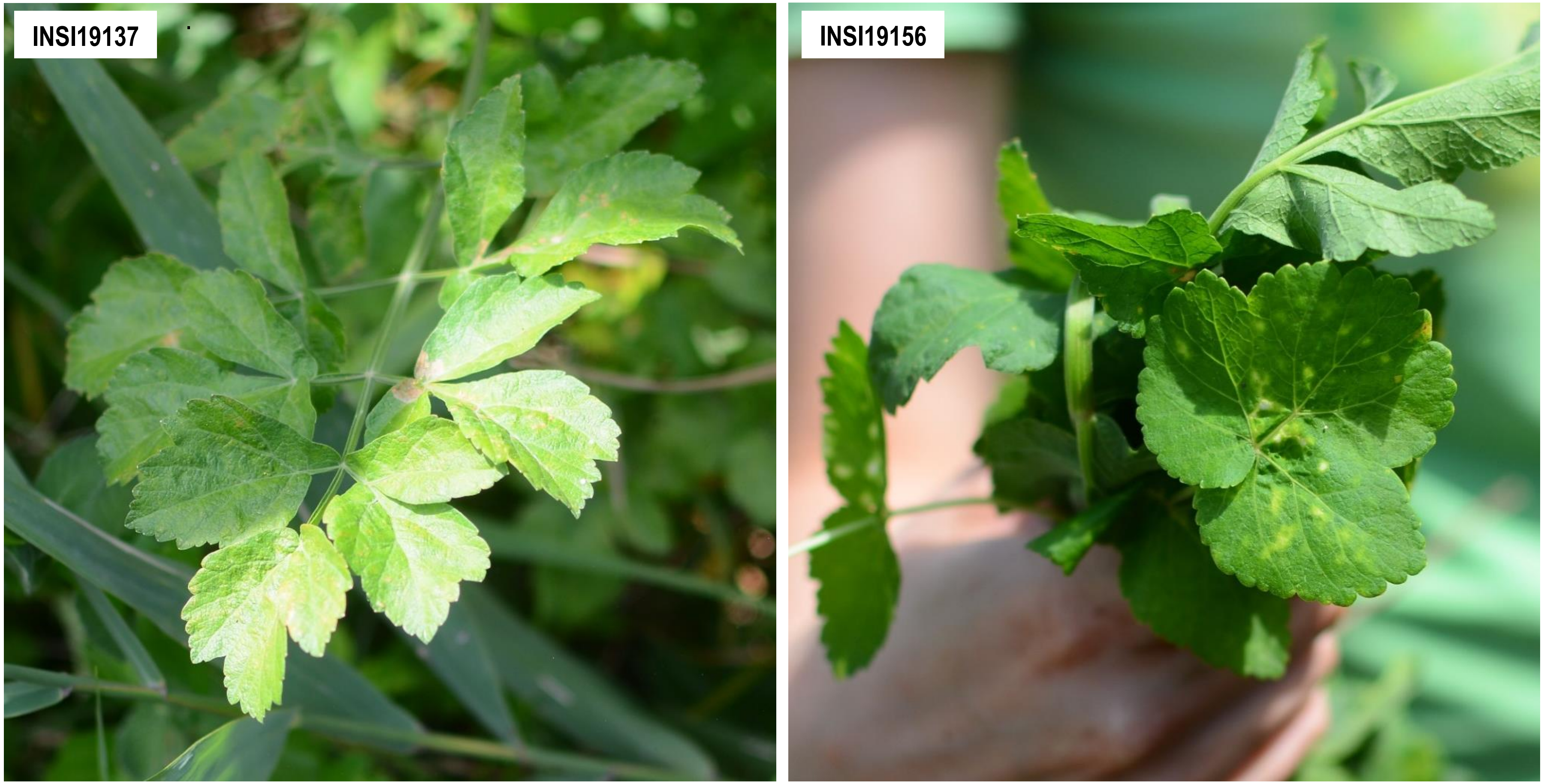


**Supplementary Figure 3-07.** Pastinaca umbravirus 1. The confirmed associated plant host(s) shown is/are *Pastinaca sativa* (Apiaceae). Sample INSI19137 is also the confirmed associated plant host of Pastinaca potexvirus 1 and Pastinaca cytorhabdovirus 1.

**Supplementary Figure 3** (continued)**.** Representative field photos of confirmed associated plant hosts of selected viruses and putative viroids. **Note**: There are other viruses detected in the sequencing pools (*i.e.* composite sample) where the samples shown belong, which implies possible co-infection of viruses on the specific sample shown. Thus, it is difficult to associate the symptom phenotype with the detection of a single specific virus. Complete information on the individual samples can be found in Supplementary Table 1 and the description of symptoms of the associated plant hosts are summarized in Supplementary Table 9.

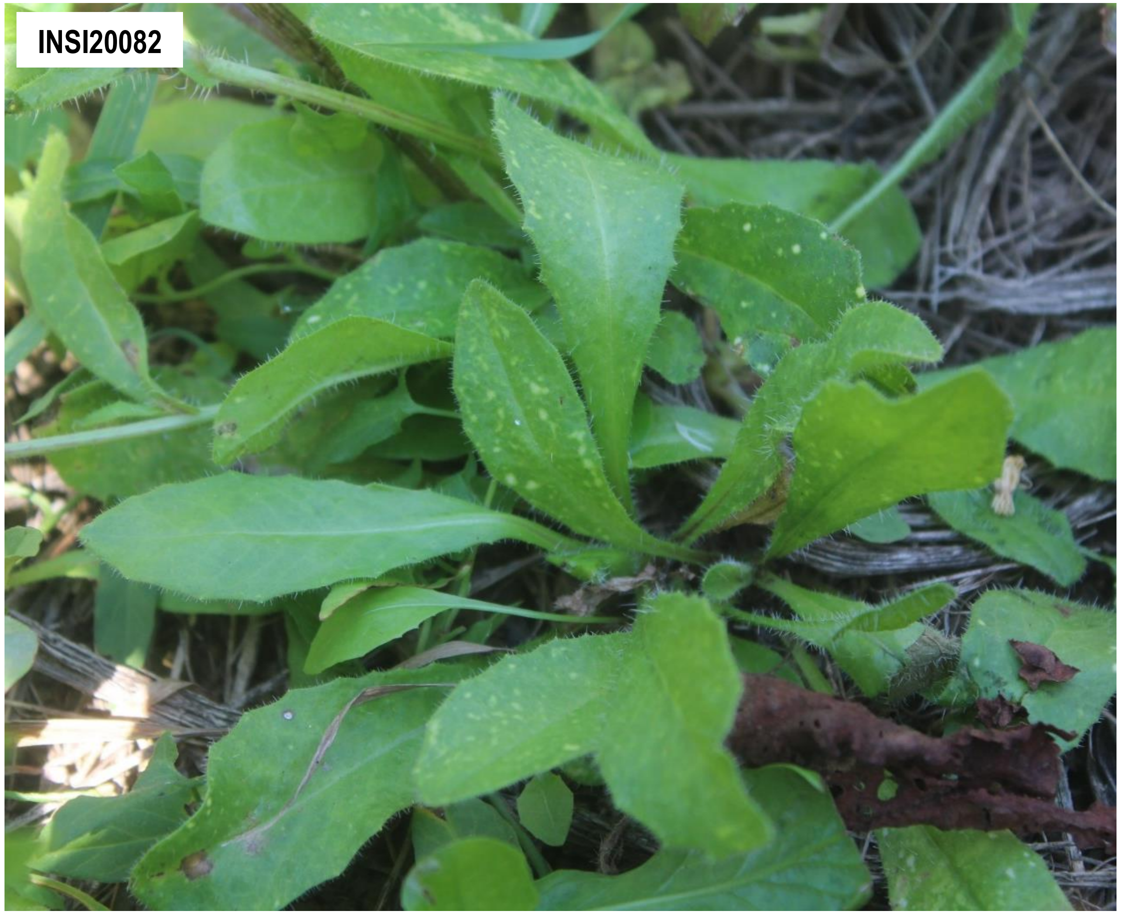


**Supplementary Figure 3-08.** Picris umbravirus 1. The confirmed associated plant host(s) shown is/are *Picris echoides* (Asteraceae). The sample shown is also confirmed associated host of Cichorium alphacarmovirus 1 and Picris cytorhabdovirus 1.

**Supplementary Figure 3** (continued)**.** Representative field photos of confirmed associated plant hosts of selected viruses and putative viroids. **Note**: There are other viruses detected in the sequencing pools (*i.e.* composite sample) where the samples shown belong, which implies possible co-infection of viruses on the specific sample shown. Thus, it is difficult to associate the symptom phenotype with the detection of a single specific virus. Complete information on the individual samples can be found in Supplementary Table 1 and the description of symptoms of the associated plant hosts are summarized in Supplementary Table 9.

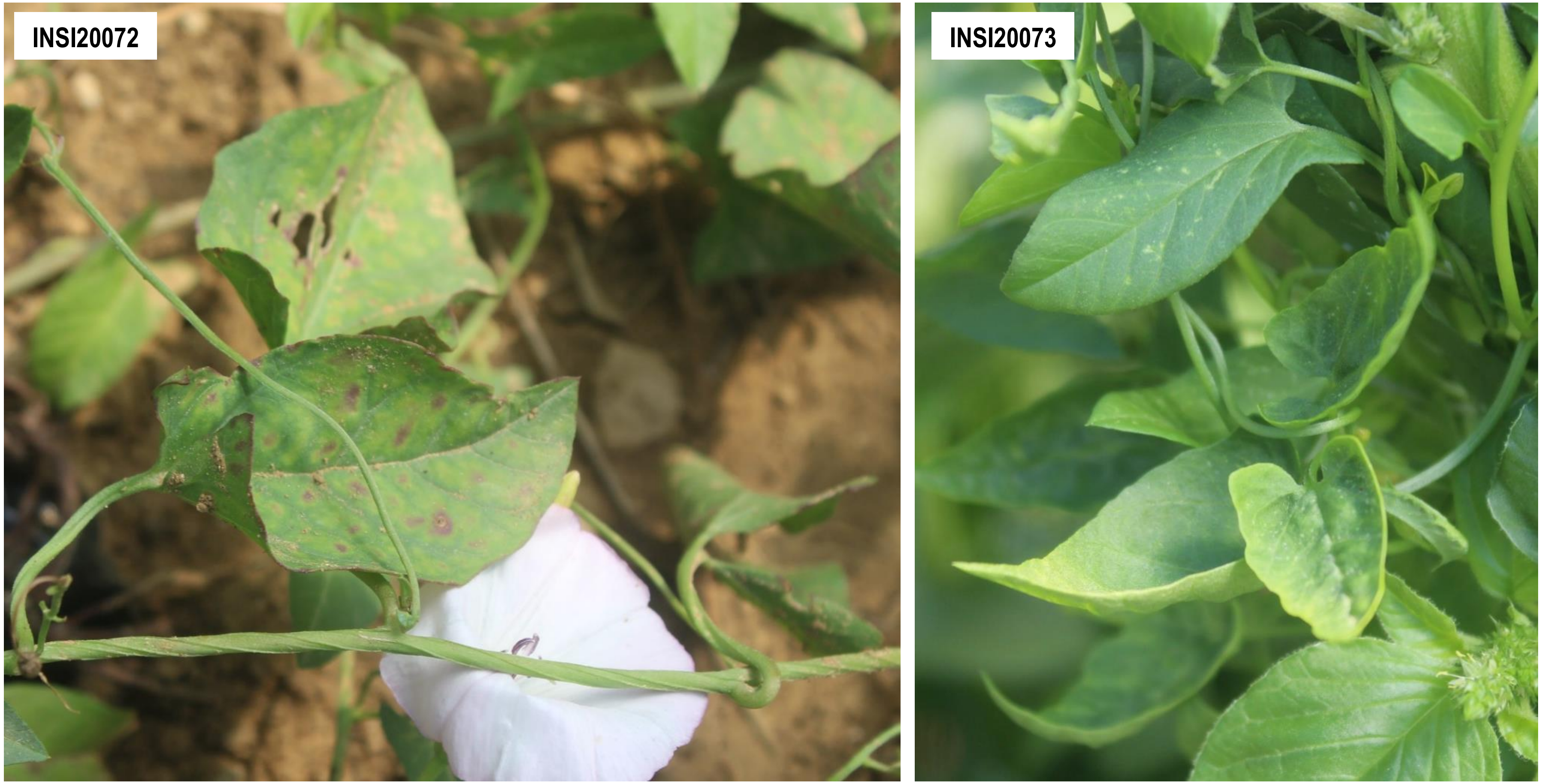


**Supplementary Figure 3-09.** Convolvulus aureusvirus 1. The confirmed associated plant host(s) shown is/are *Convolvulus arvensis* (Convolvulaceae).

**Supplementary Figure 3** (continued)**.** Representative field photos of confirmed associated plant hosts of selected viruses and putative viroids. **Note**: There are other viruses detected in the sequencing pools (*i.e.* composite sample) where the samples shown belong, which implies possible co-infection of viruses on the specific sample shown. Thus, it is difficult to associate the symptom phenotype with the detection of a single specific virus. Complete information on the individual samples can be found in Supplementary Table 1 and the description of symptoms of the associated plant hosts are summarized in Supplementary Table 9.

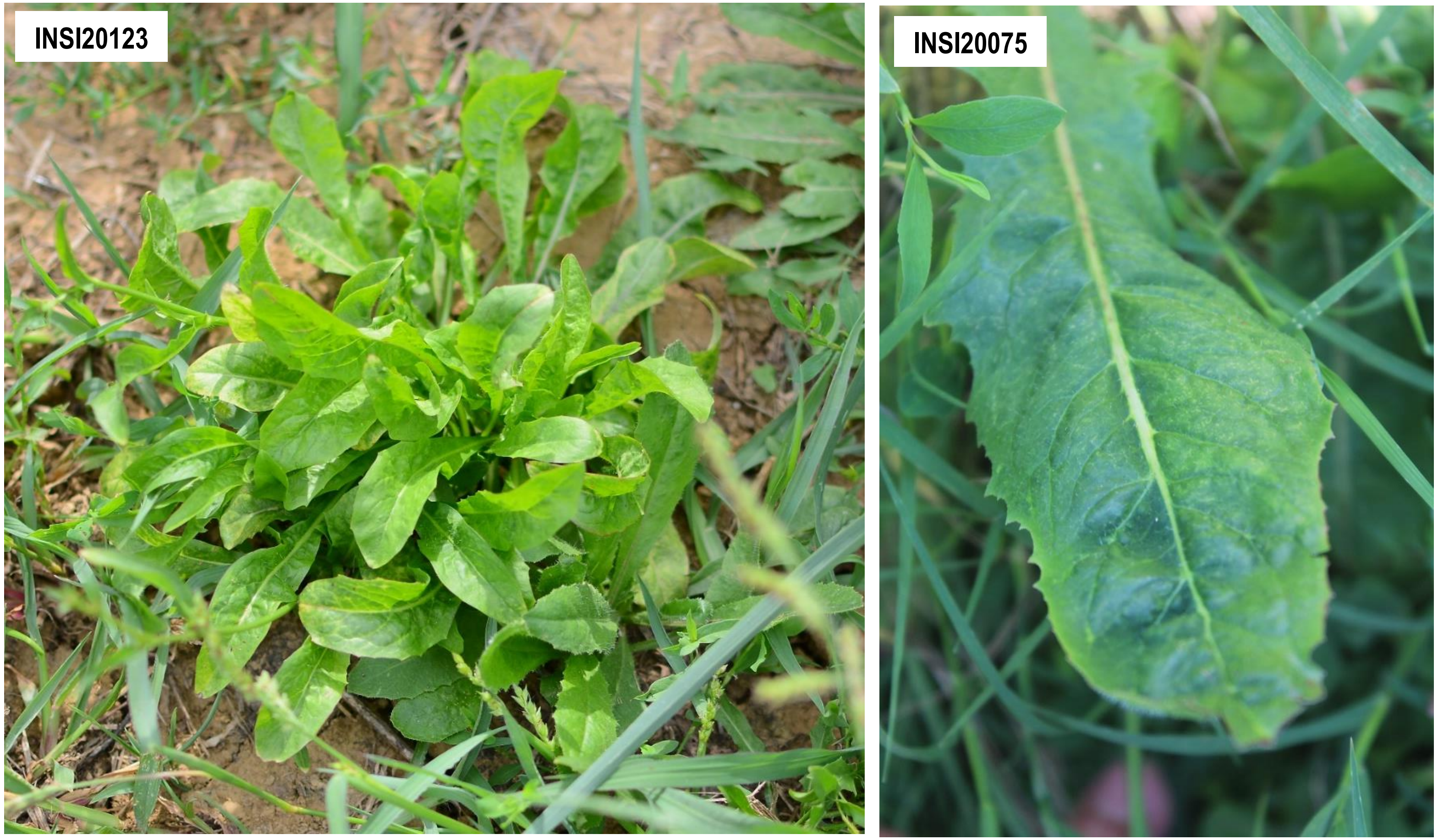


**Supplementary Figure 3-10.** Cichorium alphacarmovirus 1. The confirmed associated plant host(s) shown is/are *Calystegia* sp. (Convolvulaceae).

**Supplementary Figure 3** (continued)**.** Representative field photos of confirmed associated plant hosts of selected viruses and putative viroids. **Note**: There are other viruses detected in the sequencing pools (*i.e.* composite sample) where the samples shown belong, which implies possible co-infection of viruses on the specific sample shown. Thus, it is difficult to associate the symptom phenotype with the detection of a single specific virus. Complete information on the individual samples can be found in Supplementary Table 1 and the description of symptoms of the associated plant hosts are summarized in Supplementary Table 9.

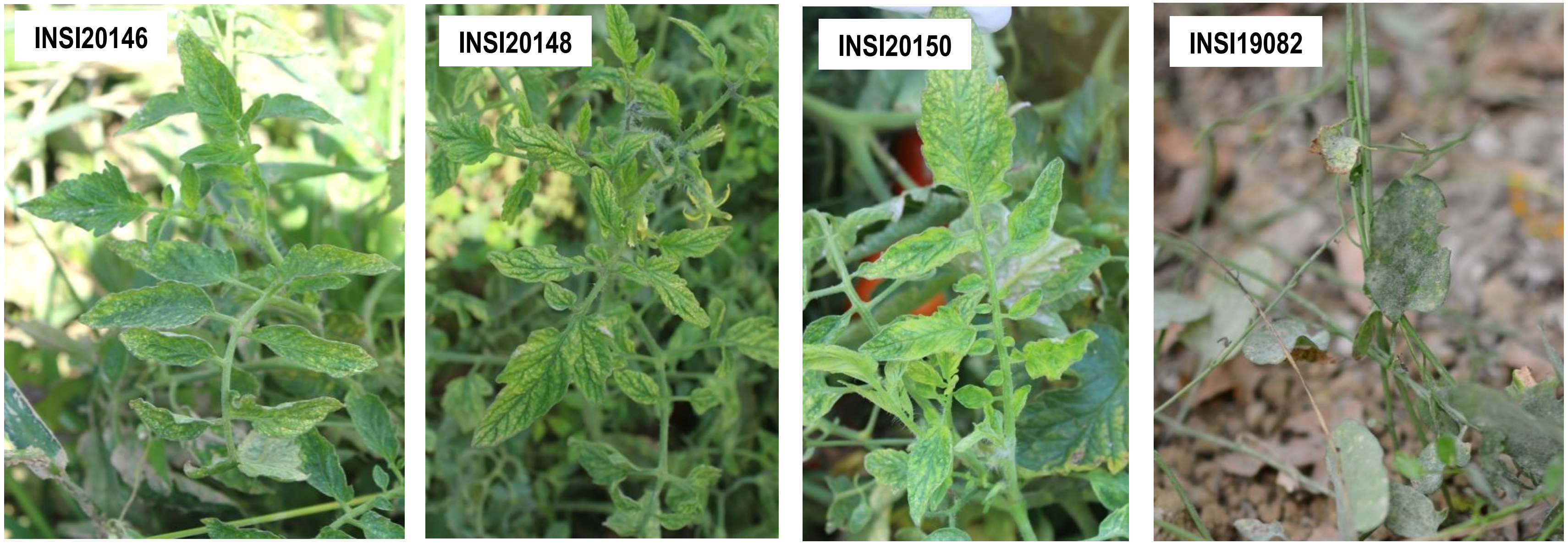


**Supplementary FigureS3-11.** Plant associated tobamo-like virus 1. The confirmed associated plant host(s) shown is/are *Solanum lycopersicum* (Solanaceae) (INSI20146, INSI20148, INSI20150) and *Convolvulus arvensis* (Convolvulaceae) (INSI19082).

**Supplementary Figure 3** (continued)**.** Representative field photos of confirmed associated plant hosts of selected viruses and putative viroids. **Note**: There are other viruses detected in the sequencing pools (*i.e.* composite sample) where the samples shown belong, which implies possible co-infection of viruses on the specific sample shown. Thus, it is difficult to associate the symptom phenotype with the detection of a single specific virus. Complete information on the individual samples can be found in Supplementary Table 1 and the description of symptoms of the associated plant hosts are summarized in Supplementary Table 9.

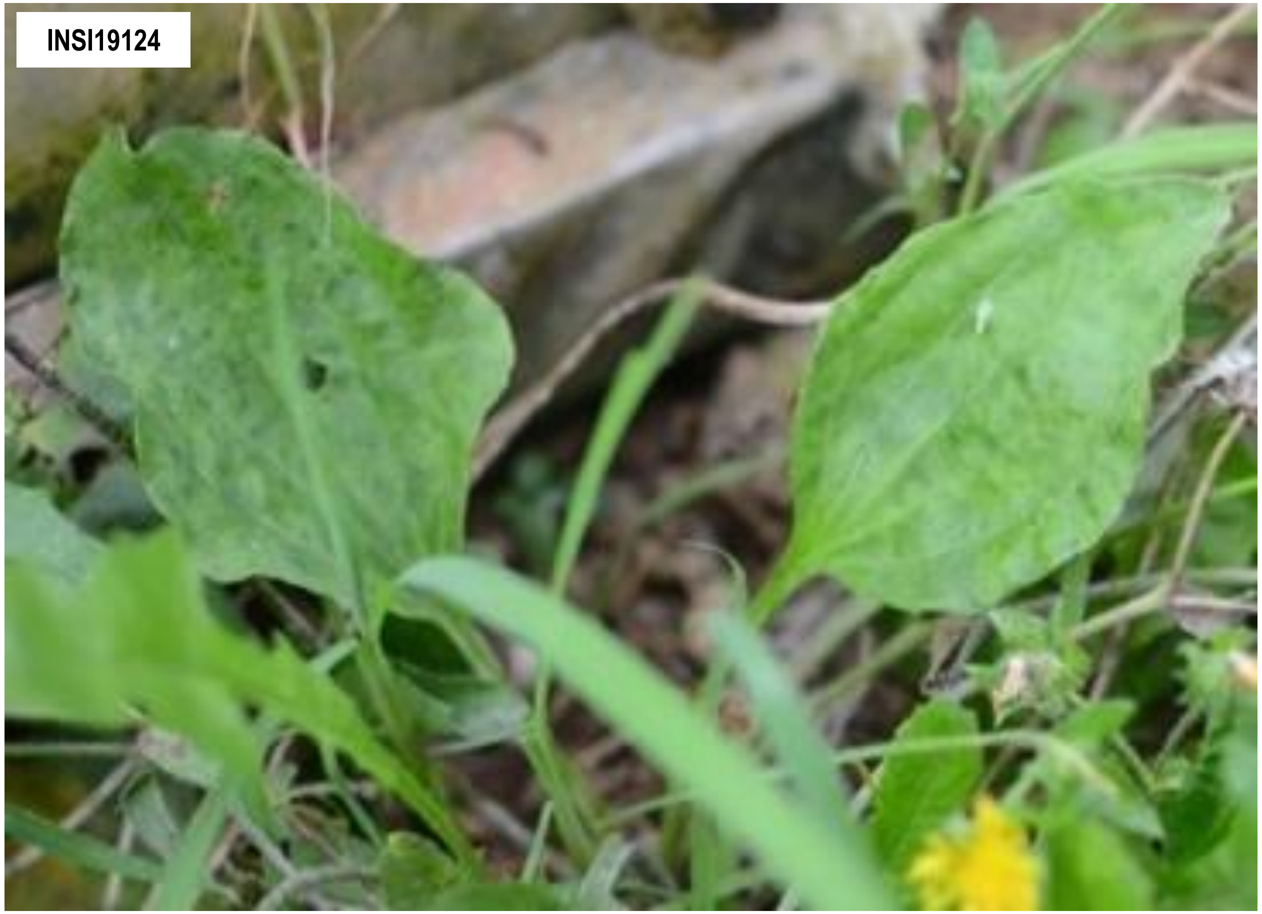


**Supplementary Figure 3-12.** Plantago tobamovirus 1. The confirmed associated plant host shown is *Plantago major* (Plantaginaceae).

**Supplementary Figure 3** (continued)**.** Representative field photos of confirmed associated plant hosts of selected viruses and putative viroids. **Note**: There are other viruses detected in the sequencing pools (*i.e.* composite sample) where the samples shown belong, which implies possible co-infection of viruses on the specific sample shown. Thus, it is difficult to associate the symptom phenotype with the detection of a single specific virus. Complete information on the individual samples can be found in Supplementary Table 1 and the description of symptoms of the associated plant hosts are summarized in Supplementary Table 9.

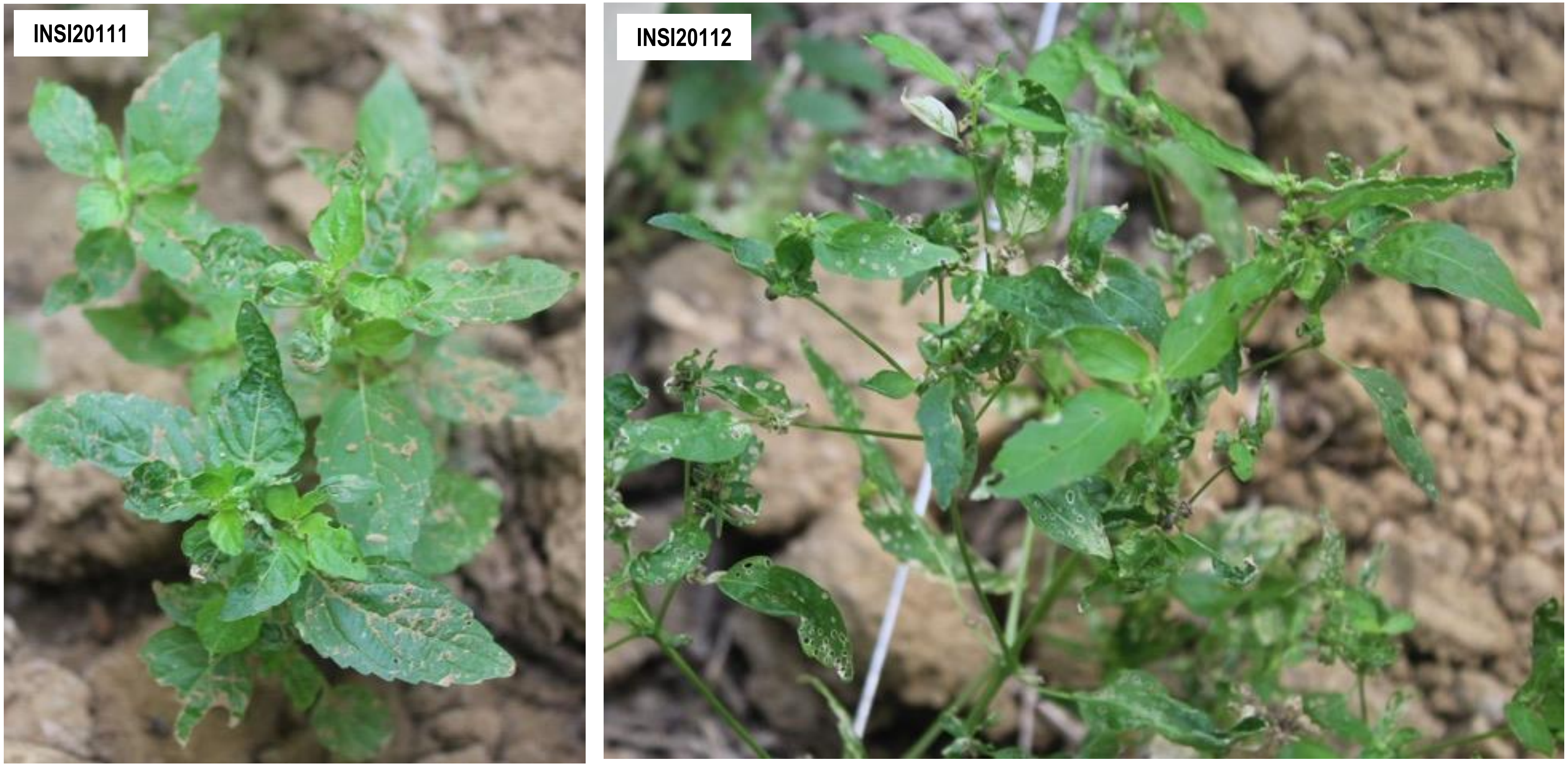


**Supplementary Figure 3-13.** Mercurialis orthotospovirus 1. The confirmed associated plant hosts shown are *Mercurialis annua*.

**Supplementary Figure 3** (continued)**.** Representative field photos of confirmed associated plant hosts of selected viruses and putative viroids. **Note**: There are other viruses detected in the sequencing pools (*i.e.* composite sample) where the samples shown belong, which implies possible co-infection of viruses on the specific sample shown. Thus, it is difficult to associate the symptom phenotype with the detection of a single specific virus. Complete information on the individual samples can be found in Supplementary Table 1 and the description of symptoms of the associated plant hosts are summarized in Supplementary Table 9.

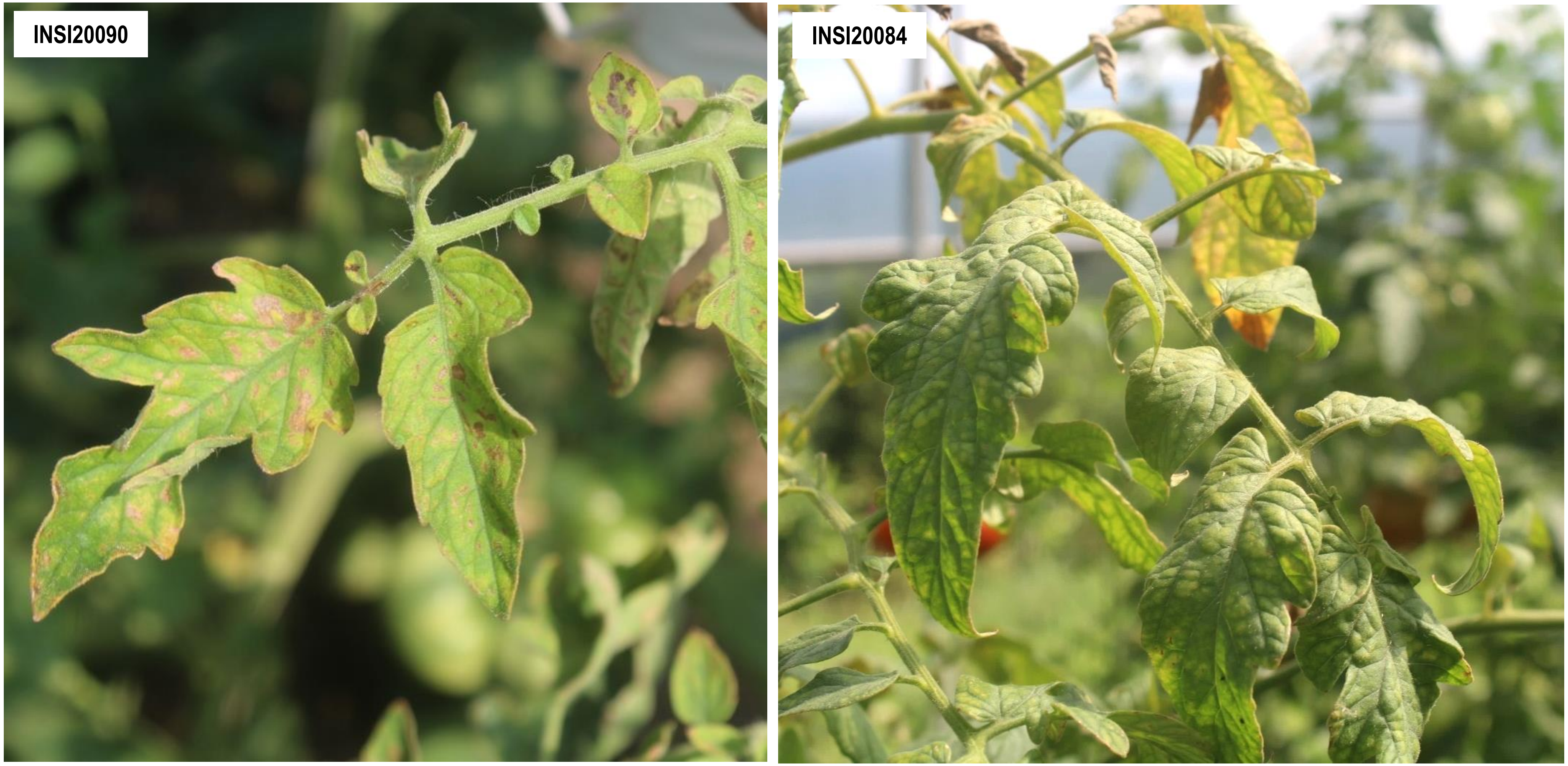


**Supplementary Figure 3-14.** Tomato associated bunya-like virus 1. The confirmed associated plant host shown is *Solanum lycopersicum*.

**Supplementary Figure 3** (continued)**.** Representative field photos of confirmed associated plant hosts of selected viruses and putative viroids. **Note**: There are other viruses detected in the sequencing pools (*i.e.* composite sample) where the samples shown belong, which implies possible co-infection of viruses on the specific sample shown. Thus, it is difficult to associate the symptom phenotype with the detection of a single specific virus. Complete information on the individual samples can be found in Supplementary Table 1 and the description of symptoms of the associated plant hosts are summarized in Supplementary Table 9.

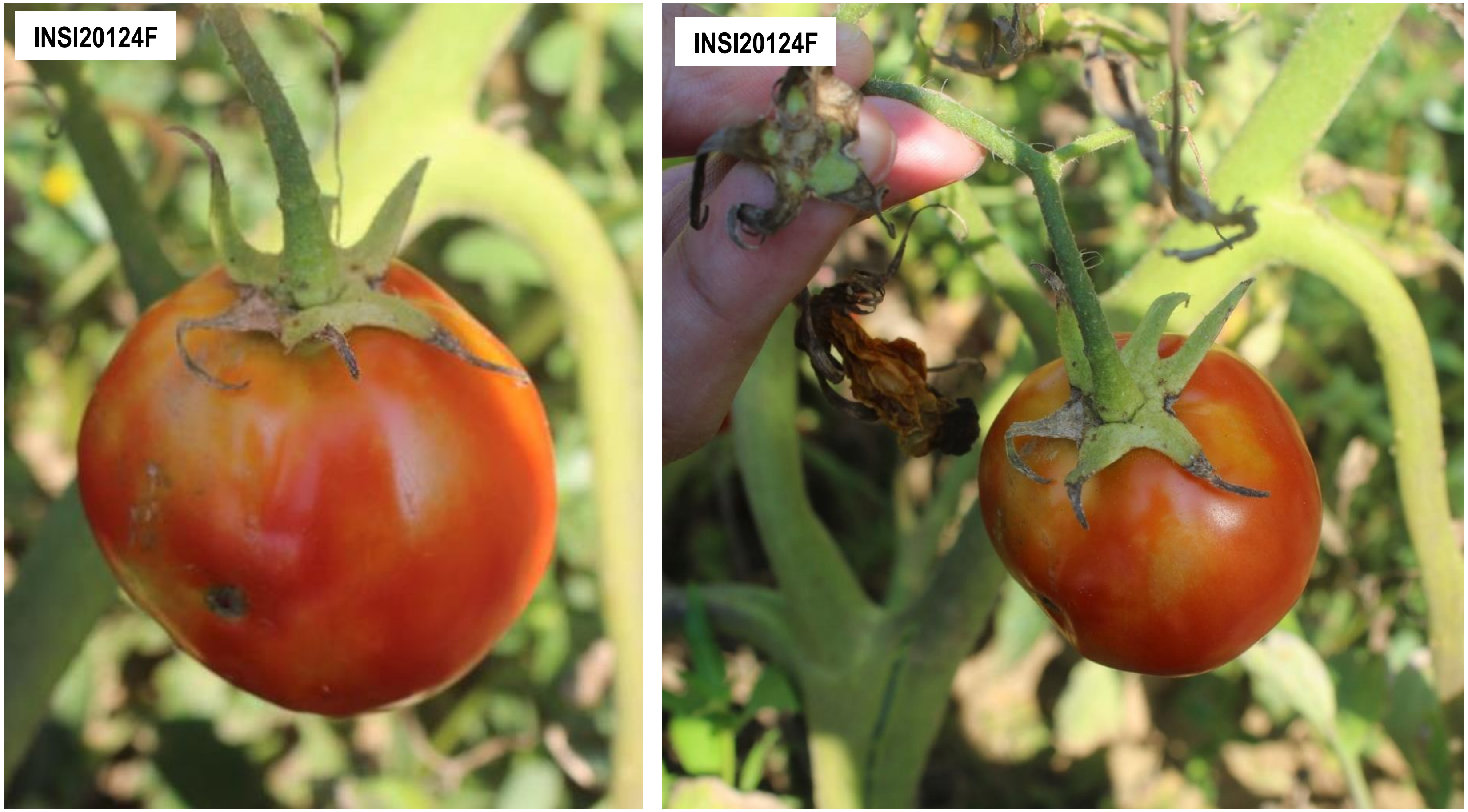


**Supplementary Figure 3-15.** Tomato vitivirus 1. The confirmed associated plant host(s) shown is/are *Solanum lycopersicum* (Solanaceae).

**Supplementary Figure 3** (continued)**.** Representative field photos of confirmed associated plant hosts of selected viruses and putative viroids. **Note**: There are other viruses detected in the sequencing pools (*i.e.* composite sample) where the samples shown belong, which implies possible co-infection of viruses on the specific sample shown. Thus, it is difficult to associate the symptom phenotype with the detection of a single specific virus. Complete information on the individual samples can be found in Supplementary Table 1 and the description of symptoms of the associated plant hosts are summarized in Supplementary Table 9.

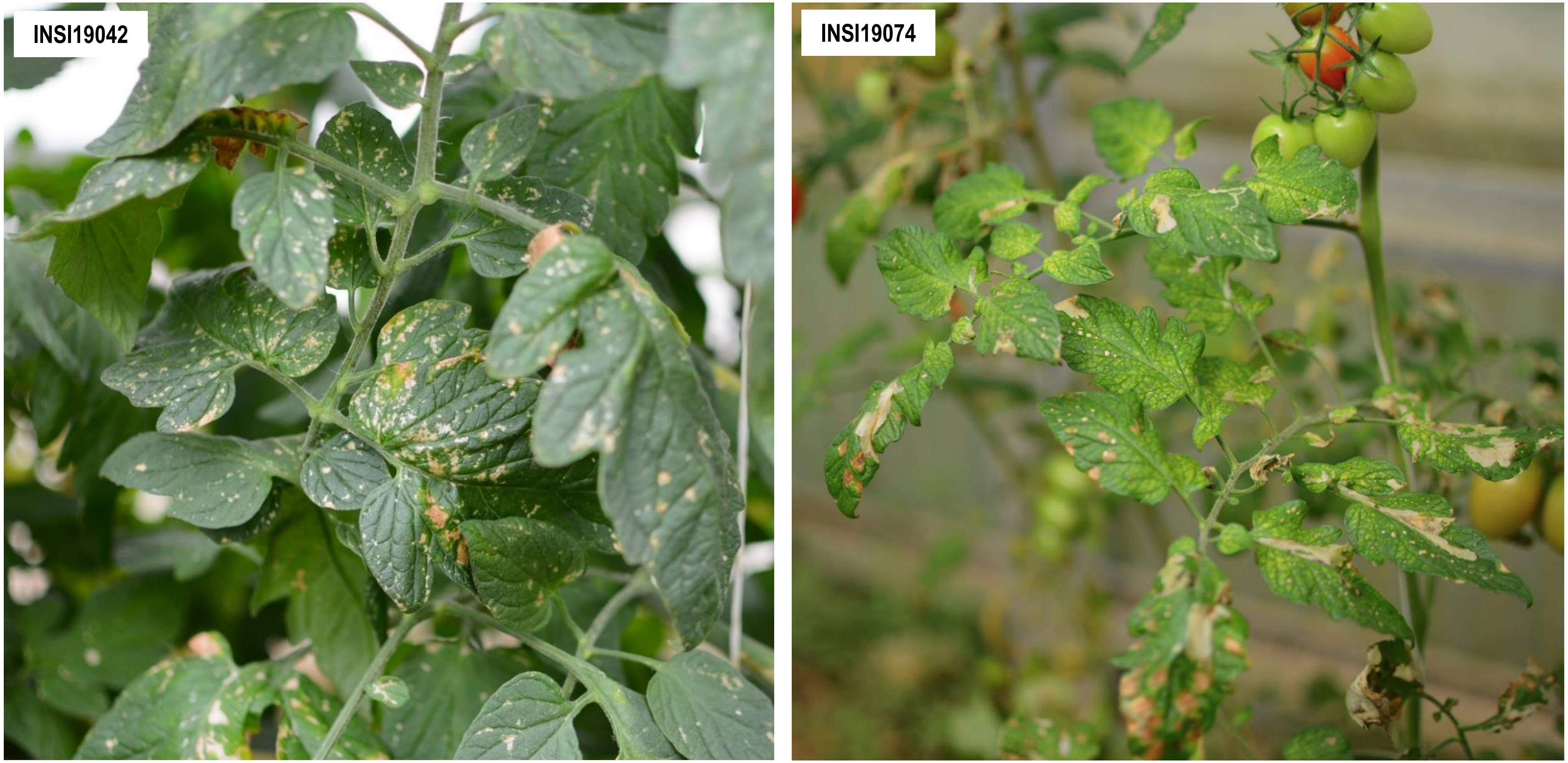


**Supplementary Figure 3-16.** Ranunculus white mottle ophiovirus. The confirmed associated plant hosts shown are tomatoes (*Solanum lycopersicum*).

**Supplementary Figure 3** (continued)**.** Representative field photos of confirmed associated plant hosts of selected viruses and putative viroids. **Note**: There are other viruses detected in the sequencing pools (*i.e.* composite sample) where the samples shown belong, which implies possible co-infection of viruses on the specific sample shown. Thus, it is difficult to associate the symptom phenotype with the detection of a single specific virus. Complete information on the individual samples can be found in Supplementary Table 1 and the description of symptoms of the associated plant hosts are summarized in Supplementary Table 9.

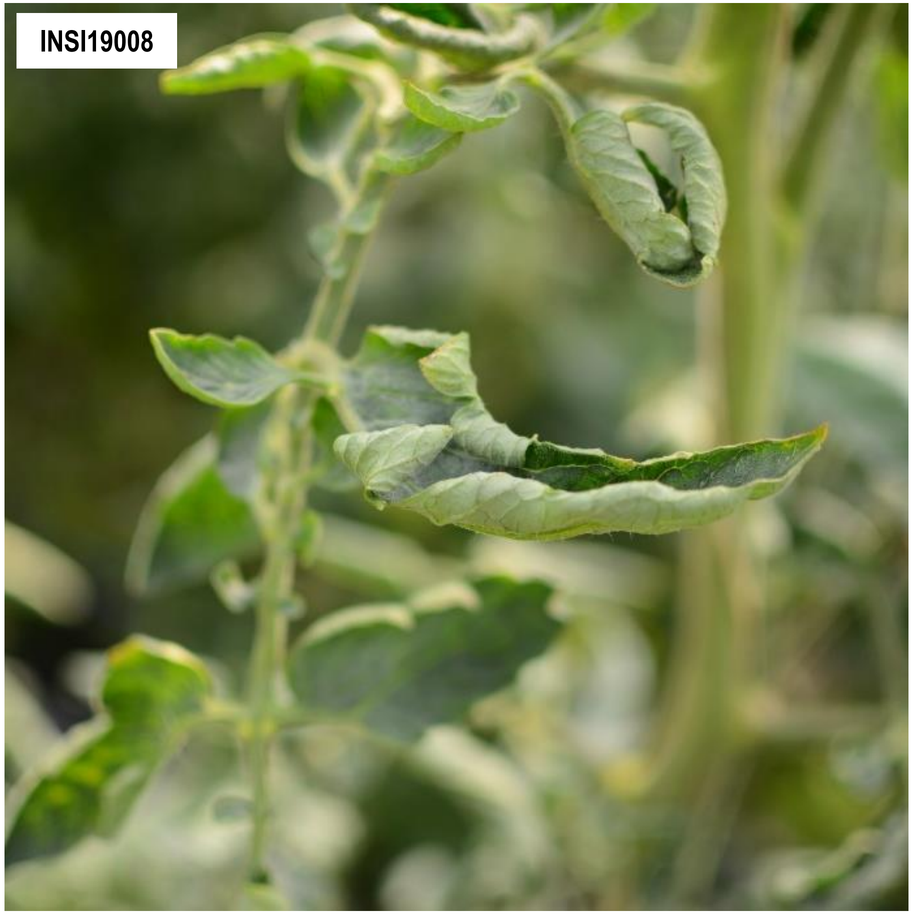

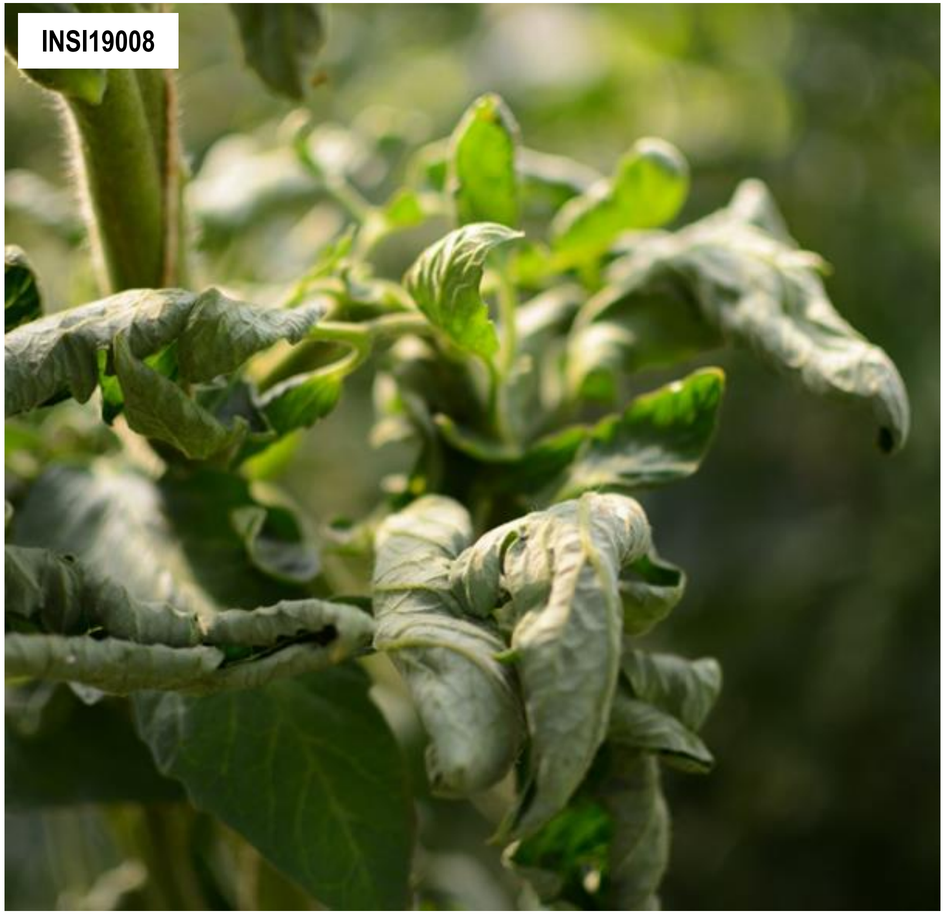


**Supplementary Figure 3-17.** Tomato betanucleorhabdovirus 1. The confirmed associated plant host shown is tomato (*Solanum lycopersicum*).

**Supplementary Figure 3** (continued)**.** Representative field photos of confirmed associated plant hosts of selected viruses and putative viroids. **Note**: There are other viruses detected in the sequencing pools (*i.e.* composite sample) where the samples shown belong, which implies possible co-infection of viruses on the specific sample shown. Thus, it is difficult to associate the symptom phenotype with the detection of a single specific virus. Complete information on the individual samples can be found in Supplementary Table 1 and the description of symptoms of the associated plant hosts are summarized in Supplementary Table 9.

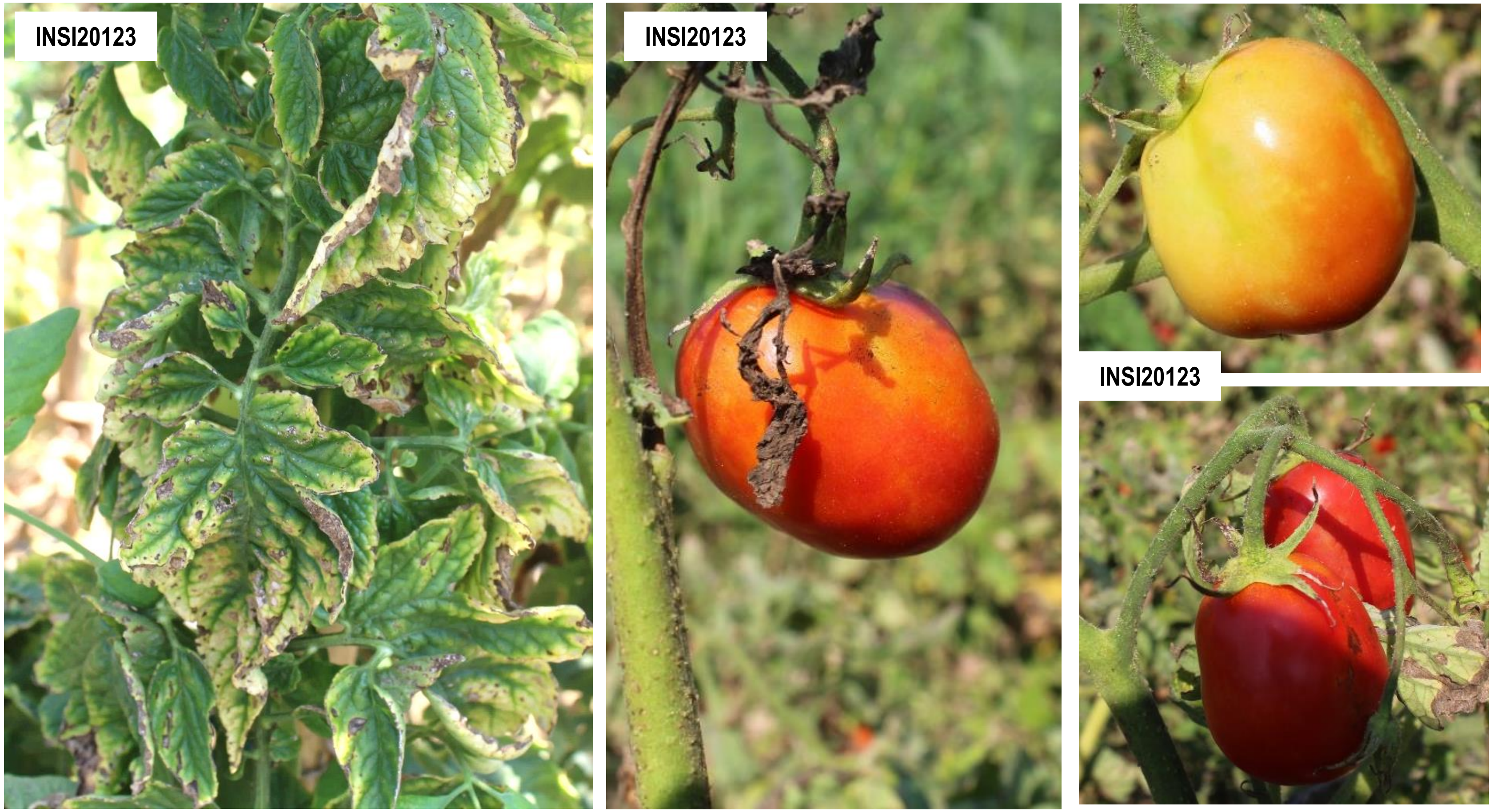


**Supplementary Figure 3-18.** Tomato betanucleorhabdovirus 2. The confirmed associated plant host shown is tomato (*Solanum lycopersicum*).

**Supplementary Figure 3** (continued)**.** Representative field photos of confirmed associated plant hosts of selected viruses and putative viroids. **Note**: There are other viruses detected in the sequencing pools (*i.e.* composite sample) where the samples shown belong, which implies possible co-infection of viruses on the specific sample shown. Thus, it is difficult to associate the symptom phenotype with the detection of a single specific virus. Complete information on the individual samples can be found in Supplementary Table 1 and the description of symptoms of the associated plant hosts are summarized in Supplementary Table 9.

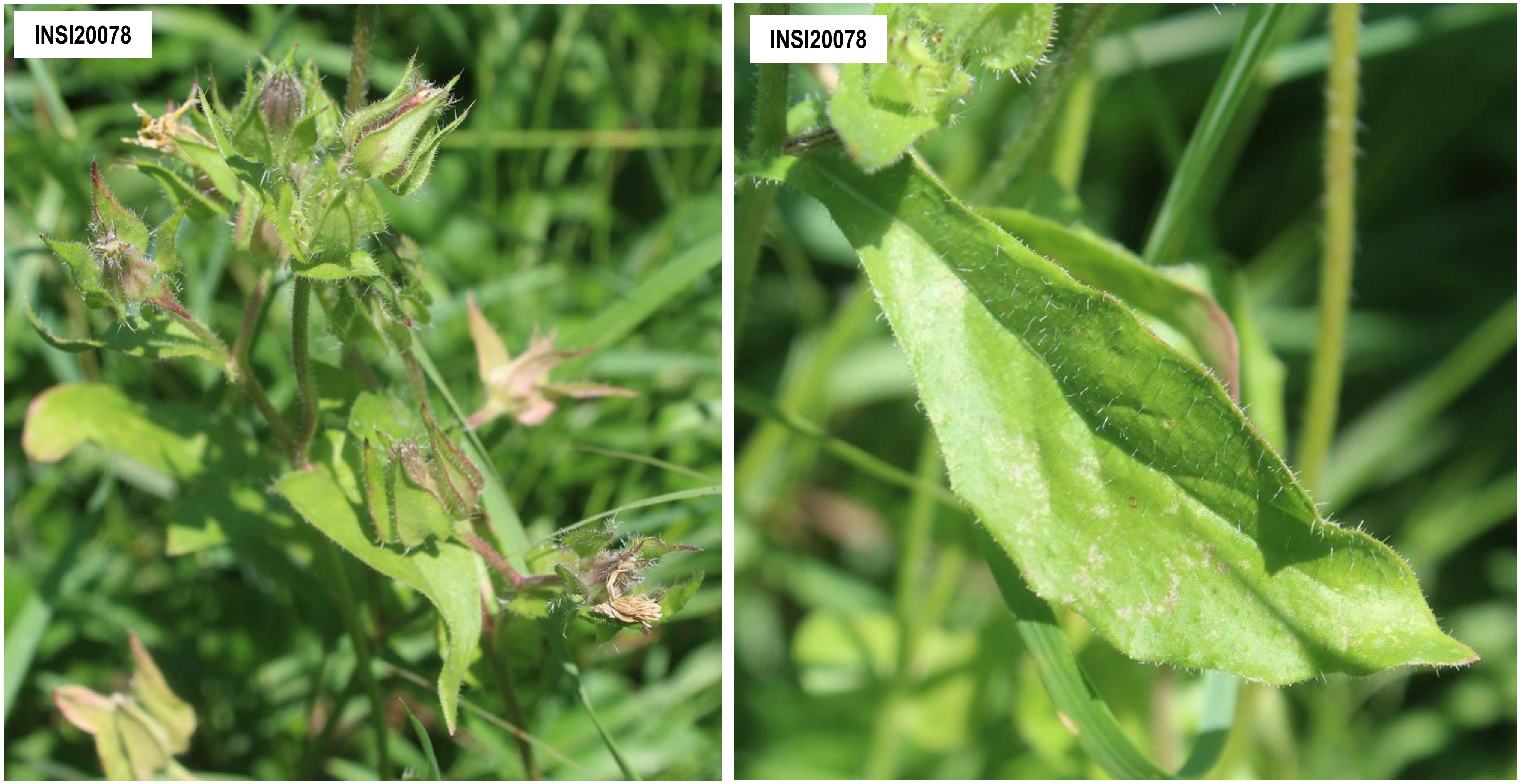


**Supplementary Figure 3-19.** Picris betanucleorhabdovirus 1. The confirmed associated plant host shown is *Picris echoides* (Asteraceae). The sample shown is also confirmed associated host of Cichorium alphacarmovirus 1 and Prunus virus I.

**Supplementary Figure 3** (continued)**.** Representative field photos of confirmed associated plant hosts of selected viruses and putative viroids. **Note**: There are other viruses detected in the sequencing pools (*i.e.* composite sample) where the samples shown belong, which implies possible co-infection of viruses on the specific sample shown. Thus, it is difficult to associate the symptom phenotype with the detection of a single specific virus. Complete information on the individual samples can be found in Supplementary Table 1 and the description of symptoms of the associated plant hosts are summarized in Supplementary Table 9.

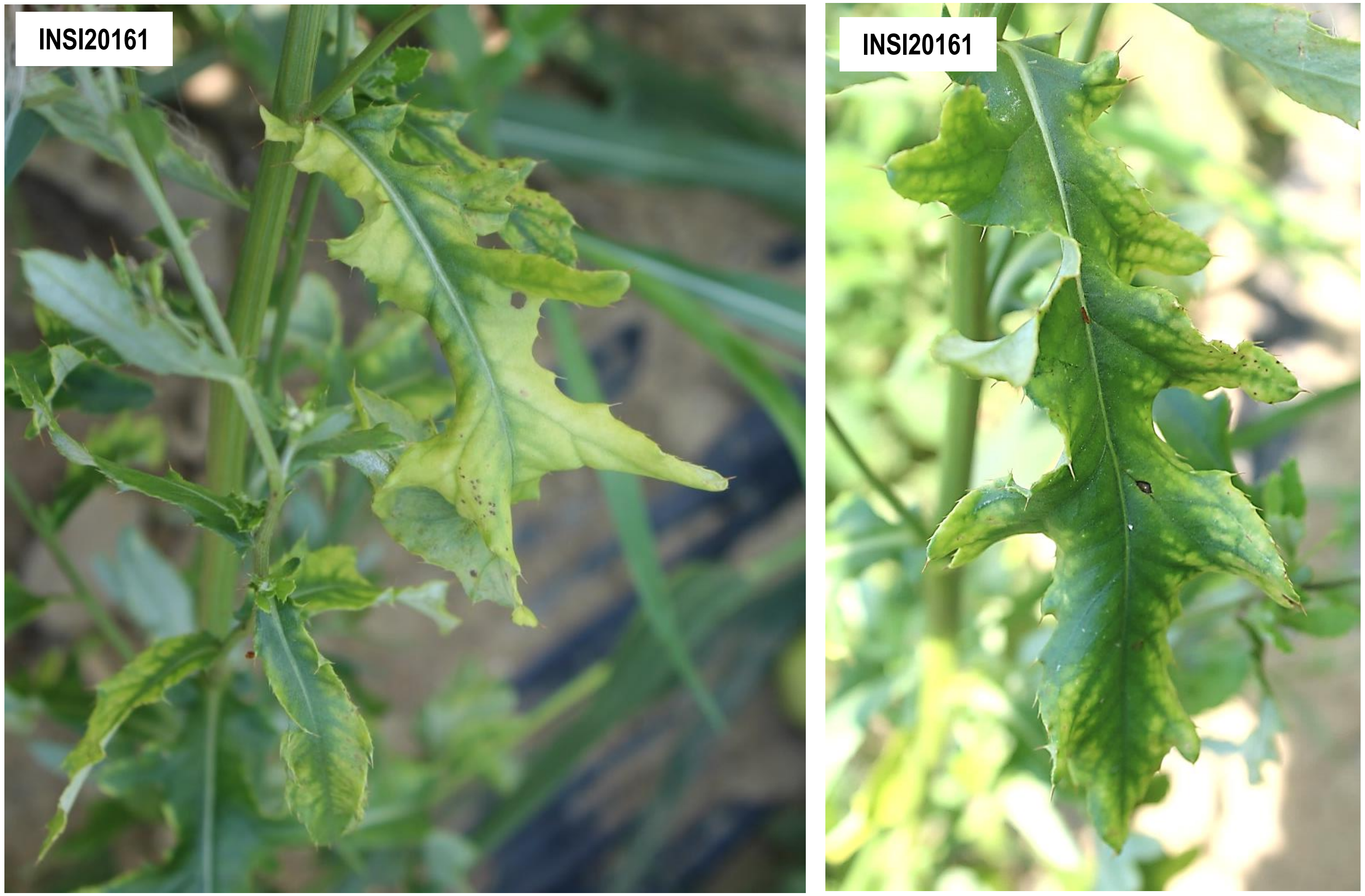


**Supplementary Figure 3-20.** Cirsium cytorhabdovirus 1. The confirmed associated plant host shown is *Cirsium arvense* (Asteraceae).

**Supplementary Figure 3** (continued)**.** Representative field photos of confirmed associated plant hosts of selected viruses and putative viroids. **Note**: There are other viruses detected in the sequencing pools (*i.e.* composite sample) where the samples shown belong, which implies possible co-infection of viruses on the specific sample shown. Thus, it is difficult to associate the symptom phenotype with the detection of a single specific virus. Complete information on the individual samples can be found in Supplementary Table 1 and the description of symptoms of the associated plant hosts are summarized in Supplementary Table 9.

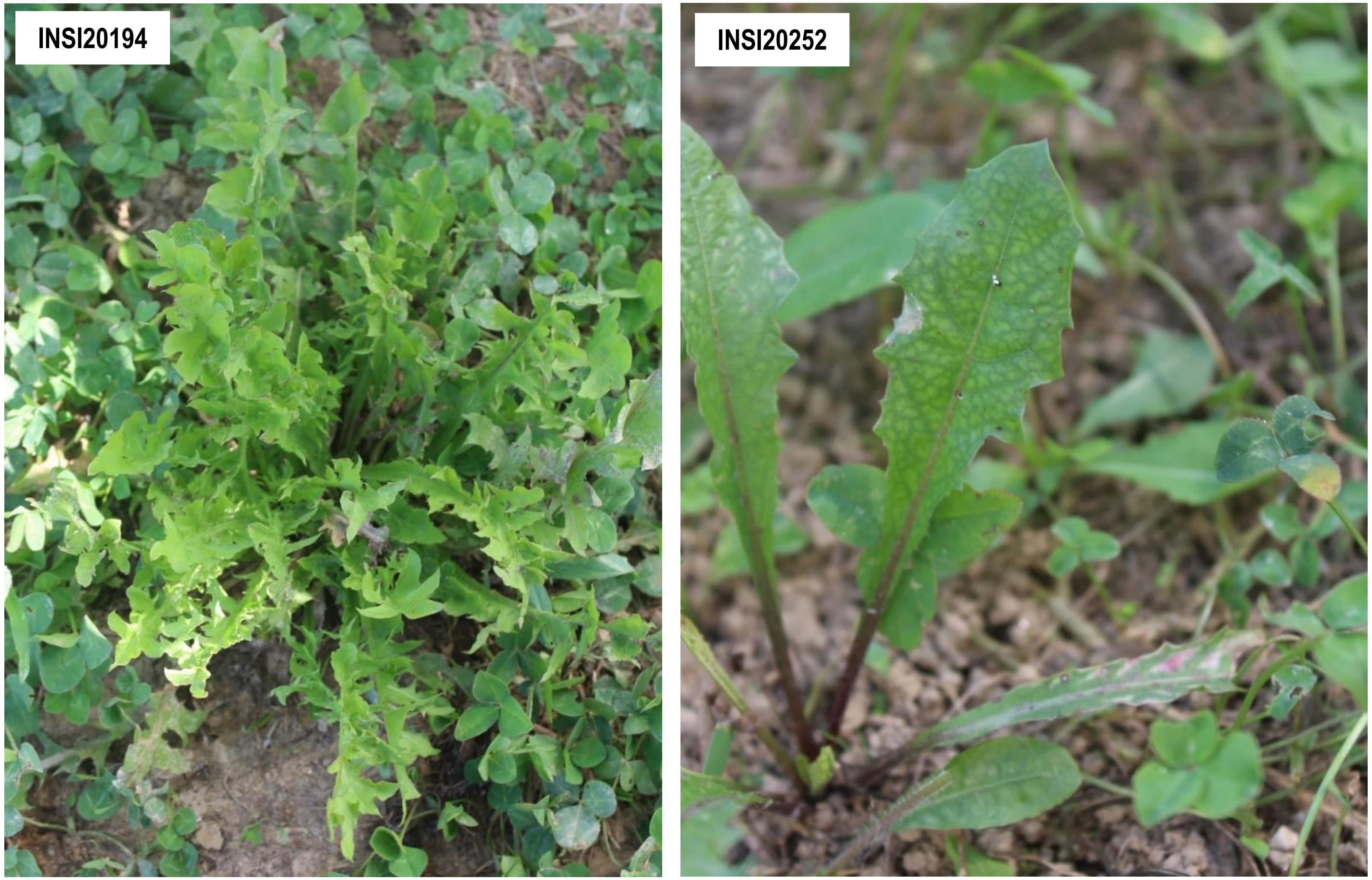


**Supplementary Figure 3-21.** Taraxacum betanucleorhabdovirus 1. The confirmed associated plant host shown is *Taraxacum officinale* (Asteraceae). Sample INSI20194 is also confirmed associated host of Taraxacum cytorhabdovirus 1, while sample INSI20252 is also confirmed associated host of Taraxacum viroid-like circular RNA 1.

**Supplementary Figure 3** (continued)**.** Representative field photos of confirmed associated plant hosts of selected viruses and putative viroids. **Note**: There are other viruses detected in the sequencing pools (*i.e.* composite sample) where the samples shown belong, which implies possible co-infection of viruses on the specific sample shown. Thus, it is difficult to associate the symptom phenotype with the detection of a single specific virus. Complete information on the individual samples can be found in Supplementary Table 1 and the description of symptoms of the associated plant hosts are summarized in Supplementary Table 9.

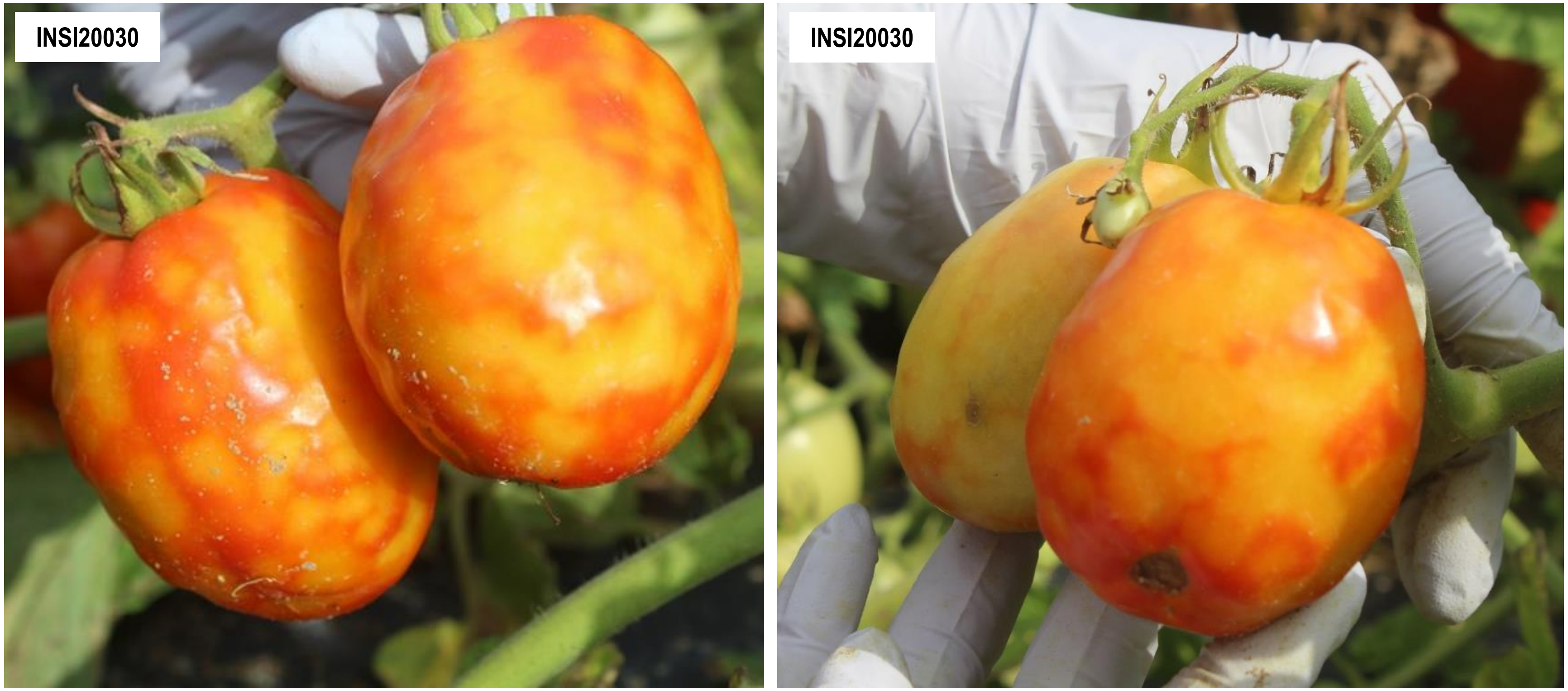


**Supplementary Figure 3-22.** Tomato alphanucleorhabdovirus 1. The confirmed associated plant host shown is tomato (*Solanum lycopersicum*).

**Supplementary Figure 3** (continued)**.** Representative field photos of confirmed associated plant hosts of selected viruses and putative viroids. **Note**: There are other viruses detected in the sequencing pools (*i.e.* composite sample) where the samples shown belong, which implies possible co-infection of viruses on the specific sample shown. Thus, it is difficult to associate the symptom phenotype with the detection of a single specific virus. Complete information on the individual samples can be found in Supplementary Table 1 and the description of symptoms of the associated plant hosts are summarized in Supplementary Table 9.

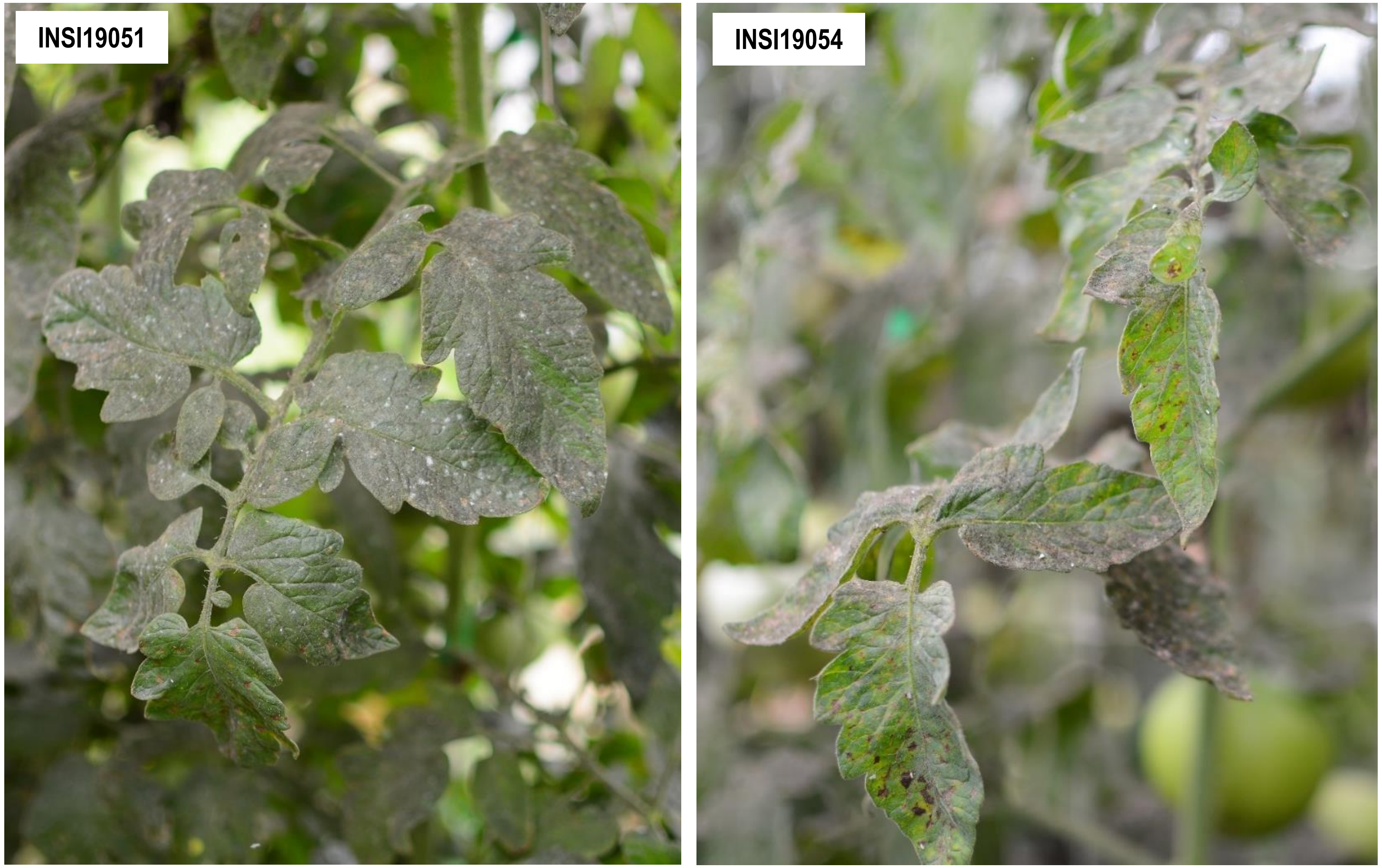


**Supplementary Figure 3-23.** Leveillula taurica associated rhabdo-like virus 1. The confirmed associated plant host shown is *Solanum lycopersicum* (Solanaceae).

**Supplementary Figure 3** (continued)**.** Representative field photos of confirmed associated plant hosts of selected viruses and putative viroids. **Note**: There are other viruses detected in the sequencing pools (*i.e.* composite sample) where the samples shown belong, which implies possible co-infection of viruses on the specific sample shown. Thus, it is difficult to associate the symptom phenotype with the detection of a single specific virus. Complete information on the individual samples can be found in Supplementary Table 1 and the description of symptoms of the associated plant hosts are summarized in Supplementary Table 9.

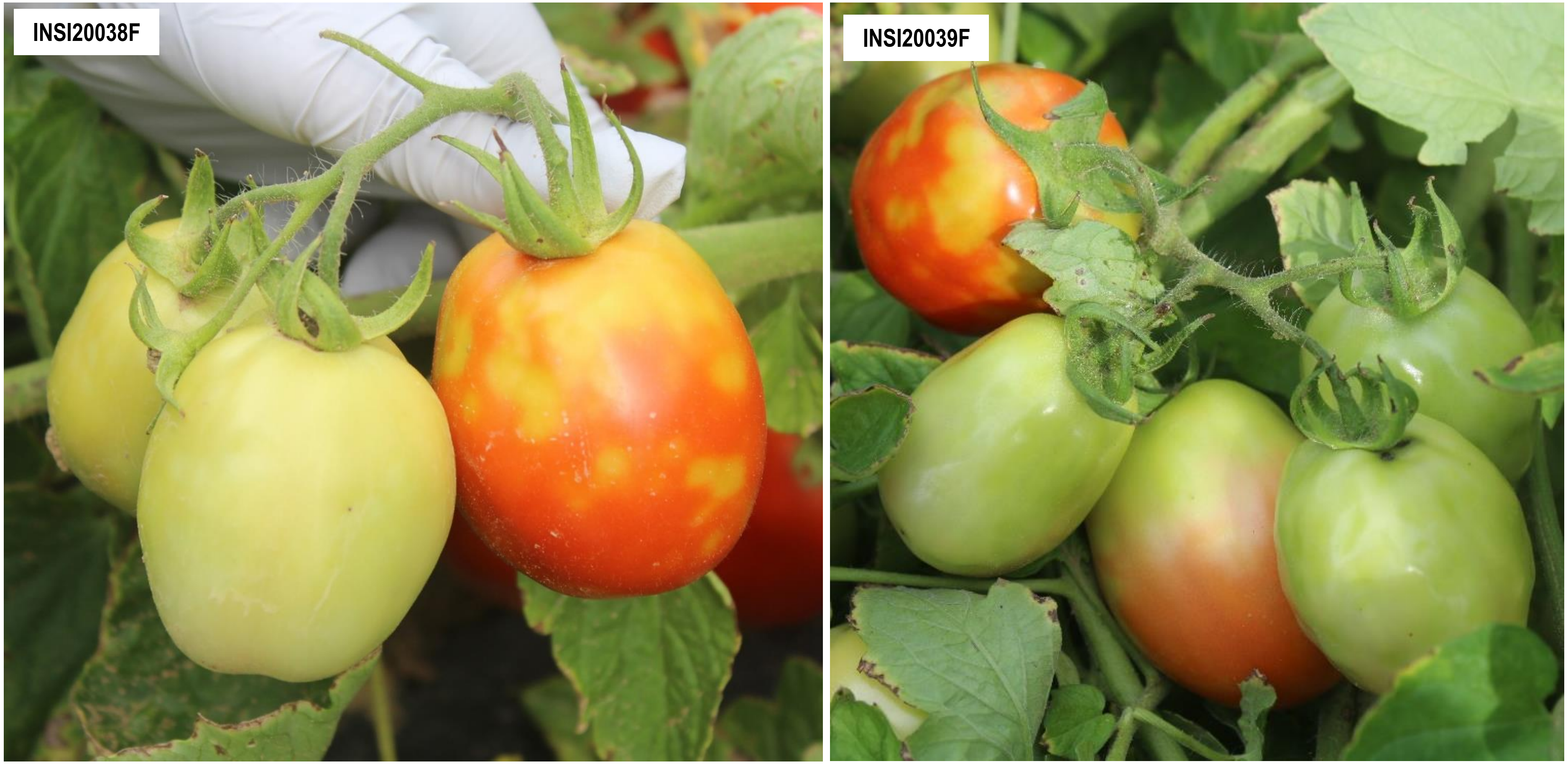


**Supplementary Figure 3-24.** Eggplant mottled dwarf alphanucleorhabdovirus. The confirmed associated plant host shown is *Solanum lycopersicum* (Solanaceae).

**Supplementary Figure 3** (continued)**.** Representative field photos of confirmed associated plant hosts of selected viruses and putative viroids. **Note**: There are other viruses detected in the sequencing pools (*i.e.* composite sample) where the samples shown belong, which implies possible co-infection of viruses on the specific sample shown. Thus, it is difficult to associate the symptom phenotype with the detection of a single specific virus. Complete information on the individual samples can be found in Supplementary Table 1 and the description of symptoms of the associated plant hosts are summarized in Supplementary Table 9.

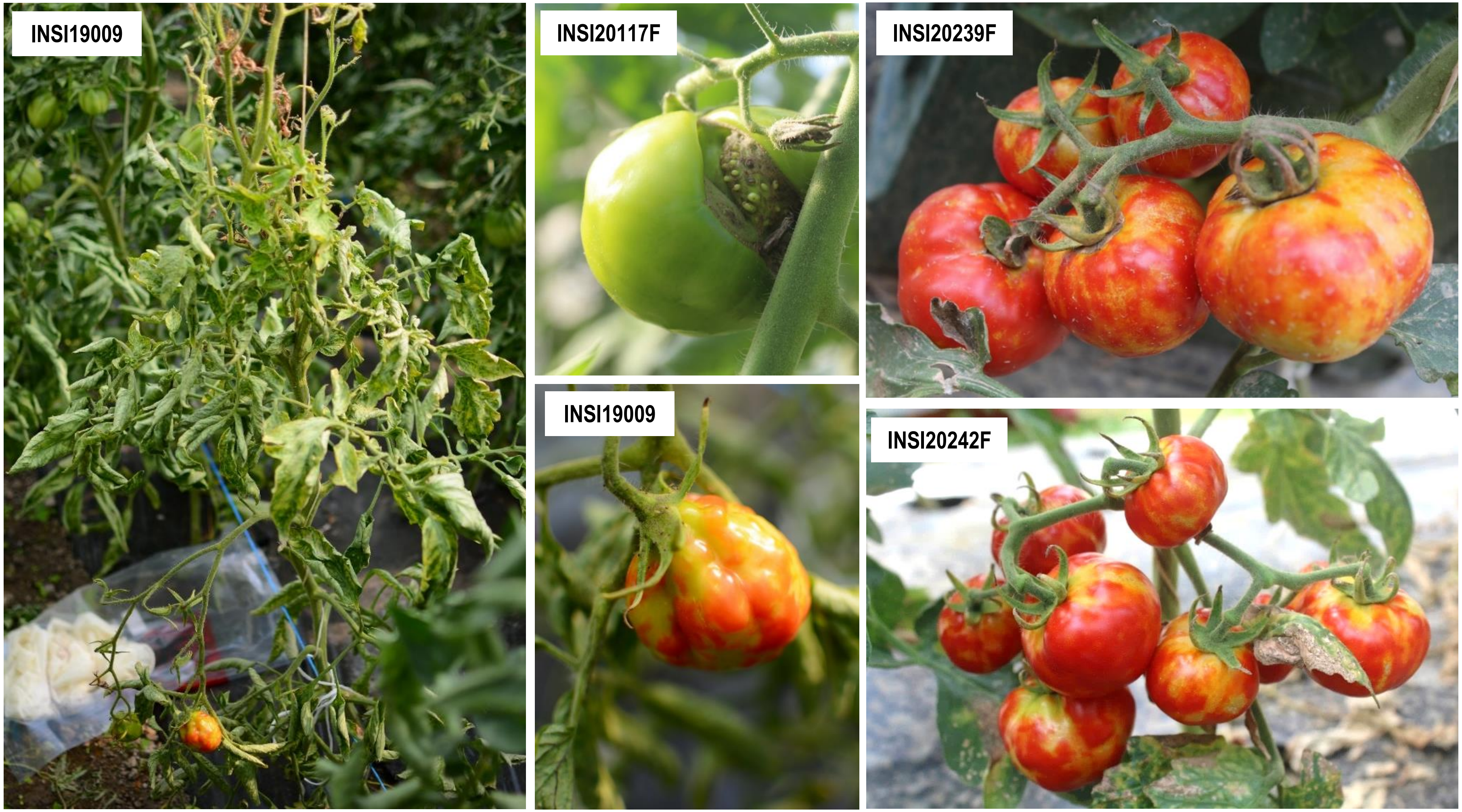


**Supplementary Figure 3-25.** Physostegia chlorotic mottle alphanucleorhabdovirus. All confirmed associated plant hosts shown are tomatoes (*Solanum lycopersicum*).

**Supplementary Figure 3** (continued)**.** Representative field photos of confirmed associated plant hosts of selected viruses and putative viroids. **Note**: There are other viruses detected in the sequencing pools (*i.e.* composite sample) where the samples shown belong, which implies possible co-infection of viruses on the specific sample shown. Thus, it is difficult to associate the symptom phenotype with the detection of a single specific virus. Complete information on the individual samples can be found in Supplementary Table 1 and the description of symptoms of the associated plant hosts are summarized in Supplementary Table 9.

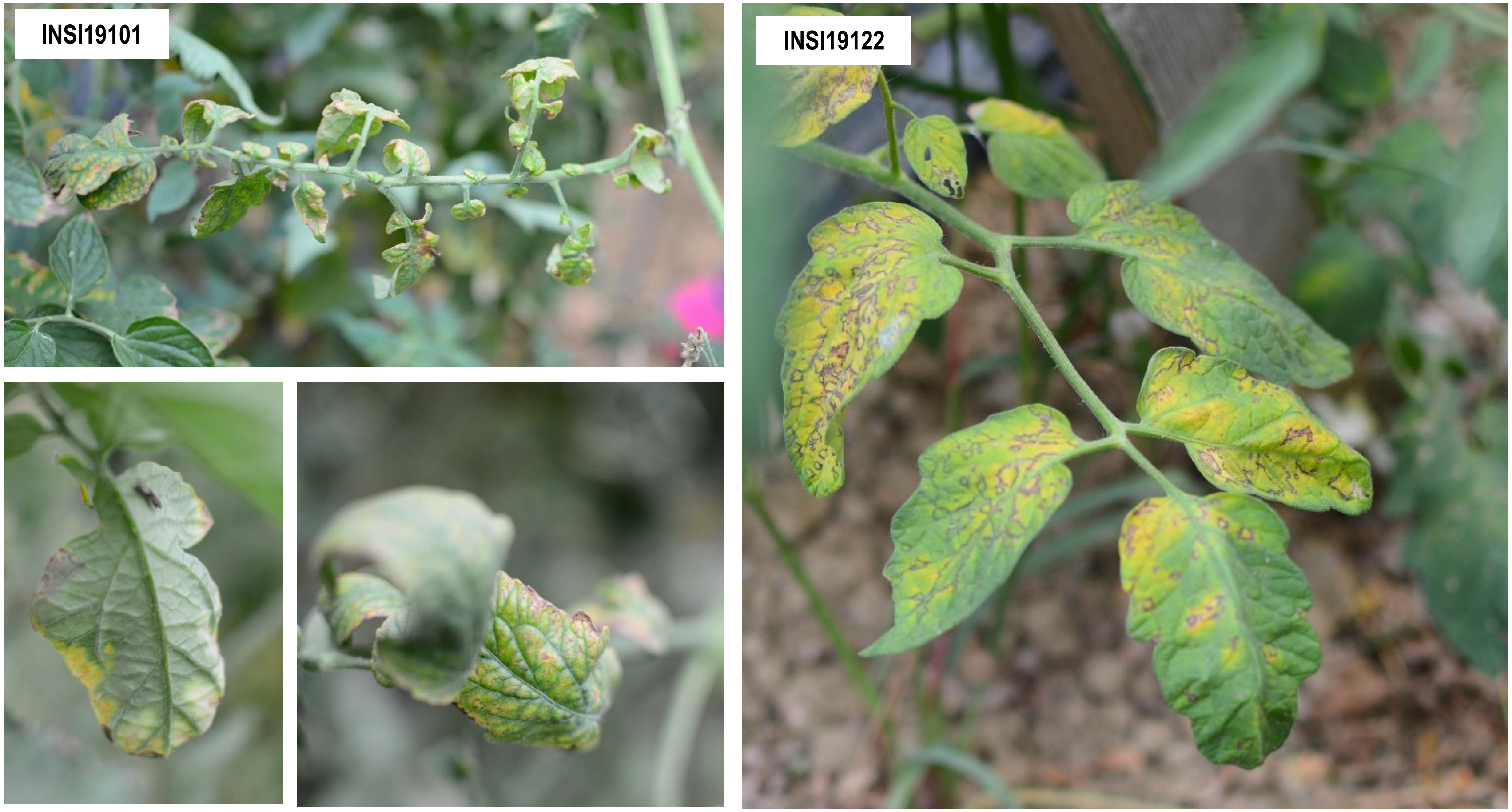


**Supplementary Figure 3-26.** Tomato fruit blotch virus and Tomato matilda virus co-infected samples.
The confirmed associated plant host shown is *Solanum lycopersicum* (Solanaceae).

**Supplementary Figure 3** (continued)**.** Representative field photos of confirmed associated plant hosts of selected viruses and putative viroids. **Note**: There are other viruses detected in the sequencing pools (*i.e.* composite sample) where the samples shown belong, which implies possible co-infection of viruses on the specific sample shown. Thus, it is difficult to associate the symptom phenotype with the detection of a single specific virus. Complete information on the individual samples can be found in Supplementary Table 1 and the description of symptoms of the associated plant hosts are summarized in Supplementary Table 9.

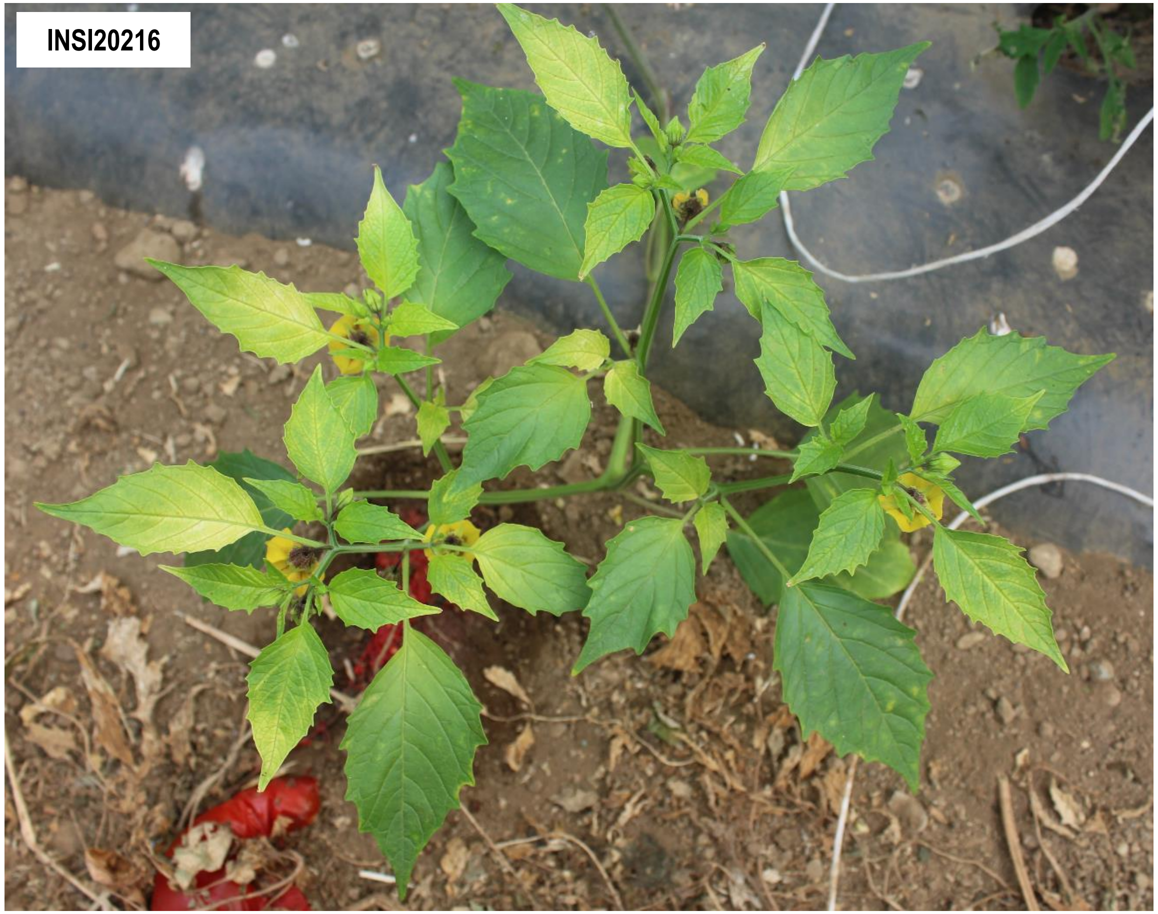


**Supplementary Figure 3-27.** Solanum nigrum ilarvirus 1. The confirmed associated plant host shown is *Physalis* sp. (Solanaceae).

**Supplementary Figure 3** (continued)**.** Representative field photos of confirmed associated plant hosts of selected viruses and putative viroids. **Note**: There are other viruses detected in the sequencing pools (*i.e.* composite sample) where the samples shown belong, which implies possible co-infection of viruses on the specific sample shown. Thus, it is difficult to associate the symptom phenotype with the detection of a single specific virus. Complete information on the individual samples can be found in Supplementary Table 1 and the description of symptoms of the associated plant hosts are summarized in Supplementary Table 9.
