## Additional-File-06_Phylogenetic-Analyses-Trees for "In-depth study of tomato and weed viromes reveals undiscovered plant virus diversity in an agroecosystem"

### SUPPLEMENTARY INFORMATION

### Additional File 06

**Supplementary Table 10.** Model selection and other parameters used for phylogenetic analyses. **Supplementary Figure 4.** Phylogenetic trees up to the virus family or order level showing relationships of identified known and novel virus species in this study.

**Supplementary Table 10.** Model selection and other parameters used for maximum likelihood phylogenetic analyses.

Abbreviations used: aa (amino acid), nt (nucleotide). RdRp (RNA-dependent RNA polymerase), Mtr (methyl transferase), Hel (helicase), CP (coat protein), RT (reverse transcriptase). Abbreviations of substitution models used are indicated in the succeeding paragraph.

Notes: *Substitution model was chosen based on the lowest Bayesian Information Criteria (BIC) value. [a] subtree of a larger but identical tree indicated in parentheses, [b] identical and/or collapsed version of a tree indicated in parentheses.

| Virus family / families (or virus order) | Figure(s) where tree is shown (trees in supplementary figures are numbered consecutively) | Gene or genome segment used | Number of sequences in the dataset | Alignment and trimming programs used | Number of positions in final alignment | Substitution model selected* |
|---|---|---|---|---|---|---|
| *Tospoviridae* and *Fimoviridae* (*Bunyavirales*) | (01) Supplementary Figure 4-01<br>Figure 4i [a,(01)] | RdRp | 65 | ClustalW, MAFFT (after manual trimming), trimAl | 376 aa | LG+G+I+F |
| *Aspiviridae* (*Serpentovirales*) | (02) Supplementary Figure 4-02<br>Figure 4k [a,(02)] | RdRp | 29 | ClustalW, MAFFT (after manual trimming), trimAl | 447 aa | LG+G+I |
| *Rhabdoviridae, Lispiviridae* (*Mononegavirales*) | (03) Supplementary Figure 4-03<br>Figure 5f [b,(03)] | RdRp | 51 | ClustalW, MAFFT (after manual trimming), trimAl | 510 aa | LG+G+I+F |
| *Kitaviridae* (*Martellivirales*) | (04) Supplementary Figure 4-04<br>Figure 6k [b,(04)] | RdRp | 28 | ClustalW, MAFFT (after manual trimming), trimAl | 583 aa | LG+G+I+F |
| *Bromoviridae* (*Martellivirales*) | (05) Supplementary Figure 4-05<br>Figure 6l [b,(05)] | RdRp | 63 | ClustalW, MAFFT (after manual trimming), trimAl | 327 aa | LG+G+I |
| *Closteroviridae* (*Martellivirales*) | (06) Supplementary Figure 4-06<br>Figure 6m [b,(06)] | RdRp | 48 | ClustalW, MAFFT (after manual trimming), trimAl | 219 aa | LG+G+I |
| *Endornaviridae* (*Martellivirales*) | (07) Supplementary Figure 4-07<br>Figure 6n [b,(07)] | RdRp | 43 | ClustalW, MAFFT (after manual trimming), trimAl | 278 aa | LG+G+I |

| | | | | | | |
|---|---|---|---|---|---|---|
| *Virgaviridae* (*Martellivirales*) | (08-A) Supplementary Figure 4-08-A<br>Figure 7h [b,(08-A)] (left tree) | Mtr, Hel | 48 | ClustalW, MAFFT (after manual trimming), trimAl | 542 aa | LG+G+I |
| | (08-B) Supplementary Figure 4-08-B<br>Figure 7h [b,(08-B)] (right tree) | CP | 48 | ClustalW, MAFFT (after manual trimming), trimAl | 68 aa | LG+G |
| | Figure 7i | full genome nt sequence | 41 | MUSCLE (and manual trimming of ends) | 6233 nt | GTR+G+I |
| *Potyviridae* (*Patatavirales*) | (09) Supplementary Figure 4-09<br>Figure 8g [b,(09)] | RdRp | 93 | ClustalW, MAFFT (after manual trimming), trimAl | 470 aa | LG+G+I |
| *Tombusviridae* (*Tolivirales*) | (10) Supplementary Figure 4-10<br>Figure 9h [b,(10)] | RdRp | 108 | ClustalW, MAFFT (after manual trimming), trimAl | 291 aa | LG+G+I |
| *Partitiviridae* (*Durnavirales*) | (11) Supplementary Figure 4-11 | RdRp | 57 | ClustalW, MAFFT (after manual trimming), trimAl | 326 aa | LG+G+I |
| *Geminiviridae* (*Geplafuvirales*) | (12) Supplementary Figure 4-12 | replicase (C1 gene) | 109 | ClustalW, MAFFT (after manual trimming), trimAl | 289 aa | LG+G |
| *Totiviridae* (*Ghabrivirales*) | (13) Supplementary Figure 4-13 | RdRp | 71 | ClustalW, MAFFT (after manual trimming), trimAl | 363 aa | LG+G+I |
| *Caulimoviridae* (*Ortelivirales*) | (14) Supplementary Figure 4-14 | RT | 57 | ClustalW, MAFFT (after manual trimming), trimAl | 220 aa | LG+G |
| *Iflaviridae* (*Picornavirales*) | (15) Supplementary Figure 4-15 | RdRp | 29 | ClustalW, MAFFT (after manual trimming), trimAl | 460 aa | LG+G+I |
| *Dicistroviridae* and *Iflaviridae* (*Picornavirales*) | (16) Supplementary Figure 4-16 | RdRp | 61 | ClustalW, MAFFT (after manual trimming), trimAl | 307 aa | LG+G |

| | | | | | | |
|---|---|---|---|---|---|---|
| *Secoviridae* (*Picornavirales*) | (17) Supplementary Figure 4-17 | RdRp | 63 | ClustalW, MAFFT (after manual trimming), trimAl | 364 aa | LG+G |
| *Solemoviridae* (*Sobelivirales*) | (18) Supplementary Figure 4-18 | RdRp | 47 | ClustalW, MAFFT (after manual trimming), trimAl | 191 aa | LG+G |
| *Tymovirales* | (19) Supplementary Figure 4-19 | RdRp | 75 | ClustalW, MAFFT (after manual trimming), trimAl | 293 aa | LG+G+I |
| Satellite viruses (*Albetovirus, Aumaivirus*) | (20) Supplementary Figure 4-20 | CP | 10 | ClustalW, MAFFT (after manual trimming), trimAl | 109 aa | LG+G |

**Details of multiple sequence alignments and phylogenetic tree construction:**

Representative viral RefSeq genomes of ICTV-recognized species, which were translated to amino acid sequences, and selected best hit in BLASTp searches (E-val<$10^{-4}$) of the viruses of interest, were gathered for the phylogenetic analyses. When applicable, outgroup virus sequences were chosen based on known phylogenetic relationships to the closest related taxa (*i.e.* species of a closely related family), and with reference to phylogenetic trees published in peer-reviewed articles. Multiple sequence alignments were initially done using ClustalW (Thompson et al., 1994), and after manual trimming of unaligned ends, MAFFT alignment was implemented (Katoh and Standley, 2013). Thereafter, further trimming of the alignments with trimAl (Capella-Gutiérrez et al., 2009) was performed. Highly divergent sequences (*i.e.* <30%) based on overall pairwise amino acid similarity, examined in SDT v. 1.2 (Muhire et al., 2014), were removed from the final alignment prior to further analyses. In the case of tobamoviruses, a nucleotide-based phylogenetic analyses was done on full genomes. Genomes of different isolates of turnip vein-clearing virus, ribgrass mosaic virus, wasabi mosaic virus, youcai mosaic virus and Plantago tobamovirus 1, were gathered from GenBank and viral RefSeq database and aligned using MUSCLE (Edgar, 2004). The unaligned ends were trimmed, then checked for possible recombinants before proceeding with the phylogenetic analyses with the alignments free of recombinants. Substitution model selection was done on the final alignments, with either amino acid or nucleotide alignments. The best model was chosen, based on the lowest Bayesian Information Criteria (BIC) value (Schwarz, 1978). Selected amino acid substitution models indicated in the table above are as follows: Le_Gascuel_2008 model (Le and Gascuel, 2008) with discrete Gamma distribution with 5 rate categories (LG+G), which may also assume that a certain fraction of sites are evolutionarily invariable (LG+G+I) and may account for amino acid frequencies (LG+G+I+F). For the tobamovirus dataset, a General Time Reversible (GTR) model was selected, with discrete gamma distribution (+G) and evolutionarily invariable sites (+I). The final phylogenetic trees after the analyses have the highest log likelihood value inferred with 1000 bootstrap replicates (Felsenstein, 1985). In the succeeding phylogenetic trees shown in **Supplementary Figure 4**, the percentage of replicate trees in which the associated taxa clustered together in the bootstrap test is shown next to the branches, with only >0.6 (>60%) support shown. The trees are drawn to scale for easier visualization, with branch lengths (scale bar is shown) measured in the number of amino acid (or nucleotide) substitutions per site. ClustalW alignment, manual trimming and pairwise similarity analysis were done in CLC Genomics Workbench v. 20.0 (Qiagen). MAFFT was executed in EMBL-EBI website (www.ebi.ac.uk/Tools/msa/mafft/), and trimAl was downloaded from (www.trimal.cgenomics.org), and was executed in command line, with 'automated1' option. Substitution model testing and phylogenetic tree construction were done in MEGA X (Kumar et al., 2018), and the final phylogenetic trees were annotated in Interactive Tree of Life (iToL) (Letunic and Bork, 2021).

**References for substitution models, online tools and programs used in the phylogenetic analyses:**


Capella-Gutiérrez, S., Silla-Martínez, J. M., and Gabaldón, T. (2009). trimAl: A tool for automated alignment trimming in large-scale phylogenetic analyses. *Bioinformatics* 25, 1972–1973. doi:10.1093/bioinformatics/btp348.

Edgar, R. C. (2004). MUSCLE: multiple sequence alignment with high accuracy and high throughput. *Nucleic Acids Res.* 32, 1792–1797. doi:10.1093/nar/gkh340.

Felsenstein, J. (1985). Confidence Limits on Phylogenies: An Approach Using the Bootstrap. *Evolution (N. Y).* 39, 783. doi:10.2307/2408678.

Katoh, K., and Standley, D. M. (2013). MAFFT multiple sequence alignment software version 7: Improvements in performance and usability. *Mol. Biol. Evol.* 30, 772–780. doi:10.1093/molbev/mst010.

Kumar, S., Stecher, G., Li, M., Knyaz, C., and Tamura, K. (2018). MEGA X: Molecular Evolutionary Genetics Analysis across Computing Platforms. *Mol. Biol. Evol.* 35, 1547–1549. doi:10.1093/molbev/msy096.

Le, S. Q., and Gascuel, O. (2008). An improved general amino acid replacement matrix. *Mol. Biol. Evol.* 25, 1307–1320. doi:10.1093/molbev/msn067.

Letunic, I., and Bork, P. (2021). Interactive Tree Of Life (iTOL) v5: an online tool for phylogenetic tree display and annotation. *Nucleic Acids Res.* 49, W293–W296. doi:10.1093/nar/gkab301.

Muhire, B. M., Varsani, A., and Martin, D. P. (2014). SDT: A Virus Classification Tool Based on Pairwise Sequence Alignment and Identity Calculation. *PLoS One* 9, e108277. doi:10.1371/journal.pone.0108277.

Schwarz, G. (1978). Estimating the Dimension of a Model. *Ann. Stat.* 6. doi:10.1214/aos/1176344136.

Thompson, J. D., Higgins, D. G., and Gibson, T. J. (1994). CLUSTAL W: Improving the sensitivity of progressive multiple sequence alignment through sequence weighting, position-specific gap penalties and weight matrix choice. *Nucleic Acids Res.* 22, 4673–4680. doi:10.1093/nar/22.22.4673.

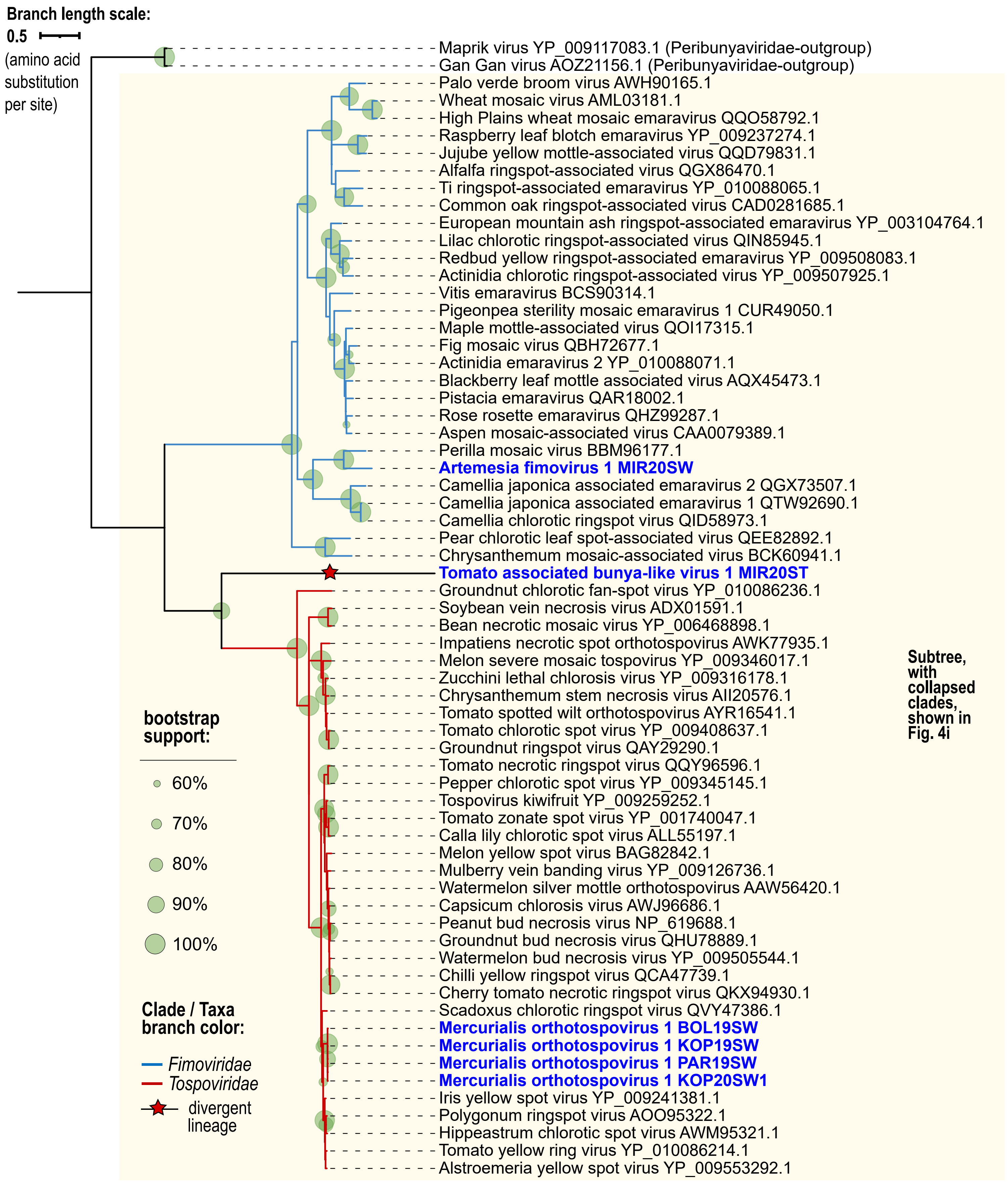


**Supplementary Figure 4-01.** Maximum likelihood phylogenetic tree based on the alignment of conserved RNA-dependent RNA polymerase amino acid sequence of virus species under *Tospoviridae* and *Fimoviridae* (*Bunyavirales*).

**Supplementary Figure 4.** Phylogenetic trees up to the virus family (or order) level showing relationships of identified known (in bold black font) and novel (in bold blue font) virus species from this study, with the isolate names indicated. GenBank accession numbers for the viruses from this study can be found in Supplementary Table 5, while GenBank accession number (protein) are indicated for known viruses in the tree. All details of phylogenetic analyses can be found above in Supplementary Table 10.

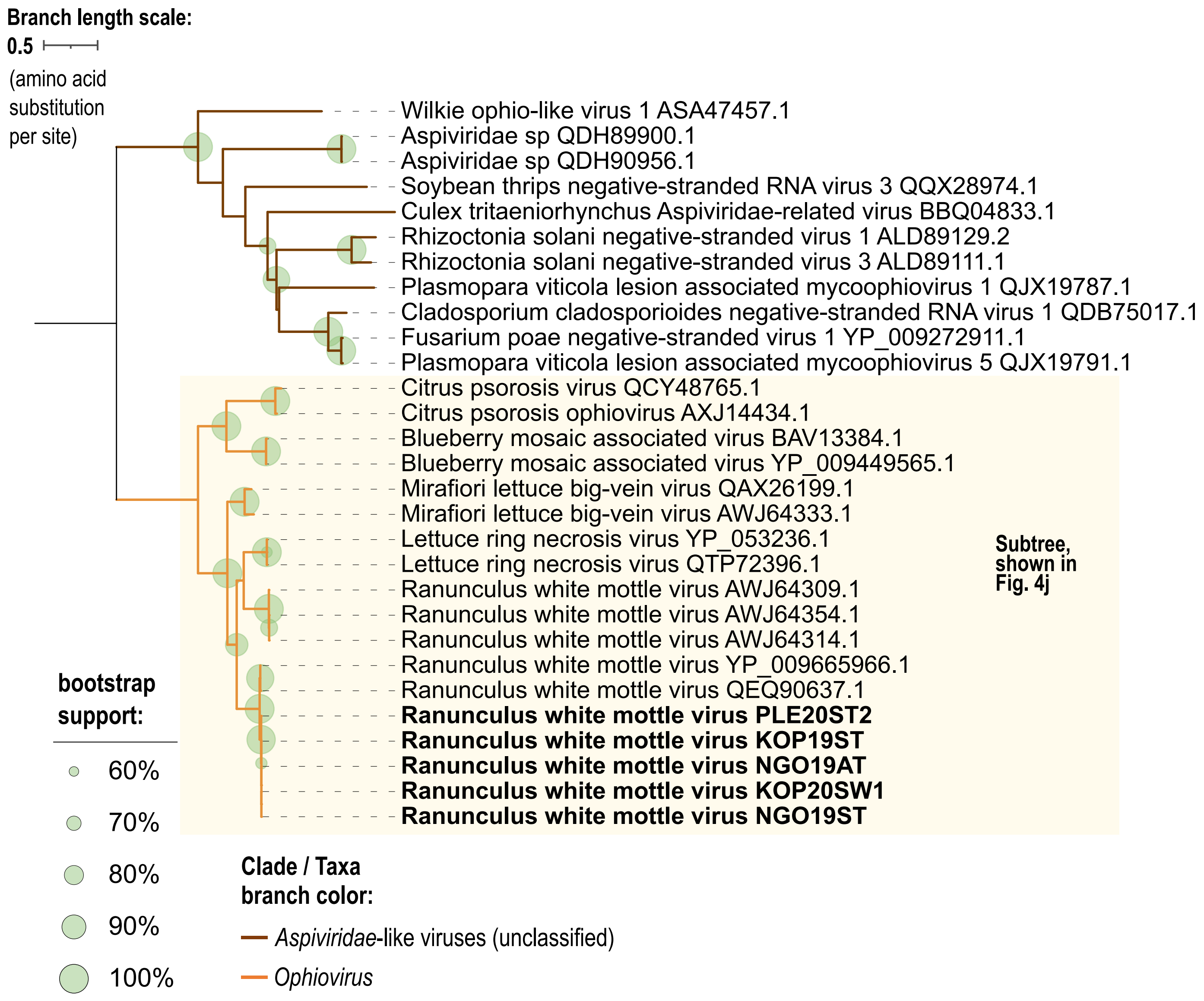


**Supplementary Figure 4-02.** Maximum likelihood phylogenetic tree based on the alignment of conserved RNA-dependent RNA polymerase amino acid sequence of virus species under family *Aspiviridae* (*Serpentovirales*).

**Supplementary Figure 4.** Phylogenetic trees up to the virus family (or order) level showing relationships of identified known (in bold black font) and novel (in bold blue font) virus species from this study, with the isolate names indicated. GenBank accession numbers for the viruses from this study can be found in Supplementary Table 5, while GenBank accession number (protein) are indicated for known viruses in the tree. All details of phylogenetic analyses can be found above in Supplementary Table 10.

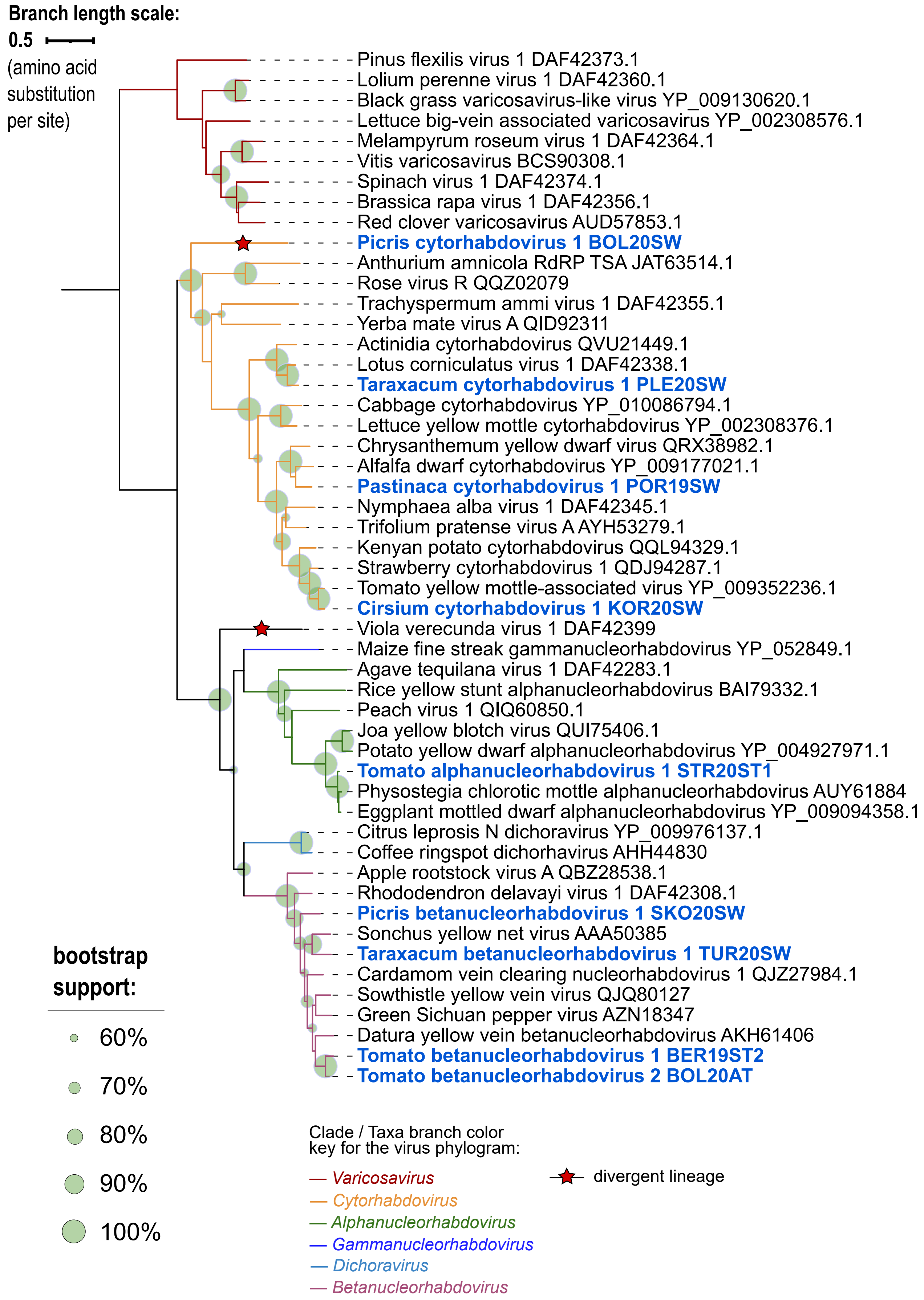


**Supplementary Figure 4-03.** Maximum likelihood phylogenetic tree based on the alignment of conserved RNA-dependent RNA polymerase amino acid sequence of virus species under family *Rhabdoviridae* (*Mononegavirales*).

**Supplementary Figure 4.** Phylogenetic trees up to the virus family (or order) level showing relationships of identified known (in bold black font) and novel (in bold blue font) virus species from this study, with the isolate names indicated. GenBank accession numbers for the viruses from this study can be found in Supplementary Table 5, while GenBank accession number (protein) are indicated for known viruses in the tree. All details of phylogenetic analyses can be found above in Supplementary Table 10.

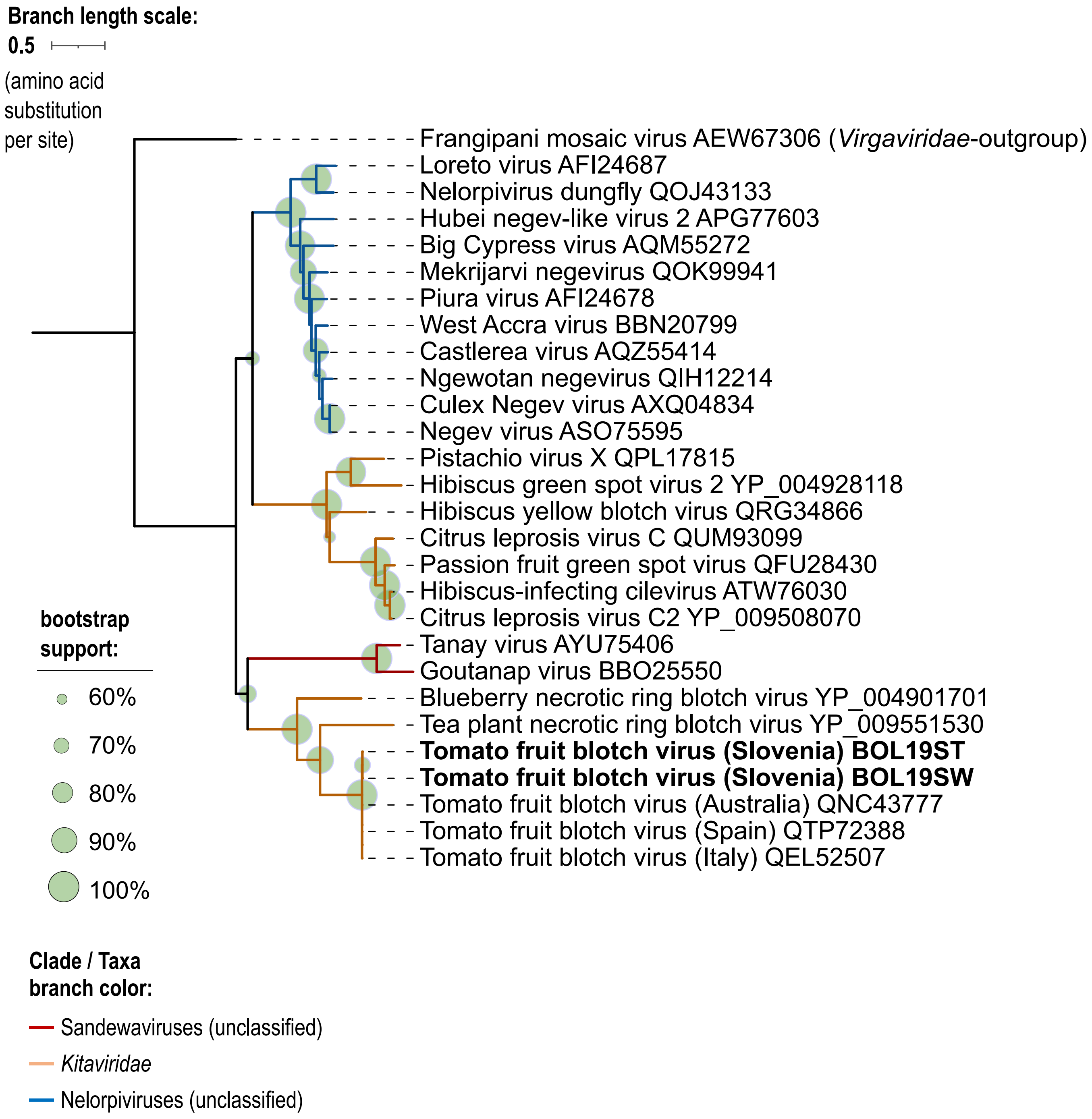


**Supplementary Figure 4-04.** Maximum likelihood phylogenetic tree based on the alignment of conserved RNA-dependent RNA polymerase amino acid sequence of virus species under family *Kitaviridae* (*Martellivirales*).

**Supplementary Figure 4.** Phylogenetic trees up to the virus family (or order) level showing relationships of identified known (in bold black font) and novel (in bold blue font) virus species from this study, with the isolate names indicated. GenBank accession numbers for the viruses from this study can be found in Supplementary Table 5, while GenBank accession number (protein) are indicated for known viruses in the tree. All details of phylogenetic analyses can be found above in Supplementary Table 10.

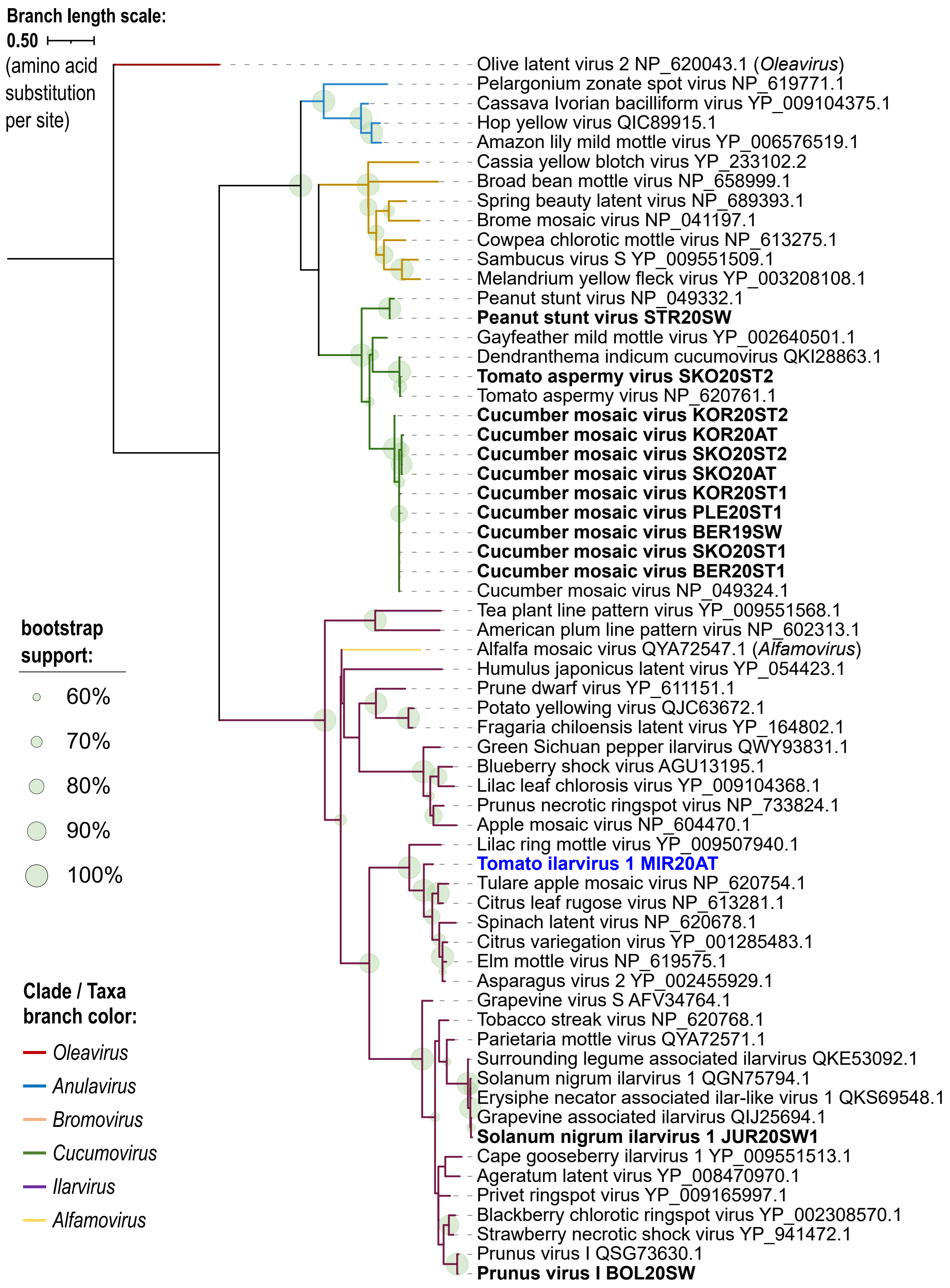


**Supplementary Figure 4-05.** Maximum likelihood phylogenetic tree based on the alignment of conserved RNA-dependent RNA polymerase amino acid sequence of virus species under family *Bromoviridae* (*Martellivirales*).

**Supplementary Figure 4.** Phylogenetic trees up to the virus family (or order) level showing relationships of identified known (in bold black font) and novel (in bold blue font) virus species from this study, with the isolate names indicated. GenBank accession numbers for the viruses from this study can be found in Supplementary Table 5, while GenBank accession number (protein) are indicated for known viruses in the tree. All details of phylogenetic analyses can be found above in Supplementary Table 10.

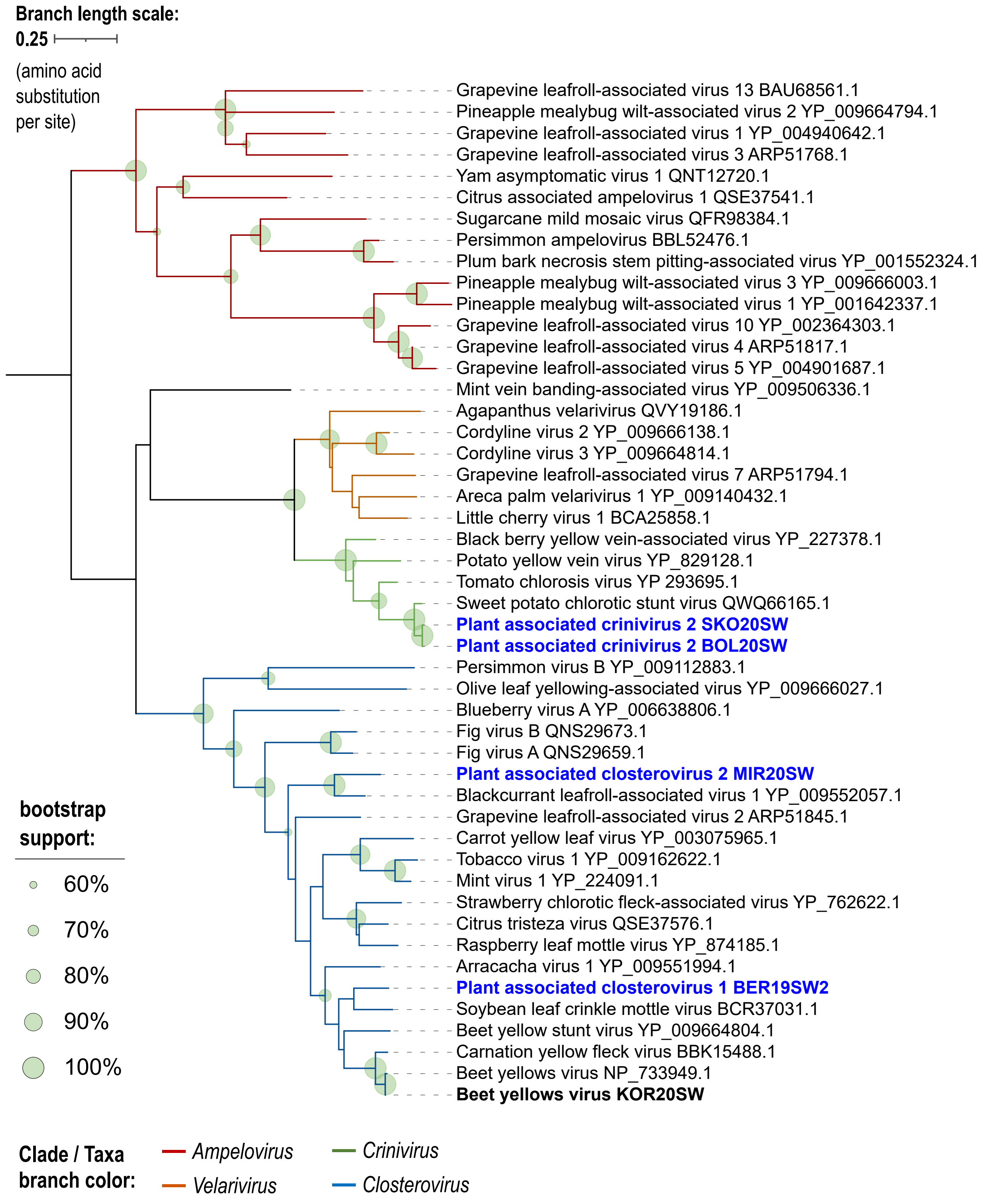


**Supplementary Figure 4-06.** Maximum likelihood phylogenetic tree based on the alignment of conserved RNA-dependent RNA polymerase amino acid sequence of virus species under family *Closteroviridae* (*Martellivirales*).

**Supplementary Figure 4.** Phylogenetic trees up to the virus family (or order) level showing relationships of identified known (in bold black font) and novel (in bold blue font) virus species from this study, with the isolate names indicated. GenBank accession numbers for the viruses from this study can be found in Supplementary Table 5, while GenBank accession number (protein) are indicated for known viruses in the tree. All details of phylogenetic analyses can be found above in Supplementary Table 10.

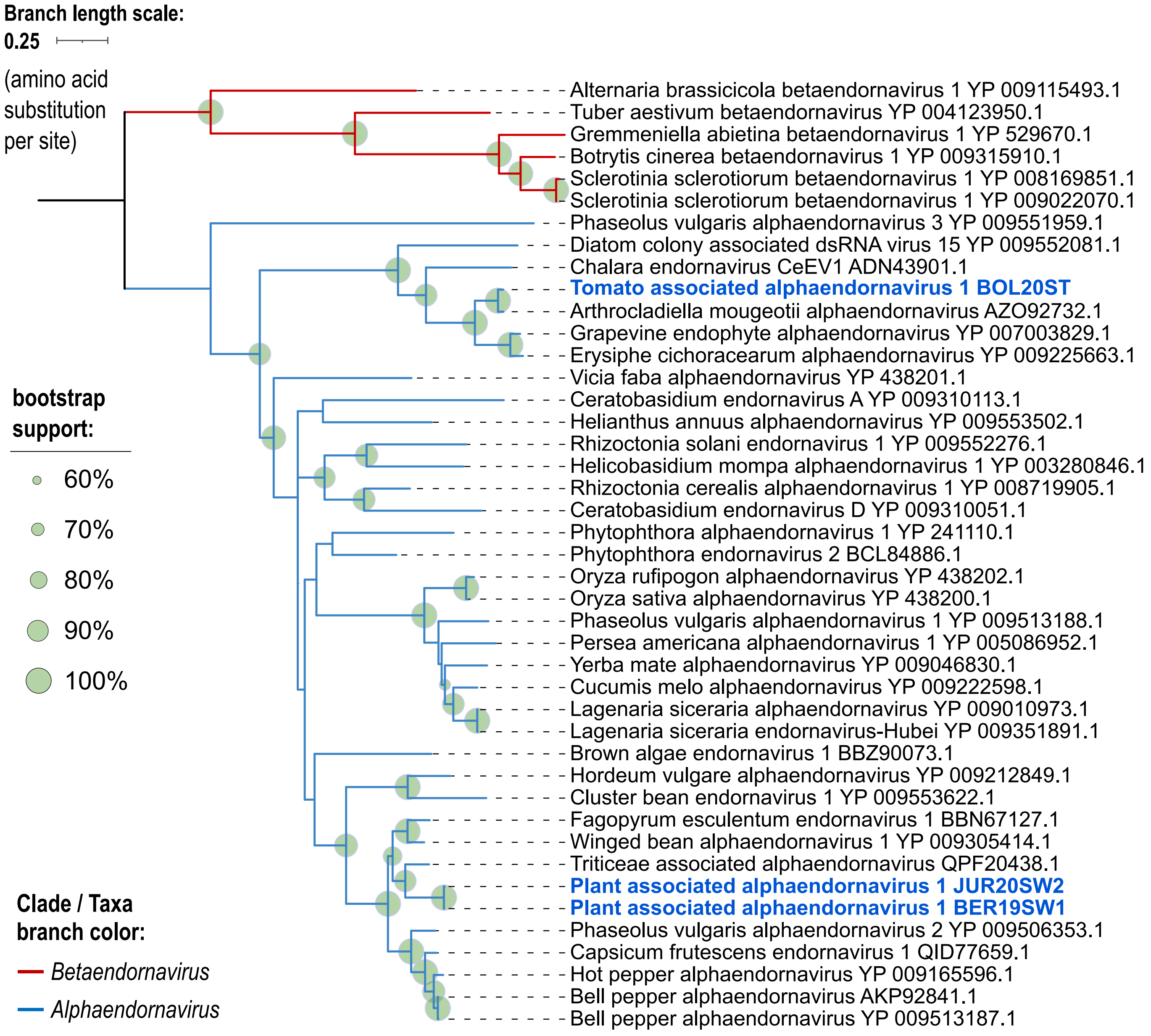


**Supplementary Figure 4-07.** Maximum likelihood phylogenetic tree based on the alignment of conserved RNA-dependent RNA polymerase amino acid sequence of virus species under family *Endornaviridae* (*Martellivirales*).

**Supplementary Figure 4.** Phylogenetic trees up to the virus family (or order) level showing relationships of identified known (in bold black font) and novel (in bold blue font) virus species from this study, with the isolate names indicated. GenBank accession numbers for the viruses from this study can be found in Supplementary Table 5, while GenBank accession number (protein) are indicated for known viruses in the tree. All details of phylogenetic analyses can be found above in Supplementary Table 10.

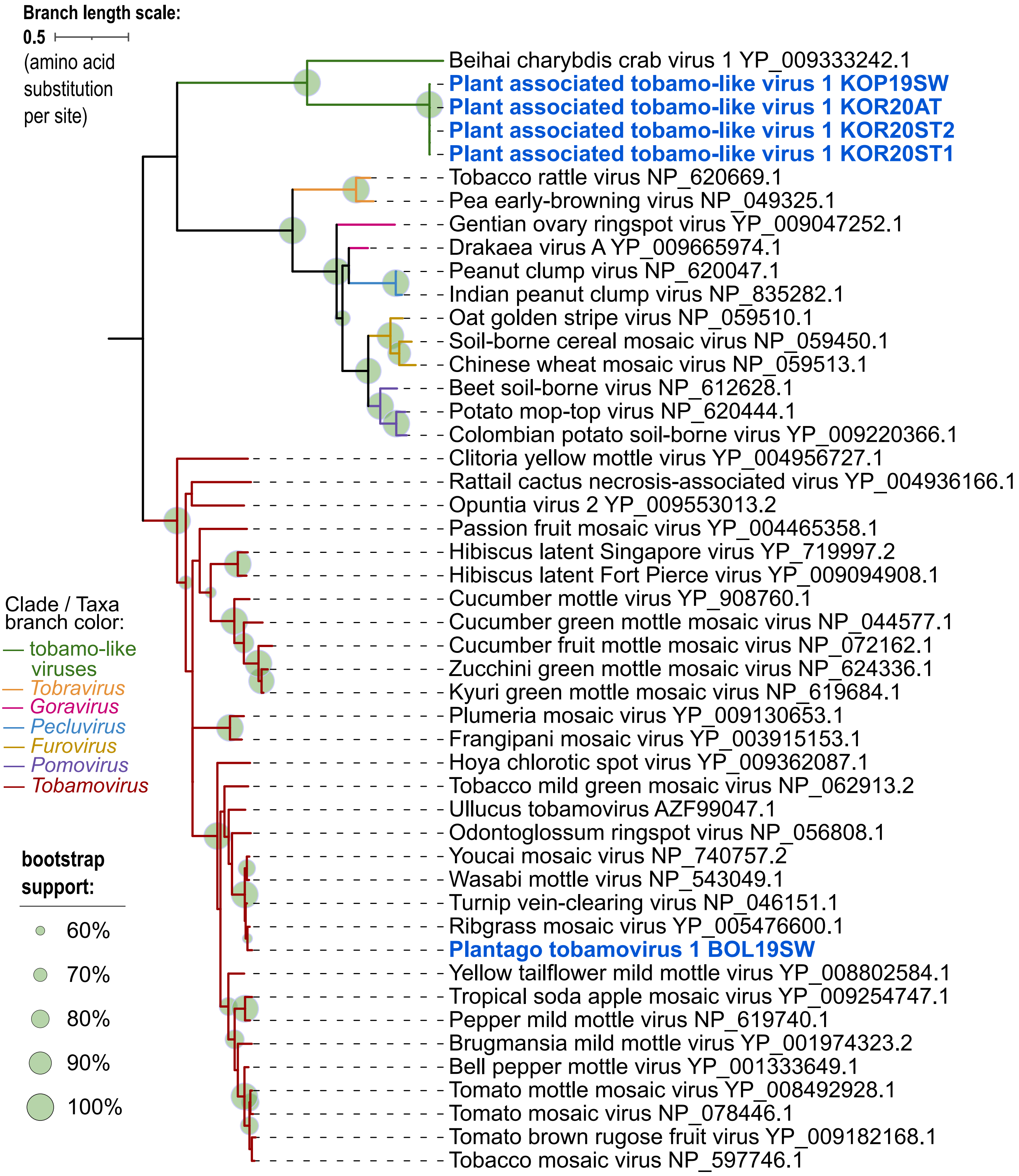


**Supplementary Figure 4-08-A.** Maximum likelihood phylogenetic tree based on the alignment of conserved methyl transferase-helicase amino acid sequence of virus species under family *Virgaviridae* (*Martellivirales*).

**Supplementary Figure 4.** Phylogenetic trees up to the virus family (or order) level showing relationships of identified known (in bold black font) and novel (in bold blue font) virus species from this study, with the isolate names indicated. GenBank accession numbers for the viruses from this study can be found in Supplementary Table 5, while GenBank accession number (protein) are indicated for known viruses in the tree. All details of phylogenetic analyses can be found above in Supplementary Table 10.

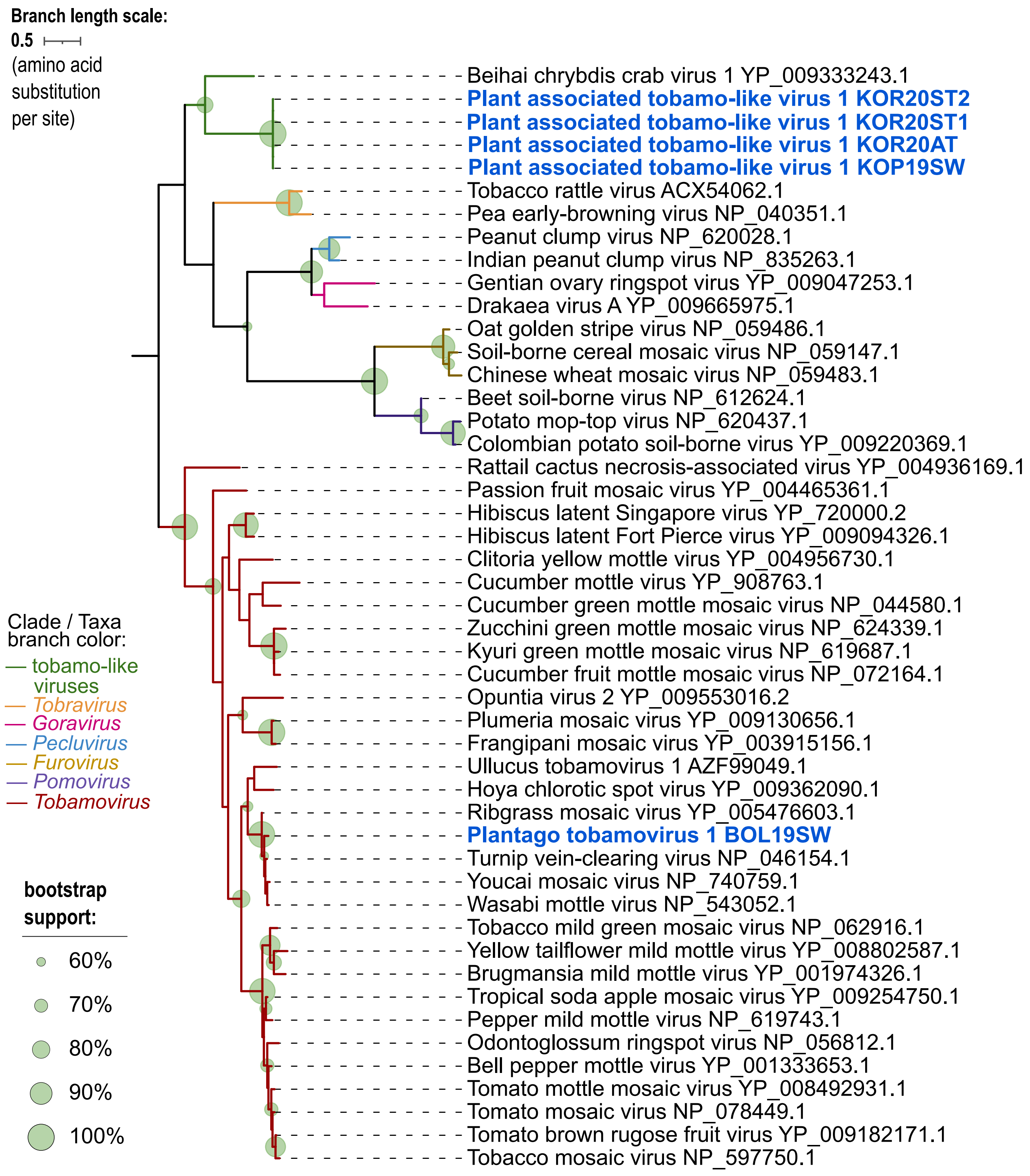


**Supplementary Figure 4-08-A.** Maximum likelihood phylogenetic tree based on the alignment of conserved coat protein amino acid sequence of virus species under family *Virgaviridae* (*Martellivirales*).

**Supplementary Figure 4.** Phylogenetic trees up to the virus family (or order) level showing relationships of identified known (in bold black font) and novel (in bold blue font) virus species from this study, with the isolate names indicated. GenBank accession numbers for the viruses from this study can be found in Supplementary Table 5, while GenBank accession number (protein) are indicated for known viruses in the tree. All details of phylogenetic analyses can be found above in Supplementary Table 10.

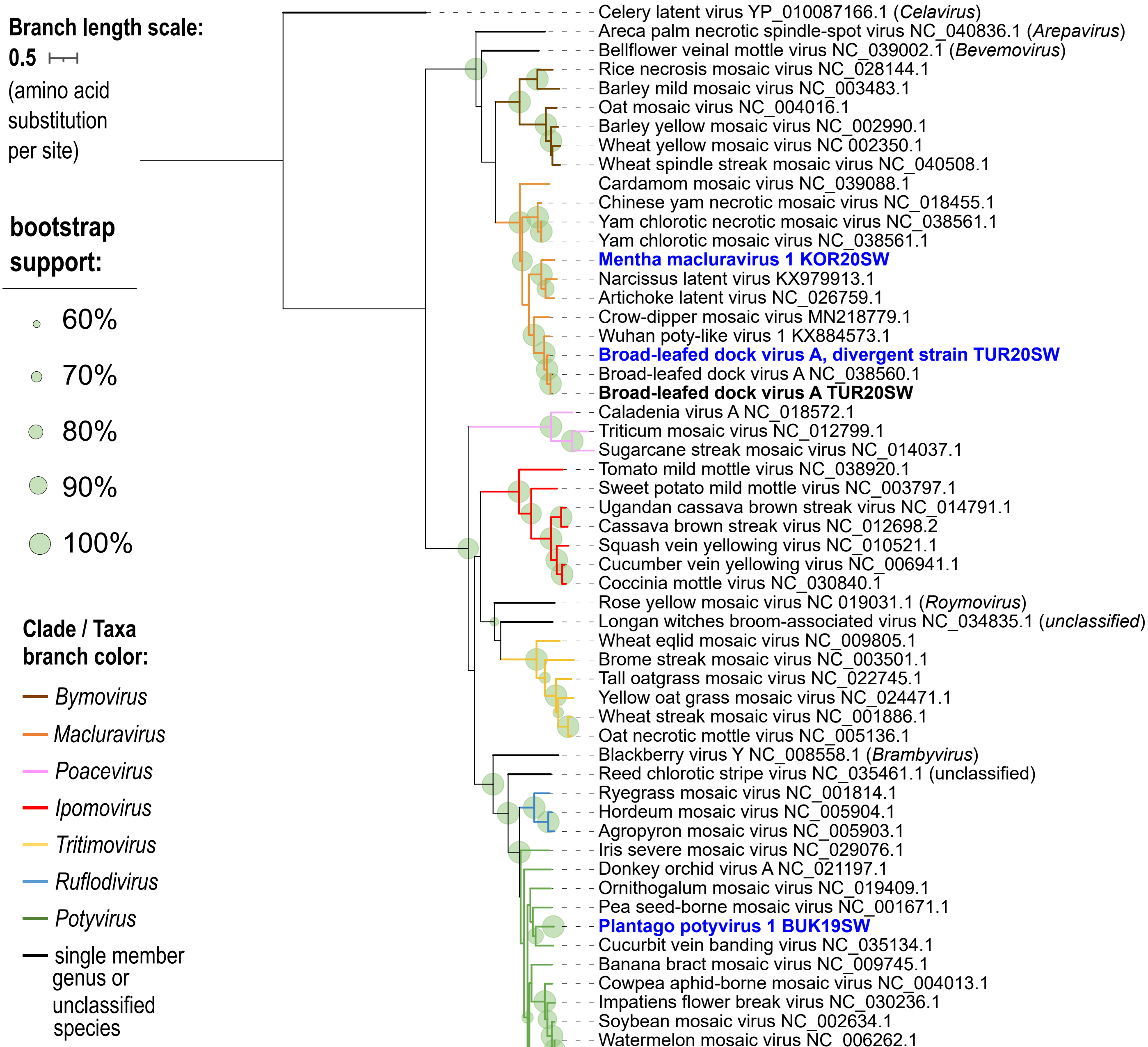

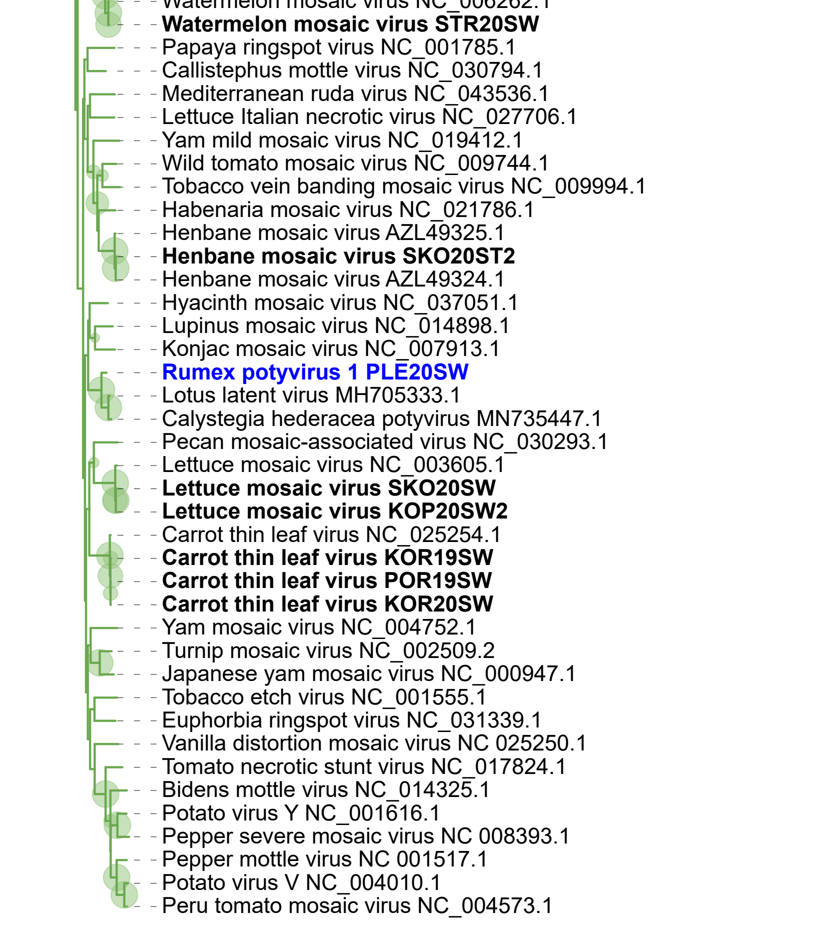


**Supplementary Figure 4-09.** Maximum likelihood phylogenetic tree based aon the alignment of conserved RNA-dependent RNA polymerase amino acid sequence of virus species under family *Potyviridae* (*Patatavirales*).

**Supplementary Figure 4.** Phylogenetic trees up to the virus family (or order) level showing relationships of identified known (in bold black font) and novel (in bold blue font) virus species from this study, with the isolate names indicated. GenBank accession numbers for the viruses from this study can be found in Supplementary Table 5, while GenBank accession number (protein) are indicated for known viruses in the tree. All details of phylogenetic analyses can be found above in Supplementary Table 10.

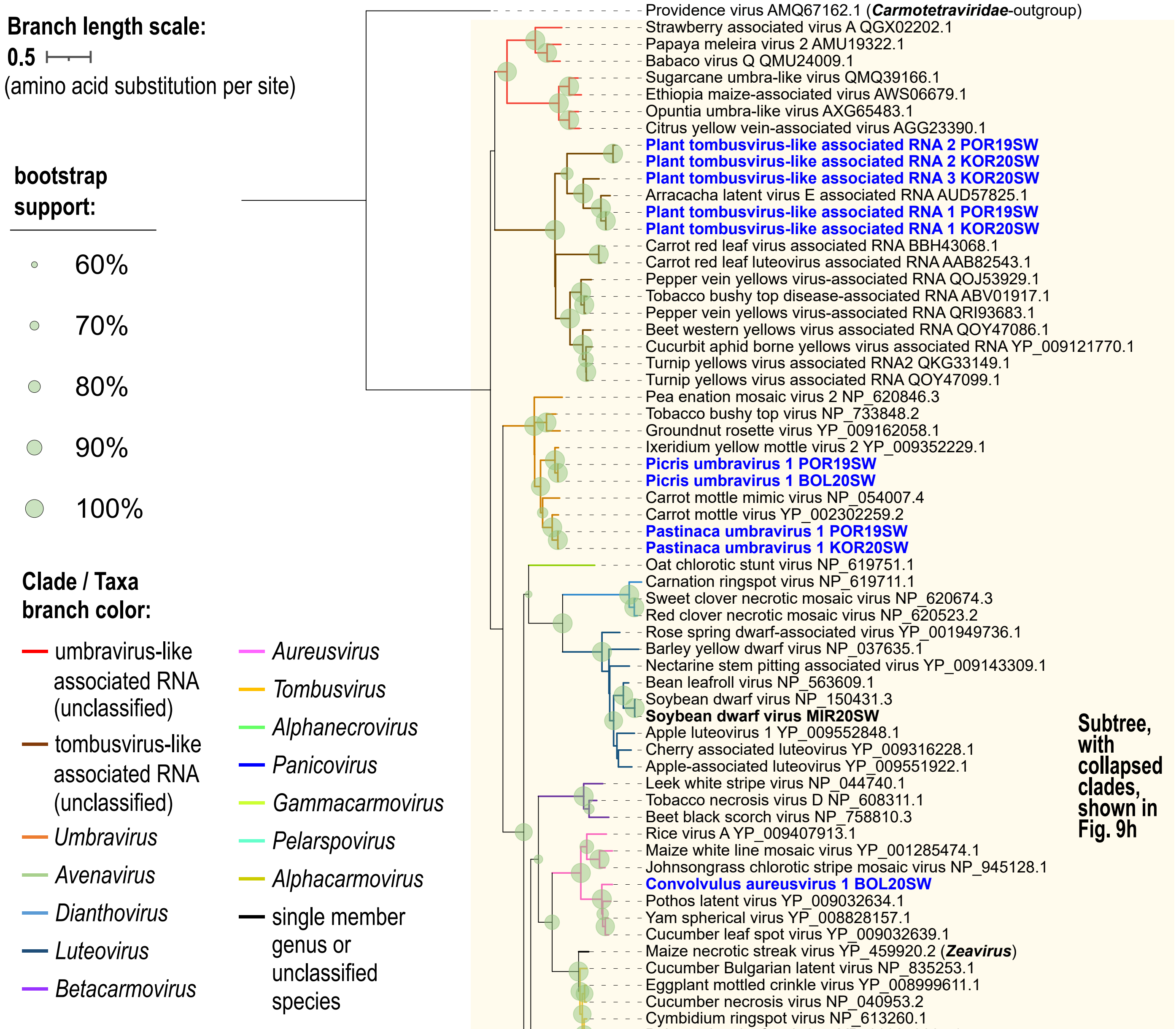


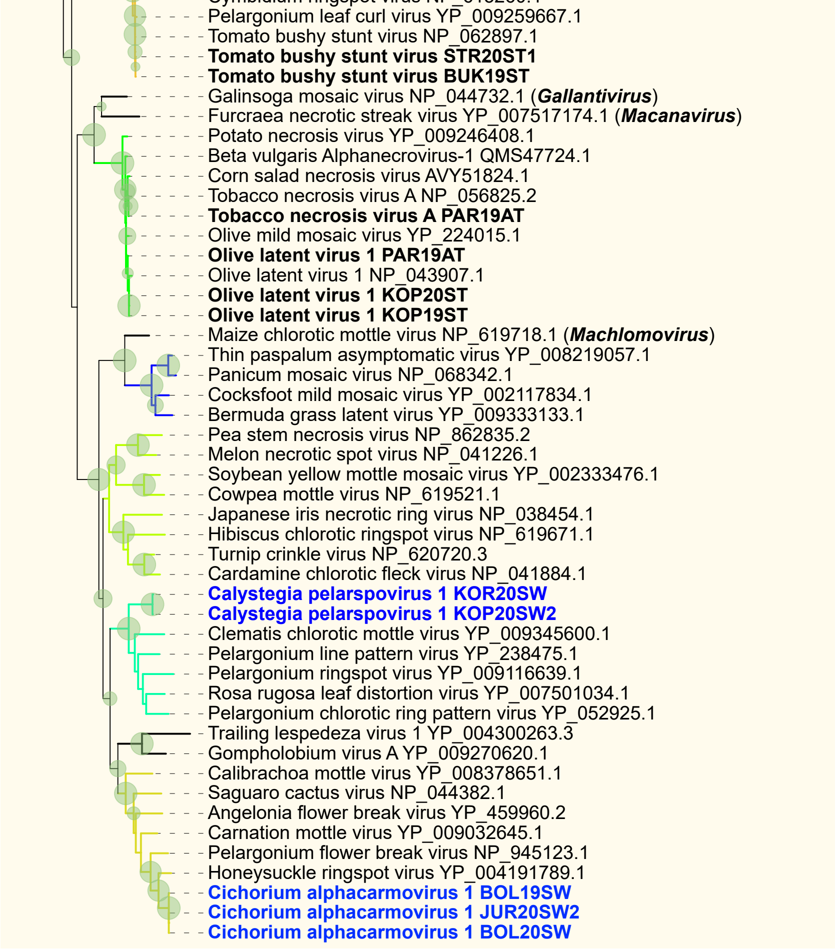


**Supplementary Figure 4-10.** Maximum likelihood phylogenetic tree based aon the alignment of conserved RNA-dependent RNA polymerase amino acid sequence of virus species under family *Tombusviridae* (*Tolivirales*).

**Supplementary Figure 4.** Phylogenetic trees up to the virus family (or order) level showing relationships of identified known (in bold black font) and novel (in bold blue font) virus species from this study, with the isolate names indicated. GenBank accession numbers for the viruses from this study can be found in Supplementary Table 5, while GenBank accession number (protein) are indicated for known viruses in the tree. All details of phylogenetic analyses can be found above in Supplementary Table 10.

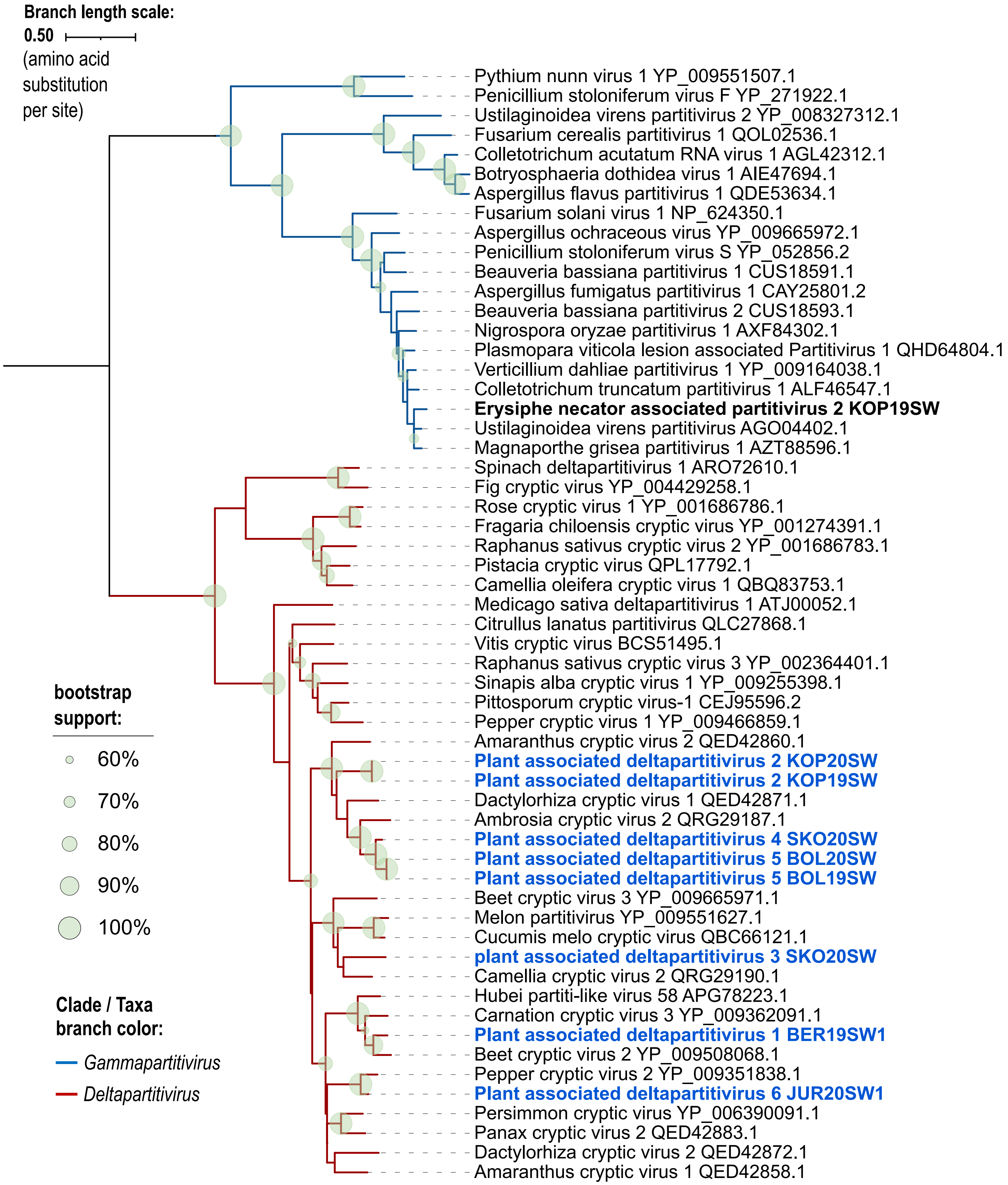


**Supplementary Figure 4-11.** Maximum likelihood phylogenetic tree based on the alignment of conserved RNA-dependent RNA polymerase amino acid sequence of virus species under family *Partitiviridae* (*Durnavirales*).

**Supplementary Figure 4.** Phylogenetic trees up to the virus family (or order) level showing relationships of identified known (in bold black font) and novel (in bold blue font) virus species from this study, with the isolate names indicated. GenBank accession numbers for the viruses from this study can be found in Supplementary Table 5, while GenBank accession number (protein) are indicated for known viruses in the tree. All details of phylogenetic analyses can be found above in Supplementary Table 10.

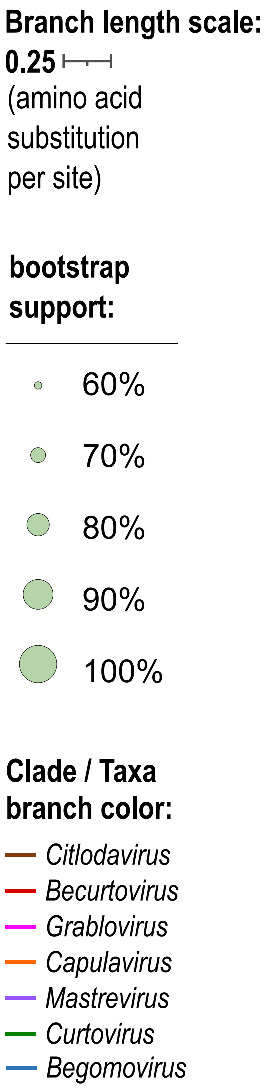


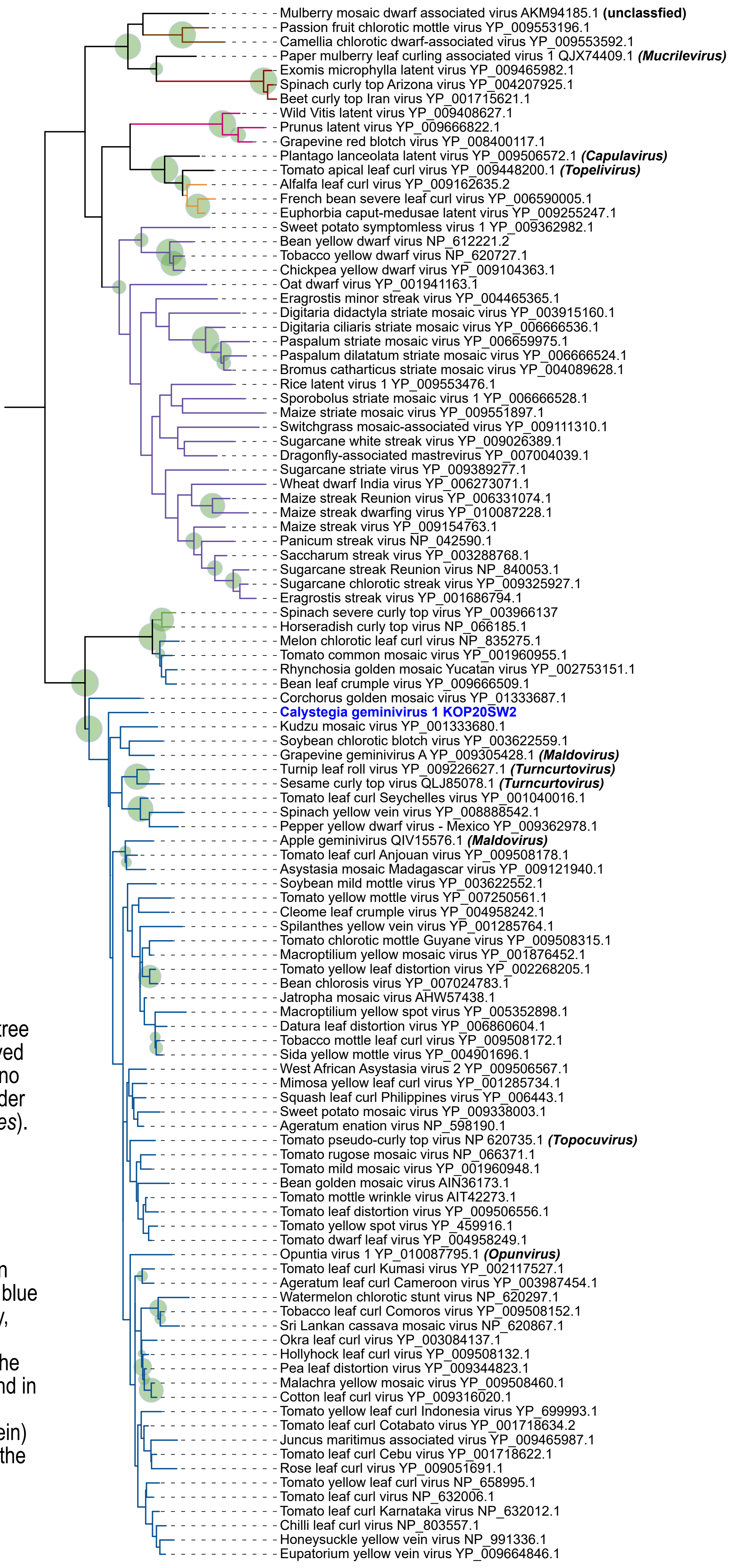


**Supplementary Figure 4-12.** Maximum likelihood phylogenetic tree based on the alignment of conserved replicase (C1 gene in DNA-A) amino acid sequence of virus species under family *Geminiviridae* (*Geplafuvirales*).

**Supplementary Figure 4.** Phylogenetic trees up to the virus family (or order) level showing relationships of identified known (in bold black font) and novel (in bold blue font) virus species from this study, with the isolate names indicated. GenBank accession numbers for the viruses from this study can be found in Supplementary Table 5, while GenBank accession number (protein) are indicated for known viruses in the tree. All details of phylogenetic analyses can be found above in Supplementary Table 10.

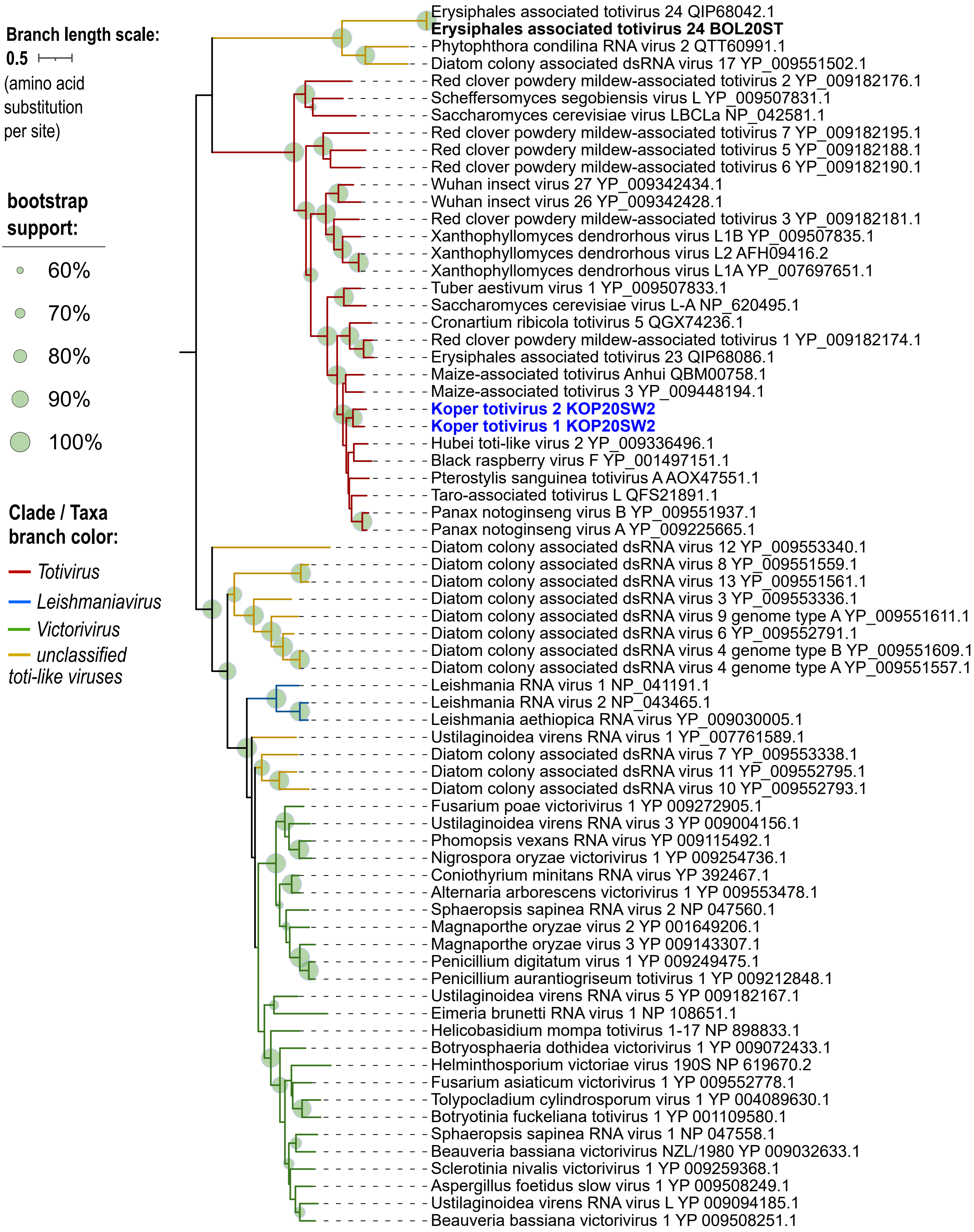


**Supplementary Figure 4-13.** Maximum likelihood phylogenetic tree based on the alignment of conserved RNA-dependent RNA polymerase amino acid sequence of virus species under family *Totiviridae* (*Ghabrivirales*).

**Supplementary Figure 4.** Phylogenetic trees up to the virus family (or order) level showing relationships of identified known (in bold black font) and novel (in bold blue font) virus species from this study, with the isolate names indicated. GenBank accession numbers for the viruses from this study can be found in Supplementary Table 5, while GenBank accession number (protein) are indicated for known viruses in the tree. All details of phylogenetic analyses can be found above in Supplementary Table 10.

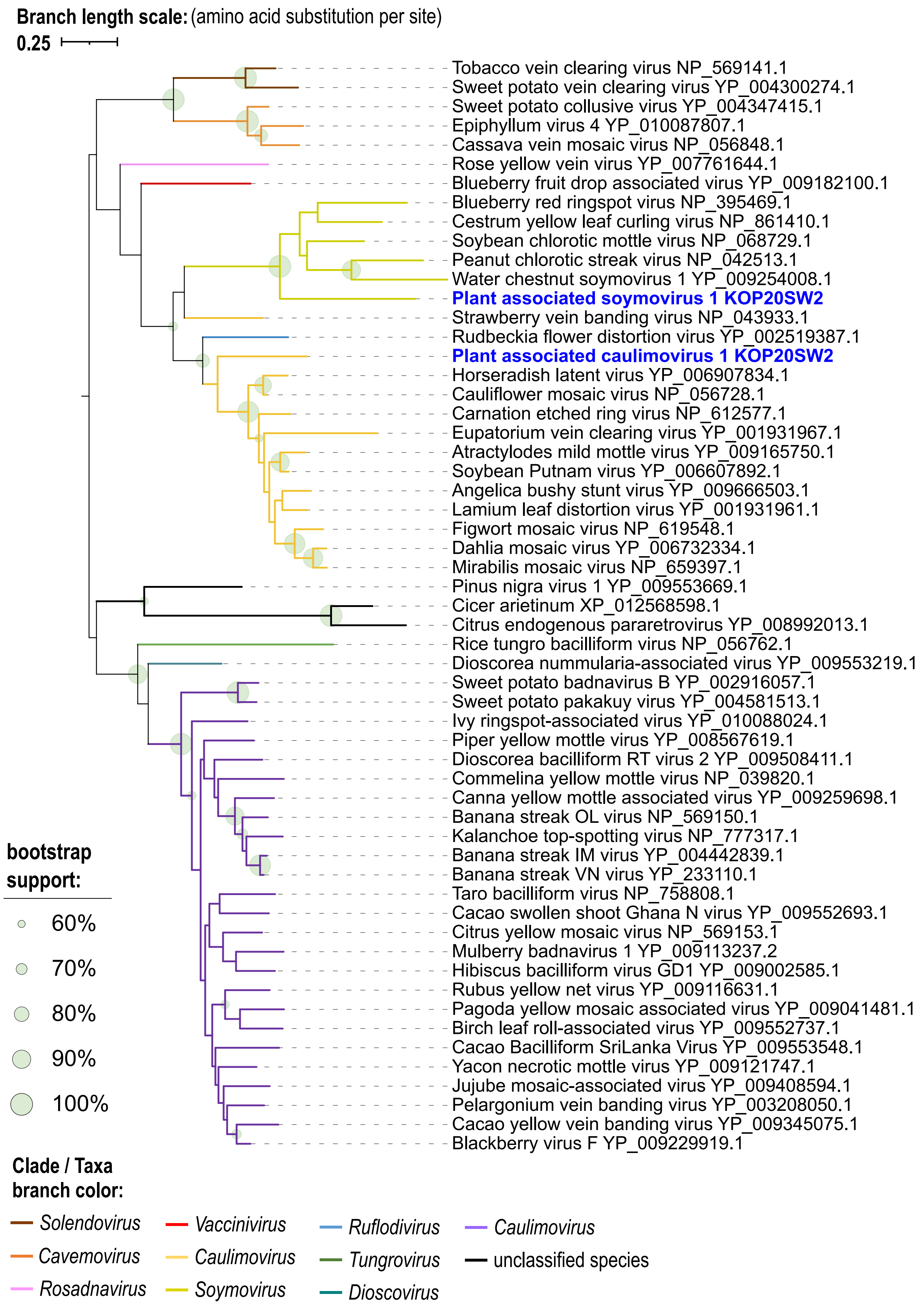


**Supplementary Figure 4-14.** Maximum likelihood phylogenetic tree based on the alignment of conserved reverse transcriptase (RT) amino acid sequence of virus species under family *Caulimoviridae* (*Ortelivirales*).

**Supplementary Figure 4.** Phylogenetic trees up to the virus family (or order) level showing relationships of identified known (in bold black font) and novel (in bold blue font) virus species from this study, with the isolate names indicated. GenBank accession numbers for the viruses from this study can be found in Supplementary Table 5, while GenBank accession number (protein) are indicated for known viruses in the tree. All details of phylogenetic analyses can be found above in Supplementary Table 10.

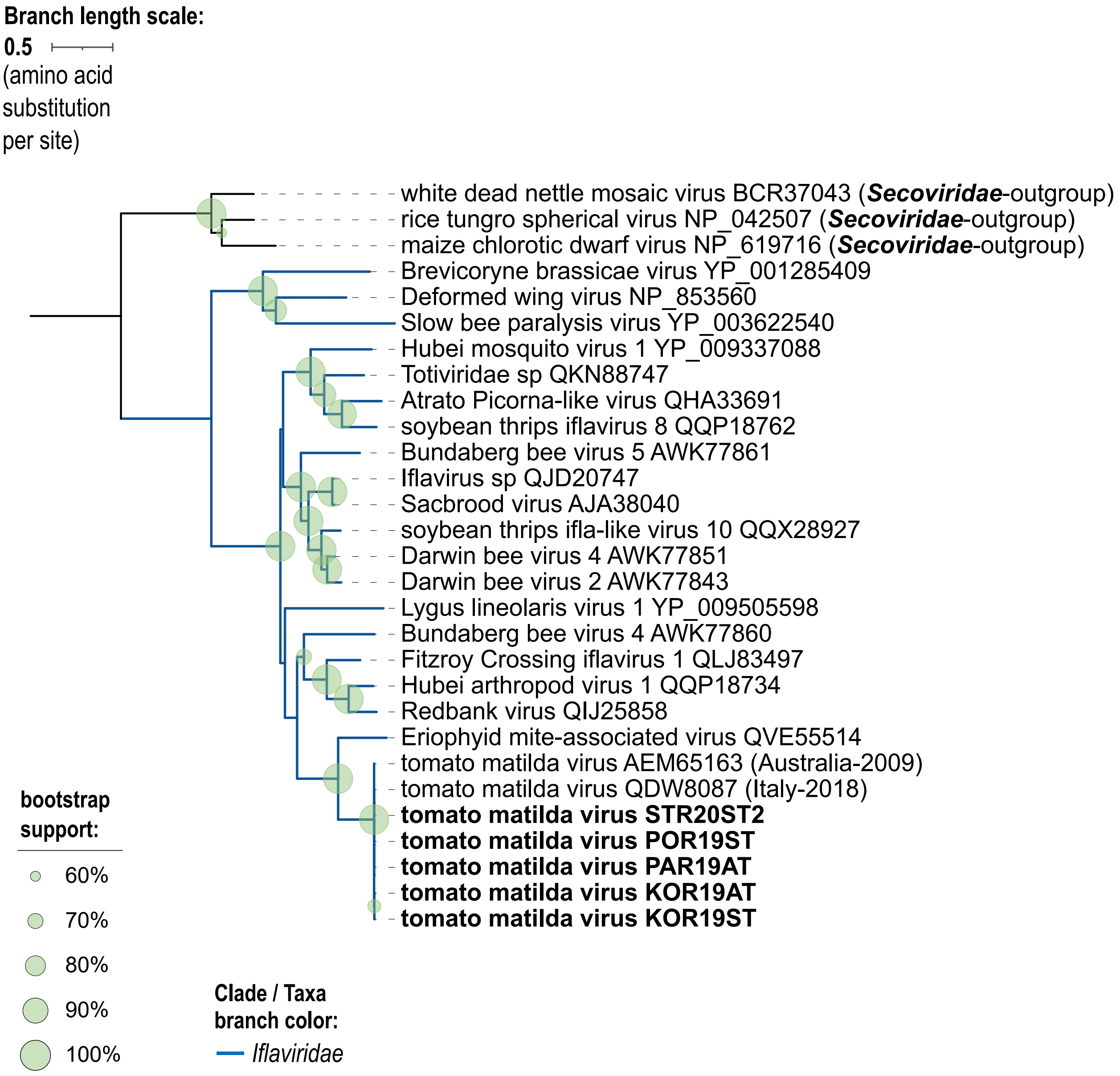


**Supplementary Figure 4-15.** Maximum likelihood phylogenetic tree based on the alignment of conserved RNA-dependent RNA polymerase amino acid sequence of virus species under *Iflaviridae* (*Picornavirales*), highlighting isolates of Tomato matilda virus from this study.

**Supplementary Figure 4.** Phylogenetic trees up to the virus family (or order) level showing relationships of identified known (in bold black font) and novel (in bold blue font) virus species from this study, with the isolate names indicated. GenBank accession numbers for the viruses from this study can be found in Supplementary Table 5, while GenBank accession number (protein) are indicated for known viruses in the tree. All details of phylogenetic analyses can be found above in Supplementary Table 10.

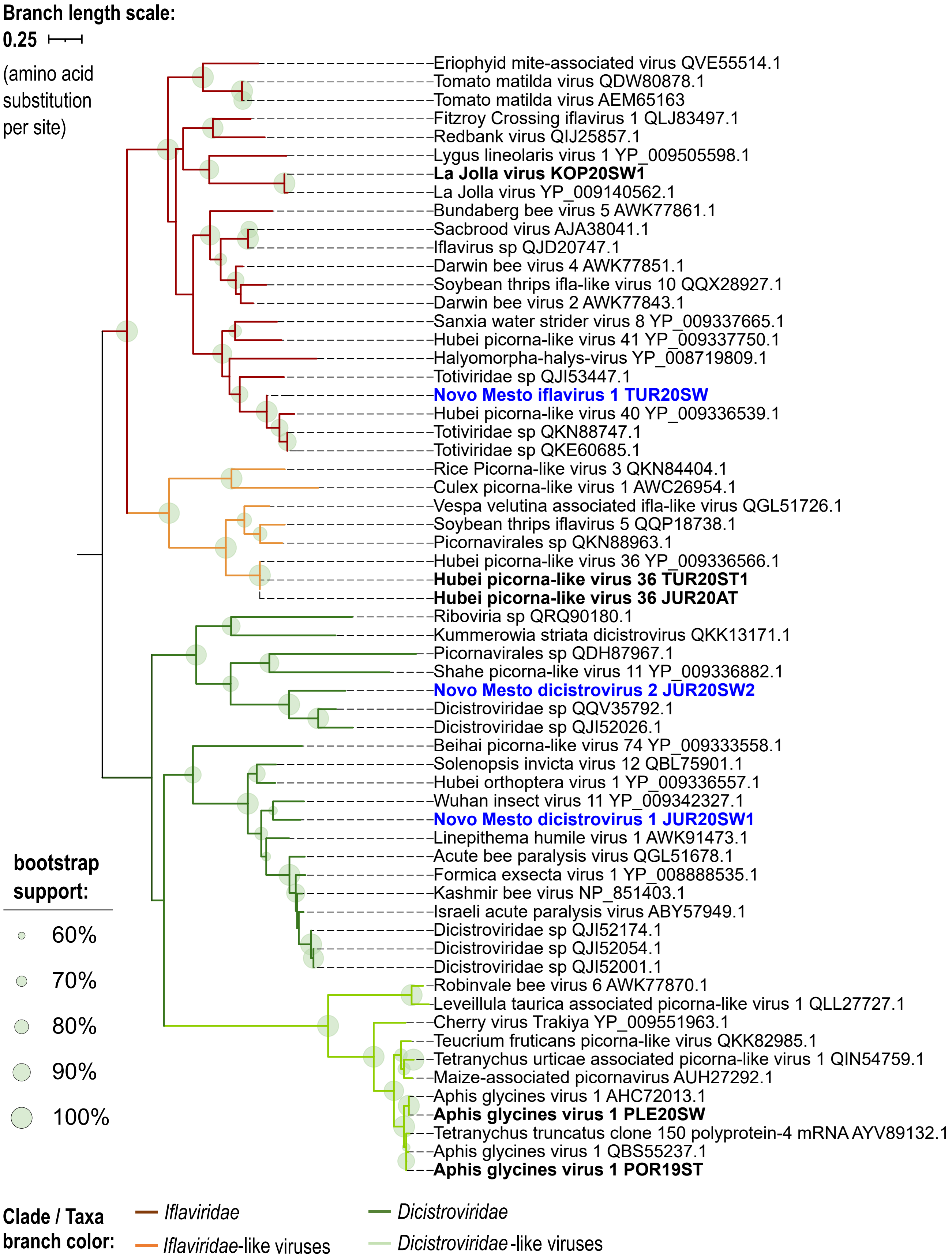


**Supplementary Figure 4-16.** Maximum likelihood phylogenetic tree based on the alignment of conserved RNA-dependent RNA polymerase amino acid sequence of virus and virus-like species classifiable under family *Dicistroviridae* and *Iflaviridae* (*Picornavirales*).

**Supplementary Figure 4.** Phylogenetic trees up to the virus family (or order) level showing relationships of identified known (in bold black font) and novel (in bold blue font) virus species from this study, with the isolate names indicated. GenBank accession numbers for the viruses from this study can be found in Supplementary Table 5, while GenBank accession number (protein) are indicated for known viruses in the tree. All details of phylogenetic analyses can be found above in Supplementary Table 10.

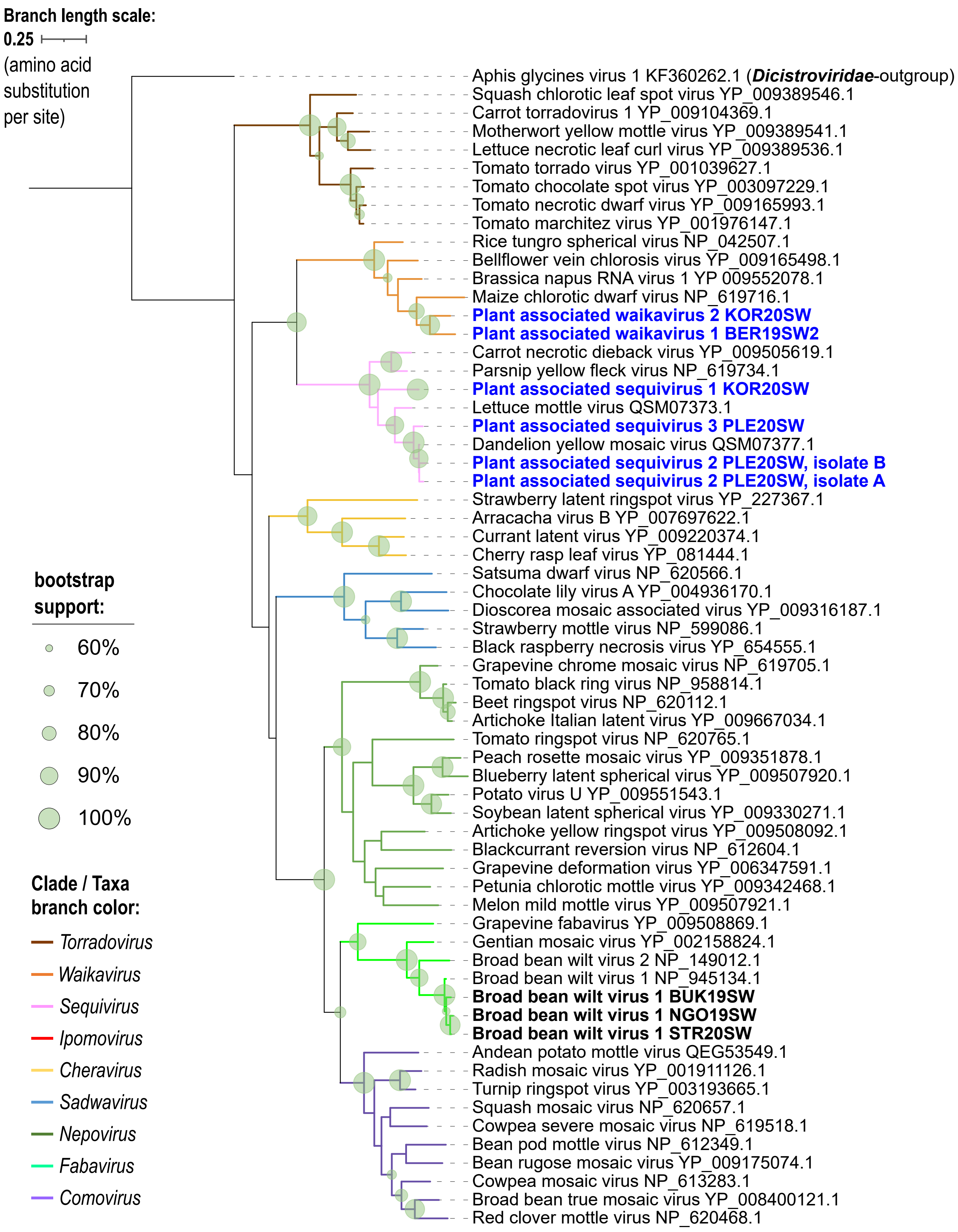


**Supplementary Figure 4-17.** Maximum likelihood phylogenetic tree based on the alignment of conserved RNA-dependent RNA polymerase amino acid sequence of virus species under family *Secoviridae* (*Picornavirales*).

**Supplementary Figure 4.** Phylogenetic trees up to the virus family (or order) level showing relationships of identified known (in bold black font) and novel (in bold blue font) virus species from this study, with the isolate names indicated. GenBank accession numbers for the viruses from this study can be found in Supplementary Table 5, while GenBank accession number (protein) are indicated for known viruses in the tree. All details of phylogenetic analyses can be found above in Supplementary Table 10.

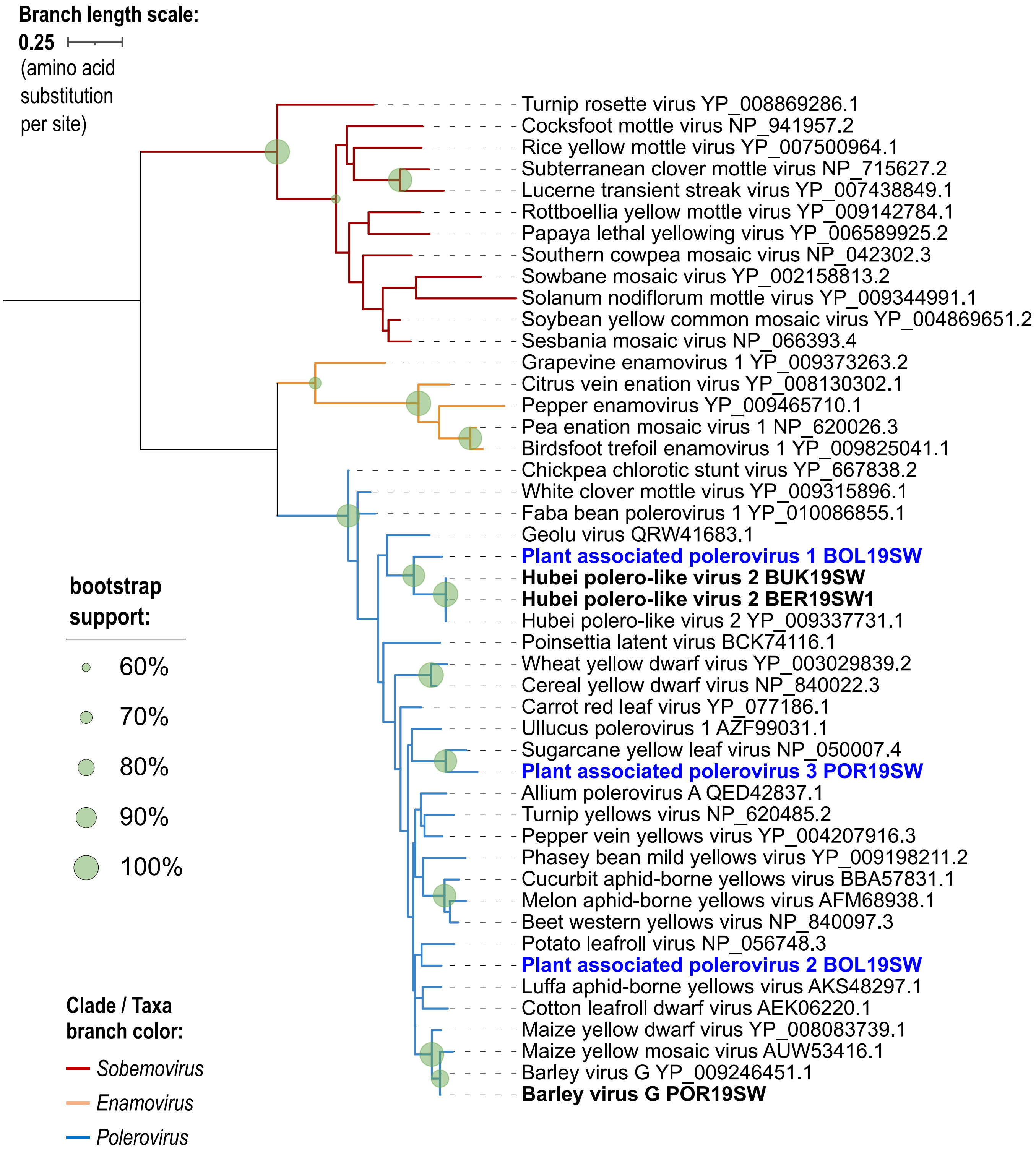


**Supplementary Figure 4-18.** Maximum likelihood phylogenetic tree based on the alignment of conserved RNA-dependent RNA polymerase amino acid sequence of virus species under family *Solemoviridae* (*Sobelivirales*).

**Supplementary Figure 4.** Phylogenetic trees up to the virus family (or order) level showing relationships of identified known (in bold black font) and novel (in bold blue font) virus species from this study, with the isolate names indicated. GenBank accession numbers for the viruses from this study can be found in Supplementary Table 5, while GenBank accession number (protein) are indicated for known viruses in the tree. All details of phylogenetic analyses can be found above in Supplementary Table 10.

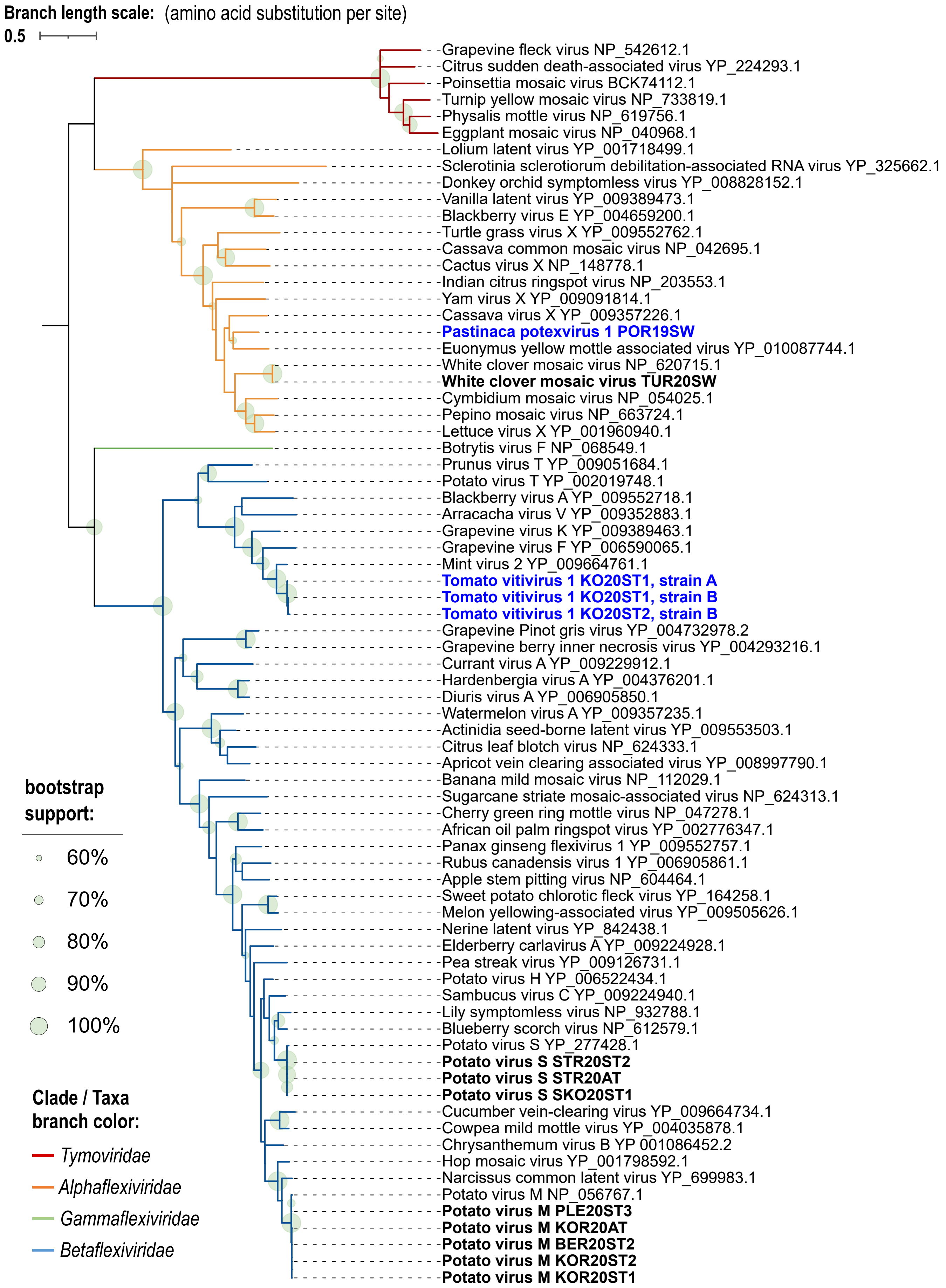


**Supplementary Figure 4-19.** Maximum likelihood phylogenetic tree based on the alignment of conserved RNA-dependent RNA polymerase amino acid sequence of virus species under order *Tymovirales*.

**Supplementary Figure 4.** Phylogenetic trees up to the virus family (or order) level showing relationships of identified known (in bold black font) and novel (in bold blue font) virus species from this study, with the isolate names indicated. GenBank accession numbers for the viruses from this study can be found in Supplementary Table 5, while GenBank accession number (protein) are indicated for known viruses in the tree. All details of phylogenetic analyses can be found above in Supplementary Table 10.

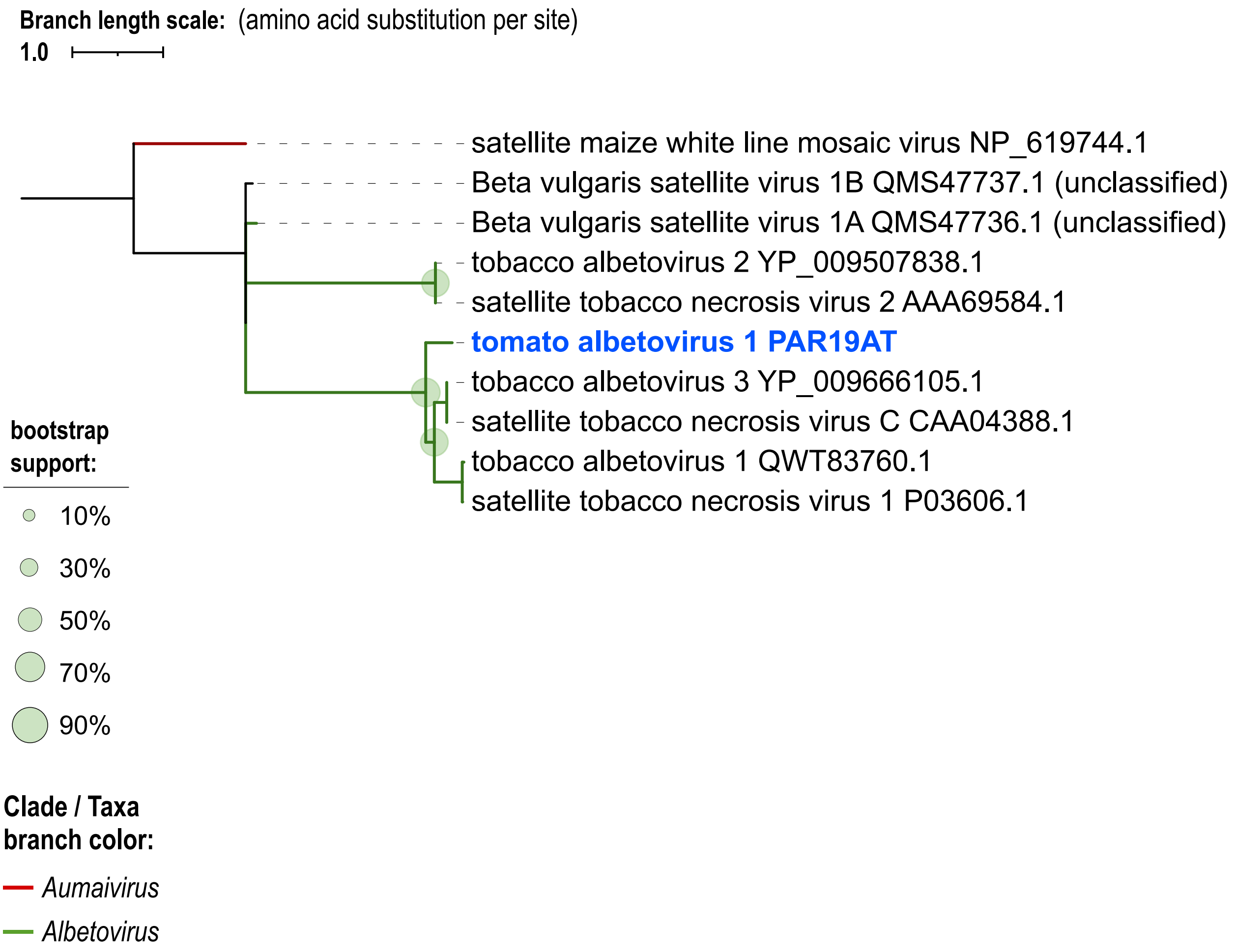


**Supplementary Figure 4-20.** Maximum likelihood phylogenetic tree based on the alignment of conserved coat protein amino acid sequence of satellite virus species with emphasis on genus *Albetovirus*.

**Supplementary Figure 4.** Phylogenetic trees up to the virus family (or order) level showing relationships of identified known (in bold black font) and novel (in bold blue font) virus species from this study, with the isolate names indicated. GenBank accession numbers for the viruses from this study can be found in Supplementary Table 5, while GenBank accession number (protein) are indicated for known viruses in the tree. All details of phylogenetic analyses can be found above in Supplementary Table 10.
