## Supplementary material for "In-depth study of tomato and weed viromes reveals undiscovered plant virus diversity in an agroecosystem": Additonal-File-10_Supp-Figure-5_RT-PCR-tests-HTS-PTV1

### SUPPLEMENTARY INFORMATION

### Additional File 10

**Supplementary Figure 5.** RT-PCR, TEM and nanopore sequencing results on the confirmation of infectivity of Plantago tobamovirus 1 (PTV1) on Solanaceae hosts.

**a**

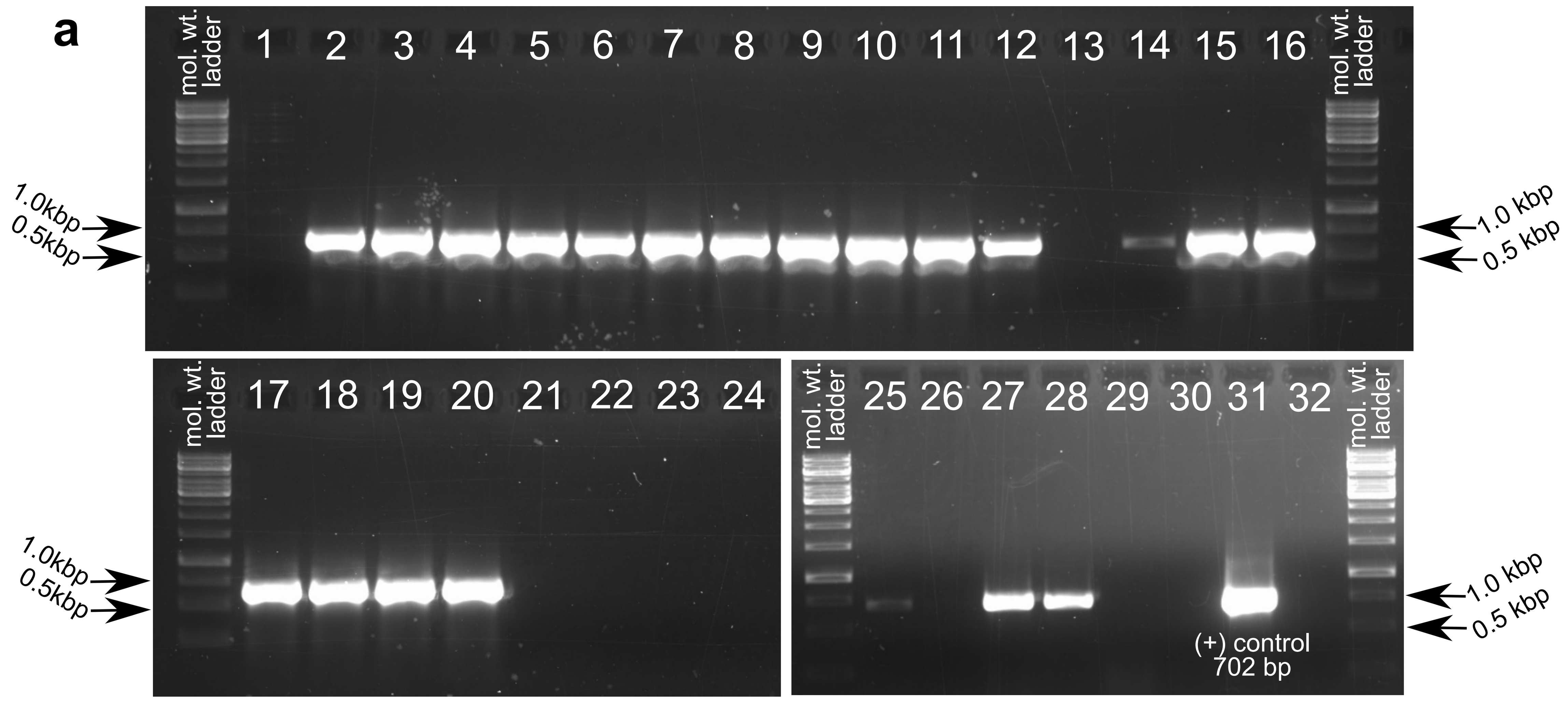


Note: kbp - kilo base pairs

**b**

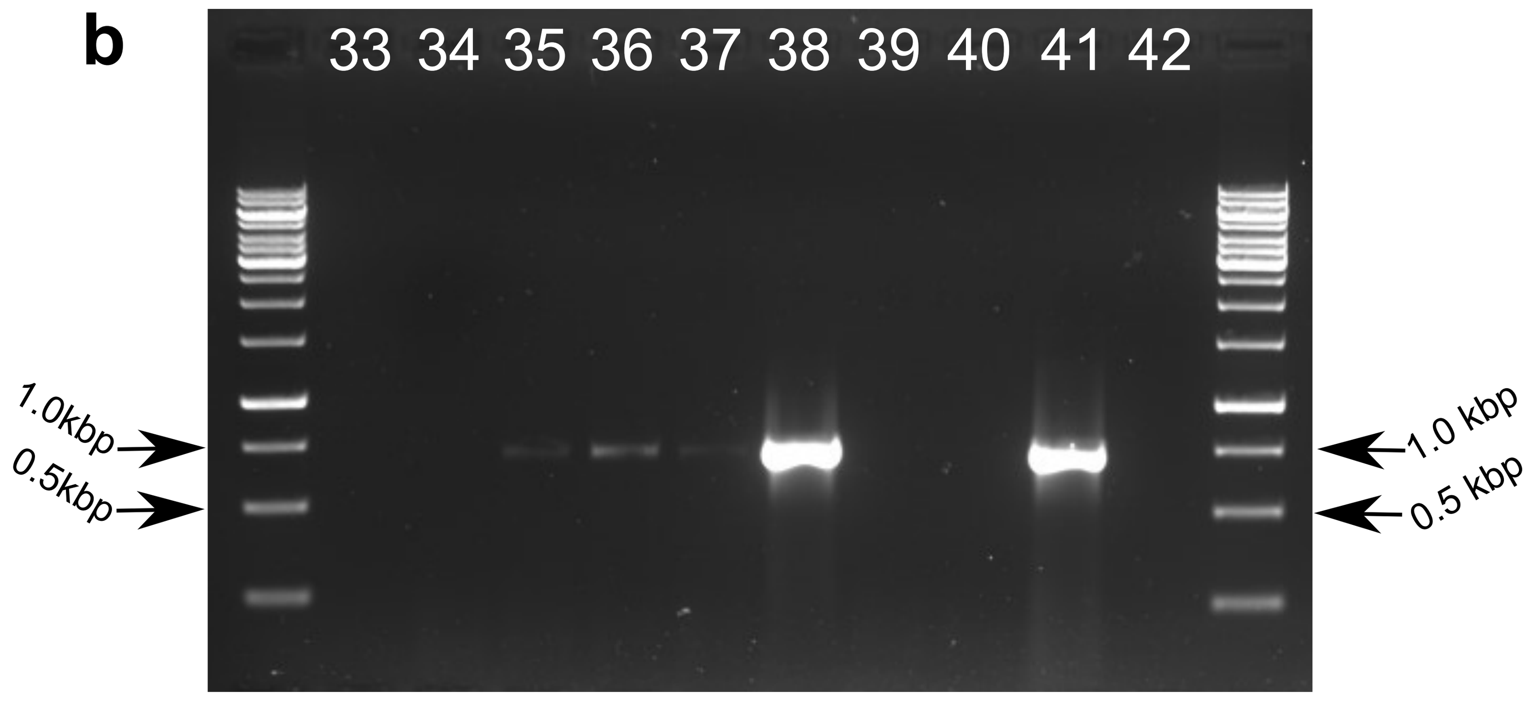


Note: kbp - kilo base pairs

Samples tested in the RT-PCR assay shown in **a:**

Assay controls in **a**:

1 - non-template control 1
13 - blank 1
21 - negative control of RNA isolation 1
22 - negative control of RNA isolation 2
23 - non-template control 1
24 - blank 2
31 - positive control (PTV1 RNA)
32 - negative control of RNA isolation 3

| Inoculated plant | 7dpi pool of inoculated leaves | 14 dpi pool of inoculated leaves | 14 dpi pool of young apical (systemic) leaves | 21 dpi pool of young apical (systemic) leaves | 28 dpi pool of young apical (systemic) leaves | 35 dpi pool of young apical (systemic) leaves |
|---|---|---|---|---|---|---|
| *Solanum lycopersicum* | 2[asym] | 8[asym] | 14[asym] | 25[a,asym]<br>26[b,asym] | 27[a,asym]<br>28[b,asym] | 29[a,asym]<br>30[b,asym] |
| *Nicotiana benthamiana* | 3[sym] | 9[sym] | 15[sym] | 17[sym] | 19[sym] | not tested |
| *Nicotiana clevelandii* | 4[sym] | 10[sym] | 16[sym] | 18[sym] | 20[sym,HTS(+)] | not tested |

Note: dpi - days post inoculation, [a]10x diluted RNA, [b]100x diluted RNA, [asym]asymptomatic, [sym]symptomatic, [HTS(+)]was used in HTS where presence of PTV1 was confirmed. Other inoculated plants tested: 7dpi/14dpi pool of inoculated leaves: *Nicotiana tabacum* (lane 5/lane11), *Nicotiana glutinosa* (lane 6/lane12), *Chenopodium quinoa* (lane 7/-);

Samples tested in the RT-PCR assay shown in **b:**

Assay controls in **b**:

33 - non-template control
41 - positive control (PTV1 RNA)
42 - negative control of RNA isolation

| Inoculated plant | 28 dpi young apical (systemic) leaves from individual plants | | | | | 49 dpi pool of young apical (systemic) leaves | 56 dpi pool of young apical (systemic) leaves |
|---|---|---|---|---|---|---|---|
| | plant 1 | plant 2 | plant 3 | plant 4 | plant 5 | | |
| *Solanum lycopersicum* | 34[asym] | 35[asym] | 36[asym] | 37[asym] | 38[asym] | 39[asym] | 40[asym] |

Note: RNA extracts used in this assay (**b**) were diluted 1:200, except for the positive control.
dpi - days post inoculation. [asym]asymptomatic. [sym]symptomatic.

**c**

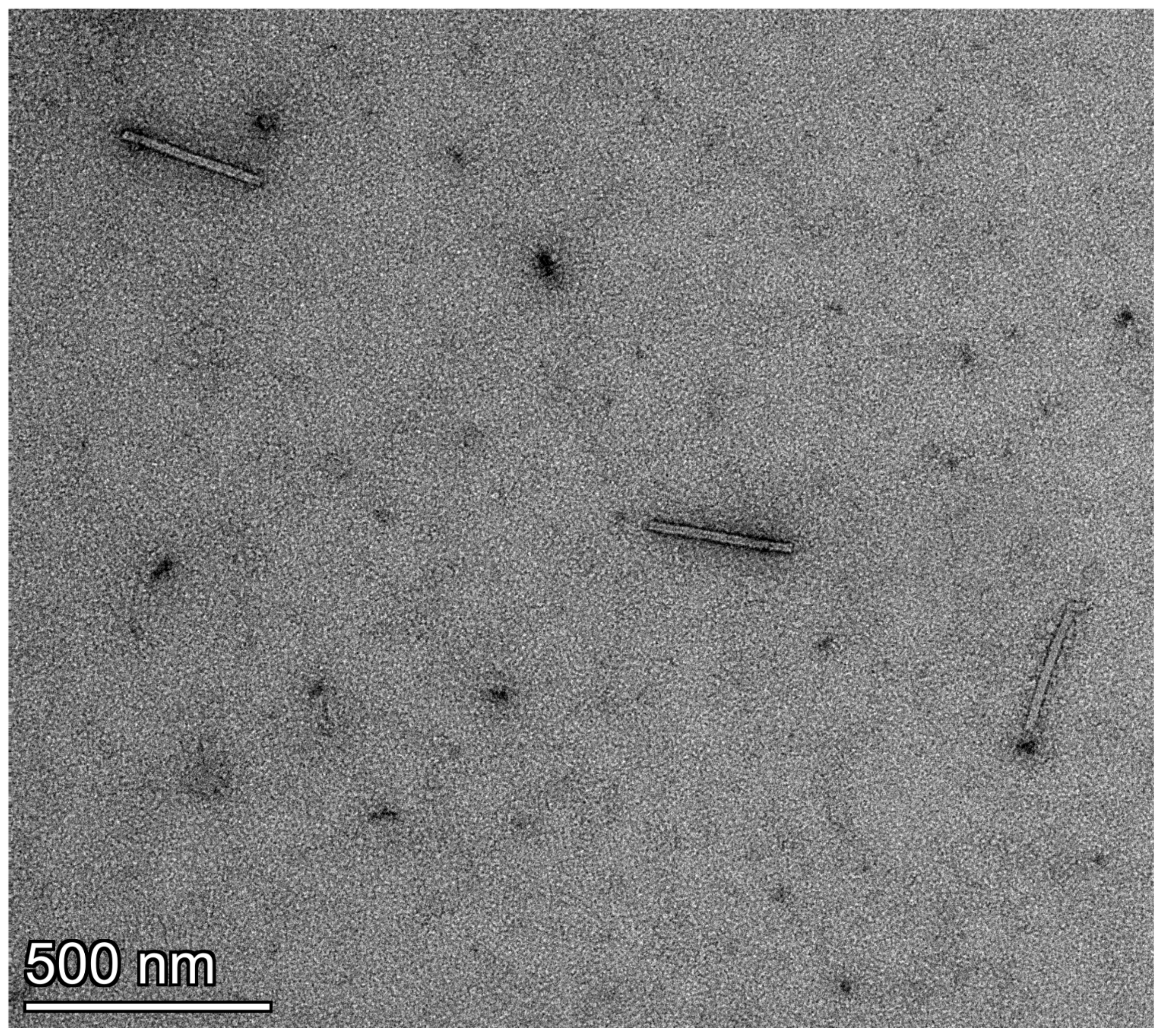


**d**

| Test plant / collection dpi / individual or pooled? | No. of quality-controlled (QC) reads | No. of reads mapping to PTV1 genome | Percentage of QC reads mapping to PTV1 genome |
|---|---|---|---|
| *Solanum lycopersicum* / 28 dpi / individual plant[a] | 157191 | 0 | 0.00% |
| *Solanum lycopersicum* / 28 dpi / individual plant[b] | 160438 | 0 | 0.00% |
| *Nicotiana clevelandii* / 21 dpi / pooled plants[c] | 159570 | 14216 | 8.91% |

Note: [a]lane 35 in (**b**). [b]lane 37 in (**b**). [c]lane 18 in (**a**).

**Supplementary Figure 5.** RT-PCR, TEM and nanopore sequencing results on the confirmation of infectivity of Plantago tobamovirus 1 (PTV1) on Solanaceae hosts. **a, b** RT-PCR assays for the detection of PTV1 in inoculated and young apical (systemic) leaves at different time points. **c** Transmission electron micrograph showing three virions of PTV1 from crude preparations of infected *Nicotiana clevelandii* tissues. **d** Number and proportion of long reads mapping to PTV1 genome, derived from selected inoculated plants.
